# Integrating computer-aided engineering and computer-aided design for DNA assemblies

**DOI:** 10.1101/2020.05.28.119701

**Authors:** Chao-Min Huang, Anjelica Kucinic, Joshua A. Johnson, Hai-Jun Su, Carlos E. Castro

**Affiliations:** Department of Mechanical and Aerospace Engineering, The Ohio State University, Columbus, Ohio 43210, USA; Department of Chemical and Biomolecular Engineering, The Ohio State University, Columbus, Ohio 43210, USA; Biophysics Graduate Program, The Ohio State University, Columbus, Ohio 43210, USA

## Abstract

Functional properties of modern engineering products result from merging the geometry and material properties of underlying components into sophisticated overall assemblies. The foundation of this design process is an integration of computer aided design (CAD) tools that allow rapid geometric modifications with robust simulation tools to guide design iterations (i.e. computer-aided engineering, CAE). Recently, DNA has been used to make nanodevices for a myriad of applications across fields including medicine, nanomanufacturing, synthetic biology, biosensing, and biophysics. However, currently these self-assembled DNA nanodevices rely primarily on geometric design, and hence, they have not demonstrated the same sophistication as real-life products. We present an iterative design pipeline for DNA assemblies that integrates CAE based on coarse-grained molecular dynamics with a versatile CAD approach that combines top-down automation with bottom-up control over geometry. This intuitive framework redefines the scope of structural complexity and enhances mechanical and dynamic design of DNA assemblies.

---

Combining computer aided design (CAD) with computer aided engineering (CAE)^1^ (i.e. iterative design guided by simulation) into Integrated Computational Materials Engineering (ICME) frameworks^2,3^ is essential to integrate consideration of material properties and geometric design across multiple length scales. ICME has been well studied for tailoring performance metrics of traditional engineering materials such as alloys and composites^4^. In contrast, integrating CAD and CAE for biomolecular self-assembly has remained elusive. Computationally-guided design of proteins is well-established^5^, but the diversity of structures and complexity of the interactions that govern self-assembly impede the development of geometric CAD. On the other hand, CAD tools that capture the structure and interactions of DNA have been essential to facilitating structural DNA nanotechnology^6–9^, but currently these approaches rely purely on geometric design. The recent emergence of high fidelity coarse-grained molecular dynamics (MD) simulation tools for DNA nanostructures^10–15^ provide an opportunity to realize CAE for DNA-based design to enable systems with new levels of structural complexity that can also be tailored for functional properties such as reconfiguration, mechanical properties, or stimulus response. Here we present an ICME approach for DNA assemblies that relies on a custom CAD tool with several features that enhance the scope of geometric design and facilitate tight integration with coarse-grained MD models^10–12^ to enable CAE for complex DNA assemblies.

The precise control over geometry of DNA assemblies^16–19^ make them highly attractive for applications such as drug delivery^20^, templating a variety of materials or molecules^21–25^, nanoscale measurement tools^26,27^, and molecular robotics^28–33^. However, DNA-based design approaches have largely overlooked material properties, which limits the structural, mechanical, and functional complexity. Currently, DNA assemblies are primarily designed using bottom-up approaches^6,7^ where strands are manually arranged and connected. In particular, caDNAno^6^ was transformative in simplifying the design process and broadening the use of DNA origami, but the largely manual routing is a slow process that is a challenge for non-experts and limits designs to small number of components with simple connectivity. To lower the barrier and speed up the design process, recent efforts have developed top-down approaches^8,9,34^ that take standard line models as inputs and utilize routing algorithms to partially or fully automate the design of wireframe structures. These tools are simple and fast, but, lack user interfaces to collect design parameters; hence, they are all limited to only static wireframe geometries. In summary, current bottom-up methods provide user control over geometry but limited complexity and relatively slow manual design, and top-down methods offer rapid and simple approaches to design complex shapes while sacrificing structural diversity and the ability to design features beyond a static shape (Supplementary Fig. 1). Hence, a rapid and versatile design approach is still needed to harness the potential of CAE for DNA assemblies.

**Figure 1.**
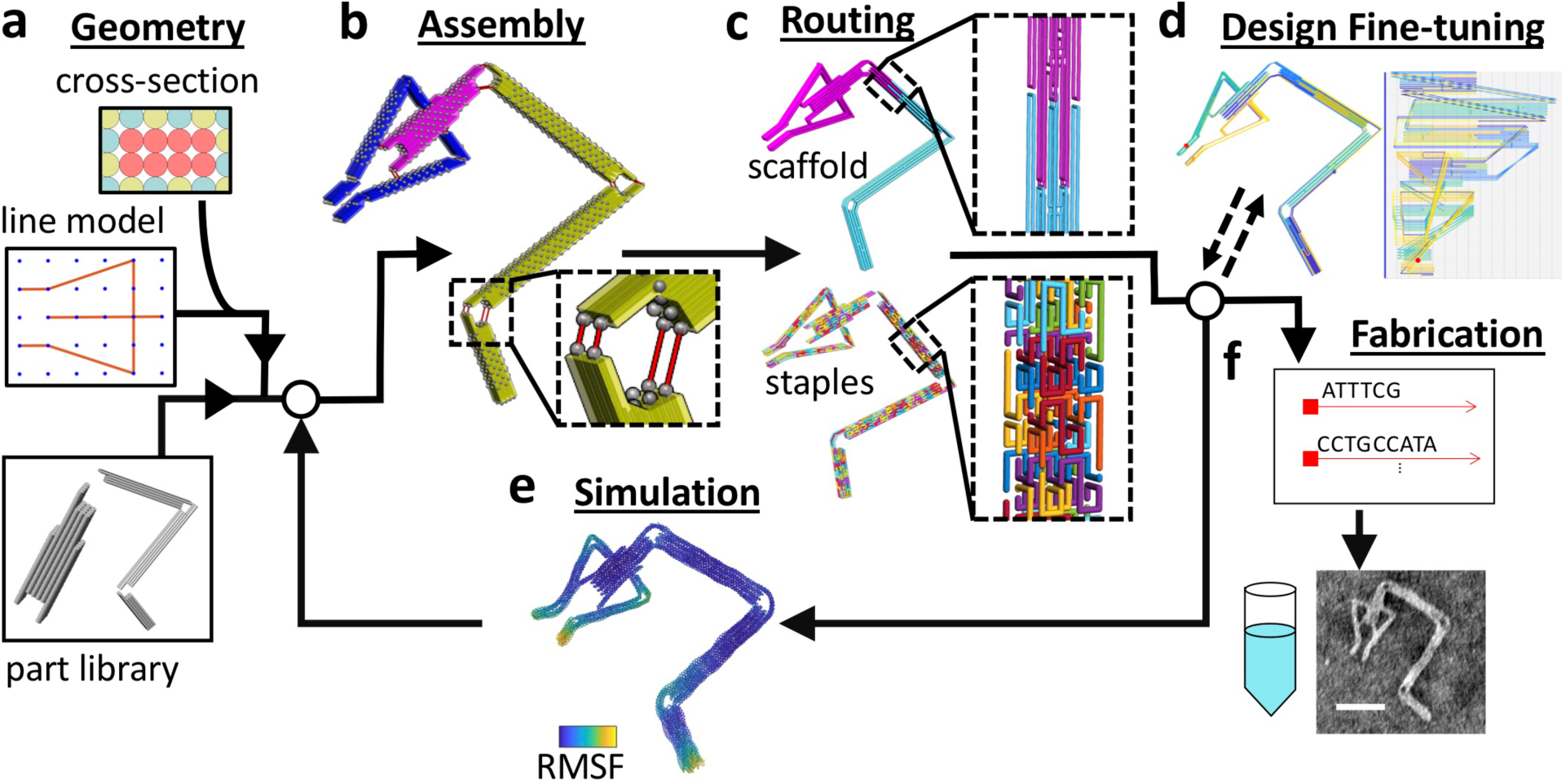
Schematic of the proposed design framework for multi-component DNA origami assemblies. **a**, To define the initial overall geometry users can take a top-down approach using a line model (.STEP file or MagicDNA sketch GUI) and specifying the length and cross-section of each line to create a full cylinder model of the assembly. Alternatively, individual or groups of components can be imported from a part library to build up assemblies. **b**, For assembly, each component can be subjected to translation or rotation to arrange the desired configuration. Users can specify connectivity between components manually or specify the number and type (e.g. end-to-end, end-to-side) and allow the program to automatically search for the closest sites (potential connection sites indicated by gray dots). **c**, Routing of scaffold and staple strands is automated including the capability to incorporate multiple scaffolds. **d**, Details of the strand routing can be visualized in a 3D structure and 2D diagram, and there is a two-way interface with the software caDNAno^6^ for fine modification of routing. **e**, Input files for simulation in oxDNA^10,43^ are automatically generated for virtual prototyping with built-in analysis including calculating the average shape and root-mean-squared fluctuations (RMSF). **f**, Once desired design metrics are achieved, the corresponding DNA sequences are automatically generated for fabrication and verification as shown by TEM. Scale bar = 50 nm.

Here we introduce a new hybrid design methodology for DNA assemblies that merges bottom-up and top down methods to provide a high level of structural diversity, expand the scope of complex design, and enable engineering of mechanical and dynamic properties. This hybrid framework accommodates design at multiple scales spanning the single nucleotide level to large and complex DNA assemblies. We implemented this approach through a GUI-based software called <u>M</u>ulti-component <u>A</u>ssembly in a <u>G</u>raphical <u>I</u>nterface guided by <u>C</u>omputation for <u>DNA</u> assemblies (MagicDNA) (Supplementary Fig. 2) that integrates simple user inputs, intuitive 3D visualization, and straightforward interfacing with molecular dynamics simulation tools for rapid CAE including assessment of properties like mobility and stiffness. We present 66 designs with simulation results and selected 14 structures with a range of complexity for experimental validation. Our results demonstrate that this framework simplifies and accelerates the design process, significantly expands the design domain for more applications, and enhances the robustness of DNA-based design.

**Figure 2.**
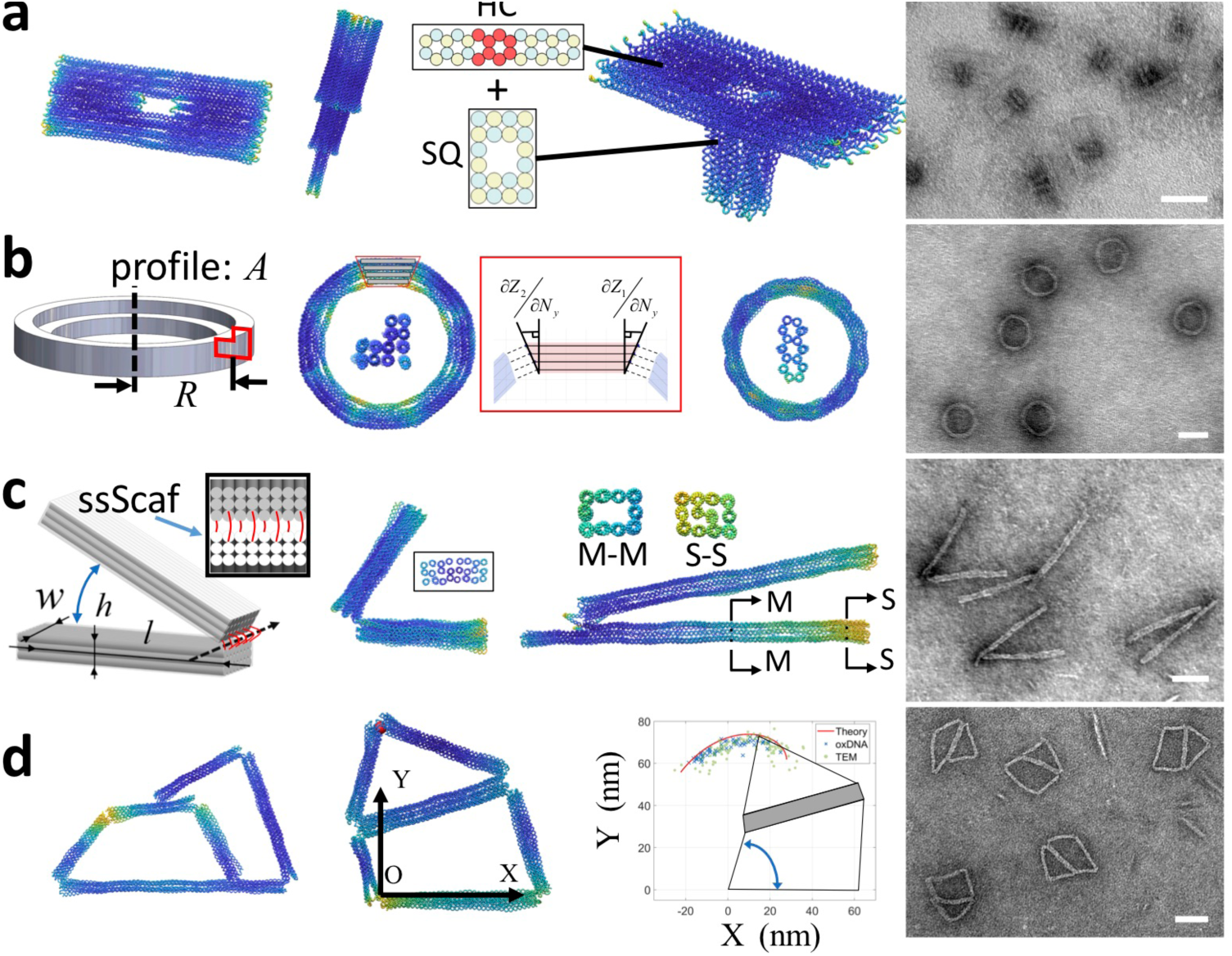
Parametric design of functional nano-devices. Structures depicted are the average oxDNA configuration with color-coded RMSF values. **a**, Horizontal, vertical, and 3D hybrid lattice nanopore structures are presented from left to right. **b**, The ring series are approximated by a polygon of straight bundles with gradients at the ends. **c**, The hinge devices are formed by two stiff arms joined by ssDNA scaffold connections to form a flexible rotational joint. **d**, The linkage designs implement multiple hinge joints to achieve a desired motion path. simulated motion closely matches the experimental data. TEM images illustrate well-folded structures with high yield. Scale bars = 50 nm.

## Iterative design process with simulation feedback

Modern CAD software supports geometric modeling for single component-design, and assembly-modeling for design of machinery with multiple components. Mimicking this widely accepted framework, we introduce a hybrid top-down and bottom-up design process for DNA assemblies based on scaffolded DNA origami^18,19^, where a long scaffold strand is folded into a compact structure by base-pairing with many shorter strands in a piecewise manner. Expanding on prior design approaches^6,8,9^, we introduce an intermediate component level for design, in addition to the nucleotide (bottom-up) and overall assembly (top-down) levels, into the framework where components are bundles of two or more dsDNA helices (Supplementary Fig. 3). Introducing this component level provides a convenient intermediate to design a wide range of static and dynamic assemblies with simple user inputs in an interactive 3D visualization environment. In MagicDNA, GUI tools and algorithms are implemented at the nucleotide, component, or assembly level, to enable seamless data exchange from lower to higher-levels (i.e. bottom-up) and communication from higher to lower levels (i.e. top-down). “Part” design is carried out among the nucleotide and component levels, and “assembly” modeling is carried out among the component and assembly levels.

**Figure 3.**
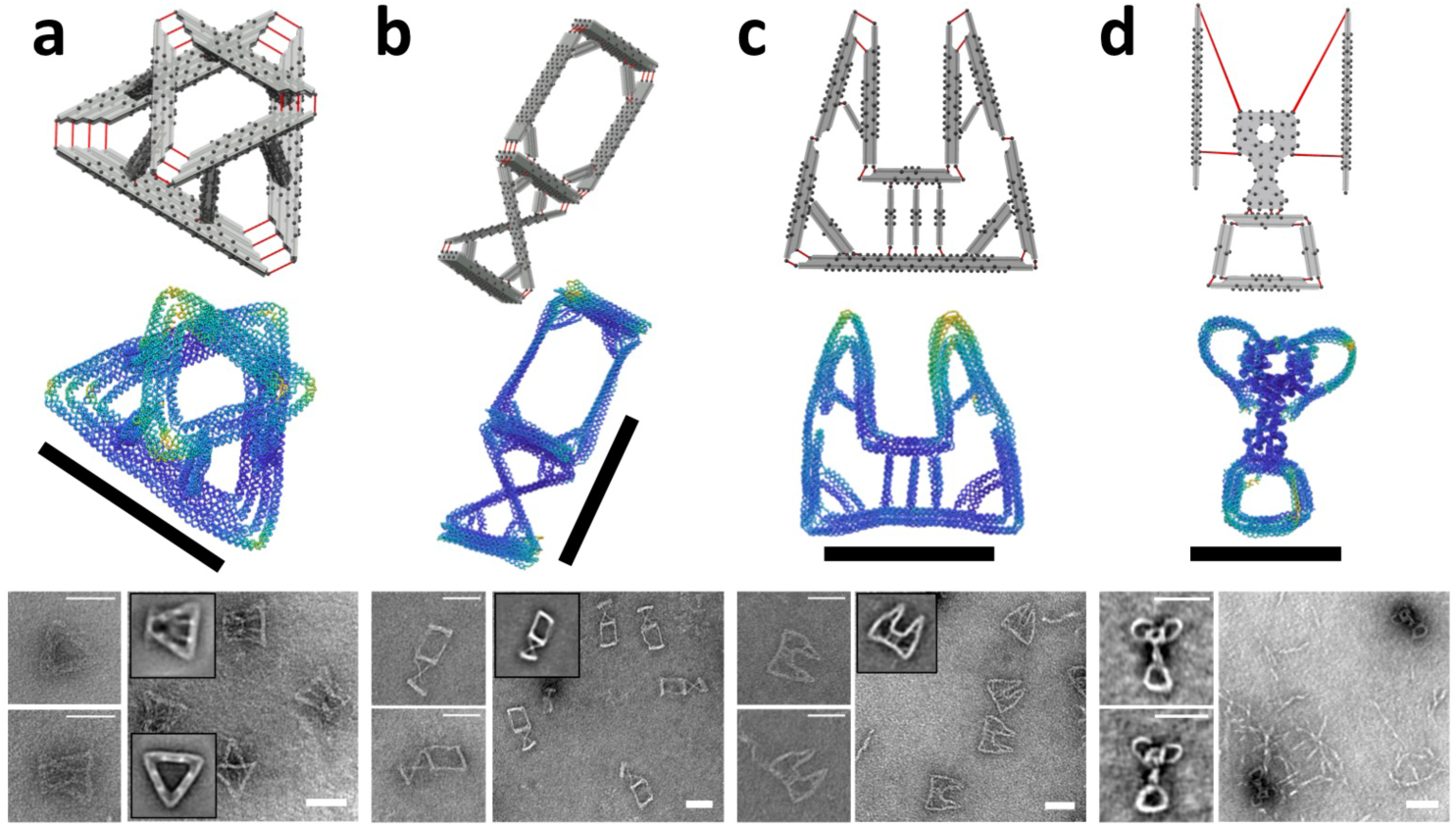
Design of multi-component complex structures. The first row shows assembly models from MagicDNA. Red lines represent joint connections, and black dots represent potential connection locations. The second row shows the average structure from oxDNA simulations with color-coded RMSF. The third row shows TEM images with inset image averages (except for trophy due to low yield, Supplementary Fig. 43). **a**, The Stewart platform consists of top and bottom triangular plates and six 2×2 square-lattice connecting limbs. **b**, The compound joint incorporates a compliant sliding joint on the top with two parallel blades and a compliant rotational joint at the bottom with cross-diagonal blades. Vertices on both joints are reinforced with struts. **c**, The gripper has a total of 15 square-lattice bundles, some using end gradients to create sharp corners. Seven struts were introduced to reinforce the overall shape. **d**, The trophy consists of a 62-helix bundle with a honeycomb lattice cross-section. The connections between the two single-layer “handles” and the central component were manually assigned to create the curved shapes. Scale bars = 50nm.

This collection of algorithms and GUI tools enables the systematic design workflow illustrated in Fig. 1. The first step of this workflow is inputting the overall assembly geometry (Fig. 1a). The top-down approach converts a 3D line model (imported from .STEP file or using built-in sketch interface) to components with user-defined cross-section and length. Alternatively, users can use a bottom-up approach by inserting components from a part library to build the assembly. In either case, users can modify various aspects of component geometries (Supplementary Figs. 5 and 6) to expand the design space, and components can be manipulated to arrange a desired assembly configuration (Supplementary Fig. 7). Connections between components (Fig. 1b) can be introduced either at the ends of components or on the surface where the scaffold is at an outward facing helical orientation. By specifying which components are connected to which (i.e. defining the connectivity matrix), connections can be automatically formed based on minimal distances between potential connections points (Supplementary Fig. 8). Alternatively, users can manually select locations for connections. In either case, each connection is formed by a double-scaffold crossover, and the length of these connections can be easily adjusted to tune the geometry or mechanical properties of the joint^35^.

With this approach it is straightforward to design assemblies with many interconnected components, making manually routing the DNA exceedingly challenging. To circumvent manual routing, recent top-down approaches developed automated routing algorithms^8,9,34,36^; however, those are limited to static wireframe structures with uniform components. To enable robust design of complex and dynamic assemblies with diverse geometries at the component level, we developed a general scaffold routing algorithm based on double-scaffold crossovers and a spanning tree algorithm. Details of the algorithm are provided in Supplementary Figs 9-12. Briefly, components are sub-divided into pairs of neighboring helices connected by external scaffold crossovers at their ends to form a scaffold cycle, and cycles corresponding to pairs of helices that connect across a joint are merged to reach a total number of *N* scaffold cycles. A spanning tree algorithm is used to find *N-*1 internal crossovers to reduce to a single cycle. We extended this algorithm to incorporate multiple scaffolds (Fig. 1c, discussed below), and we adopted a staple routing algorithm similar to prior work^6^ (details in Supplementary Figs 13-15) with added functionality for designing actuation or higher-order assembly. MagicDNA provides convenient GUIs for defining routing parameters and visualizing scaffold and staple routing in 3D (Fig. 1c and Supplementary Figs 16-19). We also added a feature to export a .JSON file for finer modification in caDNAno,^6^ which also enables the use of computational tools such as cando^37,38^ and COSM^13^ (Supplementary Fig. 20), which run caDNAno files, and modified routings can be uploaded back into MagicDNA.

To realize an ICME approach for DNA materials, we incorporated rapid virtual prototyping in MagicDNA with CAE simulation feedback (Supplementary Figs. 21 and 22) to fine-tune structural and functional properties while avoiding costs associated with multiple experimental fabrications. In particular, we incorporated tools to interface with the coarse-grained MD model oxDNA^10,11^, which is frequently used to predict the shape, mechanical properties, and motion of DNA nanostructures.^14,15,39,40^ We automated the generation of oxDNA simulation input files and integrated tools that calculate the average shape and fluctuations (Fig. 1e) or track key parameters (e.g. angles), which can guide design modifications. Once the desired design metrics are achieved, the design can be fabricated and verified experimentally (e.g. by TEM, Fig. 1f).^37,41,42^

### Top-down parametric design of functional devices

To demonstrate the versatility of our hybrid design process and the ability to rapidly adjust parameters building of the line model, we present several examples with the design workflow consisting of: 1) sketching the line model, 2) specifying component properties, 3) 3D manipulation of components to arrange a desired assembly configuration, 4) specifying ssDNA connections between components, and 5) running automated routing algorithms (Supplementary Movie 1). This process can be completed within ∼10 minutes, and then nucleotide (e.g. ssDNA connection length) or component (e.g. cross-section geometry) parameters can be adjusted within seconds. Specifically, we designed nanopores, ring structures, hinge devices, and 4-bar mechanisms, and selected one example of each for fabrication (Fig. 2). These are all design concepts that have demonstrated for the various applications such as detecting or probing biomolecules^27,44,45^, templating nanoparticles or other objects^46,47^, or as platforms for biomedical applications^48^. These examples also illustrate features of our CAD framework. For each case, we generated multiple designs for simulation (10^7^ steps in oxDNA) and chose one for experimental verification, but CAE iteration was not required for these relatively simple designs.

For the nanopore, we generated four designs (Supplementary Figs 23 and 24, three designs shown in Fig. 2a). The one chosen for fabrication (Fig. 2a, right) consists of a honeycomb lattice platform and a square lattice central pore, demonstrating distinct geometries for individual components rigidly connected in 3D. For the ring, we generated three designs (Supplementary Figs 25 and 26, two designs shown in Fig. 2b) starting with a polygonal line model, and we approximated the local curvature by incorporating gradients along the ends of the bundles (i.e. difference in length between layers of helices) to form angled vertex connections. Linear end gradients, as in the ring, can be input directly as a component property, and non-linear or discrete end gradients can be specified in a bottom-up manner by extruding helices with base-pair resolution. The ring illustrates the capability to design vertices and approximate curvature in 3D structures, both useful features in DNA-based design^49,50^.

A key goal of this framework is to simplify design of reconfigurable assemblies, since currently no automated tools address this class of emerging DNA systems. To demonstrate this capability, we designed three versions of a dynamic hinge (Supplementary Figs. 27 and 28, two designs shown in Fig. 2c). A GUI tool in MagicDNA allows visualization of the local 3D helical structure to assign joint connections at desired helical orientations and specify appropriate lengths to form an axis of rotation (Supplementary Fig. 27). We selected one hinge for fabrication that has a non-uniform cross-section (i.e. hollow in the middle) and exhibits flexible angular motion (Fig. 2c, right). Finally, we designed three mechanisms based on 4-bar linkages (Supplementary Figs. 29 and 30, two designs shown in Fig. 2d).We selected one for fabrication and used a longer oxDNA simulation (3×10^8^ steps) to track the motion of the top vertex. The simulations closely matched conformations measured by TEM (Fig. 2d, middle right), demonstrating the ability to design functional properties beyond shape, such as mobility.

### Top-down iterative design for complex structures

The ability to create complex multi-component designs enabled by our ICME framework, GUI-based CAD tool, algorithms, and interfacing with coarse-grained model, makes CAE simulation feedback essential to design verification and improvement. To reduce the number of iterations for complex designs, we take a modular approach by first optimizing sub-systems consisting of a few components. This reduces simulation time and allows more extensive study on how design parameters affect design metrics (e.g. shape, stiffness, configuration etc.). For example, a vertex joint can be flexible or rigid, or have sharp or rounded corners (Supplementary Fig. 31). This initial simulation is to identify of structural instability or excessive deformation, which can occur with small-cross sections or short lengths (Supplementary Fig. 32). Additionally, simulation feedback can be used to create curved features by connecting components with mismatched length and stiffness (Supplementary Fig. 33). While a modular approach is efficient, one can still iterate sub-component designs within a larger assembly (e.g. birthday cake structure in Supplementary Fig. 34).

We term this approach “hierarchical design”, where users can fine-tune component or sub-assembly geometry by adjusting design parameters based on simulation feedback, and then add those components into a larger-scale assembly. We applied this hierarchical approach to design four complex structures: a Stewart platform, a compound compliant joint, a gripper, and a trophy, all inspired by macroscopic counterparts. Due to the complex features (many components, 3D geometry and connectivity, vertices, curvature, hybrid lattice, etc.), these structures would not be practical to design with prior software tools. The top-down approach is still convenient for assigning approximate geometric parameters to a large number of components in the first iteration. Then component and nucleotide level parameters are specified in MagicDNA to complete a design. In all these designs we used information from sub-assembly simulations (Supplementary Figs 31-34) to guide the designs of specific features. The simulation trajectory of the full design was also used to guide structural adjustment such as editing bundle geometry or adding trusses to enhance mechanical properties of vertices (Fig. 3 and Supplementary Figs 35-43). The platform, compound joint, gripper, and trophy, required 2, 4, 13, and 5 iterations, respectively, with the final designs illustrated in Fig. 3.

### Bottom-up and hierarchical design of reconfigurable assemblies

Functional materials are often comprised of many similar structural units, as in structural metamaterials^51^, which is well-suited to the hierarchical approach. Here, we demonstrate reconfigurable assemblies comprised of multiple similar units, specifically a deployable mechanism (Fig. 4a, serial tetrahedron) and a rotational mechanism (Fig. 4b, the butterfly). We generated the initial structural unit design using the top-down approach and optimized the unit design with simulation feedback. Then, we used bottom-up assembly to import and integrate these basic units into a desired pattern.

**Figure 4.**
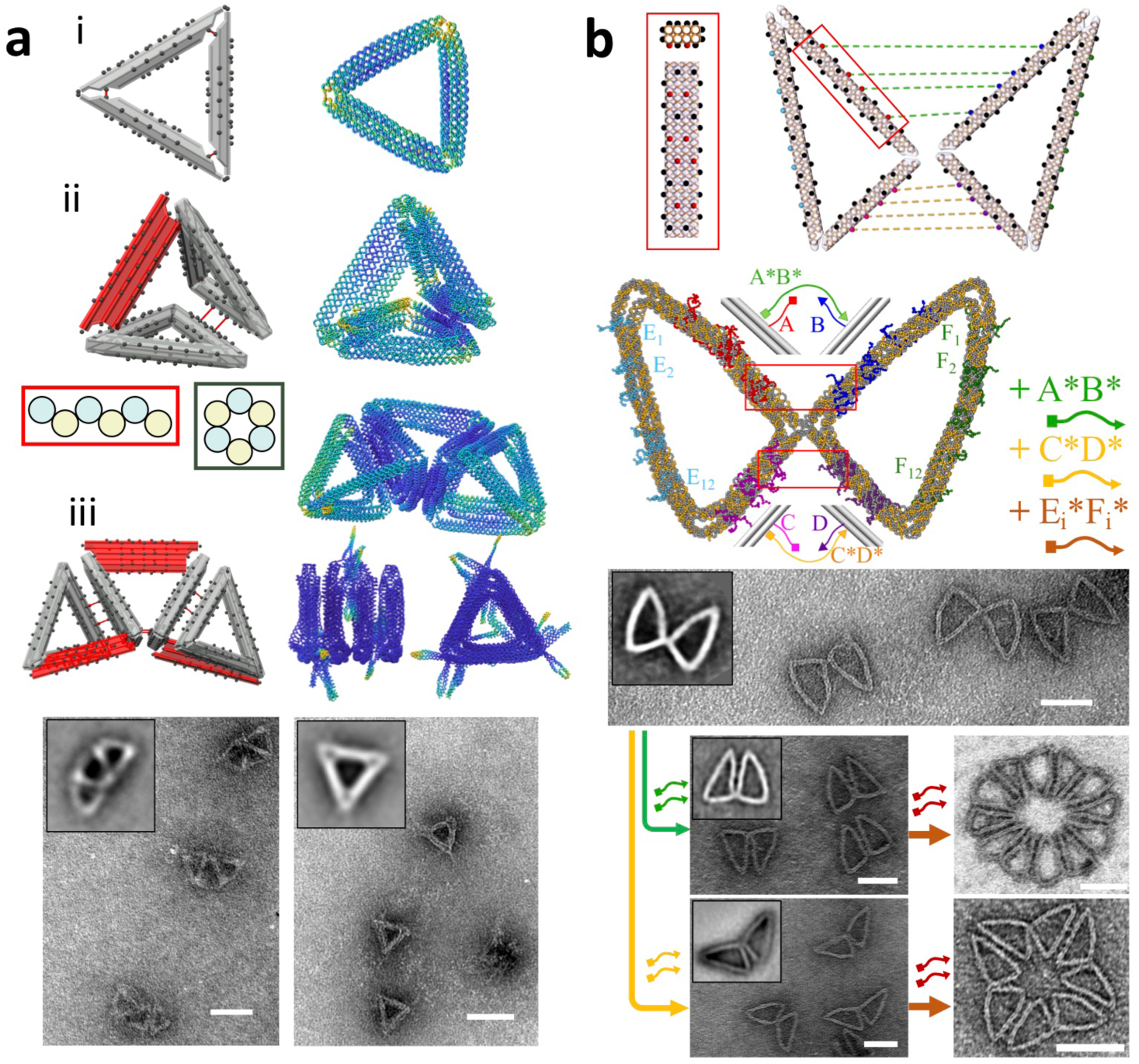
Reconfigurable devices by hierarchical design. **a**, A deployable mechanism formed by a serial chain of three tetrahedrons. From top to bottom: i) a triangular plate was validated with oxDNA and ii) duplicated to form a tetrahedron with an extra blade component (red). iii) The verified tetrahedron was duplicated to form a serial chain with deployed and compact configurations as validated with TEM images. Insets show image averages **b**, The butterfly mechanism is made of two triangles connected by a hinge joint. There are overhangs on the upper, lower, and outer edges to actuate into different configurations and then polymerized into distinct circular assemblies. Scale bars = 50nm.

**Figure 5.**
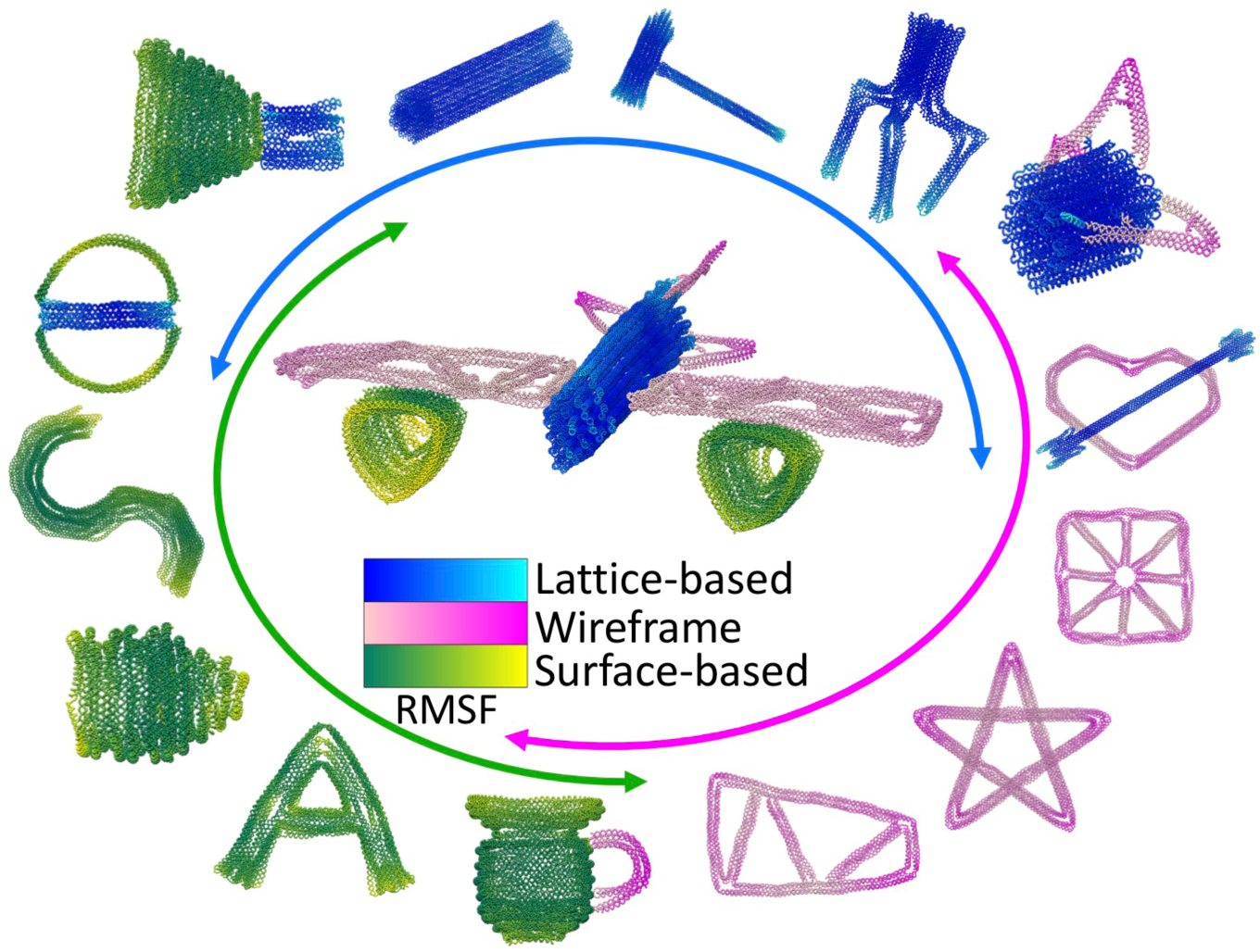
Broadening the design spectrum by integrating wireframe, lattice, and surface-based components. The three classes of geometric modeling are supported by various built-in GUIs in MagicDNA. The stiffer lattice-based components (relative RMSF shown blue to cyan) are made by assigning an initial geometry to lines (top-down) and extruding helices in the bundle editor GUI (bottom-up). Surface-based (or shell) components (relative RMSF shown green to yellow) are designed using multiple segments with end gradients to approximate features with curvature in multiple directions. The wireframe models (RMSF shown pink to magenta) are similar to lattice but with small cross-section (e.g. 2×2) and many can be easily connected in space. The airplane (∼33 kbps) exploits all three types (wireframe wings, surface-based turbine, and lattice-based fuselage).

**Figure 6.**
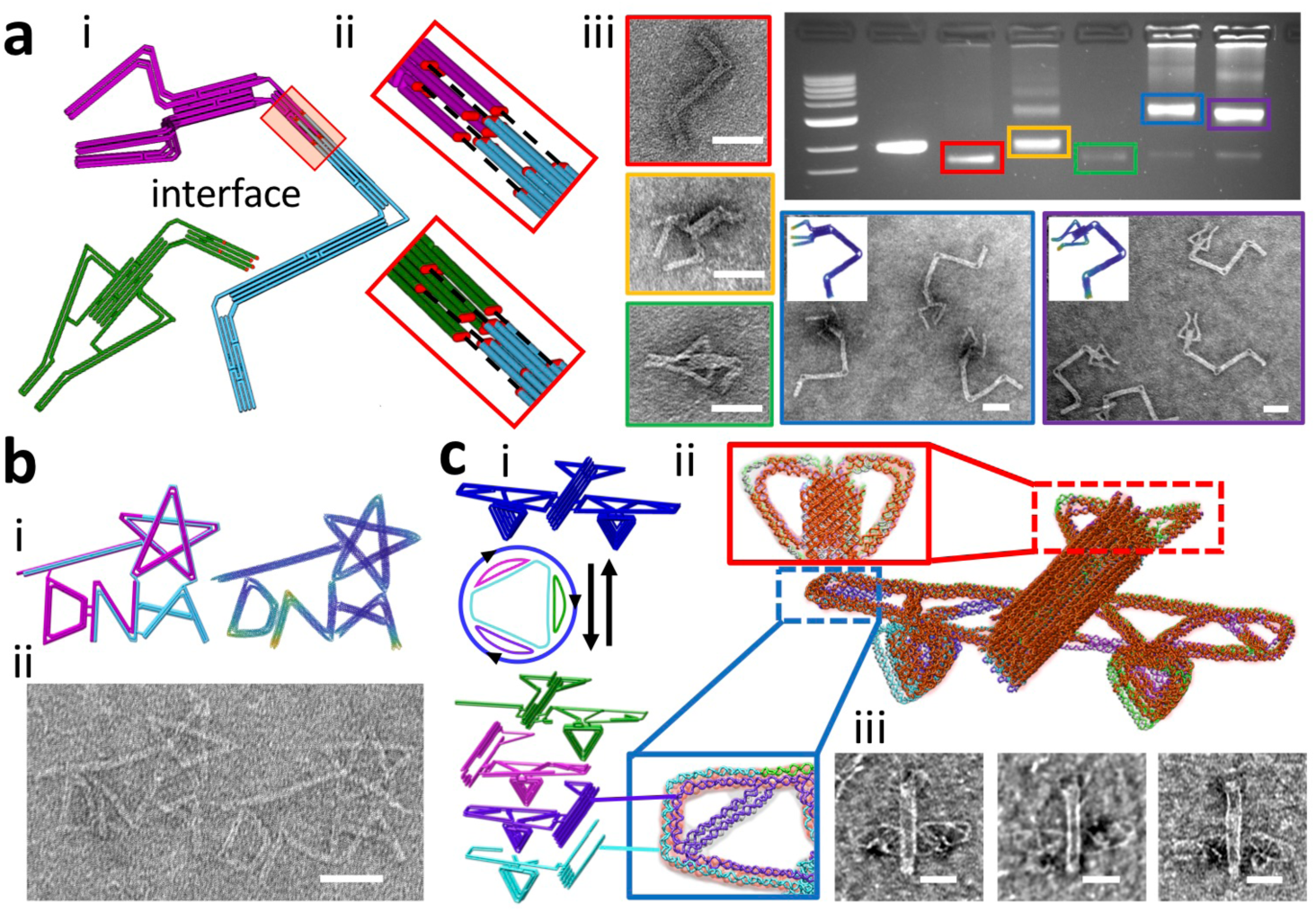
Multi-scaffold designs. **a**, One of two multi-scaffold methods to achieve devices with interchangeable parts by specifying the interface. (i) The two-scaffold robot arm design has an exchangeable End-Of-Effector, claw (magenta) or tweezer (green), and (ii) shows a zoom-in of the user-defined interface specified by adding crossovers across the interface and then forming a single scaffold cycle on either side of the interface using a spanning forest algorithm with the interface defined as the separation between trees. (iii) Individual and combined structures were folded, each in a single-pot reaction, and validated by gel electrophoresis and TEM. **b**, (i) The second multi-scaffold approach applies *K*-1 crossovers to split a single cycle scaffold into *K* cycles, as demonstrated here for the MagicDNA logo where *K*=2. (ii) TEM image of the logo. **c**, (i) This approach was also used for the airplane to add three crossovers to split the original scaffold into four cycles (*K*=4) of desired lengths. (ii) The airplane design comprises 4 orthogonal scaffolds with a total of 682 staples. 462 of these staples connect at least two scaffolds (depicted as red staples, and the staples that bind to a single scaffold are grey and transparent), showing most of the structural components are populated by staples that connect at least two scaffolds. (iii) TEM images of the airplane. Scale bars = 50nm.

To illustrate the ICME process within a hierarchical assembly to scale the dimensions and function, the deployable tetrahedron was first designed by sketching three lines and converting them into bundles with end gradients to form a basic triangular shape, confirmed with oxDNA (Fig. 4a, i and Supplementary Fig. 44). The verified triangle unit was duplicated into two instances, which were arranged to form two sides of a tetrahedron (Fig. 4a, ii). A blade component, which controls the open or closed configuration, was added to complete the tetrahedron. Our normal scaffold algorithm does not allow connections to multiple components on a single node, since in double-scaffold crossovers both strands must connect between the same two components. Since multi-way junctions are required for the tetrahedron design, we fine-tuned the routing in caDNAno (Supplementary Fig. 44) and uploaded to MagicDNA. The blade can be folded into a straight or contracted configuration to convert the structure into the deployed or compact states (Supplementary Fig. 44). Both open and closed configurations of the single tetrahedron were validated by oxDNA (Fig. 4a and Supplementary Fig. 44). The verified tetrahedron was duplicated into three instances while removing one of the triangular plates on the two outer tetrahedrons. The final designs in both deployed and compact configurations were verified by simulation and experimental fabrication (Fig. 4a and Supplementary Figs 46 and 47). We also demonstrated a deployable umbrella mechanism following this hierarchical design (Supplementary Figs 48-50).

Another common actuation strategy for dynamic devices is to add strands that connect multiple overhangs to latch two components together,^32,33^ which we demonstrate here for a butterfly mechanism (Figs. 4b and Supplementary Figs 51-54). We created an overhang design GUI in MagicDNA where users input parameters (e.g. length) and specify locations and connections directly on the 3D model. The staple routing algorithm then satisfies these inputs and optimizes overhang sequences to minimize complementarity to the scaffold.^46^ We used the bottom-up duplication of two triangles for the butterfly design and fine-tuned the scaffold routing at the joint with caDNAno (Supplementary Fig. 51). Next, we used the overhang design GUI to specify 28 pairs of overhangs (Supplementary Fig. 4b, top), including 8 pairs with an identical sequence to close the wings along the upper edges, 8 pairs with a second identical sequence to close the wings along the lower edges, and 12 pairs all with unique sequences to assemble multiple butterflies together along the outer edges. By adding complementary strands to the overhangs in a specific order, we show different actuation/assembly pathways result in two distinct high-order assemblies (Fig. 4b).

### Expanding the design domain of complex DNA assemblies

So far, we have shown the versatility (varying cross-section, end gradients, hybrid lattices, mobility, etc.) of our hybrid top-down and bottom-up design framework implemented in MagicDNA (Supplementary Fig. 55). Here we generalize to an even wider spectrum of design by integrating wireframe, surface-based, and lattice-based models (Figs. 5 and Supplementary Figs 56-68) into complex assemblies. These types of structures have been demonstrated individually^17^; however, no current CAD tools integrate these classes of structures into a single assembly.

We demonstrated this capability to integrate lattice, surface, and wireframe components into a DNA airplane assembly design (Fig. 5 center). We started with a top-down line model for the whole system to establish approximate sizes for the six sub-systems (Supplementary Fig. 69). Then, the sub-systems were individually optimized and then combined into a single assembly consisting of a lattice-based main body, wireframe wings and tail, and surface-based turbines. The airplane design totals 28 bundle components, ∼33 kbps, and more than 800 staples, showing the design framework, algorithms, and software essentially have no limit for scale. Simulation times increase, but continued efforts in coarse-grained molecular dynamics^12^ are addressing this challenge (Supplementary Fig. 22).

### Multi-scaffold and modular designs

Increasing design complexity and size generally requires more material. There have been multiple recent advances in making higher order DNA assemblies that integrate multiple structures^52,53^ to make well-controlled micron-scale assemblies; however, these are usually carried out through multiple reaction steps and have only demonstrated assembly of similar and relatively simple structures. To improve yields and enhance complexity while also simplifying the design process, we sought to enable design of large assemblies with many unique components that could be fabricated in a single-pot reaction. Prior work demonstrated single-pot folding of large DNA structures using up to ∼50 kb scaffolds^54^, integrating multiple orthogonal sequence scaffolds^42,50^, or using exclusively brick strands^55,56^(similar to staple strands). We chose to focus on using orthogonal scaffolds based on a recent breakthrough demonstrated for simpler assemblies^42^, which was demonstrated using relatively simple geometries. But this approach enables assemblies with many distinct structural components, asymmetric and fully addressable shape, and programmable mechanical and dynamic properties. Also, the use of multiple scaffolds allows for modular design where single scaffold sub-assemblies can be re-used in multiple higher order assemblies.

We implemented two approaches based on a spanning forest algorithm for multi-scaffold design in MagicDNA. The first is intended for modular design, allowing users to add well-defined interfaces between structures by: 1) adding internal crossovers to form a desired interface, and 2) ignoring potential scaffold crossovers in this region during automated scaffold routing to ensure formation of separate cycles (additional details in Supplementary Figs 9 and 70-72). This approach is demonstrated by the robotic manipulator with an exchangeable end effector (Fig. 6a, Supplementary Figs 73-78). The arm comprises three components connected via compliant joints^35^, and at one end, the arm is connected to either a claw-like end effector or a tweezer-like end effector with the two structures interlocked at the interface, which was shown to improve yield ^42^. Our gel and TEM results demonstrate quality folding and high yield of this robotic manipulator with exchangeable end effectors as an example of modular robots. Although it is not the focus here, this approach with defined bundle connectivity and interfaces also allows for direct design and simulation of multiple structures bound together intended for hierarchical multi-pot assembly (Supplementary Figs. 79-80)^53,57^.

In the second approach for multi-scaffold designs, which is intended for complex asymmetric shapes, we split the full single scaffold cycle into *K* cycles by applying *K-*1 crossovers (Supplementary Fig. 81). The algorithm searches a list of internal scaffold crossovers in the initial single scaffold until it finds crossovers that break the scaffold into *K* cycles of the desired lengths. To facilitate finding solutions, we include some tolerance for the cycle lengths to be somewhat shorter than the full length (default = 10%). We used this approach to design a wireframe MagicDNA logo using a bottom-up process to assemble a wand with a stick and star (Supplementary Fig. 58) and the “DNA” script (Supplementary Fig. 65). The MagicDNA logo was folded with an M13-derived p8064^19^ scaffold and a CS4-7559 scaffold^42^, and verified with TEM (Figs. 6b, and Supplementary Figs 82 and 83). In the case of the airplane (Fig. 6c) before splitting, there are about ∼30 kb and roughly two thousand possible crossovers to split the scaffold. We used a custom and stochastic search algorithm to identify three (*K*-1) crossovers splitting the *K*=4 scaffolds with the desired scaffold lengths (M13-derived p8064 and CS3_L_7560, CS4_7557, CS5_7559^42^). To ensure stable attachment between the four scaffolds during folding, we developed a heuristic optimization with the objective of maximizing the number of staples that connect at least two scaffolds (Supplementary Fig. 84). Detailed design data showing connectivity between the scaffolds and staples are visualized in GUIs (Supplementary Figs. 85 and 86 and Movie 2). The resulting four scaffold routings for the airplane are shown in Fig. 6c where 69% of staples connect at least two scaffolds (depicted as red staples in Fig. 6c, top right model).

## Discussion

We demonstrated a versatile framework that combines the benefits of top-down, bottom-up, and hierarchical design implemented in the MATLAB-based software MagicDNA, which offers: 1) scaffold and staple routing algorithms to automate design operations that are tedious, time-consuming and error-prone; 2) built-in GUIs for intuitive editing of 3D geometric models of complex geometries at nucleotide, component, and assembly levels; and 3) interfacing with coarse-grained simulation for rapid CAE including consideration of properties beyond shape. Compared with bottom-up design tools^6,7^, MagicDNA has routing algorithms and component and assembly level manipulations to allow for rapid construction of large many-component designs directly from 3D models with simple user inputs (Supplementary Fig. 55). Compared with top-down tools^8,9,34,36^, our framework significantly enhances user control over geometric, mechanical and dynamic properties of assemblies, and enables actuation, higher order assembly, and multi-scaffold capabilities (Supplementary Fig. 55). These advances significantly extend the current-state-of-the-art for design of DNA assemblies.

Our design framework and the MagicDNA tool evolved from a heuristic design process for dynamic DNA origami mechanisms (DOMs)^29,32,33,40^ as well as recently emerging techniques such as hierarchical multi-structure assembly^52,53^, multi-scaffold assembly^42,50^, and recent advances in simulation of DNA nanodevices^10–12,14^. A key foundation for our design process is the ability to control stiffness of the DNA material by leveraging highly flexible ssDNA, semi-flexible double-stranded helices, and stiff compact bundles to match the desired local properties and overall function. This is particularly useful for prescribing the mobility and reconfiguration of assemblies. Furthermore, our robust algorithms and intuitive 3D visualization facilitate designs with many components connected in 3D. Finally, the CAE component is essential for this new frontier of complex design. Hence, this integrated CAD/CAE framework can accelerate the development of next generation molecular robots.

In addition, the recent emergence of controlled templating of gold, silver, or silica on DNA assemblies^21,23,46^, and the use of DNA “masks”^22^ for lithography provide avenues to exploit the new levels of size and geometric complexity demonstrated here, especially the design of large multi-scaffold structures and higher order multi-structure assemblies. Furthermore, we took an initial step to integrate functional considerations beyond shape into the design process based on CAE, and with the continued development of simulation tools we anticipate a powerful capability to rapidly design devices for targeted functions. Finally, this new regime of fabrication opens new fundamental questions about folding pathways, kinetics, and thermodynamics for these complex (many-component, multi-scaffold, hybrid lattice, etc.) assemblies.

## Supporting information

MagicDNA software manual

Supplemental Table - DNA staple sequences

## Acknowledgments

This work was supported by National Science Foundation grants 1536862 to H.-J.S and C.E.C. and grant 1921955 to C.E.C. We acknowledge Floris Engelhardt and Hendrik Dietz for providing custom scaffolds, Tural Aksel and Shawn Douglas for sharing the caDNAno toolkit, Christopher Maffeo and Aleksei Aksimentiev for supporting interface to MrDNA, Tara MacCulloch and Nickolas Stephanopoulos for providing K-10 peptide. We thank the Campus Microscopy and Imaging Facility (CMIF) of The Ohio State University for imaging support. We also thank Wolfgang Pfeifer and Christopher Maffeo for critiques on the manuscript and supplement material.

## Author contributions

C.-M.H. developed the software and the algorithm, designed and simulated all the structures, analyzed the data, prepared the tutorial, and supported experiments. A.K. conducted the majority of experiments and analyzed the experimental results. J.A.J was an initial user and provided critical early feedback on software features, interface, and instructions. H.-J.S. supervised the development of the software and interpreted the data. C.E.C supervised the experimental validation and the entire study and interpreted the data. C.-M.H, A.K., H.-J.S, and C.E.C. wrote the manuscript. All authors commented on and edited the manuscript.

## Data and materials availability

All data presented in the main text and the supplementary materials are available upon request. The software package is available from the link provided in the supplementary materials. The design files of the structures that were fabricated for experimental validation are included in the software package.

## Supplementary Materials

Materials and Methods

Supplementary Text

Supplementary Figures 1-87

Supplementary Table 1

Supplementary Movies 1 and 2

The pdf file for the software user manual

The excel sheets for the staple list of the 14 structures for fabrication

Supplementary References 1 to 30

## References

1. Chang, K.-H. e-Design: Computer-Aided Engineering Design. (Academic Press, 2016).

2. Olson, G. B. Computational Design of Hierarchically Structured Materials. Science 277, 1237–1242 (1997).

3. Panchal, J. H., Kalidindi, S. R. & McDowell, D. L. Key computational modeling issues in Integrated Computational Materials Engineering. Computer-Aided Design 45, 4–25 (2013).

4. Backman, D. G. et al. ICME at GE: Accelerating the insertion of new materials and processes. JOM 58, 36–41 (2006).

5. Huang, P.-S., Boyken, S. E. & Baker, D. The coming of age of de novo protein design. Nature 537, 320–327 (2016).

6. Douglas, S. M. et al. Rapid prototyping of 3D DNA-origami shapes with caDNAno. Nucleic Acids Res 37, 5001–5006 (2009).

7. Williams, S. et al. Tiamat: A Three-Dimensional Editing Tool for Complex DNA Structures. in DNA Computing 90–101 (Springer, Berlin, Heidelberg, 2008). doi: 10.1007/978-3-642-03076-5_8.

8. Benson, E. et al. DNA rendering of polyhedral meshes at the nanoscale. Nature 523, 441–444 (2015).

9. Veneziano, R. et al. Designer nanoscale DNA assemblies programmed from the top down. Science 352, 1534–1534 (2016).

10. Doye, J. P. K. et al. Coarse-graining DNA for simulations of DNA nanotechnology. Phys. Chem. Chem. Phys. 15, 20395–20414 (2013).

11. Snodin, B. E. K. et al. Introducing improved structural properties and salt dependence into a coarse-grained model of DNA. J. Chem. Phys. 142, 234901 (2015).

12. Maffeo, C. & Aksimentiev, A. MrDNA: a multi-resolution model for predicting the structure and dynamics of DNA systems. Nucleic Acids Res doi: 10.1093/nar/gkaa200.

13. Reshetnikov, R. V. et al. A coarse-grained model for DNA origami. Nucleic Acids Res 46, 1102–1112 (2018).

14. Sharma, R., Schreck, J. S., Romano, F., Louis, A. A. & Doye, J. P. K. Characterizing the Motion of Jointed DNA Nanostructures Using a Coarse-Grained Model. ACS Nano (2017) doi: 10.1021/acsnano.7b06470.

15. Shi, Z., Castro, C. E. & Arya, G. Conformational Dynamics of Mechanically Compliant DNA Nanostructures from Coarse-Grained Molecular Dynamics Simulations. ACS Nano 11, 4617–4630 (2017).

16. Seeman, N. C. Nucleic acid junctions and lattices. Journal of Theoretical Biology 99, 237–247 (1982).

17. Seeman, N. C. & Sleiman, H. F. DNA nanotechnology. Nat Rev Mater 3, 1–23 (2017).

18. Rothemund, P. W. K. Rothemund, P. W. (2006). Folding DNA to create nanoscale shapes and patterns. Nature, 440(7082), 297–302. Nature 440, 297–302 (2006).

19. Douglas, S. M. et al. Self-assembly of DNA into nanoscale three-dimensional shapes. Nature 459, 414–418 (2009).

20. Jiang, Q. et al. DNA Origami as a Carrier for Circumvention of Drug Resistance. J. Am. Chem. Soc. 134, 13396–13403 (2012).

21. Sun, W. et al. Casting inorganic structures with DNA molds. Science 346, 1258361 (2014).

22. Shen, B. et al. Plasmonic nanostructures through DNA-assisted lithography. Science Advances 4, eaap8978 (2018).

23. Liu, X. et al. Complex silica composite nanomaterials templated with DNA origami. Nature 559, 593–598 (2018).

24. Bayrak, T. et al. DNA-Mold Templated Assembly of Conductive Gold Nanowires. Nano Lett. 18, 2116–2123 (2018).

25. Johnson, J. A. et al. The path towards functional nanoparticle-DNA origami composites. Materials Science and Engineering: R: Reports 138, 153–209 (2019).

26. Shaw, A. et al. Spatial control of membrane receptor function using ligand nanocalipers. Nature Methods 11, 841–846 (2014).

27. Le, J. V. et al. Probing Nucleosome Stability with a DNA Origami Nanocaliper. ACS Nano 10, 7073–7084 (2016).

28. Lauback, S. et al. Real-time magnetic actuation of DNA nanodevices via modular integration with stiff micro-levers. Nature Communications 9, 1446 (2018).

29. Gerling, T., Wagenbauer, K. F., Neuner, A. M. & Dietz, H. Dynamic DNA devices and assemblies formed by shape-complementary, non–base pairing 3D components. Science 347, 1446–1452 (2015).

30. Thubagere, A. J. et al. A cargo-sorting DNA robot. Science 357, eaan6558 (2017).

31. Douglas, S. M., Bachelet, I. & Church, G. M. A Logic-Gated Nanorobot for Targeted Transport of Molecular Payloads. Science 335, 831–834 (2012).

32. Marras, A. E., Zhou, L., Su, H.-J. & Castro, C. E. Programmable motion of DNA origami mechanisms. PNAS 112, 713–718 (2015).

33. Zhou, L., Marras, A. E., Huang, C.-M., Castro, C. E. & Su, H.-J. Paper Origami-Inspired Design and Actuation of DNA Nanomachines with Complex Motions. Small 0, 1802580 (2018).

34. Jun, H. et al. Automated Sequence Design of 3D Polyhedral Wireframe DNA Origami with Honeycomb Edges. ACS Nano (2019) doi: 10.1021/acsnano.8b08671.

35. Zhou, L., Marras, A. E., Su, H.-J. & Castro, C. E. DNA Origami Compliant Nanostructures with Tunable Mechanical Properties. ACS Nano 8, 27–34 (2014).

36. Matthies, M., Agarwal, N. P. & Schmidt, T. L. Design and Synthesis of Triangulated DNA Origami Trusses. Nano Lett. 16, 2108–2113 (2016).

37. Castro, C. E. et al. A primer to scaffolded DNA origami. Nat Meth 8, 221–229 (2011).

38. Kim, D.-N., Kilchherr, F., Dietz, H. & Bathe, M. Quantitative prediction of 3D solution shape and flexibility of nucleic acid nanostructures. Nucleic Acids Res 40, 2862–2868 (2012).

39. Snodin, B. E. K., Schreck, J. S., Romano, F., Louis, A. A. & Doye, J. P. K. Coarse-grained modelling of the structural properties of DNA origami. Nucleic Acids Res 47, 1585–1597 (2019).

40. Huang, C.-M., Kucinic, A., Le, J. V., Castro, C. E. & Su, H.-J. Uncertainty quantification of a DNA origami mechanism using a coarse-grained model and kinematic variance analysis. Nanoscale 11, 1647–1660 (2019).

41. Wagenbauer, K. F. et al. How We Make DNA Origami. ChemBioChem 18, 1873–1885 (2017).

42. Engelhardt, F. A. S. et al. Custom-Size, Functional, and Durable DNA Origami with Design-Specific Scaffolds. ACS Nano 13, 5015–5027 (2019).

43. Ouldridge, T. E., Louis, A. A. & Doye, J. P. K. Structural, mechanical, and thermodynamic properties of a coarse-grained DNA model. The Journal of Chemical Physics 134, 085101 (2011).

44. Wei, R., Martin, T. G., Rant, U. & Dietz, H. DNA Origami Gatekeepers for Solid-State Nanopores. Angewandte Chemie International Edition 51, 4864–4867 (2012).

45. Xu, W. et al. A Programmable DNA Origami Platform to Organize SNAREs for Membrane Fusion. J. Am. Chem. Soc. 138, 4439–4447 (2016).

46. Johnson, J. A., Dehankar, A., Winter, J. O. & Castro, C. E. Reciprocal Control of Hierarchical DNA Origami-Nanoparticle Assemblies. Nano Lett. (2019) doi: 10.1021/acs.nanolett.9b02786.

47. Kuzyk, A. et al. Reconfigurable 3D plasmonic metamolecules. Nat Mater 13, 862–866 (2014).

48. Ponnuswamy, N. et al. Oligolysine-based coating protects DNA nanostructures from low-salt denaturation and nuclease degradation. Nature Communications 8, 15654 (2017).

49. Dietz, H., Douglas, S. M. & Shih, W. M. Folding DNA into Twisted and Curved Nanoscale Shapes. Science 325, 725–730 (2009).

50. Zhang, F. et al. Complex wireframe DNA origami nanostructures with multi-arm junction vertices. Nature Nanotechnology 10, 779–784 (2015).

51. Bertoldi, K., Vitelli, V., Christensen, J. & van Hecke, M. Flexible mechanical metamaterials. Nature Reviews Materials 2, 1–11 (2017).

52. Wagenbauer, K. F., Sigl, C. & Dietz, H. Gigadalton-scale shape-programmable DNA assemblies. Nature 552, 78 (2017).

53. Tikhomirov, G., Petersen, P. & Qian, L. Fractal assembly of micrometre-scale DNA origami arrays with arbitrary patterns. Nature 552, 67–71 (2017).

54. Marchi, A. N., Saaem, I., Vogen, B. N., Brown, S. & LaBean, T. H. Toward Larger DNA Origami. Nano Lett. 14, 5740–5747 (2014).

55. Ong, L. L. et al. Programmable self-assembly of three-dimensional nanostructures from 10,000 unique components. Nature 552, 72–77 (2017).

56. Ke, Y. et al. DNA brick crystals with prescribed depths. Nature Chemistry 6, 994–1002 (2014).

57. Tikhomirov, G., Petersen, P. & Qian, L. Triangular DNA Origami Tilings. J. Am. Chem. Soc. 140, 17361–17364 (2018).

