## Supplemental Information for "Integrating computer-aided engineering and computer-aided design for DNA assemblies"

**This PDF file includes:**

Materials and Methods
Supplementary Text
Supplementary Figs. 1-87
Supplementary Tables 1
Captions for Supplementary Movies 1 and 2

**Other Supplementary Materials for this manuscript include the following:**

Movies S1 and S2
The pdf file for the software user manual
The excel sheets for the staple list of the 14 structures for fabrication

### Supplementary Information Contents

Supplementary Figure 34: An example of iterative design illustrating success and failure at the sub-component level. 56
Supplementary Figure 35: The design details of the Stewart Platform structure. 58
Supplementary Figure 36: Experimental validation of the Stewart platform structure. 59
Supplementary Figure 37: The design details of the compliant compound joint structure. (A) Schematic of the translation and rotation compliant joints. 60
Supplementary Figure 38: Experimental validation of the compliant compound joint structure. 61
Supplementary Figure 39: Representative designs and simulation results illustrating the iterative design process of the gripper structure. 62
Supplementary Figure 40: The design details of the gripper structure. 63
Supplementary Figure 41: Experimental validation of the gripper structure. 64
Supplementary Figure 42: The design details of the trophy structure. 65
Supplementary Figure 43: Experimental validation of the trophy structure. 66
Supplementary Figure 44: The basic unit design of the serial tetrahedron structure. 67
Supplementary Figure 45: The design details of the serial tetrahedron structure. 68
Supplementary Figure 46: Experimental validation of the deployable tetrahedrons in the open configuration. 69
Supplementary Figure 47: Experimental validation of the deployable tetrahedrons in the closed configuration. 70
Supplementary Figure 48: The design details of the umbrella structure. 71
Supplementary Figure 49: Experimental validation of the umbrella structure in the open configuration. 72
Supplementary Figure 50: Experimental validation of the umbrella structure in the close configuration. 73
Supplementary Figure 51: The design details of the butterfly mechanism. 74
Supplementary Figure 52: Experimental validation of the butterfly mechanism without actuation. 76
Supplementary Figure 53: Experimental validation of the butterfly structure after actuation. 77
Supplementary Figure 54: Experimental validation of polymerization of the butterfly mechanism 78
Supplementary Figure 55: The functionality and features in MagicDNA to broaden the design spectrum. 81
Supplementary Figure 56: Examples of lattice-based structures. 83
Supplementary Figure 57: Example of surface-based structures. 84
Supplementary Figure 58: Examples of wireframe structures. 85
Supplementary Figure 59: Examples of hybrid structures. 86
Supplementary Figure 60: Other complex design examples. 87
Supplementary Figure 61: Examples of designs with interlocking features. 88
Supplementary Figure 62: Examples of wireframe structures with complex 3D arrangement and connectivity. 89
Supplementary Figure 63: Other examples. 90
Supplementary Figure 64: Other examples. 91
Supplementary Figure 65: Examples of written designs. 92
Supplementary Figure 66: Other examples. 93
Supplementary Figure 67: Other examples. 94
Supplementary Figure 68: Other examples. 95

#### Materials and Methods

##### Multi-component assembly design and software availability

MagicDNA is an open-source software available at https://github.com/cmhuang2011/MagicDNA. It was coded in MATLAB 2017a and is compatible with newer versions of MATLAB. Detailed descriptions for installing the software are in the software user manual. Additional material including tutorial movies can be accessed through the Supplementary Material or the YouTube channel "MagicDNA software". Output files from MagicDNA include the staple sequence list for ordering staples, caDNAno .JSON files for fine-tuning of strand routing details, and oxDNA topology and configuration files for validating the design profile with coarse-grained simulation.

##### Typical Design workflow in MagicDNA

The general design workflow consists of four steps (Fig. S3): 1) Define the overall geometry and the geometries of each component, 2) Assemble the components by forming stiff or flexible joints between them, 3) Use the routing algorithms and fine-tune the routing if necessary, and 4) Generate topology and configuration files for coarse-grained simulations. For the top-down approach in geometry, either sketching lines in MagicDNA or importing a line model through .STEP files is needed to convert lines to bundles with also inputting design parameters like cross-section and lengths. Alternatively, one can remove or insert components to the assembly using a bottom-up approach. The next assembly step includes manipulations of each component or a set of grouped components to arrange a desired 3D assembly configuration, connecting the components by specifying the connectivity matrix and/or using the optional manual mode to specify locations of connections between components, and finally specifying the single-stranded scaffold lengths. Once the routing algorithms receive the design parameters from the geometry and assembly steps, the scaffold and staple routings are automatically generated with the option of fine-tuning in caDNAno[1]. Lastly, using the automatically generated simulation files to conduct the coarse-grained simulations allows users to evaluate the design and provides feedback to guide modifications in the next iteration if needed. This design process is illustrated in detail for a hinge example in Supplementary Movie S1.

##### Coarse-grained MD simulation

The topology and initial configuration files were generated directly from MagicDNA. The relaxation algorithm was similar to our previous study[2] adapted from standard oxDNA relaxation protocols[3]. The relaxation is carried out in three steps: oxDNA1, oxDNA2 relaxations with gradually increasing coefficients, and a short simulation, all with mutual traps between paired scaffold and staple bases. After relaxation, the oxDNA2 interaction model was used to conduct coarse-grained simulations without applying any mutual traps. For most simulations, a total of $10^7$ steps with GPU acceleration were used. For the 4-bar mechanism, $3\times10^8$ steps were used to get a better depiction of the motion. The simulation time for each step was set to 15.15 fs. Simulation parameters included an Anderson-like thermostat, temperature at 30 °C, and monovalent salt concentration at 0.5 M, all standard conditions in oxDNA simulations[4,5]. The frequency to save the current configuration into the trajectory was set as either $10^6$ or $5\times10^5$ steps. The processes mentioned above were executed through a shell script for all structures in this study in a Linux computer equipped with a NVIDIA GeForce 1080Ti graphics card. The trajectory file was later analyzed in MATLAB, including visualization of configurations, root-mean-square deviation (RMSD), and root-mean-squared fluctuations (RMSF). The average configurations were exported to the UCSF Chimera[6] software and rendered to high-quality images.

**Assembly and Fabrication of DNA Origami structures**

All DNA staple strands were ordered and synthesized with salt-free purification and in 10 nmole scale from Eurofins (Louisville, KY), except the staple strands of the ring structure in 25 nmole scale from IDT (Coralville, IA). Scaffolds for single-scaffold structures were made in-house as described in[7] or purchased from Guild Biosciences (Dublin, OH) for M13mp18 derived scaffolds. Scaffolds for multi-scaffold structures were kindly provided by the Dietz lab at Technische Universität München[8]. Each structure was folded (thermal cycler from Bio-Rad, Hercules, CA) and optimized for solution conditions (i.e. salt, scaffold, and staple concentrations), annealing ramp protocol, and in some cases gel running conditions (i.e. salt concentration in gel). Single-scaffold structures were folded with 200 nM staples and 20 nM scaffold. Multi-scaffold structures and single-components of multi-scaffold structures were folded with 110 nM staples and 10 nM scaffold or folded with 100 nM staples and 10 nM scaffold. Each folding reaction contained a buffer solution consisting of 5 mM Tris, 5 mM NaCl (pH 8), 1 mM EDTA, and varying $MgCl_2$ conditions found in the respecting supplemental figure captions. Folding conditions varied by structure, and specific details for all structures are provided in Supplementary Table 1. Thermal annealing ramps were also tailored for individual structures (details also in Supplementary Table 1). The different annealing ramps used included a two-and-a-half-day fold[7] starting with a 1 hr/°C from 65-61°C melt, followed by, 2 hr/°C from 60-40°C anneal, and a cool step 30 min/°C from 39-4°C ; a four-and-a-half-day fold starting with a 1hr/°C from 65-62°C melt, 2hr/°C from 61-59°C anneal, 5hr/°C from 58-46°C anneal, 2hr/°C 45-40°C cooling , and a final cooling step 1hr/°C from 39-4°C. Single-scaffold structures folded in rapid folds[9] all include a 15min 65°C melt: then an anneal 4hr/°C in non-linear increments from 60-40°C , and a 4hr/°C from 56-50°C anneal. The multi-scaffold structures folded using an annealing protocol described by Engelhardt et al.[8] starting with a 65°C melt for 15 minutes, followed by an anneal 3hr/°C from 60-40°C then a cool at 4°C.

**Purification of DNA Origami**

Each DNA origami structure was purified and analyzed post-folding reaction via agarose gel electrophoresis. Buffer conditions included 0.5x TBE (45 mM Boric acid, 45 mM Tris base, and 1 mM EDTA) with either 5.5 mM or 11 mM $MgCl_2$ and agarose gels from 1.5-2% agarose and 0.5μg/mL ethidium bromide. 1.5% agarose gels with 0.5x TBE and 5.5 mM $MgCl_2$ buffer[8] were used for all multi-scaffold DNA origami structures and components as well as the umbrella closed configuration and trophy. All other structures were purified with 2% agarose gels and 0.5x TBE and 11 mM $MgCl_2$ running buffer. Each gel was run at 90V for 90-120 minutes in an ice water bath. Gels were imaged on a UV table using a FotoDyne Express FOTO/Analyst system. Details for gel purification are also summarized in Supplementary Table 1.

**Actuation and polymerization of DNA Origami**

The butterfly mechanism was actuated post-fold and gel purification. The structure concentration was quantified via Nanodrop as ~3 nM. 10μL of gel-purified structure was then combined with 2μL of actuation staples for a final concentration of 2.5 nM structure, 25 nM actuation staples and 10 mM $MgCl_2$ (10x excess concentration of actuation staples relative to the concentration of the structure). The mixture was incubated at 37°C for 2 hours. After actuation of structures, polymerization staples were added at 150 nM. The final solution contains 10μL of structure at 2 nM, 2 μL of actuation staples at 21 nM, and 2 μL of polymerization staples at 21 nM and ~8 mM $MgCl_2$. The solution was then incubated again at 37C for 2 hours..

**DNA Origami Analysis and Imaging via TEM**

Structures were suspended in respective running buffer conditions post purification with concentrations between 1-5 nM depending on structure yield. The trophy and Stewart platform structures (see Supplementary Table 1) were additionally incubated with the peptide K10 (kindly provided by the Stephanopoulos Lab at Arizona State University) at a ratio of 0.5N:P for ~30 mins prior to preparing TEM samples to improve contrast[10]. A sample volume of 4 μL was deposited onto a plasma-cleaned Formvar-coated 400 mesh copper grid (Ted Pella, Inc.) with incubation times between 4-8 minutes prior to wicking away the solution with filter paper. For the trophy, umbrella closed configuration, robotic manipulator, logo, and airplane structures (see Supplementary Table 1), a 4μL droplet of 30 mM $MgCl_2$ was added to the plasma-cleaned grid prior to sample incubation and wicked away after 2 minutes followed by adding the sample drop to enhance surface deposition. After wicking away the sample drop, a 10μL droplet of staining solution consisting of 2% uranyl formate + 25 mM NaOH was added to the grid, immediately wicked away, followed by adding a 20μL droplet of the same staining solution incubated for 40 seconds and finally wicking away the stain solution. Samples were allowed to dry for at least at least 20 minutes before imaging. The structures were imaged at the OSU Campus Microscopy and Imaging Facility on a FEI Tecnai G2 Spirit TEM with an acceleration of 80kV.

EMAN2[11] and ImageJ[12] were used for post-processing and analysis of raw TEM images. Old Particle Picker (e2boxer_old.py) in EMAN2 was used to select particles from raw TEM TIF files. At least 300 particles and up to 900 particles were used to create particle sets for image averages. Particle sets were then built and 2D analysis with 4 ncls (number of classes) and 3-8 iterations were performed for image averaging. Particles from the 4-bar mechanism were used separately in a MATLAB code for a 5-point analysis with manual selection. ImageJ set scale function was used for scale bars on TIF files and brightness/contrast/FFT bandpass filter were applied in ImageJ analysis.

**Supplementary Table 1. Summary of the experimental conditions for all fabricated structures.**

| Structure | Scaffold (bases) | No. of Staples | Folding Ramp* | $MgCl_2$ [mM]** | Scaffold [nM] | Staples [nM] | Gel % $MgCl_2$ | TEM grid prep |
|---|---|---|---|---|---|---|---|---|
| Nanopore | p7249 | 155 | 2.5 day fold | 20 | 20 | 200 | 2% 11 mM | standard |
| Ring | p7560 | 183 | 50°C 4hr/°C anneal | 20 | 20 | 200 | 2% 11 mM | standard |
| Hollow Hinge | p8064 | 177 | 58.2°C 4hr/°C anneal | 20 | 20 | 200 | 2% 11 mM | standard |
| 4-bar Mechanism | p8064 | 206 | 2.5 day fold | 20 | 20 | 200 | 2% 11 mM | standard |
| Stewart Platform | p7249 | 157 | 2.5 day fold | 24 | 20 | 200 | 2% 11 mM | $K_{10}$ |
| Compliant Joint | p8064 | 172 | 2.5 day fold | 18 | 20 | 200 | 2% 11 mM | standard |
| Gripper | p7560 | 175 | 2.5 day fold | 22 | 20 | 200 | 2% 11 mM | standard |
| Trophy | p8064 | 209 | 2.5 day fold | 20 | 20 | 200 | 1.5% 5.5 mM | $K_{10}$ or 30 mM $MgCl_2$ |
| Tetrahedron open | p8064 | 184 | 2.5 day fold | 16 | 20 | 200 | 2% 11 mM | standard |
| Tetrahedron closed | p8064 | 178 | 2.5 day fold | 14 | 20 | 200 | 2% 11 mM | standard |
| Umbrella open | p8064 | 186 | 2.5 day fold | 22 | 20 | 200 | 2% 11 mM | standard |
| Umbrella closed | p8064 | 177 | 4.5 day fold | 22 | 20 | 200 | 1.5% 5.5 mM | 30 mM $MgCl_2$ |
| Butterfly | p7249 | 194 (5 for unused scaffold) | 53°C 4hr/°C anneal | 20 | 20 | 200 | 2% 11 mM | standard |

| | | | | | | | | |
|---|---|---|---|---|---|---|---|---|
| Robotic Manipulator Components | p8064 or CS4_7557 | 164(Arm) 175(Claw) 172(Twz) | 2.5 day fold | 20 or 16 | 10 | 110 | 1.5% 5.5 mM | standard |
| Robotic Manipulator | p8064 CS4_7557 | 339 (ArmClaw) 336 (ArmTwz) | 3hr/°C 60-40°C | 20 | 10 | 110 | 1.5% 5.5 mM | 30 mM $MgCl_2$ droplet |
| Logo | p8064 CS5 7559 | 348 | 3hr/°C 60-40°C | 10 | 10 | 110 | 1.5% 5.5 mM | 30 mM $MgCl_2$ droplet |
| Airplane | p8064 CS3_L_7560 CS4_7557 CS5_7559 | 682 (16 for unused scaffold) | 3hr/°C 60-40°C | 8 | 10 | 100 | 1.5% 5.5 mM | 30 mM $MgCl_2$ droplet |

*Each ramp had a 15min 65°C melt and ended with cooling to 4°C
**Each folding reaction contained a buffer solution consisting of 5 mM Tris, 5 mM NaCl (pH 8), 1 mM EDTA

#### Supplementary Text

##### Current design software

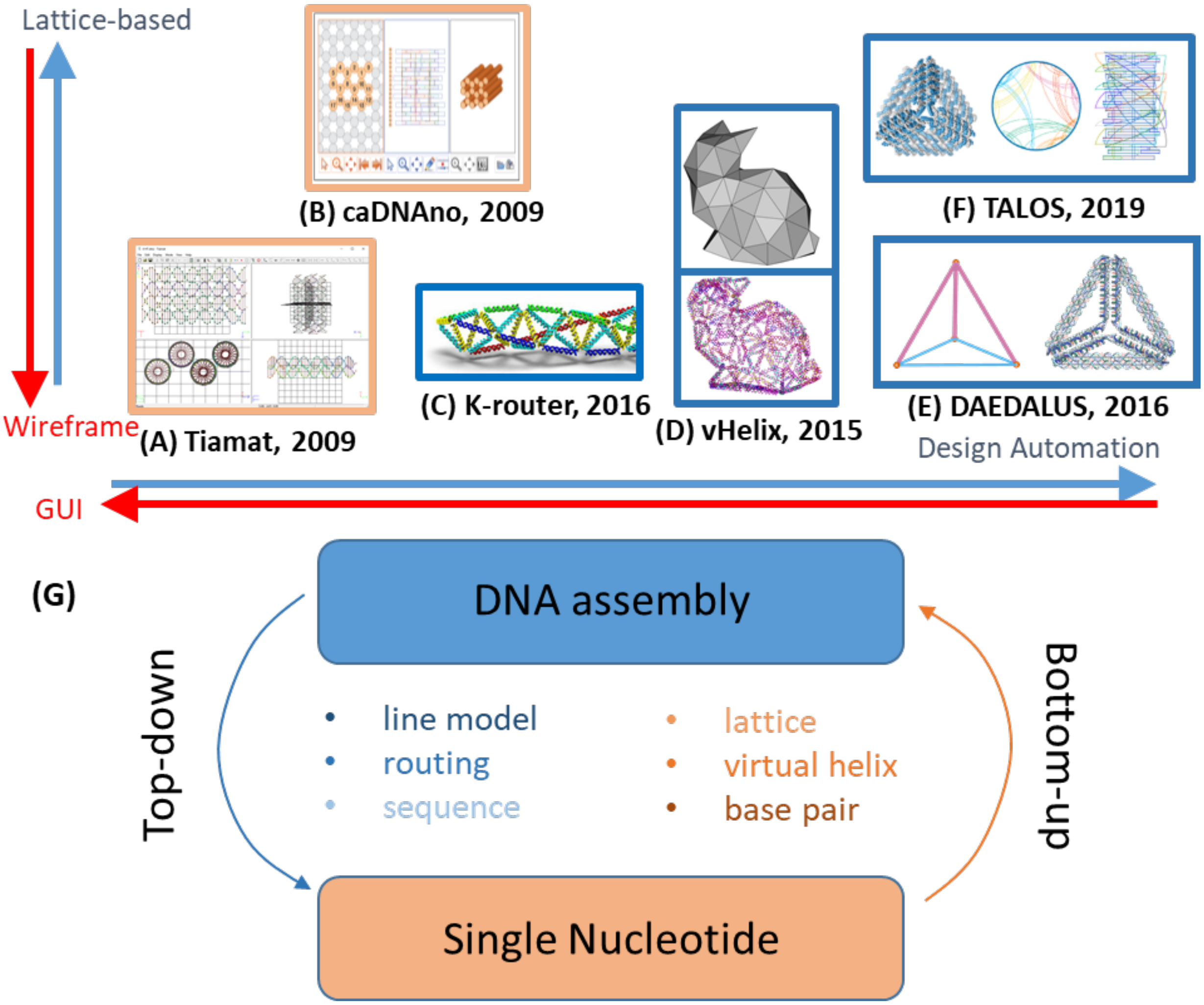

**Supplementary Figure 1: Survey of current DNA-based design software**. (A) Tiamat aimed to design large molecular systems like DNA origami using a graphical user interface workflow, especially for wireframe structures13. (B) caDNAno is the most widely used software for 2D sheet-like and 3D lattice-based DNA origami design1. (C) K-router created triangulated grid interface for users to specify the scaffold routing and export to caDNAno for further editing14. (D) vHelix is a plug-in for Autodesk Maya to render 3D polyhedral mesh, followed by BSCOR scaffold routing algorithm to solve A-trails problem15. (E) DAEDALUS is a fully automatic software to design 3D wireframe DNA origami structures by simply providing 3D polygon files from common CAD software16. (F) Enhancing the stiffness of each edge in DAEDALUS, TALOS converts each line in polygon files to 6-HB (helix bundle) honeycomb bundles to enhance mechanical stiffness17. (G) The design process in these software tools are considered either top-down (start with a model of the overall DNA assembly) or bottom-up (design carried out at the nucleotide/strand level to construct the larger DNA assembly) approaches to DNA origami design.

#### Supplementary Section 1: Iterative design process with simulation feedback

Software package, specifying the geometries, and assembly

Modern computer-aided design software packages for typical engineering have both a part (or component) design interface and an assembly design interface. We choose to create a similar hierarchical design approach using MATLAB object-oriented programming to organize the data and functions (Fig. S2). In this model, a component (Fig. S3A), as a bundle class (either square or honeycomb lattice as sub-classes), consists of geometric data (i.e. the cylinder model) along with a 3D transformation matrix for tracking the rigid-body motions. On the other hand, a mechanism class deals with the properties and the functions related to the assembly and the routing algorithm. In addition to the data structure, we also designed graphical user-interfaces (GUIs) to present the data into visualized objects that facilitate straightforward design steps including adding and organizing components into a DNA origami assembly directly in 3D space, which greatly facilitates the design of complex assemblies,.

For specifying the geometry as the first step (Figs. 1A, S3B, and S4 to S6), both top-down and bottom-up design approaches are available, similar to modern CAD software. For defining the double-stranded geometries of the bundles, the top-down approach is to use a line model like sketching a mechanism, either by importing a .STEP file exported from another CAD software or directly sketch in the line model GUI. Then the user can specify the cross-sections as profiles that will be extruded along the length of each line to create the overall cylinder model. On the other hand, once bundles are created, they can be saved as a single component or as a group of components into a library for inserting into an assembly, which we refer to as the bottom-up approach. In either case, to increase the design flexibility of each bundle at the nucleotide level, (Fig. S5) the bundle editing GUI allows users to adjust the lengths of the helices in a bundle using convenient 2D and 3D visualizations as a guide.

The second step is to assemble the bundles. The cylinder model is mathematically equivalent to an array of parallel lines with 2 nm spacing between neighboring lines, which provides a simplified representation (Fig. S7). Before starting to manipulate the bundles, the program examines the pairing of cylinders (pairs will be connected by external scaffold crossovers in the scaffold routing) to avoid infeasible structures based on constraints imposed by our scaffold routing algorithm (e.g. components must have an even number of helices). Using the data of the cylinder model, a bundle in the assembly GUI is composed of an array of cylinders as source, a tetrahedron-meshed volume representation, and allowable nodes to connect with other components. The end nodes are the extensions of pairs of cylinders to form vertex or joint designs while the side nodes occur at locations where a scaffold nick would be oriented on the surface. We omit unlikely connection sites from the inside of a bundle by excluding the side nodes inside the tetrahedron volume representation. These cylinders, volumes, and nodes are graphically displayed as a set of point clouds with topological connections, and can be translated by the vector, $t$ (Eqs. 1 and 2). In addition, the rotation along the center $G_c$ can be achieved using the rotation matrices $R$ along the global coordinates (Eqs. 3 and 4).

$$p_{3,(i+j+k)} = [p_{c,i}, p_{v,j}, p_{n,k}] \quad (1)$$

$$p' = p + t \tag{2}$$

$$[R]_x = \begin{bmatrix} 1 & 0 & 0 \\ 0 & \cos\theta & \sin\theta \\ 0 & -\sin\theta & \cos\theta \end{bmatrix}$$

$$[R]_y = \begin{bmatrix} \cos\theta & 0 & -\sin\theta \\ 0 & 1 & 0 \\ \sin\theta & 0 & \cos\theta \end{bmatrix} \tag{3}$$

$$[R]_z = \begin{bmatrix} \cos\theta & \sin\theta & 0 \\ -\sin\theta & \cos\theta & 0 \\ 0 & 0 & 1 \end{bmatrix}$$

$$p' = [R](p - G_c) + G_c \tag{4}$$

For manipulating bundle components, we also provide the option that users can translate or rotate the bundles along their local coordinates, where the bundle translates along the cylinder direction and the orthogonal axes of the cross-section, and similarly rotates about these local axes by a user-defined increment, $\phi$ (Eqs. 5-7).

$$s = (s_x, s_y, s_z)^T \tag{5}$$

$$[S] = \begin{bmatrix} 0 & -s_z & s_y \\ s_z & 0 & -s_x \\ -s_y & s_x & 0 \end{bmatrix} \tag{6}$$

$$[R] = [I] + \sin\phi[S] + (1 - \cos\phi)[S^2] \tag{7}$$

Once the bundles have been moved to a position to approximate a desired assembly configuration, users need to specify the connectivity matrix between bundles to search the connections based on the distances between nodes that define potential connection sites (Fig. S8). Currently the nodes can be only connected to one other node, which represents a double-scaffold crossover connections. The manual mode provides users freedom to specify the locations of connections providing more control over the specific nucleotide level connections. This is also useful when it is desired to make connections between nodes that are potentially distant in the assembly model, such as when intentionally bending single-layer bundles (e.g. trophy design Figs. S42-S43).

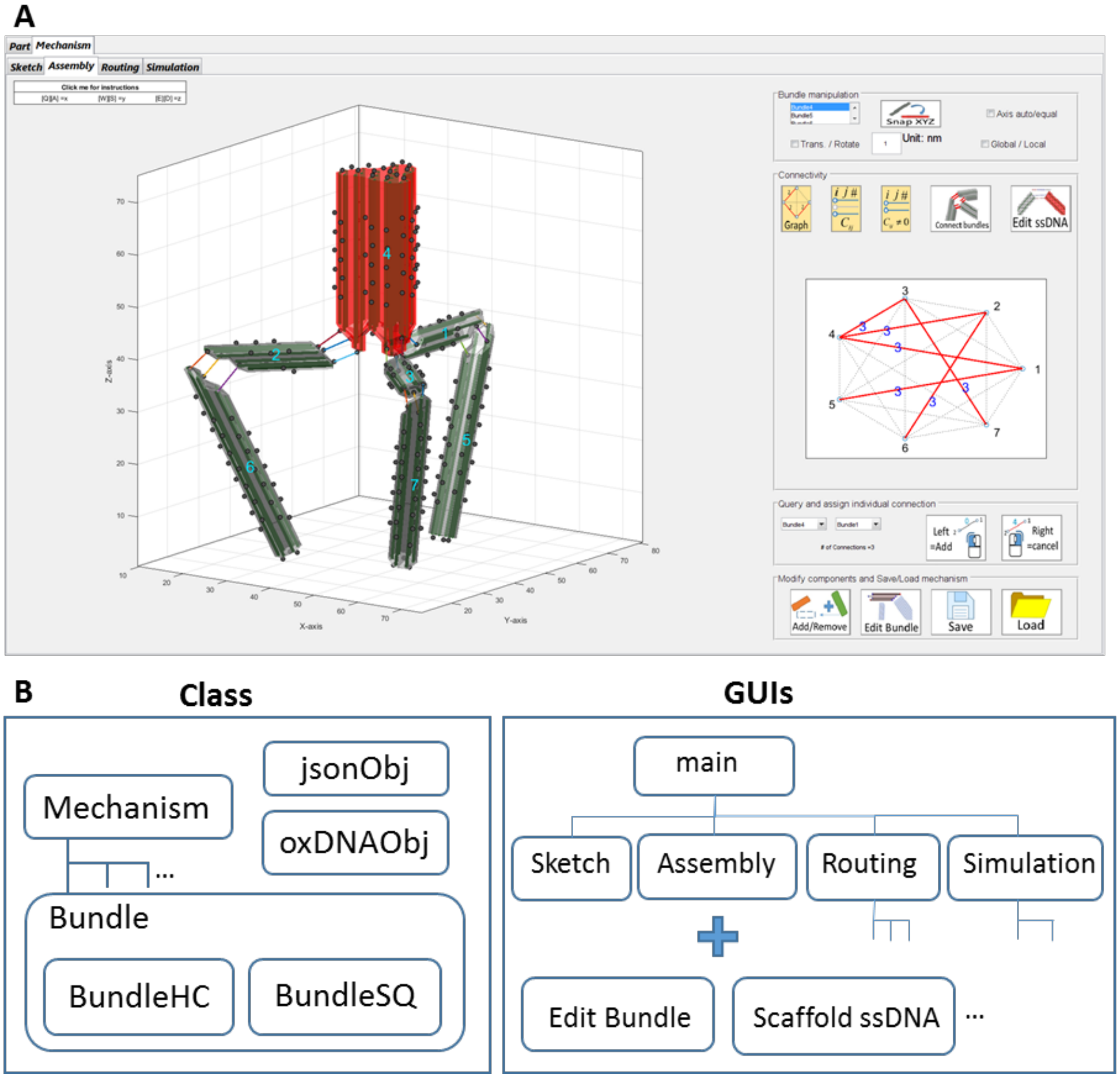

**Supplementary Figure 2: Main interface and MATLAB class and GUI structure in MagicDNA. MagicDNA is a MATLAB app with graphical user interfaces that allows users to visualize and edit a design in the 3D graphical design panel**. (A) A snapshot of the main assembly GUI where users can manipulate the bundles and search for connections based on distances between connection site nodes. (B) (Left) To organize the data, we used object-oriented programming in MATLAB, in which a DNA origami design is an instance of "Mechanism" class, which is where most of the algorithms (e.g. assembly and routings) are carried out. Bundles are either honeycomb-lattice or square-lattice objects with the corresponding geometrical data. (Right) Several sub-GUIs were created to handle different visualizations and to collect user inputs. There are four main GUIs that correspond to four major design steps: Sketch (converting lines to bundles), Assembly, Routing, and Simulation, which follows the most common main design workflow. Other local UIs were coded and implemented within the main UIs for more specific tasks, such as custom cross-sections and extrusion of cylinders.

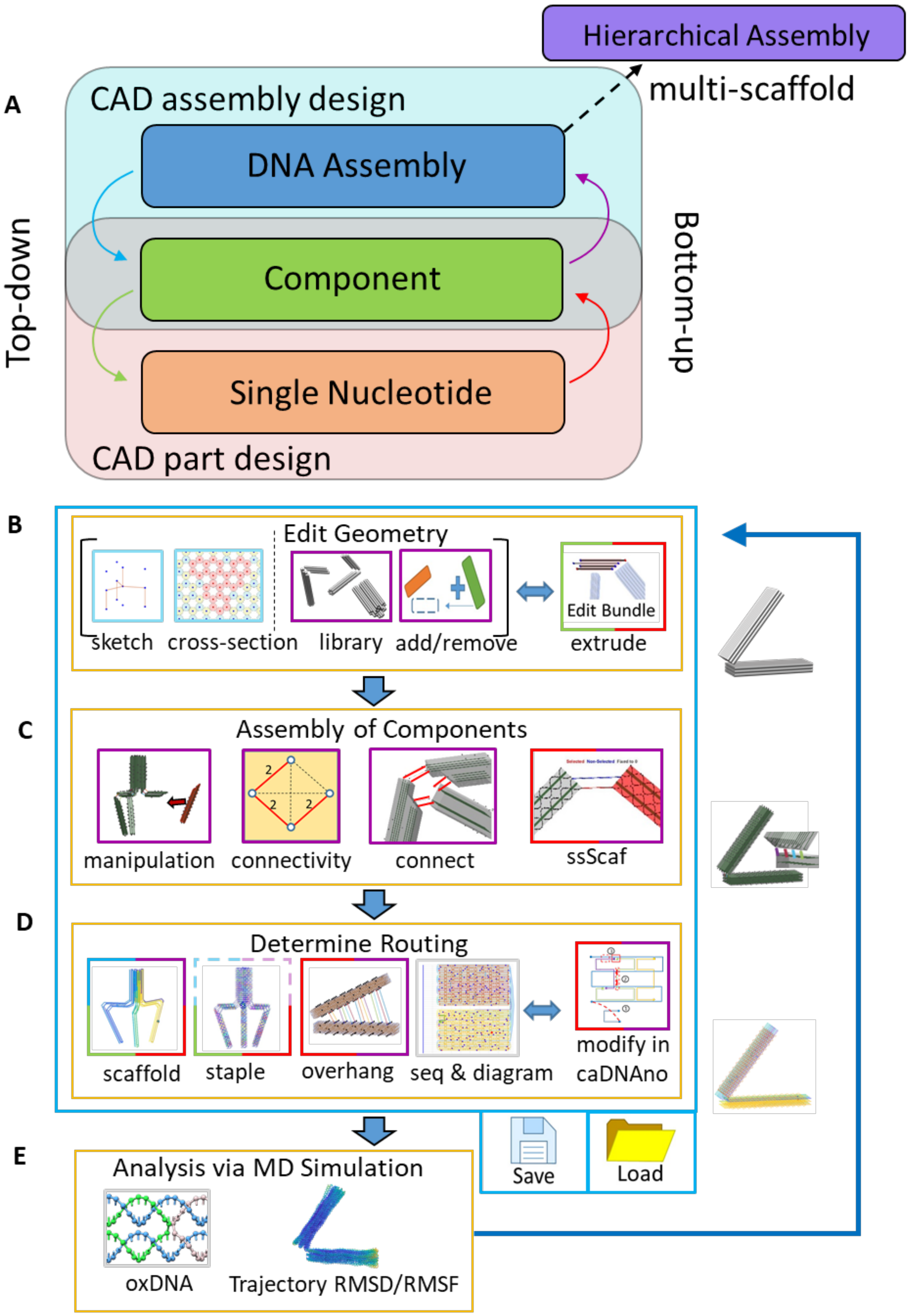
A
Hierarchical Assembly
multi-scaffold
CAD assembly design
DNA Assembly
Component
Single Nucleotide
CAD part design
Top-down
Bottom-up
B
Edit Geometry
sketch
cross-section
library
add/remove
Edit Bundle
extrude
C
Assembly of Components
manipulation
connectivity
connect
ssScaf
D
Determine Routing
scaffold
staple
overhang
seq & diagram
modify in
caDNAno
Save
Load
E
Analysis via MD Simulation
oxDNA
Trajectory RMSD/RMSF

**Supplementary Figure 3: The proposed hierarchical model for design process with GUIs at each level.** (A) Schematic of the hybrid top-down and bottom-up framework where the overall design process combines operations at the nucleotide, component (or bundle), and assembly levels. In particular, the addition of the bundle component level between the DNA assembly and nucleotide levels allows for straightforward manipulations and convenient bottom-up and top-down flow of information to enable design of complex assemblies. Many GUIs and algorithms in (B)-(E) are created for defining parameters and communicating the data across these three levels (top-down, bottom-up or hybrid). Each block in (B)-(E) represents a function in the GUI (generally a button, drop down list, or text entry box) for user inputs or data visualization. Each block can be categorized and color-coded corresponding to the data flow. An example hinge design on the right illustrates the status of key steps in the process. (B) The design process typically starts with converting lines in a sketch into cylinder bundles (Top-down). These bundles can be created, added/removed and exported into the library for importing back into a mechanism (Bottom-up). To increase design flexibility at the base level, helices within a selected bundle can be extruded to have non-uniform helix lengths in a component (e.g. for edge gradients). (C) In the assembly step, bundle component positions can be fine-tuned from their initial positions defined in the line model to the position and orientation for a desired assembly configuration with the manipulation panel and keyboard inputs that input rigid body motions. Then, a connectivity matrix is created based on the user specified number of connections between components. Users can also specify the lengths of single-stranded connections. A search algorithm is executed to automatically determine the actual connection points based on smallest distances between potential connection points guided by some user-input (e.g. number, end-to-end type, end-to-side type, side-to-side type, and manual assign). Once the double scaffold connections are specified, the length of individual single-stranded scaffold connections can be manually adjusted in the GUI with 3D visualization of the helical geometry. (D) The scaffold and staple routing are automated with built-in algorithms (Fig. S9 for scaffold and Figs. S13 for staples). Both routing algorithms allow users to specify routing parameters (i.e. poly-T ends and staple overhangs) for the sake of computational cost and design customization. The GUI for data exchange with caDNAno offers as an option to export/import designs for fine-tuning of routing, which is necessary for some features like multi-way junctions. (E) Coarse-grained simulations feedback to guide design iteration and verify final designs. Basic analysis tools like RMSD and RMSF are included in the software.

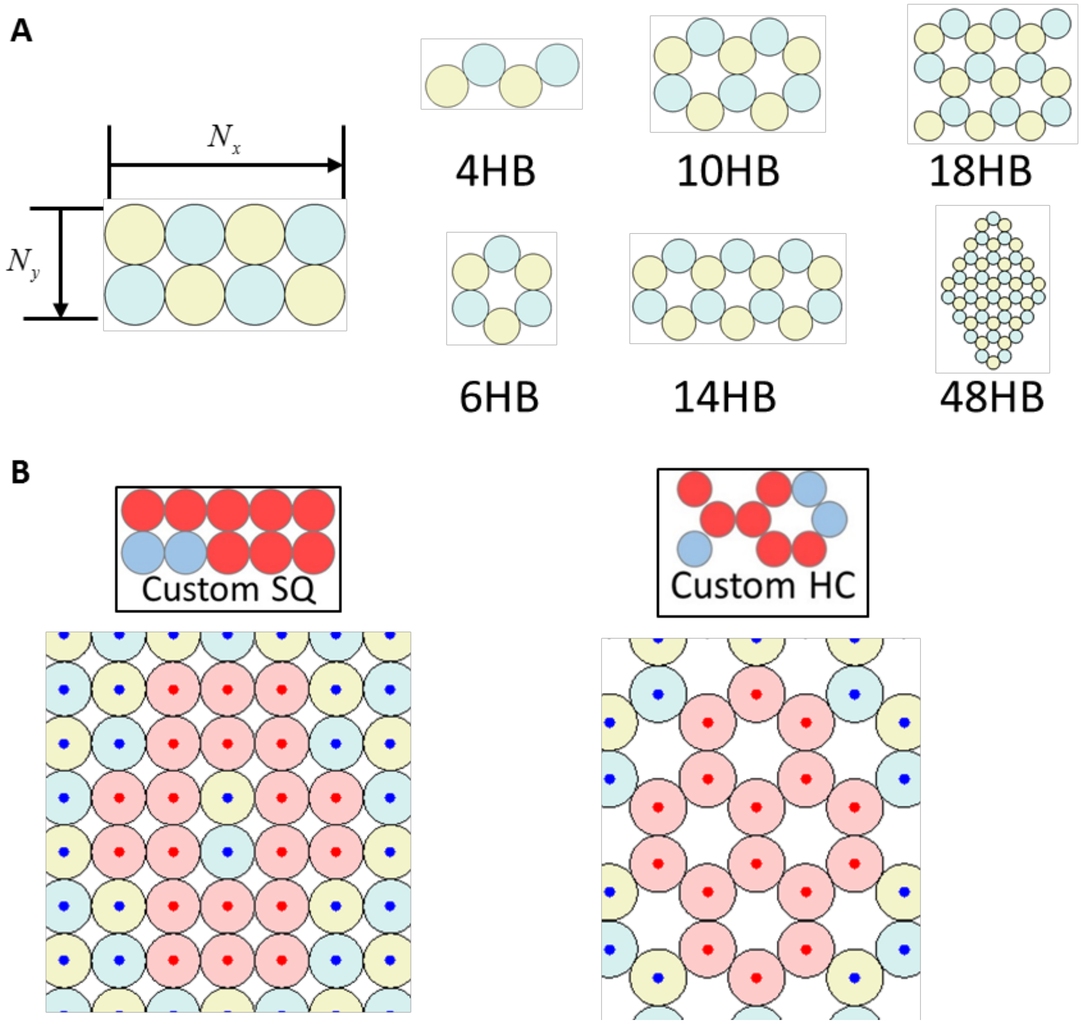

**Supplementary Figure 4: Default and custom cross-sections for square and honeycomb lattices.** (A) Cross-sectional geometry of bundles can be selected for many standard bundles (e.g. 6-Helix-Bundle) or typed in when converting line models to bundles. (B) Alternatively, custom cross-sections can be defined by selecting circles in a GUI tool for more complex cross-sections, with the constraint of having an even number of cylinders. Currently the software allows the user to store five different custom cross-sections for each lattice for future use.

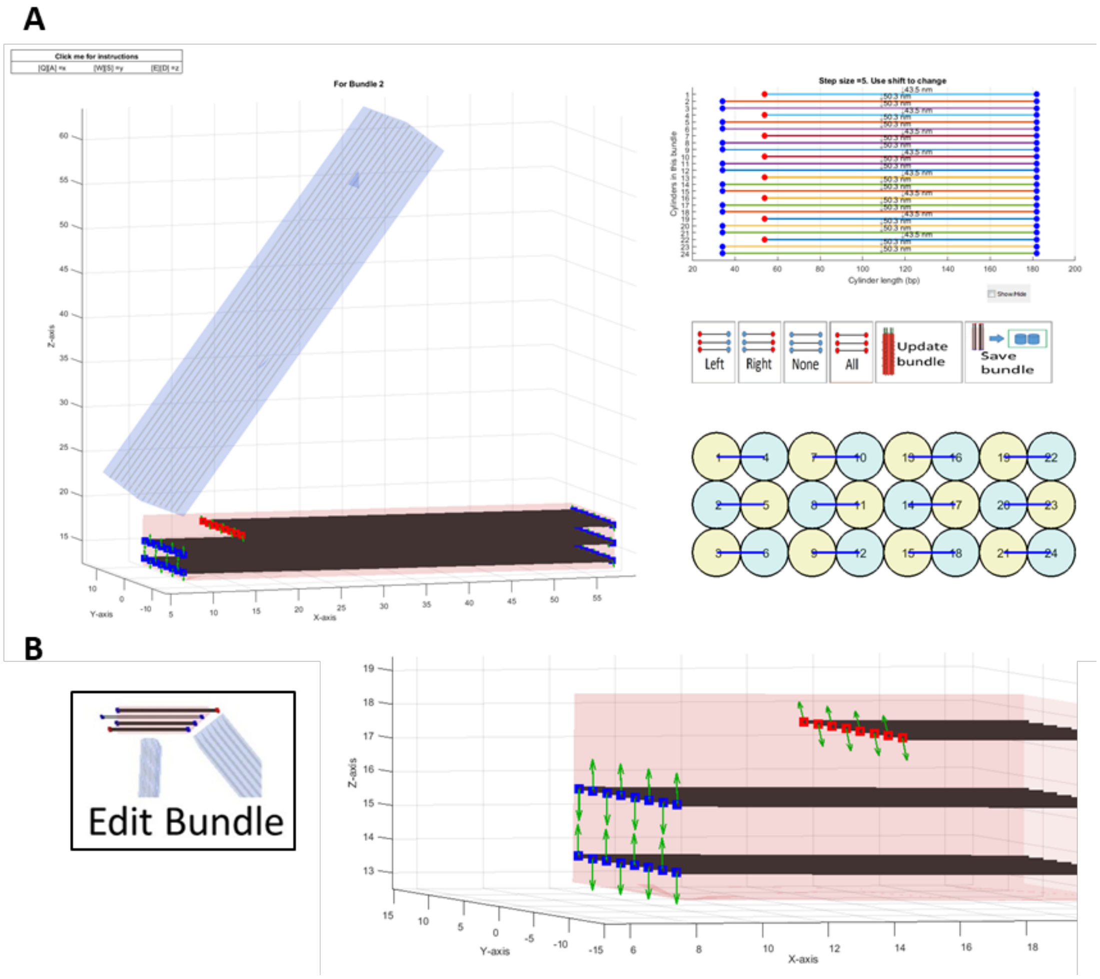

**Supplementary Figure 5: The bundle editing GUI. This GUI allows the user to extrude individual helices in a component while visualizing the cylinder model of the selected bundle.** The left side shows the 3D model while highlighting the selected bundle(s) (shown in red) with the other bundles still visualized for reference. The right-side shows a 2D path diagram (top) and the cross-section view (bottom). Note that the line or cylinder model here defines the double-helical portion. The end nodes (red dots at ends) are synchronized in the 2D and 3D views upon selection or manipulation, and one can adjust the length of each cylinder while the scaffold helical orientations at the ends are shown as the green arrows in the 3D panel (bottom right inset). The cross-section view in the bottom-right corner also shows the pairing of cylinders connected by external scaffold crossovers (blue lines in the 2D cross-section view).

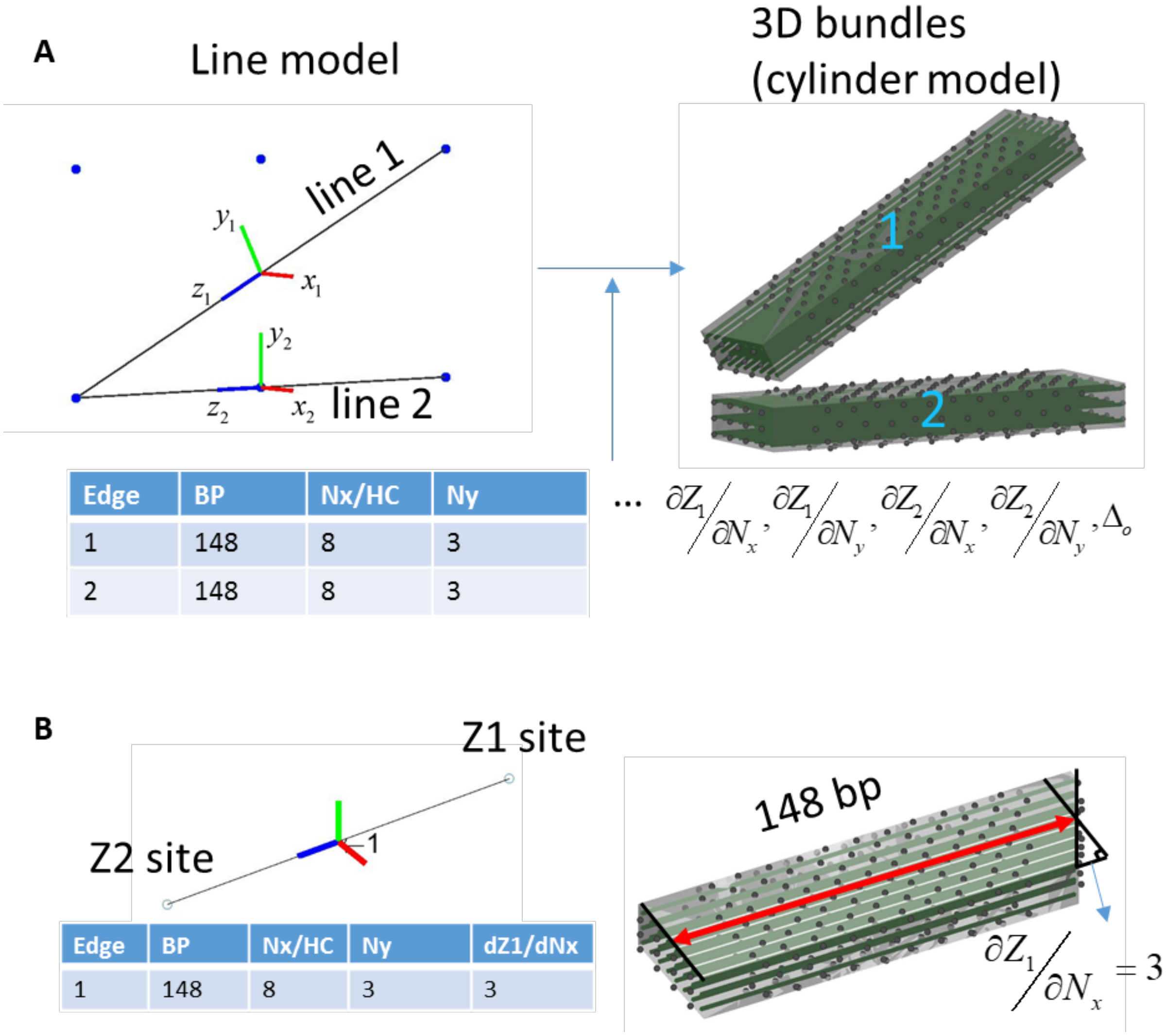

**Supplementary Figure 6: Converting lines to bundles.** (A) (Left) The line model created using the sketch tool in MagicDNA. By specifying a table with the required geometric parameters (i.e. length, cross-section, end gradient), one can convert two lines into two bundles of desired cross-section, here 8×3 square-lattice. The local coordinates of the line model (Red-Green-Blue for X-Y-Z directions) were used to provide rough positions and orientations of the bundles, which can be adjusted with rigid body motions later in the assembly step. (B) Schematic that illustrates the input of gradient properties of the line to create bundles with a spatial gradient along one end created by making different layers of helices different lengths.

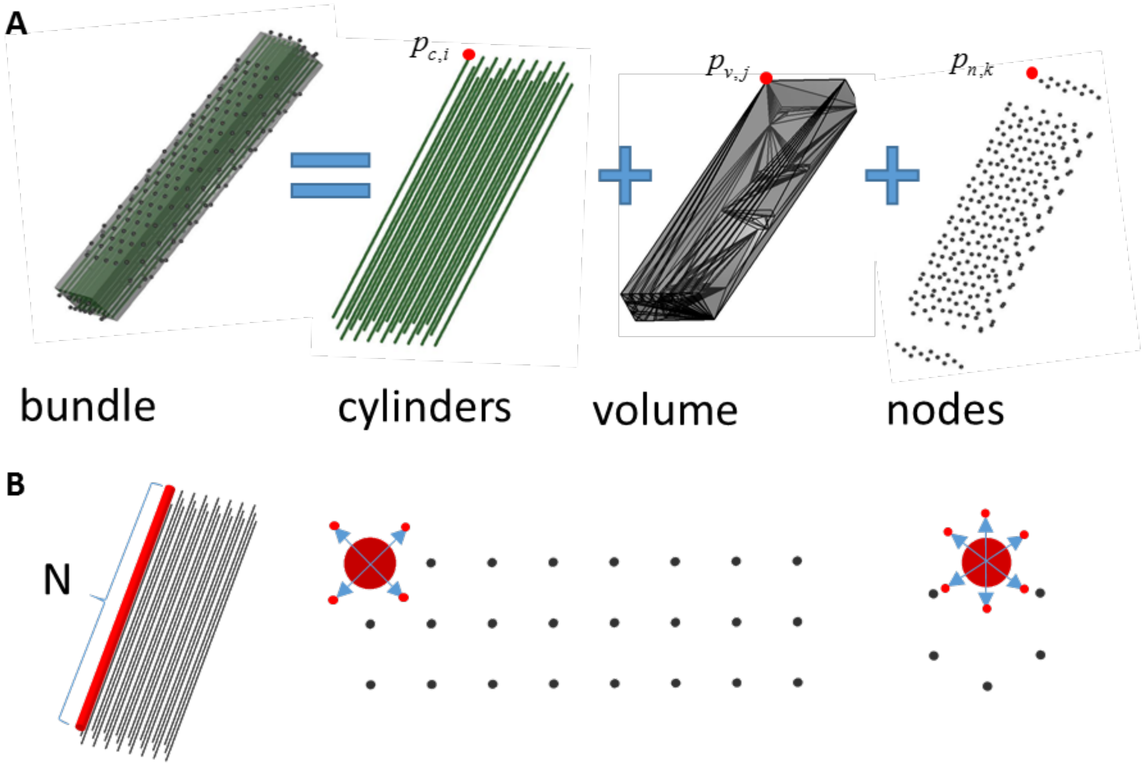

**Supplementary Figure 7: Graphical representations of point clouds and secondary objects (tetrahedron mesh and lines).** (A) In the assembly GUI, each bundle is composed of three graphical representations: cylinders, volume and nodes. Cylinders are created with a cross-section and extrusion from the sketch step. (B) The program slices each cylinder into N sections and each section extends to 4 (square lattice) or 6 (honeycomb lattice) corners as the collection of point clouds, which is later simplified into tetrahedral meshes. Those side nodes surrounding the volume are positions where nicks in the scaffold would be oriented in an outward helical position. These are positions where double-scaffold crossover connections to other bundles are allowed on the side surfaces of bundles. All nodes inside the volume are excluded for simplification.

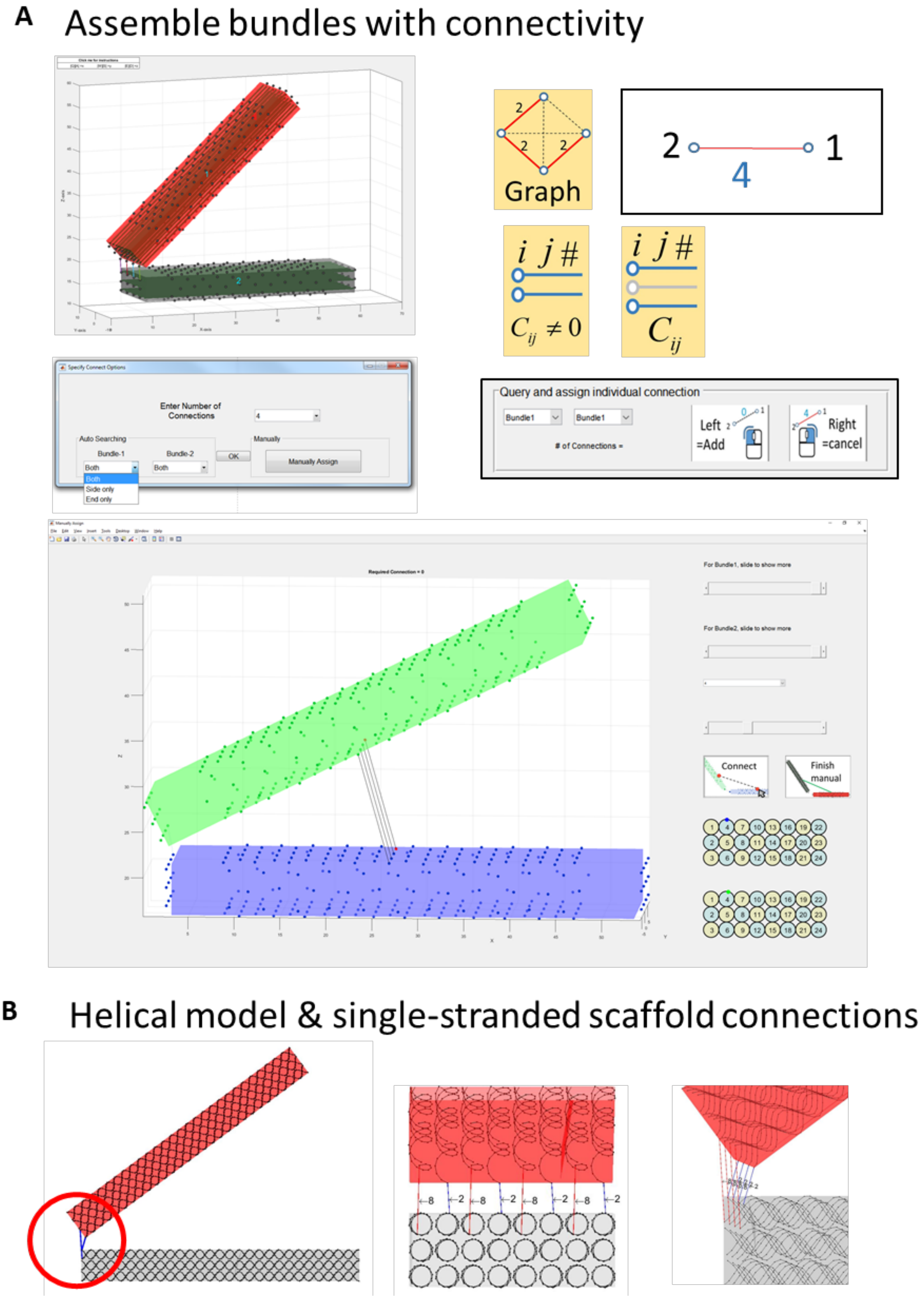

**Supplementary Figure 8: Defining the connectivity matrix between bundles and specifying the lengths of the single-stranded scaffold connections with visualization of the helical geometry for guidance.** (A) The connectivity panel visualizes the connectivity matrix into a graph where the nodes are bundles and the number label on the edges indicates the number of connections between the two nodes, or bundles (red edges in graph diagram indicate non-zero connectivity). Clicking on the line calls up a dialog box to input the number of connections, specifying whether connections should occur on end or side nodes (or either), or the user can choose to manually assign connections (shown below). As the number of bundles increases, the connectivity graph also has second-order growth and it becomes hard to click on correct lines with the mouse. Hence, we provide two other representations for the connectivity matrix as a list of connectivity between bundle *i* and bundle *j*; one table lists the connectivity between bundles with the option of excluding or including non-zero connectivity, and the other representation is a set of two popup menus that can be used to query or assign connectivity by selecting bundles on the 3D model or selecting bundles from drop down menus. In manual mode, just the two selected bundles are visualized and the user can select the nodes on each bundle to specify the connections regardless of distances. (B) Once the connectivity matrix and the connections between bundles are determined, the next step is to switch to the finer helical model and specify the lengths of single-stranded scaffold connections. Notice the external double-scaffold crossovers indicate that each connection in the assembly GUI (A) is equivalent to two scaffold connections in the helical model (B).

Scaffold routing algorithm

In DNA origami designs, the scaffold strand has to traverse the entire structure, and the staple strands are programmed to bind in a piece-wise complementary manner to the scaffold. The periodic double-crossovers between neighboring helices formed mostly by staple strands constrain the shape. The scaffold routing, which is often performed manually, is typically a barrier for beginners to this field, especially for 3D structures. Hence, recent research in DNA origami design automation has focused on developing scaffold routing algorithms to facilitate a simpler design process[15–17], such as the A-trail routing and the spanning tree approach. However, those top-down methods, either semi-automated or fully-automated, are limited to specific types of static geometries. Specifically these are designed for wireframe structures, where all components have essentially the same geometry (combination of 1 and 2 helices[15], either all two helices[16], or recently 6-helix bundles[17]), and the algorithms focused on vertex design to enable wireframe structures without incorporating general capabilities for lattice or surface-based designs.

A primary advantage of our general design process is the ability to freely assign the geometry and connectivity of components with a variety of parameters for different components within the same design. To facilitate designs with such variety, we present a hybrid top-down and bottom-up approach for scaffold routing (Fig. S9-10). The process starts with pairing cylinders in every bundle component. Each pair is connected with external scaffold crossovers at the ends of components. Then, some of those cycles are combined based on user-defined joint connections to merge two cycles into one that permeates both pairs of cylinders across the two components. This results in a total of $N$ cycles (Fig. S11B).

According to the nature of double-crossovers, applying one double-crossover to the overall routing leads to either increasing (if the two ends of the crossover are located within one cycle, this breaks the cycle.) or decreasing (if the two ends of the crossover belong to two cycles, this causes merging of these two cycles.) the total number of cycles. Hence, exactly $(N-1)$ internal double-scaffold crossover connections (DSCCs) are required, all of which connect two different cycles, to integrate $N$ cycles into a single one. Here we introduce graph theory and the spanning tree algorithm to state this problem in a rigorous way. In this graph model, each node represents a cycle in Fig. S11B and each edge means the two cycles are neighbors and can be integrated by an internal DSCC based on the lattice rule[7] (Fig. S11C inset). The actual process of determining a suitable single cycle utilizes a nested loop where the outer loop finds a solution and the inner loop adjusts distances between crossovers, if necessary, while maintaining a single cycle. The third block in the outer loop (Fig. S9) examines the $N \times N$ adjacency matrix, and the spanning tree algorithm stochastically returns a list of $(N-1)$ edges, which leads to the integration of the overall routing. Then, for each edge located on the spanning tree, the algorithm stochastically selects an internal crossover from the multiple possible crossovers that could join those two cycles, thereby merging these two corresponding cycles (or two nodes). The selected crossovers are saved into an array. Finally, these internal crossovers in the array are applied one by one to integrate cycles, while the number of the total cycles continuously decreases until they are integrated into a single cycle.

This process may result in crossovers that are close together (closer than a user-defined threshold, default is 8 bases). In this case, the inner loop will run by identifying these closely spaced pairs of crossovers, removing one of them, and then stochastically finding a new crossover to maintain a single cycle. For very complex cases, the outer loop may result in a scaffold routing that may have many crossovers that are close together. In this case, to improve computational efficiency the algorithm re-runs the outer loop to find a new single cycle. This new cycle will be different due to the stochastic steps mentioned earlier, and this outer loop can be run until a single cycle is achieved that can efficiently proceed through the inner loop.

In addition, our scaffold algorithm allows users to specify certain options to accommodate different applications, such as the lengths of single-stranded scaffold in the connections at joints and the arrangement of crossovers at free ends to allow for polymerization of structures. Fig. S9 shows the flow of the scaffold algorithm to find a single cycle(s) (illustrated for single scaffold in Figs. S10 to S12 and multi-scaffold discussed later in Figs. S70 to S72 and S81).

In summary, to accompany with various geometry definition and assembly steps in our versatile design process, this scaffold algorithm can search the scaffold routing space on the a scale of seconds, including for more complex designs as in Fig. S12. The scaffold algorithm itself is a hybrid top-down and bottom-up approach, where steps like pairing helices during creating the assembly either from the line model or importing library components and identifying the spanning tree for the entire mechanism are top-down, and steps like defining DSCCs at joint connections, placing crossovers to integrate cycles, adjusting crossover positions, and defining nucleotide level details at joints or ends (e.g. ssDNA scaffold loops) are bottom-up. After being generalized for multi-scaffold designs, MagicDNA can expand design complexity including enabling larger structures without being limited to the use of conventional scaffold strands derived from M13 that are around 7000-8000 bases.

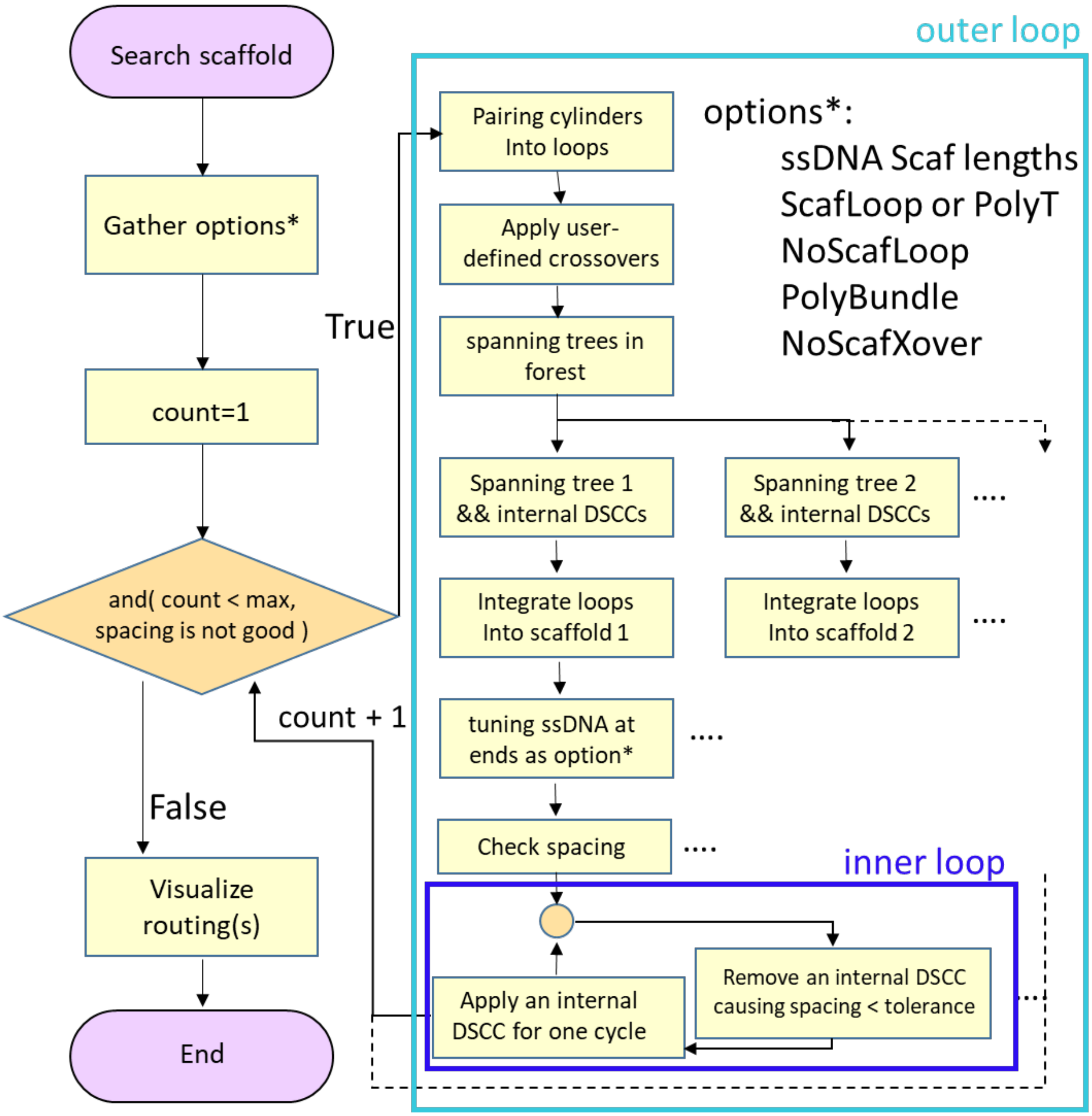

**Supplementary Figure 9: The flow of the scaffold algorithm.** The embedded algorithm is not only generalized for multi-scaffold designs but also compatible with user-specific options, such as single-stranded scaffold lengths, using scaffold loops or staple poly T overhangs to prevent base-stacking, shape-complementary stacking without scaffold loops, rounding up the scaffold loop to natural crossovers to allow for polymerizing the structure, and defining regions for no scaffold crossover in certain bundles for routing with reconfigurable components (e.g. as in the reconfigurable tetrahedron, Fig. 4A) or in multi-scaffold routing with defined interfaces between scaffolds.

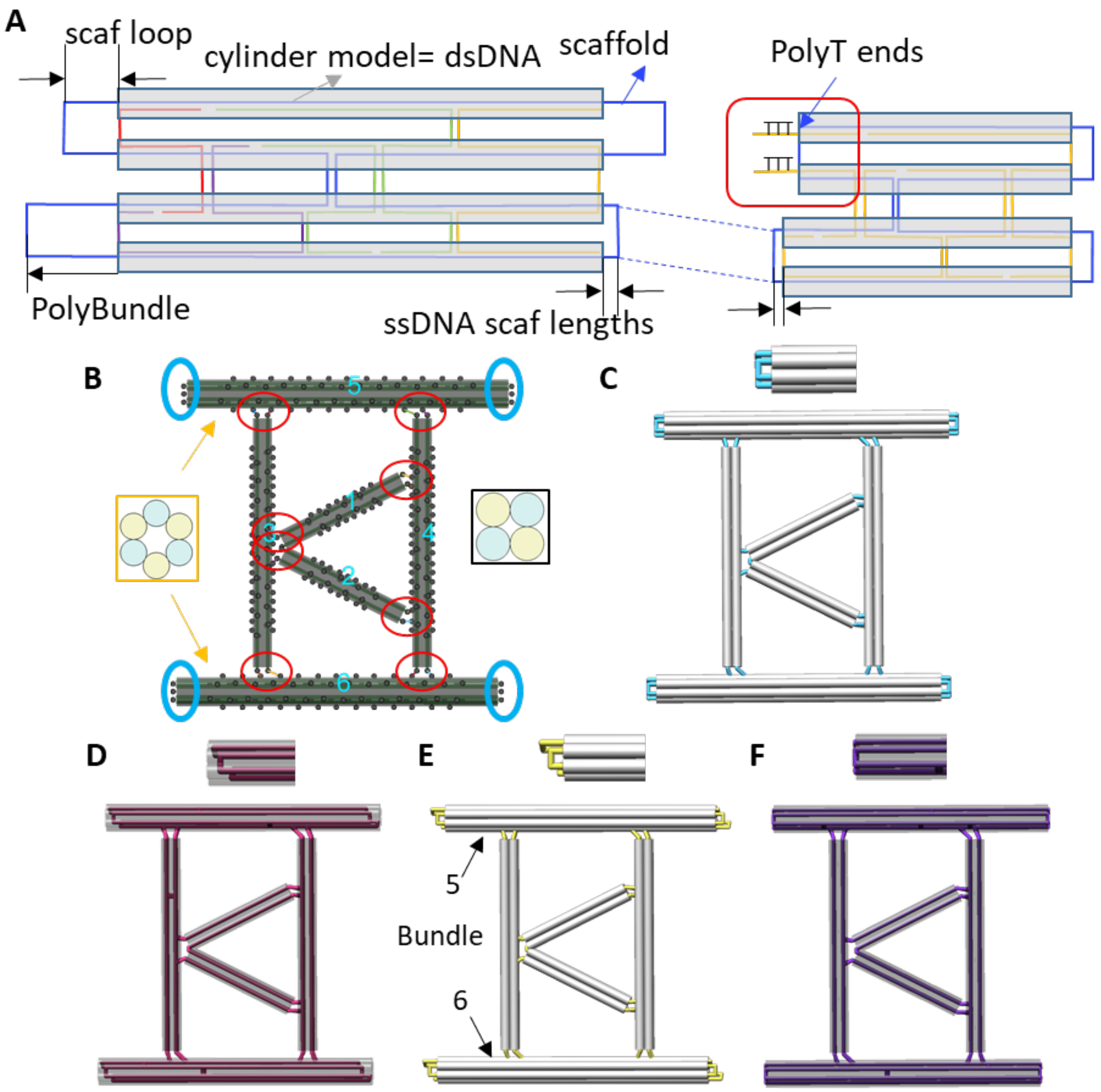

**Supplementary Figure 10: Trimming the routing at ends for user-specific options.** (A)(B) Schematic of extending the single-stranded scaffold routing beyond both ends of bundles. For connected nodes (red circles in (B)), the single-stranded scaffold is achieved by extending the scaffold routing of the cylinders on the ends. In addition, one of the most commonly used approaches to prevent base-stacking aggregation is to extend the scaffold routing from the cylinder model to leave scaffold loops (panel A, left) at open ends (blue circles in (B) indicate open ends). Another approach is to extend the staples from the cylinder model with single-stranded polyT (panel A, yellow strands on right) and shrinking the scaffold routing to the closest natural crossover position according to the helical orientation of the scaffold. For polymerization of structures, the open ends of selected bundles (labeled PolyBundle) are the interfaces between monomers where the scaffold helical direction needs to be considered and extended to the location of a natural crossover position. (C) The default setting for scaffold routing with single-stranded connections and single-stranded loops of uniform length for open ends. (D) In the case of polyT end overhangs, the scaffold is retracted from the cylinder model to the nearest natural crossover position. (E) Assign bundles 5 and 6 as polymerization bundles where the scaffold routing (yellow) is rounded up to the natural crossover positions. (F) Specify that bundles 5 and 6 do not have any scaffold loops.

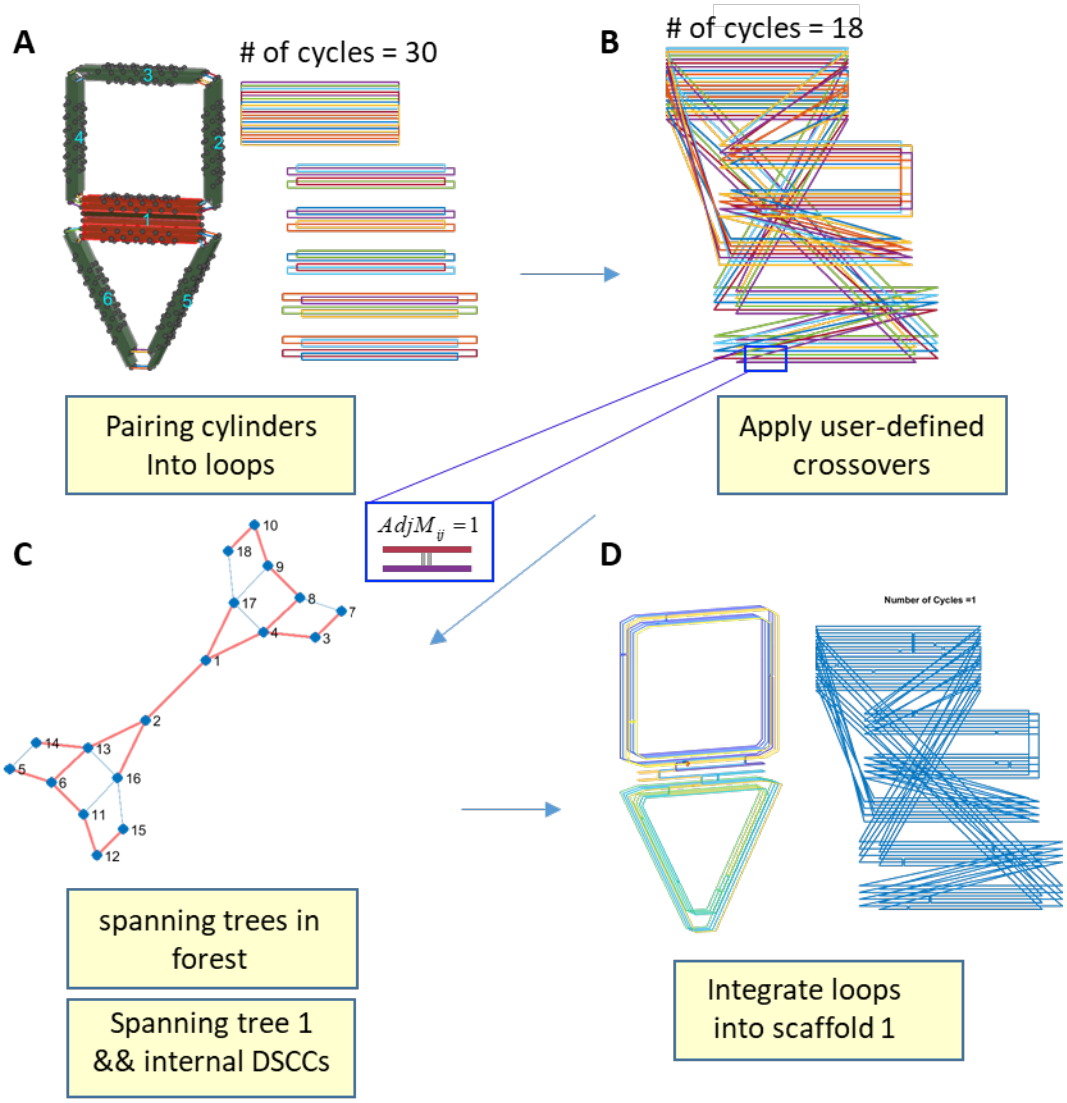

**Supplementary Figure 11: The scaffold algorithm for single-scaffold case.** (A) Pairing the cylinders within individual components. (B) Double-crossovers are applied at joint connections between components that are specified from the assembly GUI to form a total of N cycles. (C) Next, we construct the adjacency matrix where cycles are considered adjacent if there are any available internal double-scaffold crossover connections (DSCCs) that can connect the two cycles. The spanning tree algorithm assists with looking for N-1 internal crossovers to form one single-scaffold cycle that routes through the entire structure. (D) After applying those internal crossovers, the scaffold routing is visualized in both 2D and 3D representations.

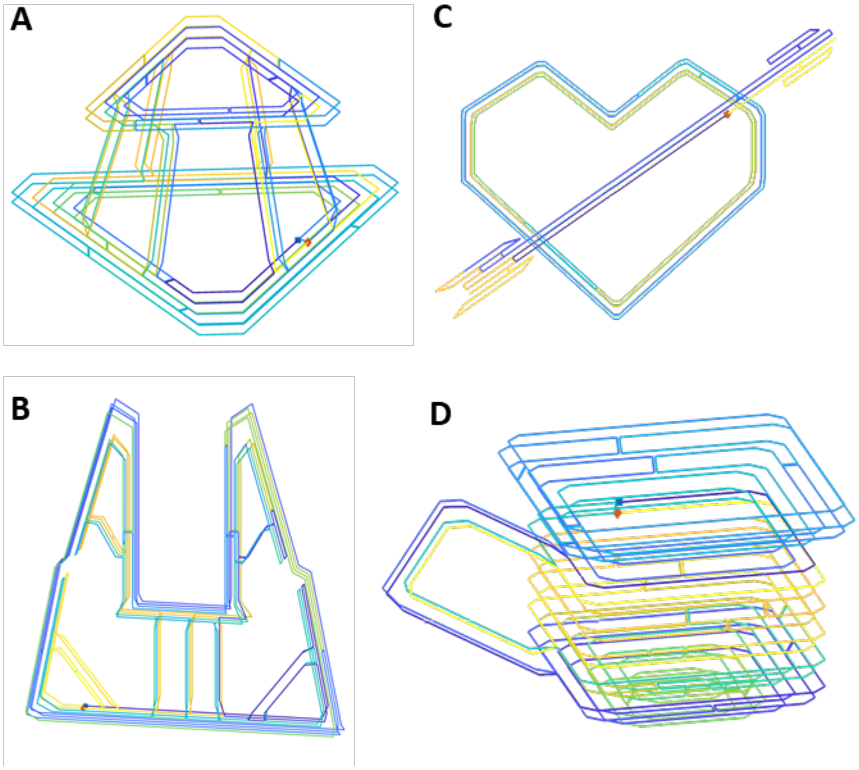

**Supplementary Figure 12: More examples of 3D scaffold routings.** The scaffold routing is color-coded along the length with blue at the virtual 5' end and yellow at the virtual 3' end. Since it is a circular strand, this follows the beginning of the sequence as typically defined in DNA origami[1]. This representation of the scaffold routing is used throughout this study, especially for single-scaffold routing for illustrating the way the scaffold weaves through the 3D structure. (A) Stewart Platform. (B) Gripper. (C) Cupid heart (D) Mug.

Staple routing algorithm

In contrast to the scaffold routing which must permeate the entire assembly, the staple routing can occur at the local component level since generally staples remain within a single component. Fig. S13 shows the flow of the staple algorithm, which is divided into two cases: without (Fig. S14) or with overhangs (Fig. S15). First, without overhang design, the algorithm creates the initial staple strands as a set of straight strands that follow along all the helices in the cylinder model with alternating directionality (5' to 3'). Initially, all possible staple crossovers between two neighboring helices (every 21 bps for honeycomb lattice and every 32 bps for square lattice) at appropriate helical positions are applied, except the staple crossovers within 10 bps of a scaffold crossover and connecting the same two helices (Fig. S14B). This process generally creates several very long staple strands that may be closed loops in the middle of 3D multi-layer components. We developed a heuristic algorithm for breaking long staple strands into appropriate lengths as a suggested option for users (Fig. S14C). In addition, based on the previous studies[2,18,19], the fraying at bundle ends is attributed as one of the major factors that affect the end-to-end lengths of bundles from oxDNA simulations. Hence, we also provide the option to apply single staple strand crossovers at the ends of bundles. We also provide an option to connect a staple across two bundle components if the length of the single-stranded scaffold at that connection is set to zero. This option was introduced as we extended MagicDNA designs to wireframe and surface-based structures. Without also using staples to connect components together in these cases, we observed higher values of RMSF at the interface of the two bundles. Details of vertex design will be discussed further subsequently (Fig. S31).

In addition to binding to the scaffold to form the double-stranded core of the structure, some staples strands can contain overhangs, which are commonly used to control the configuration of dynamic structures or used to attach objects of interest. We exploit our intuitive visualization to allow users to specify the locations of the overhangs directly on the 3D structure. Similar to the connection nodes in the assembly GUI, we define sites for possible overhangs on the surface as location where nicks in the staple strands would be oriented at the surface. These are represented as black nodes for users to select and specify the pairing for the case of actuation where a single strand should bind to two overhangs (Fig. S15B). To avoid the extremely short duplex sections around the overhangs, the spacing between overhang site nodes is every 32 bps in square-lattice and 21 bps in honeycomb lattice, instead of roughly 10 or 11 bps in the assembly GUI for scaffold connections. The placement of an overhang creates a nick with a sequence that extends out of the structure. These are similar to staple crossovers if there were an extra layer of virtual cylinders surrounding the cylinder model (a similar approach is often used to manually place overhangs in caDNAno). Once users specify the overhang locations and related parameters, the algorithm first incorporates the nicks at the desired locations, and then follows the normal process to specify staple routings (applying crossovers and breaking), and finally the overhangs are extended from the nicks (either 5' or 3' end) to virtual cylinders with the user-specified overhang lengths (Fig. S15 C).

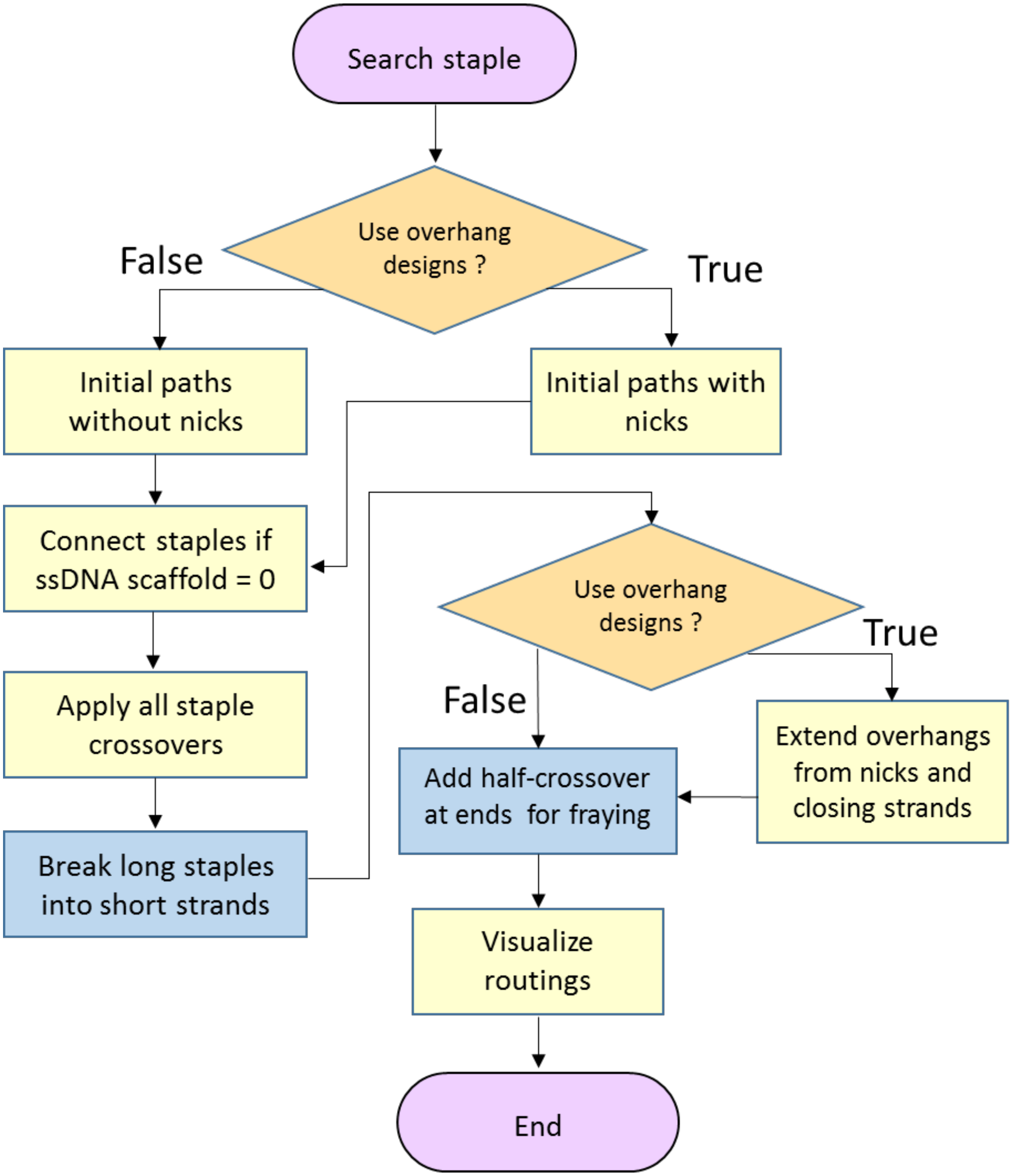

**Supplementary Figure 13: The flow of the staple algorithm.** MagicDNA offers options (blue boxes) for users to specify whether or not to execute some functions such as breaking the long staples and adding staple crossovers at ends to reduce fraying. We also created a 3D overhang design GUI as an option for actuating dynamic devices, assembling multiple structures, or functionalizing devices.

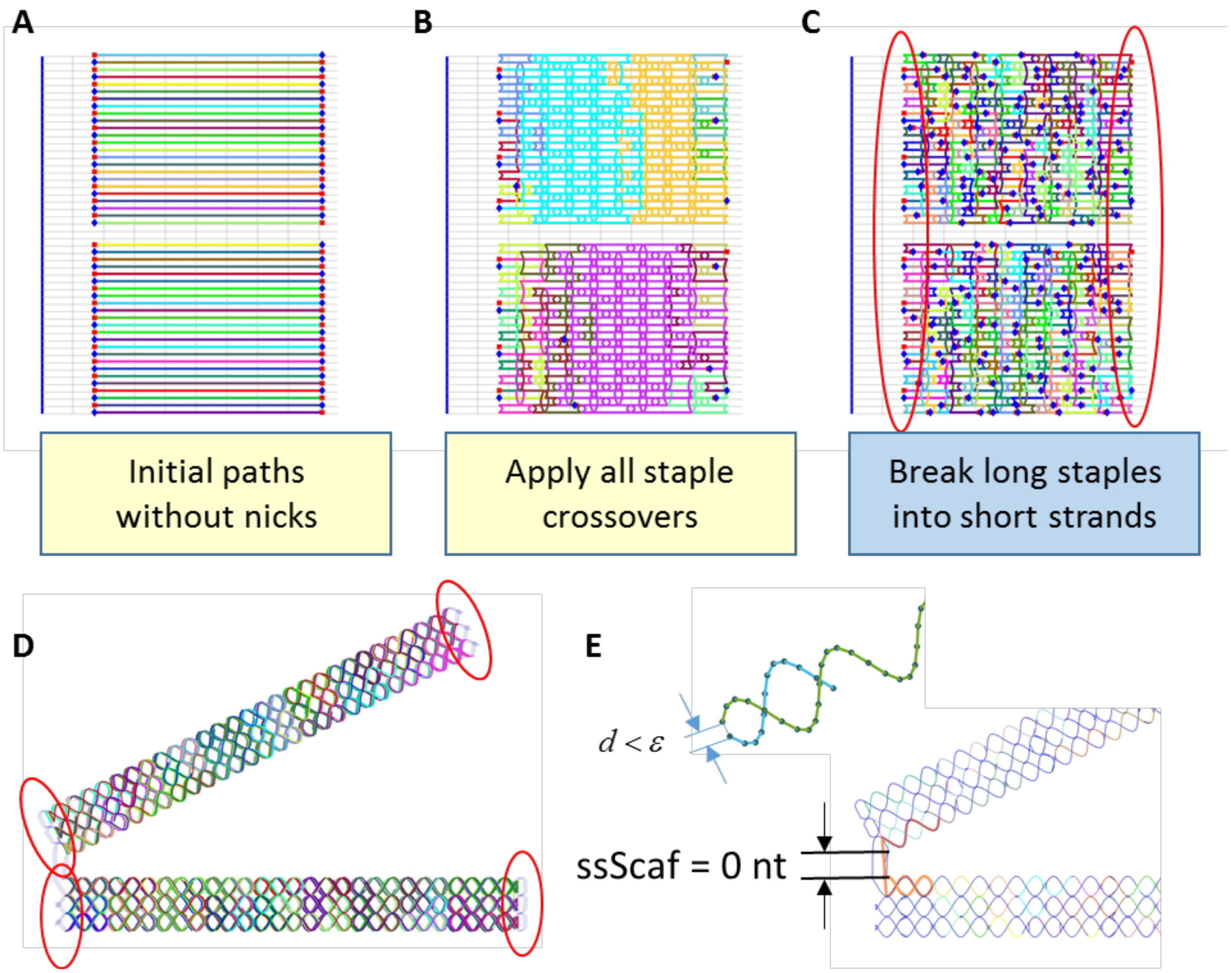

**Supplementary Figure 14: Staple routing algorithm without overhangs.** (A)The initial staple strands are based on the cylinder model as linear paths. (B) Then, all staple crossovers are applied following typical lattice rules, similar to caDNAno[1]. (C) A heuristic algorithm for breaking the long staples into target range, avoiding the bases within 3-nt of the crossovers[20]. (D)(E) Two possible adaptations on the staple algorithm. 1) To reduce the fraying at ends, users can choose to have staple crossovers at the ends if the 5' and 3' end orientations of two staple strands allow. 2) For wireframe and surface based structures, staples are connected between multiple components for more rigid interfacing of the two bundles if the single-stranded scaffold lengths are set to zeros.

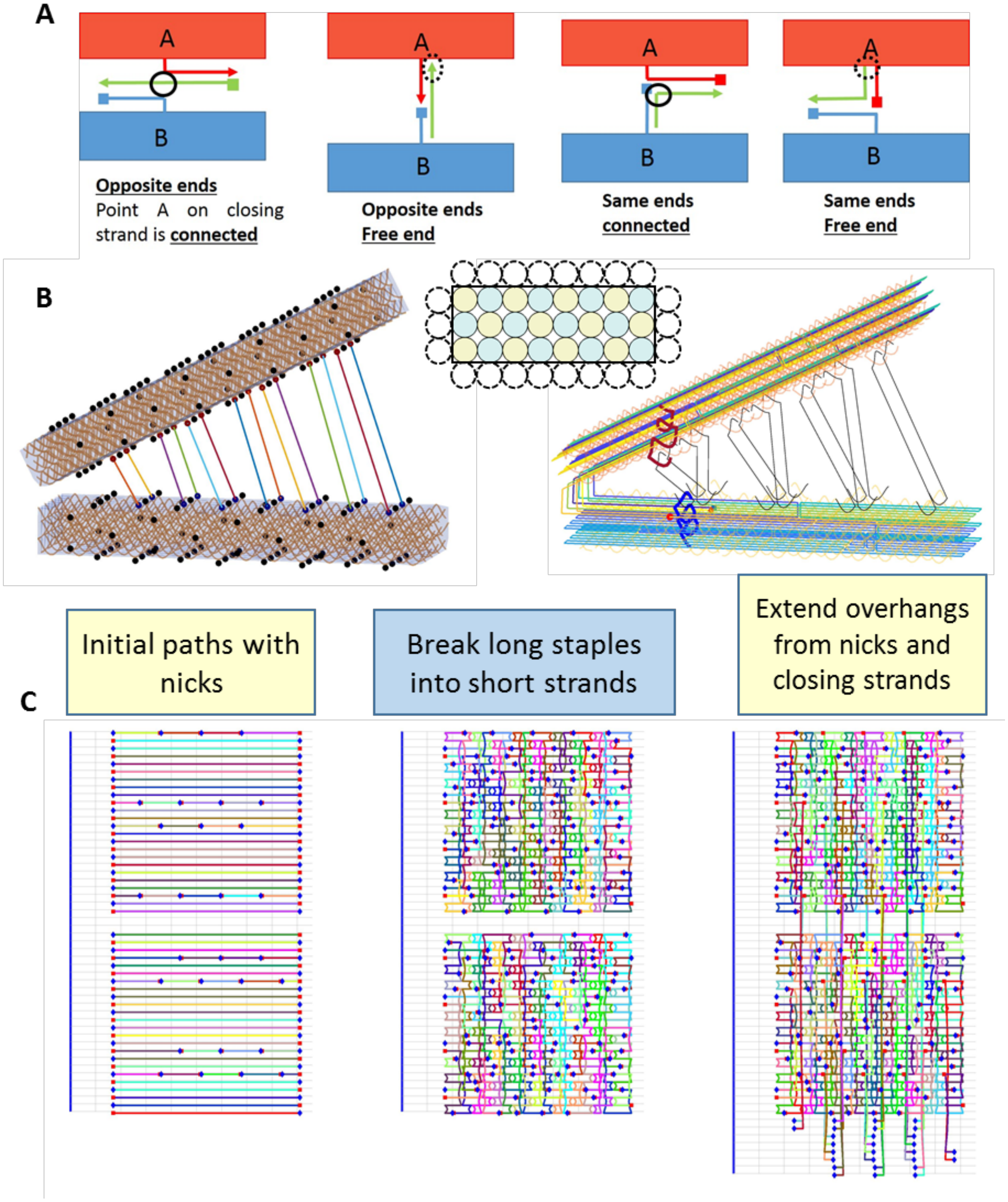

**Supplementary Figure 15: Staple routing with designing overhangs in 3D model.** (A) Schematics of several options for designing a pair of overhangs and its closing strand for actuation. (B) (Left) Similar to the assembly GUI, here we visualize each bundle with locations where staple helical orientations are at an appropriate position to add an overhang (black dots). Once two sites are chosen by the user and the desired parameters for the overhangs (3' or 5' end, length, type from options in panel (A) are specified in a table, the staple routing algorithm will take this into account and automate the staple routing with overhangs and also generate the corresponding closing strands. (C) The locations of the selected nodes will be input as nicks in the initial staple strand routing. After applying crossovers and breaking, those locations are then extended either from 5' end or 3' end at the nick with desired lengths. Alternatively, a single overhang site for functionalization can be added by specifying one of the two strands in the pair to have zero length. In this case, no nick will be placed at the site where the overhang length is defined as zero.

Interface with caDNAno and coarse-grained models

Interfacing with other DNA origami tools provides options for users to fine tune designs and access useful features in other open-source computational tools in the field. For example, the caDNAno software is well-known for its user-friendly interface to locally adjust the scaffold or staple routing. The multi-component 3D design feature in MagicDNA creates stacked 2D routing diagrams that can be exported as .JSON filed that can be opened in caDNAno. The many component 3D designs can result in complex 2D routing diagrams that are difficult to interpret. To address this problem, we created a GUI within MagicDNA that mimics the caDNAno path diagram that is juxtaposed with a 3D scaffold and staple routings for reference (Fig. S16). Hence, the desired location for modification can be located and selected in the 3D structure, and the corresponding location is highlighted in the 2D diagram. For example, a desired location for a chemical modification, such as biotin or a fluorophore, can be located in the 3D structure, and the corresponding location is highlighted in the 2D diagram in MagicDNA so it can be easily located in caDNAno to identify the staple strand for modification at the correct site. In general, by clicking on either 3D or 2D routing, MagicDNA shows the mapping locations simultaneously in the 2D and 3D depictions as well as the index of the caDNAno cylinder and position. In addition, one can also focus on a specific staple by clicking on that staple in the table with the complete staple list to highlight its locations in the 2D and 3D diagrams in MagicDNA.

Although our goal is to facilitate the routing of the complex structures with the scaffold and staple algorithms, there are cases where specific design features do not fit within our scaffold or staple algorithms. Thus, for those specific cases, an alternative solution is to provide initial nearly-complete routings in MagicDNA, which satisfy the user inputs at the component and assembly levels, and then use the interface with caDNAno to finalize the routing details at the nucleotide level. Some possible modifications in caDNAno are shown in Fig. S17A and B, including adding forced connections, applying or cancelling crossovers, shifting nicks, and non-lattice-based staple routing (e.g. to form loops in the scaffold, Fig. S17B bottom). Once those modifications are made, one can import the modified .JSON file back to MagicDNA, maintaining the updated routing (Fig. S17C), except for insertion and skip data in .JSON files. Importing the designs with updated routings back into MagicDNA is useful to leverage the 3D visualization to inspect modifications like forced-connections made in caDNAno. Another use for re-importing caDNAno designs into MagicDNA is for running oxDNA simulation, which can use the initial conformation directly from the 3D structure in MagicDNA, which likely requires some rigid body motions of components to achieve a desired configuration when converting a caDNAno .JSON file to oxDNA formats[2,21]. Also, MagicDNA allows specification and simulation of hybrid-lattice designs, which are not currently possible to design directly in caDNAno or any other software. Integration with caDNAno also allows potential use of other simulation tools like cando[7,19] and COSM[22], which take caDNAno files as input.

The interface with oxDNA can be described in two key steps: 1) exporting the topology and configuration file and 2) visualizing and analyzing the trajectory results from the simulations. First, (Fig. S21 A) the configuration file contains the positions and the orientations of all bases which can be calculated by superposing the helical representation and the cylinder model (or line passing through the center of the cylinder) representation in Fig. S16C. For the topology file, the sequences are output from MagicDNA in the sequence and diagram GUI after specifying the scaffold sequence. The connectivity between bases is also obtained from the routing in MagicDNA. Second, after the simulation is completed, one can visualize each configuration in the trajectory directly in a visualization GUI. For higher quality visualizations, one can also export the configuration as ribbon or coarse-grained model as an .BILD file to open in UCSF Chimera[6] (Fig. S21B left), or using another oxDNA visualization tool, cogli1 (Fig. S21B right). We also integrated basic analysis tools like RMSD and RMSF after aligning all configurations with the first configuration to remove global motions (Fig. S21C). In brief, the RMSD is the average spatial deviation for each base using the initial configuration as the reference along the time axis, and the RMSF is the level of the temporal fluctuation from the average configuration at each base. A useful visualization of the simulation is the average configuration color-coded with the RMSF values to present the trajectory with geometry and local flexibility in one single visualization (Fig. S21D). Regions of higher RMSF suggest more flexibility, which may be intentional or unintentional from design perspective. In addition, well-defined double-stranded bases align as clearly helical structures since the averaging all the configurations cancels out the thermal fluctuations. On the other hand, the single-stranded region and fraying double-stranded regions often appear as a series of spheres with smaller helical radii in the average configuration because of the significant thermal fluctuations.

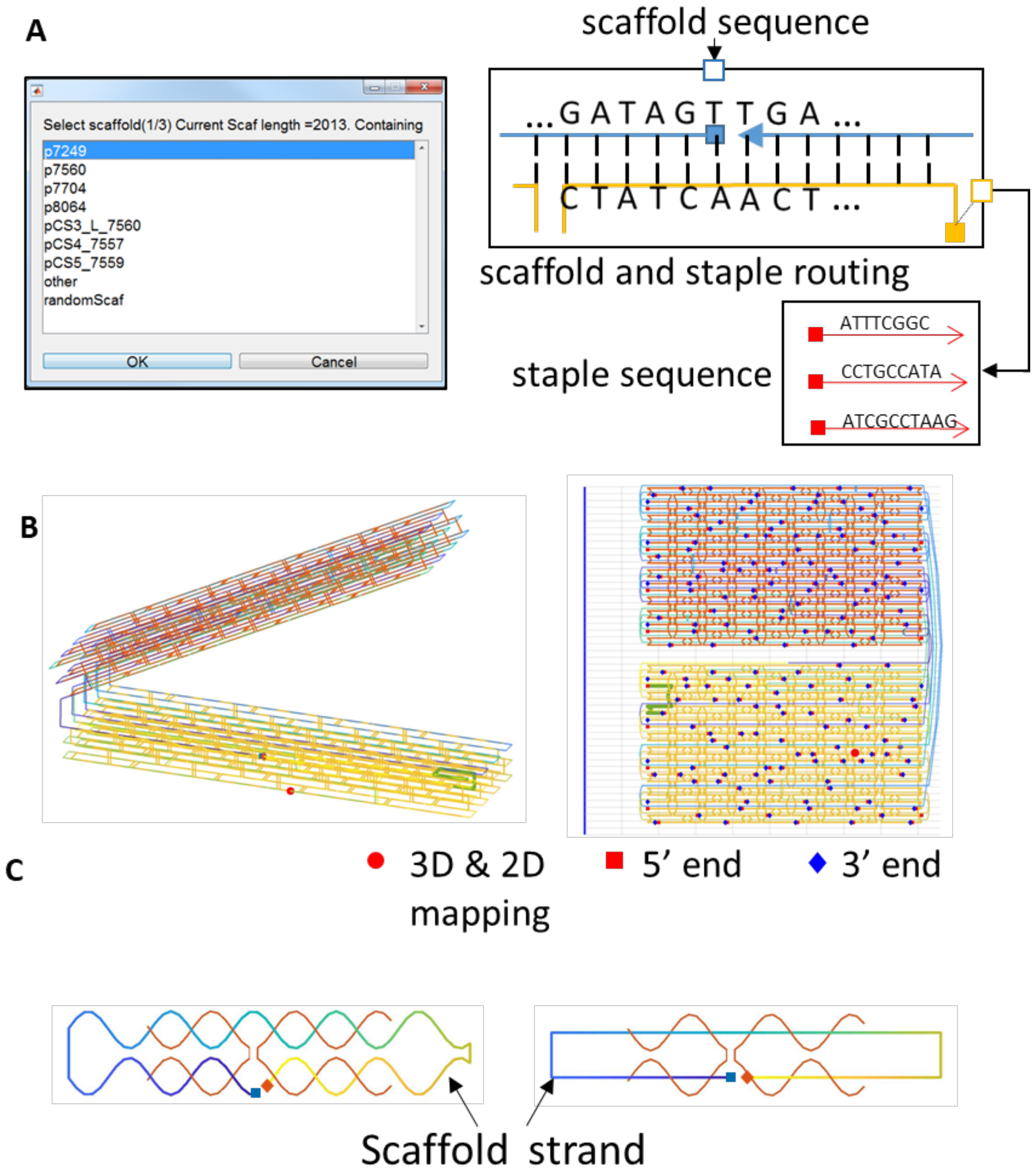

**Supplementary Figure 16: The sequence and design diagram panel GUI (I).** (A) Schematic illustrating the generation of the staple sequence list after specifying the scaffold sequence or a custom sequence. (B) (Left) The 3D visualization and (Right) its 2D design diagram of scaffold and staple routings. Here users can directly export the staple sequences or .JSON files for finer modifications in caDNAno. (C) Both 3D scaffold and staple routings can be switched between the chicken-wire representation (helices shown as lines) and helical representation in MagicDNA.

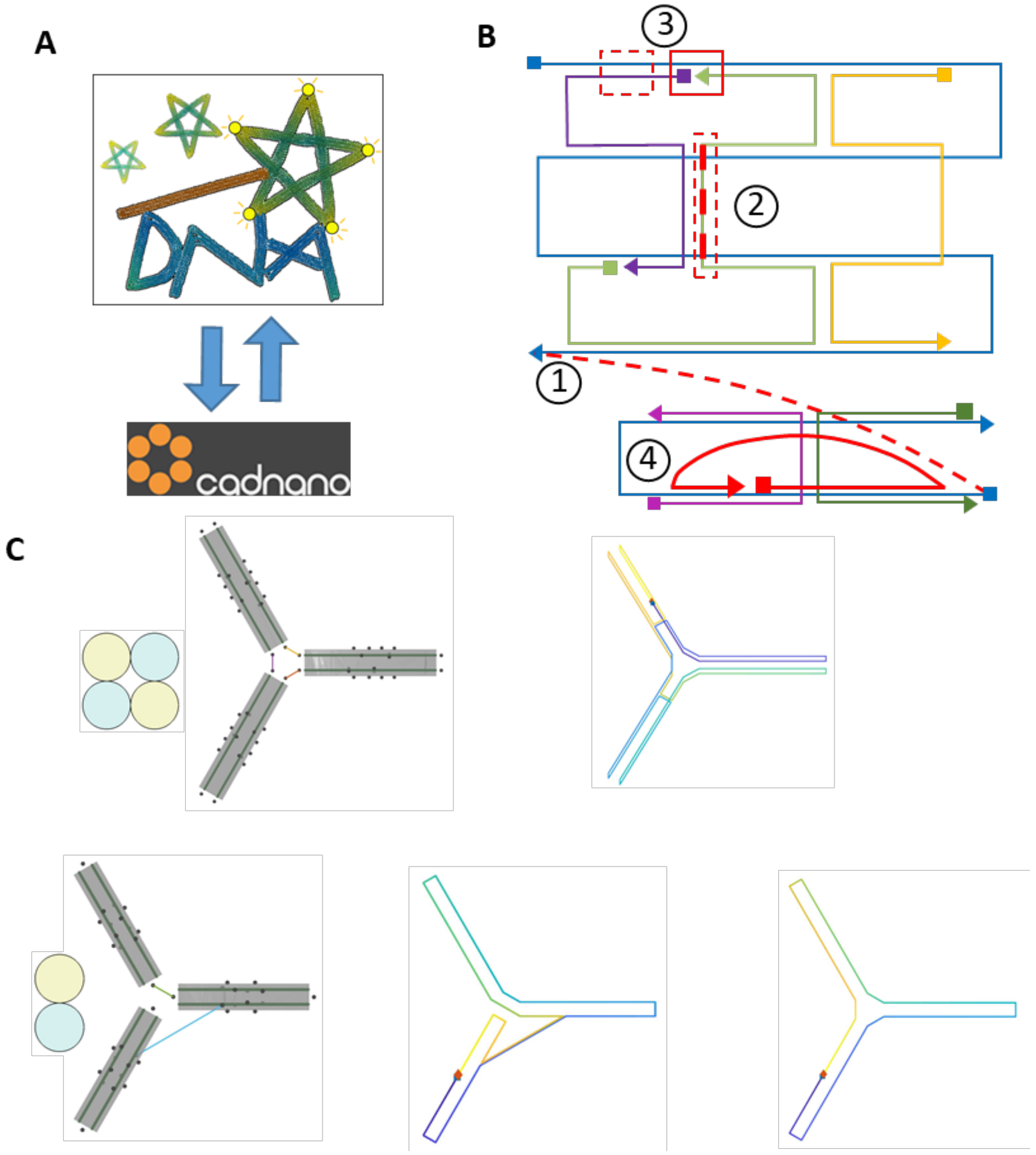

**Supplementary Figure 17: Interface and modifying design details in the caDNAno software.** (A) Schematic of two-way interfacing with caDNAno software. (B) Examples of editing in caDNAno for fully controlling the routing at more bottom level, such as forced connecting strands, adding/removing a crossover, shifting the nick position, and non-lattice reverse staple routing for multi-configuration devices. (C) Currently MagicDNA does not support the ability to connect one single node in a bundle to multiple nodes in other bundles. Hence, designing a 3-way junction with our scaffold algorithm requires at least two end nodes, or at least 4 helices. (Top) A planar 3-way junction is feasible for cross-sections with even numbers of pairs of cylinders (i.e. even numbers of end connecting nodes), such as 2×2 cross-sections. As an alternative approach, users can design a side-to-side connection, which can be moved to the junction by tuning the routing in caDNAno. (Bottom) With the single node at ends near the junction, the scaffold algorithm cannot find a scaffold routing for this joint. Here we introduced a side-to-side connection, and modification in caDNAno was used to shift the connection to the vertex. The updated routing can be imported back into MagicDNA for visualization as shown in the right.

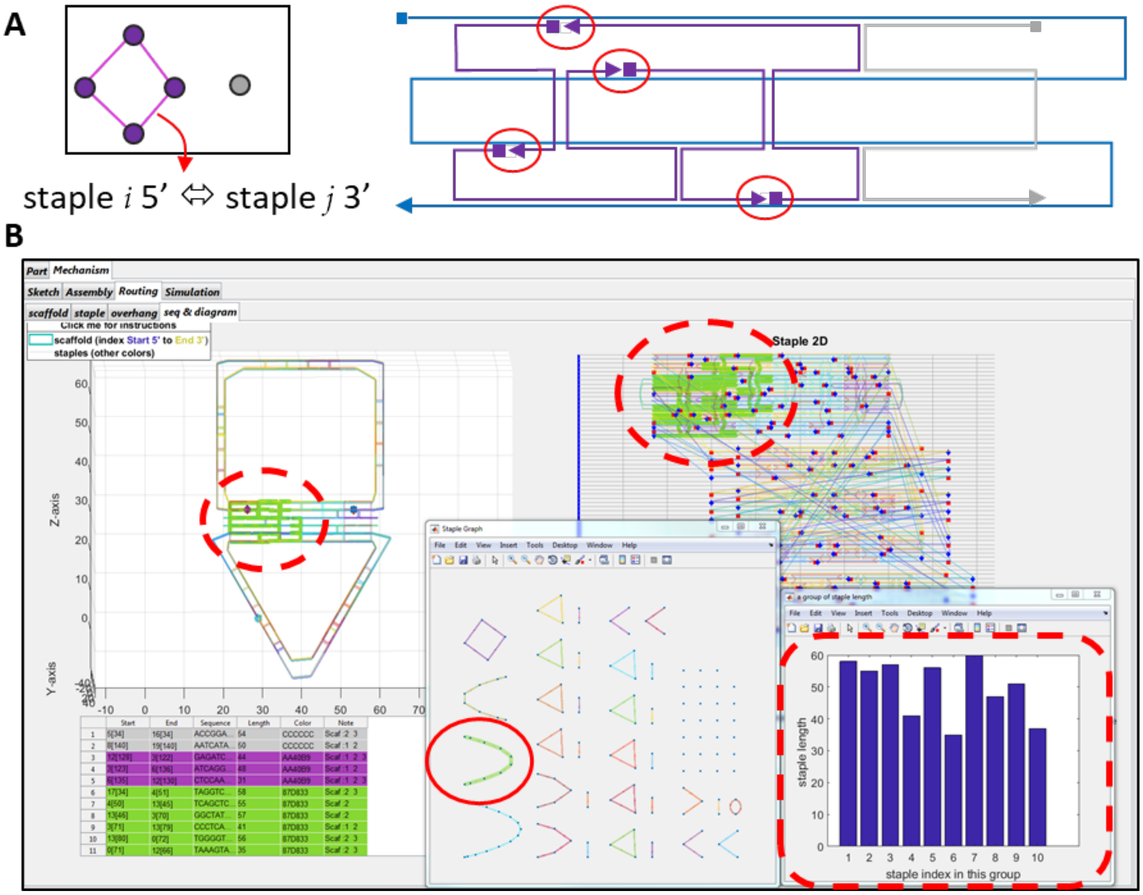

**Supplementary Figure 18: The sequence and design diagram panel UI (II):** the staple graph for adjusting staple lengths. Our staple algorithm for breaking staples is a heuristic process which may not be ideal for some specific cases. For example, prior work demonstrated folding of a concentric slider joint requires making the staple lengths longer and removing crossovers in the inner track to cause the track to fold earlier than the outer tube during annealing[23,24]. In order to inspect the staple length distribution more efficiently, this GUI was created to allow users to shift nicks to adjust two neighboring staples. (A) Schematic of the staple graph where a staple is a node and two staples are considered adjacent if their 5' and 3' ends are adjacent to each other at a nick. Staples near ends may be isolated and are shown in gray in the staple list. (B) Depending on the adjacency of staple strands, the staple graph shows a series of grouped staples as open chains or closed loops without branches. Clicking on a group triggers the functions to indicate the staple 2D and 3D locations in the routing diagram. The length distribution of this group also is shown in the lower right corner.

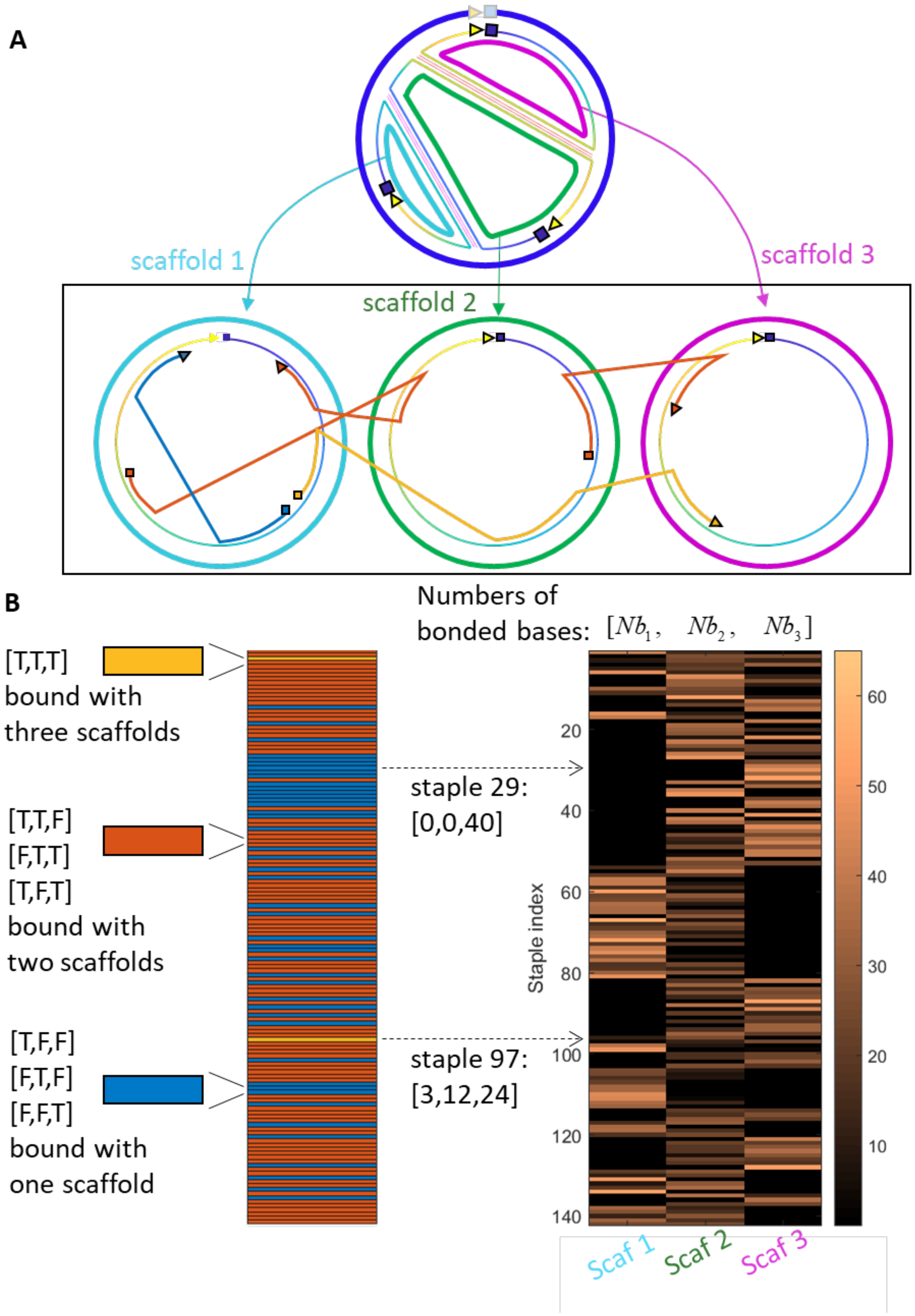

**Supplementary Figure 19: The sequence and design diagram panel GUI (III): the inspection function.** For multi-scaffold cases, the staple strands are designed to bind with one or multiple scaffolds in piece-wise manner. In other words, some staples may remain locally bound within one scaffold (e.g. blue staple strand in bottom left of panel (A)). Other staples may be bind to multiple sections on different scaffolds (e.g. orange and yellow strands in bottom panel of (A)). (B) From the perspective of the staples in this three-scaffold case, we can categorize staples into three cases depending on the number of scaffolds they bind to (either one, two, or all three). The mapping of complementary bases between staple and scaffold strands is depicted as a 2D matrix and visualized in the right for all staples in the design. In addition to the graphical representation, the GUIs in (A) and (B) are intended to show the quantitative numbers of complementary bases and staple locations on scaffolds.

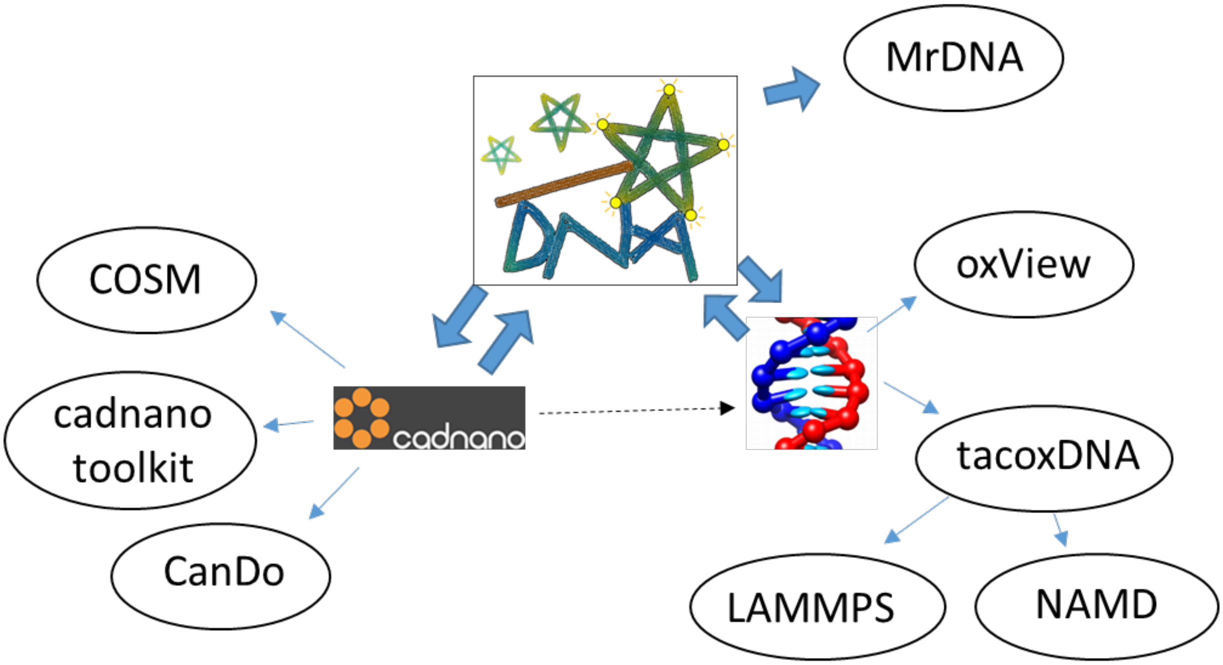

**Supplementary Figure 20: Interfaces with caDNAno and oxDNA, and extension to other computational tools in DNA nanotechnology.**

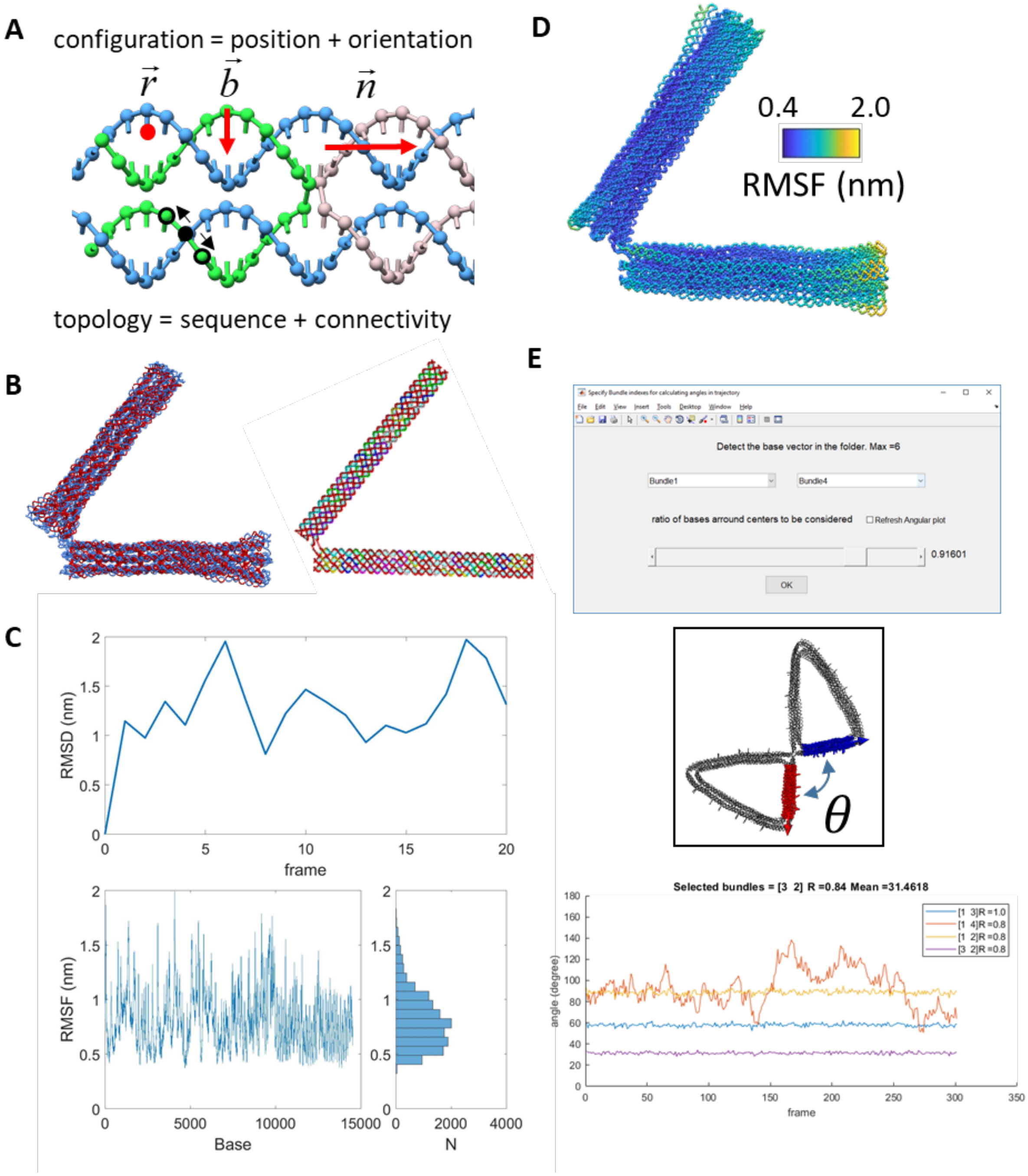

**Supplementary Figure 21: The oxDNA interface for exchanging data with oxDNA for preparing coarse-grained simulations and post simulation visualization and analysis.** (A) The schematic of the configuration and topology files for preparing oxDNA simulations. MagicDNA directly exports the 3D configuration with base orientations to the configuration file. The topology file saves the sequence information from the sequence and design diagram panel GUI (Fig. S16) and the connectivity of bases and strands. Here we also generate a force file with mutual traps between pairs of complementary scaffold and staple bases for relaxation purposes. (B) Visualization tools for oxDNA trajectory results are integrated into MagicDNA, or it is possible to export files for visualization in UCSF Chimera (Left) and cogli1 (Right). (C) Tool for calculating the RMSD and RMSF are included in MagicDNA package. (D) The average configuration over the entire trajectory file is frequently used to show double-helical DNA origami structures, color-coded with the RMSF values at the base level. (E) To extract and calculate the angle between bundles in the trajectory, we create a GUI to specify the bundle indices which are used to identify the indices of the bases in oxDNA model.

According to Sharma et al[25] the splaying at the ends may not reflect the angle between components. Hence, a slider is created for excluding bases from the ends when calculating the bundle direction. For those bases taken into account (red and blue in inset), the normal versor (unit vector)[4,5] property is the direction of the helixes. After flipping the directions of half of the bases (i.e. due to antiparallel binding of dsDNA), the average direction of these bases is considered as the bundle direction and the angle between components can be calculated for each frame in the trajectory.

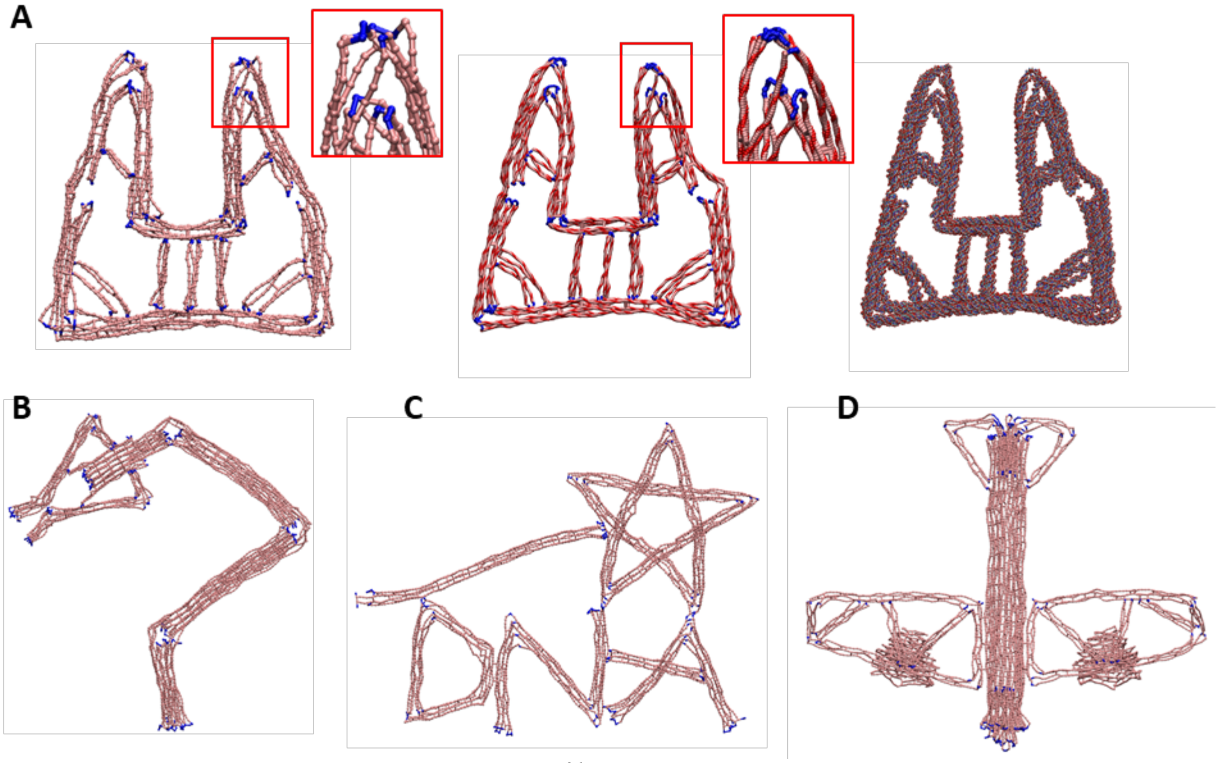

**Supplementary Figure 22: Interfaces with MrDNA[26], a multi-resolution coarse-grained model**. (A) From left to right, the resolutions of the gripper design increasing from 5-bp/bead, to 2-bp/bead, and to an atomic model. (B) Coarse-resolution models of multi-scaffold structures: the robotic manipulator with the tweezer end-effector, the MagicDNA logo, and the airplane design.

#### Supplementary Section 2: Top-down parametric design for functional devices

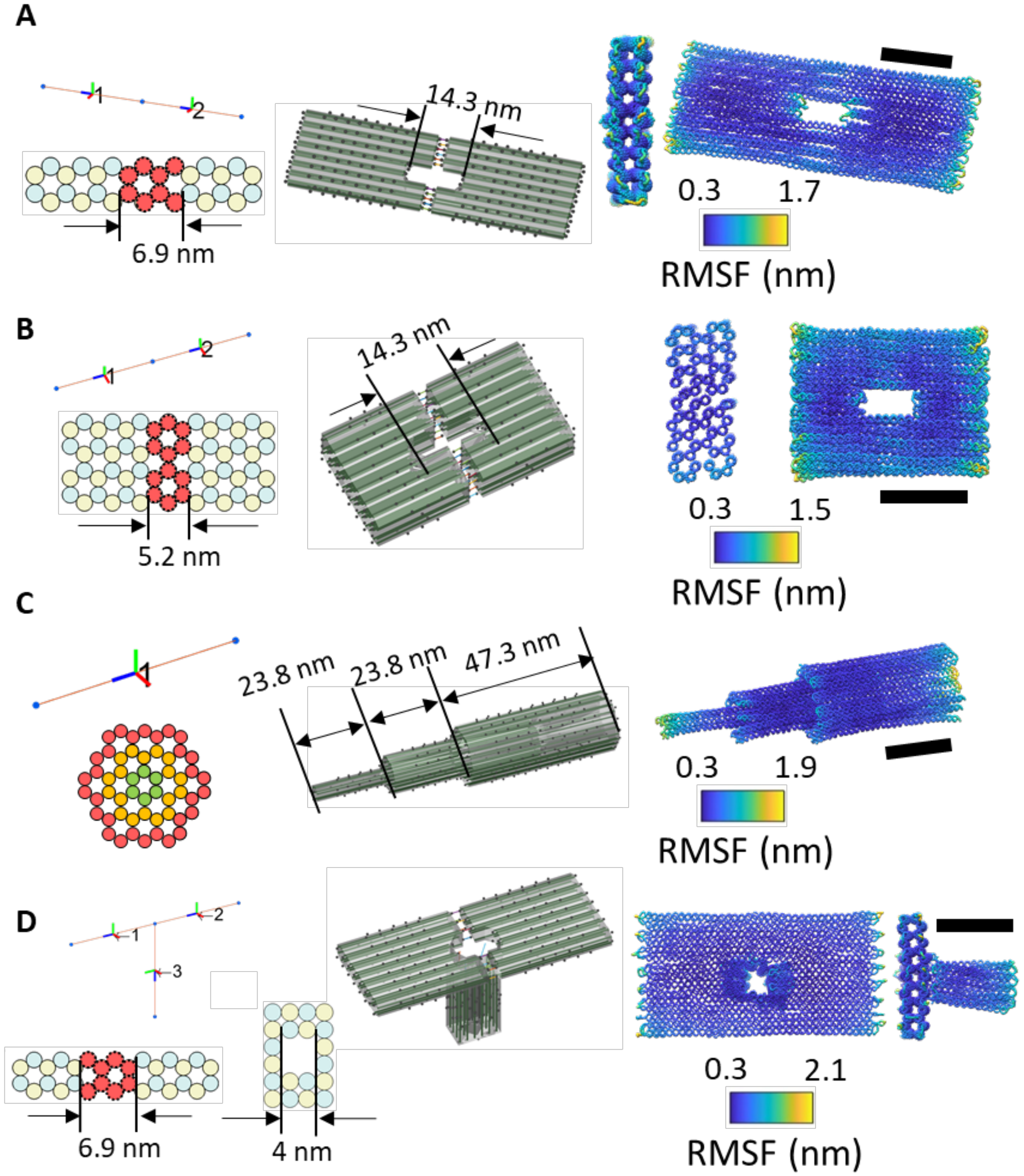

**Supplementary Figure 23: Top-down parametric design of the nanopores.** (A) To create a hole in the middle of a horizontal plate, we used 2 bundles and shrunk certain cylinders in each bundle. The number of connections between the two bundles is equal to the number of unshrunk cylinders which connect to the other bundle without any single-stranded scaffold bases. Hence, the staple algorithm also forms continuous staples at the interface to create a continuous plate without concentrated nicks at the interface. (B) Another example of a horizontally oriented nanopore which has a deeper pore due to more layers. (C) A vertical nanopore made by one bundle with cylinders in each layer shifted to corresponding positions. (D) A combination of both horizontal and vertical versions of a nanopore. This combined nanopore was designed with three bundles where the vertical portion has a different lattice for the cross-section (cross-sections depicted at left). Scale bars = 20 nm.

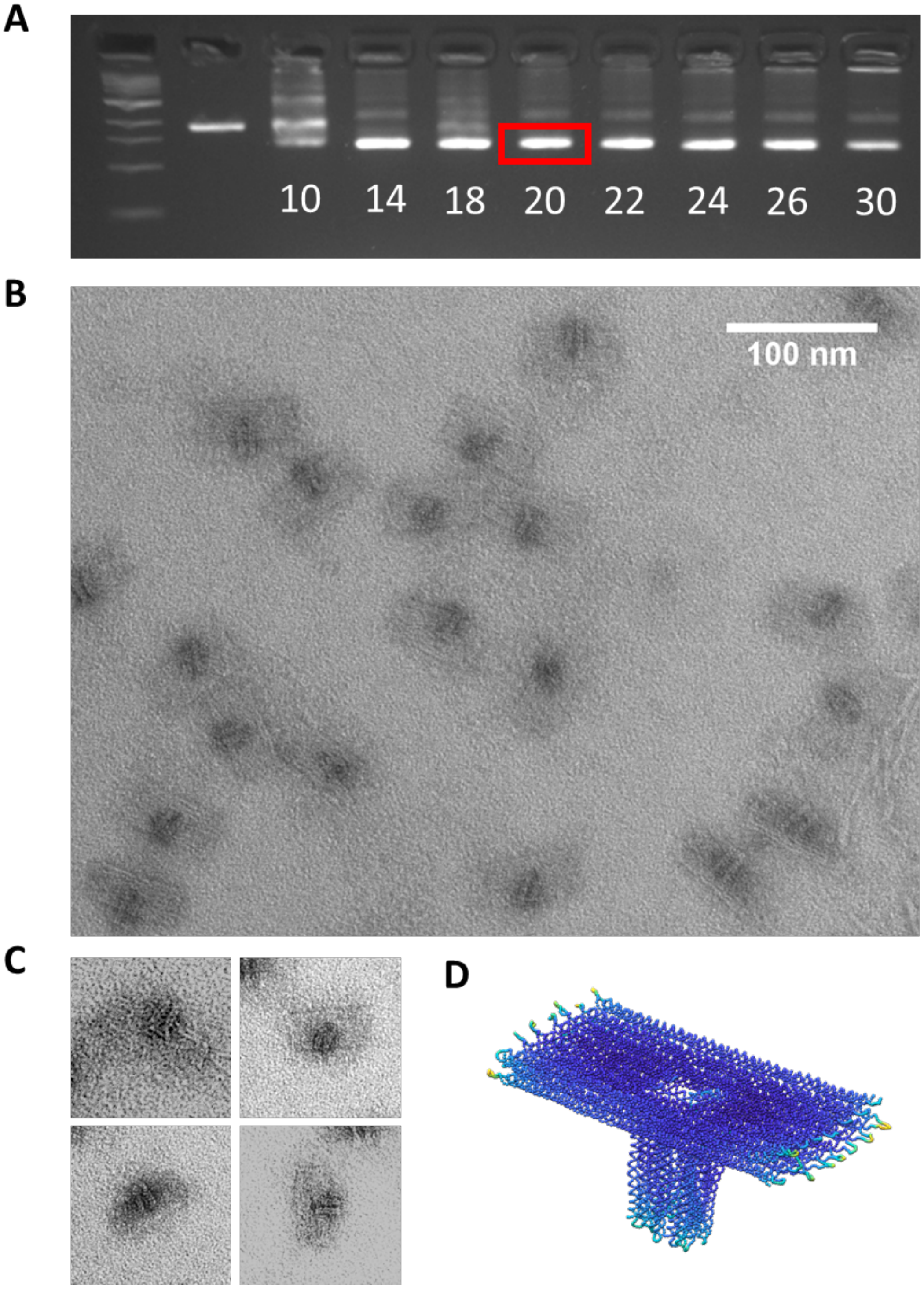

**Supplementary Figure 24: Experiments of the nanopore structure**. (A) Nanopore structure folded using a two-and-a-half-day thermal annealing protocol while testing a range of magnesium chloride concentrations from 10-30 mM and purified using agarose gel electrophoresis (buffer conditions 0.5xTBE 11 mM $MgCl_2$). (B) Zoomed-out TEM of gel-purified nanopore structure folded at 20 mM $MgCl_2$. (C) Zoomed-in TEM images of gel-purified nanopore structure folded at 20 mM $MgCl_2$ (D) OxDNA simulation result of nanopore structure.

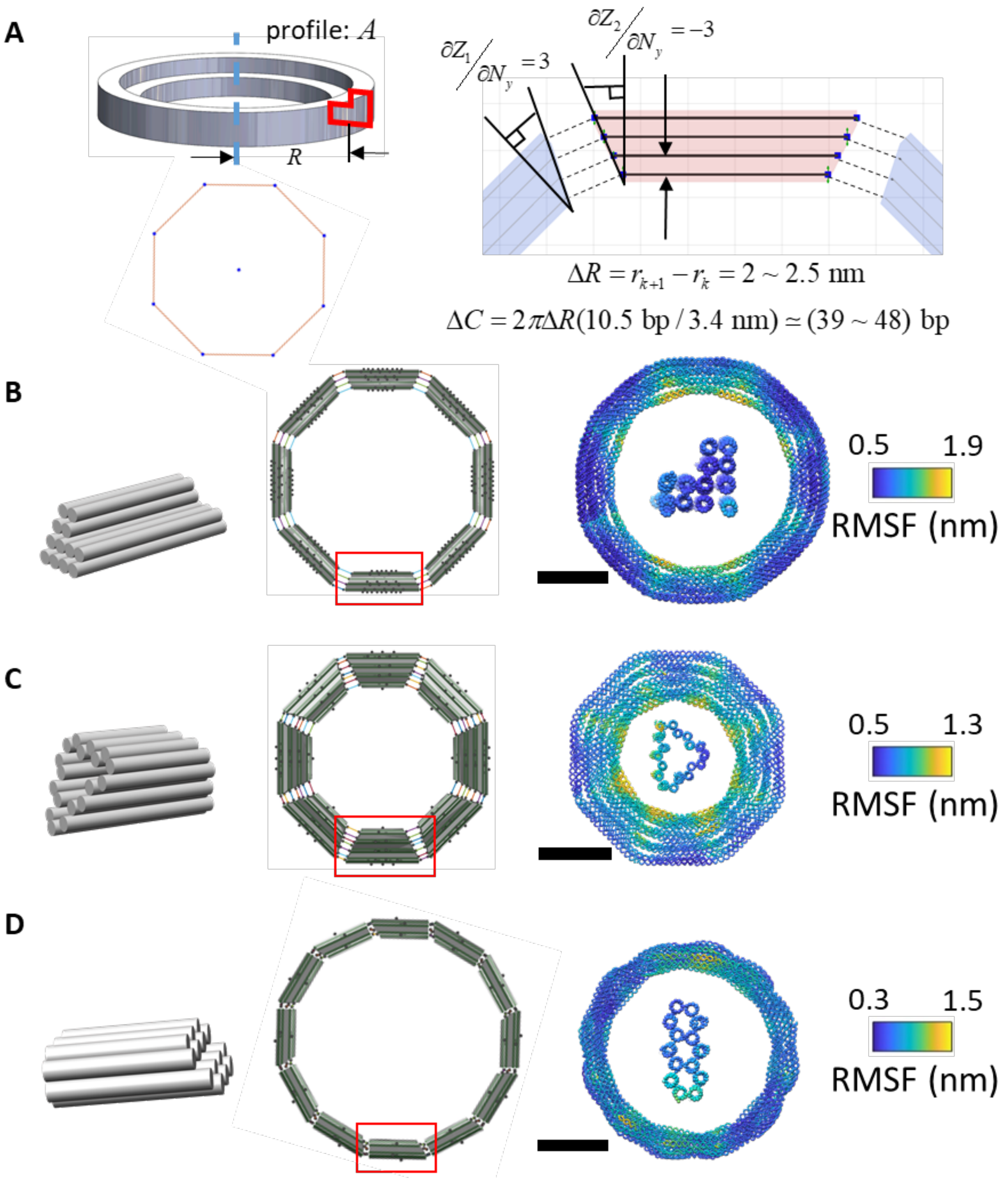

**Supplementary Figure 25: Top-down parametric design of rings.** (A) To approximate a ring, multiple components with gradients at their ends can be used to form a closed loop. For example, here we sketched 8 lines and converted them into 8 tilted bundles. The gradient values can be calculated according to DNA helical geometries[27]. Assuming the spacing between helices is 2.5 nm, the accumulated difference between any two in-plane layers should be 48 bps. Hence, for a ring made by 8 bundles, the gradient values at both ends for all bundles were assigned as ±3 (bp/layer) in the parametric table. (B) An example of a ring with 8 customized square-latticed bundles. (C) An example of ring with 8 shell-type (i.e. hollow) honeycomb bundles. (D) A ring made by 12 bundles with 14-HB honeycomb cross-sections. Scale bars = 20 nm.

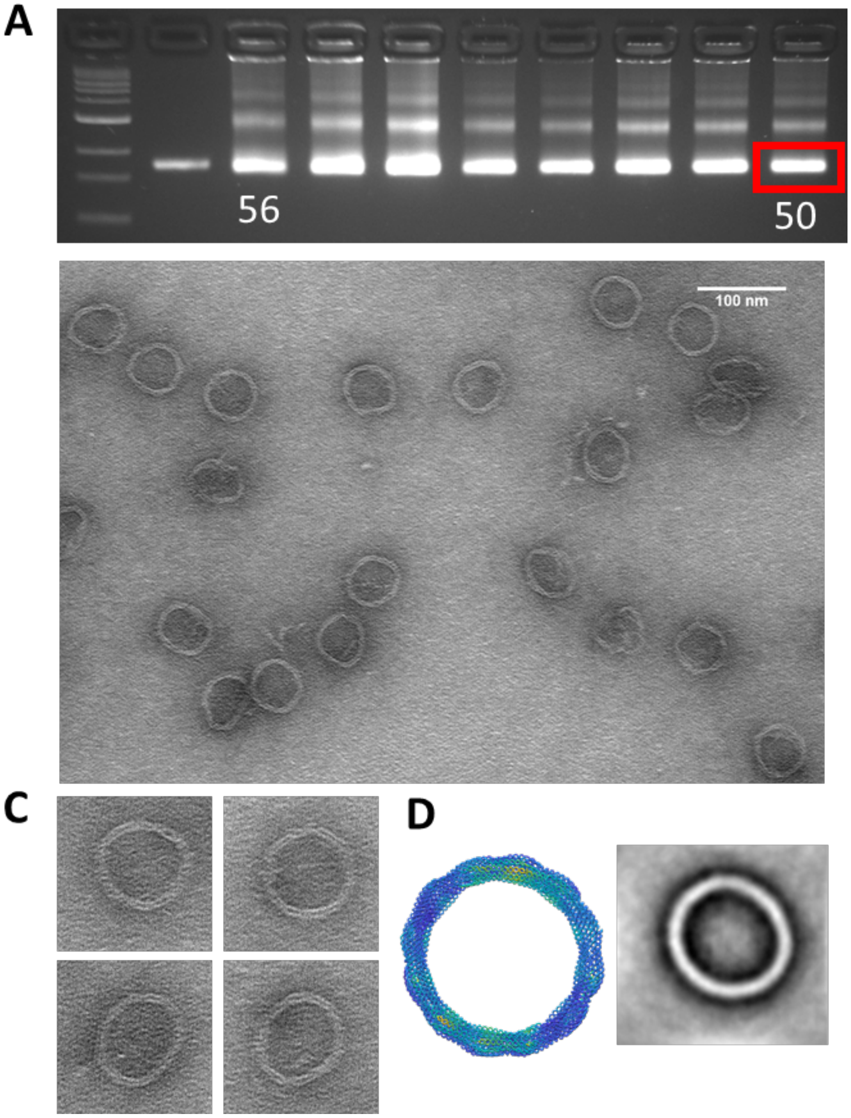

**Supplementary Figure 26: Experimental validation of the ring structure.** (A) The ring structure was folded using a 4-hour folding protocol testing a range of annealing temperatures from 56°C-50°C with 20 mM magnesium chloride concentration and purified using agarose gel electrophoresis (buffer conditions 0.5xTBE 11 mM $MgCl_2$). (B) Zoomed-out TEM image of gel-purified ring structures folded at 50°C with 20 mM $MgCl_2$. (C) Zoomed-in TEM images of gel-purified ring structure folded at 50°C with 20 mM $MgCl_2$. (D) OxDNA simulation result of ring structure (Left) and image average of gel-purified ring structure folded at 50°C with 20 mM $MgCl_2$ (Right).

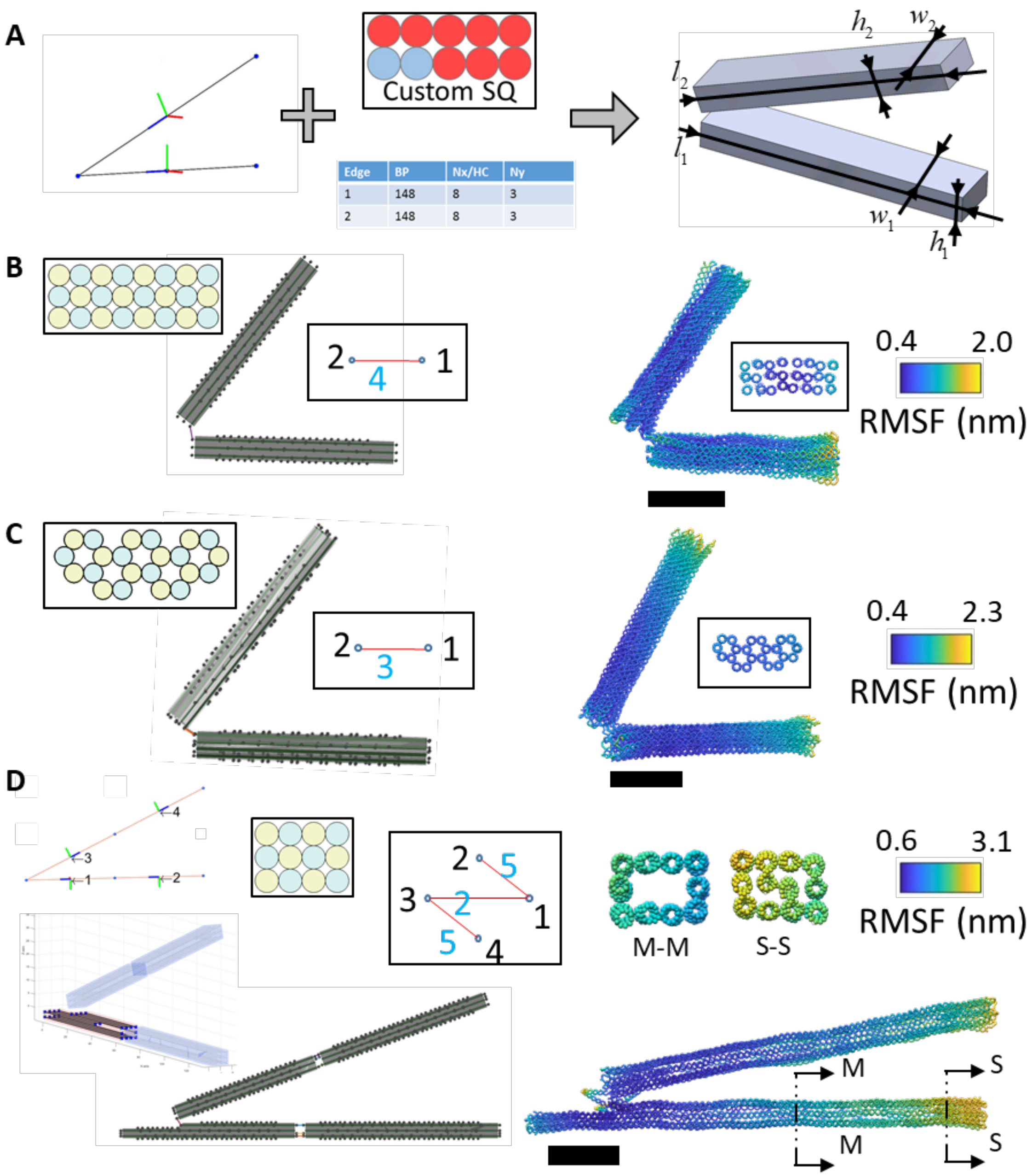

**Supplementary Figure 27: Top-down parametric design of the hinges.** (A) A hinge is formed by two arms. Thus, one can first sketch two lines in the MagicDNA line model interface and later specify the geometric parameters in a table to embody the lines, including real lengths in base pairs and cross-sections. (B) An example of hinge made by two 8×3 square-lattice arms using a sketch with two lines, connected at the inner layer with 4 external double scaffold crossovers. (C) Another example of hinge made by two 22-HB honeycomb-lattice arms and connected with 3 external double scaffold crossovers. (D) A hinge design with hollow cross-sections in the middle of the two arms. Here we used 4 bundles and the bundle editing GUI to adjust the cylinder lengths after converting. The center of the two cylinders of each bundle was shifted at one side while the other ends remained sealed. This hollow hinge was fabricated and validated by agarose gel electrophoresis and TEM. The hollow feature of the design is shown in the left and oxDNA simulation in the right. Scale bars = 20 nm.

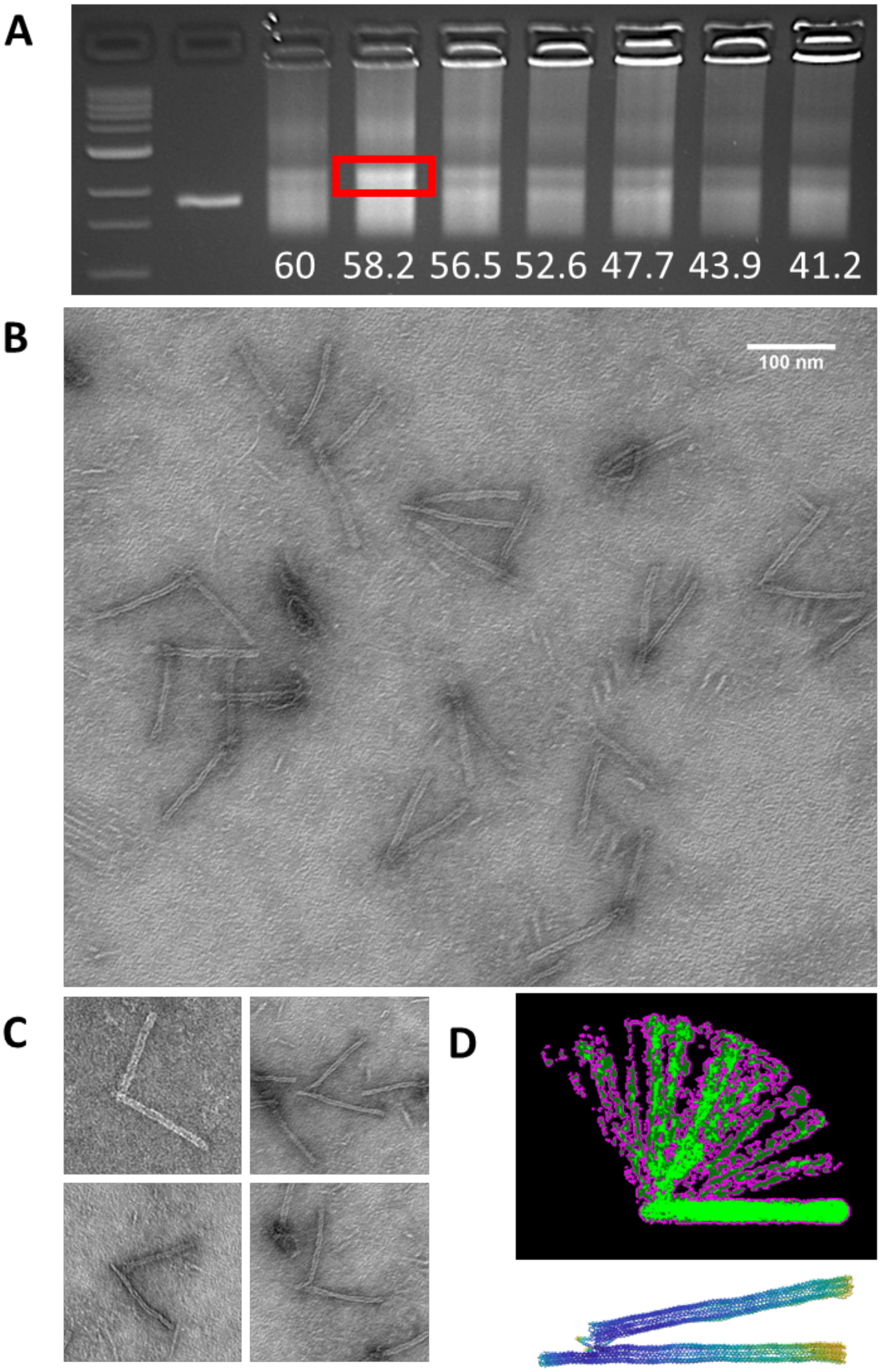

**Supplementary Figure 28: Experimental validation of the hollow hinge.** (A) Hollow hinge structures were folded using a 4-hour folding protocol testing a range of annealing temperatures from 60°C-41°C with 20 mM magnesium chloride concentration and purified using agarose gel electrophoresis (buffer conditions 0.5xTBE 11 mM $MgCl_2$. (B) TEM zoom-out of gel-purified hollow hinge structures folded at 58.2°C with 20 mM $MgCl_2$. (C) Single TEM images of gel-purified hollow hinge folded at 58.2°C with 20 mM $MgCl_2$. (D) The superposed hollow hinge TEM images of different configurations to show the range of motion and oxDNA simulation result.

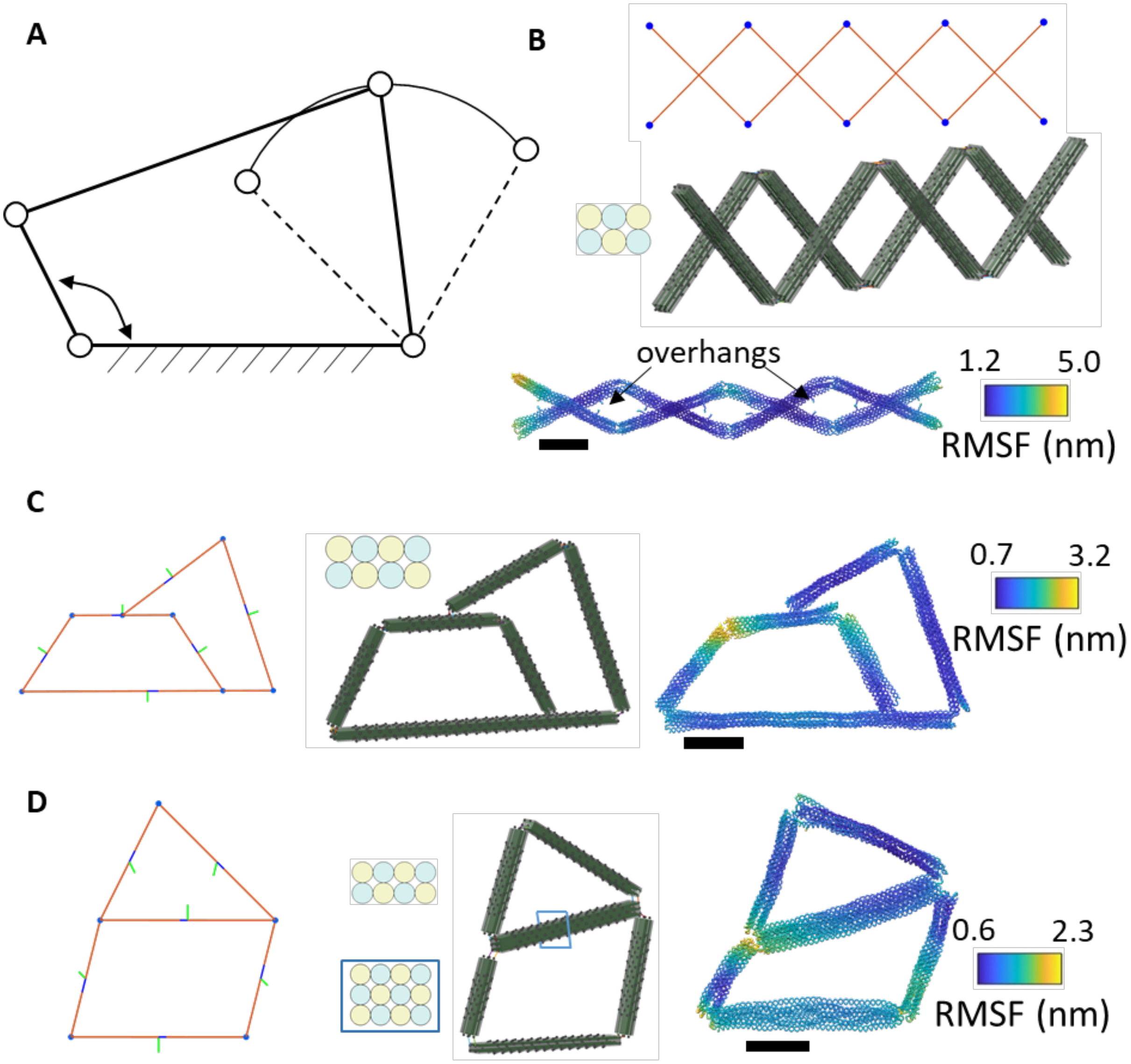

**Supplementary Figure 29: Top-down parametric design of the linkages.** (A) Schematic of a four-bar linkage connected with four revolute joints to create the desired motion. (B) An example of the linkage, embodied with a scissor mechanism, which is made of 8 3×2 square-lattice bundles. The connections between bundles include end-to-end connections like the hinge devices and side-to-side connections between the top and bottom layers. This design is similar to our previous work[28] and the overhang design tool was used to illustrate the placement of overhangs. (C) A six-bar mechanism made of six 4×2 square-lattice bundles. (D) Another four-bar mechanism with one bar formed by a triangular truss. Except for the link in the middle with 4×3 cross-sections, the other links are the bundles with 4×2 square-lattice cross-sections. Scale bars = 20 nm.

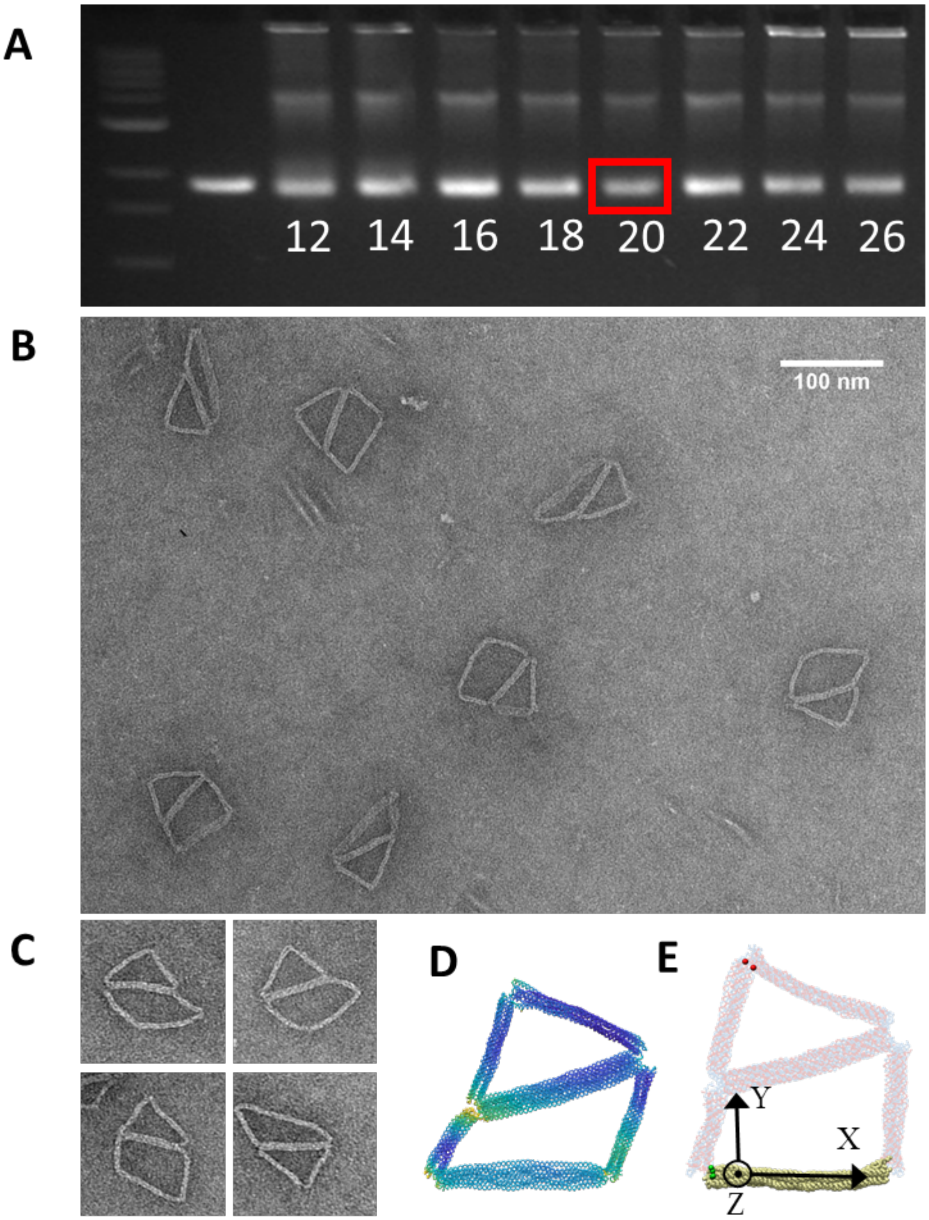

**Supplementary Figure 30: Experimental validations of the 4-bar mechanism.** (A) 4-bar mechanism structure was folded using a two-and-a-half-day thermal annealing protocol while testing a range of magnesium chloride concentrations from 12-26 mM and purified using agarose gel electrophoresis (buffer conditions 0.5xTBE 11 mM $MgCl_2$). (B) Zoomed-out TEM image of gel-purified 4-bar mechanism structure folded at 20 mM $MgCl_2$. (C) Zoomed-in TEM images of gel-purified 4-bar mechanism structure folded at 20 mM $MgCl_2$. (D) oxDNA average configuration with RMSF of 4-bar mechanism structure. (E) For extracting the tip position in the oxDNA trajectory, the local coordinates of each frame were defined such that the bases on the ground link were used to define the X direction by the normal versor (unit vector) similar to Fig. S21. Then the positions of all bases were used to calculate the out-of-plane Z direction using principle component analysis (PCA). Once the orientation was defined by X-Y-Z coordinates, the reference bases were assigned at the 2-nt hinge connection in the bottom-left corner (green spheres) to calculate the tip position (also on the 2-nt hinge connection, red spheres).

### Supplementary Section 3: Top-down iterative design for complex structures

#### Modular design process for the relation between metrics and design parameters

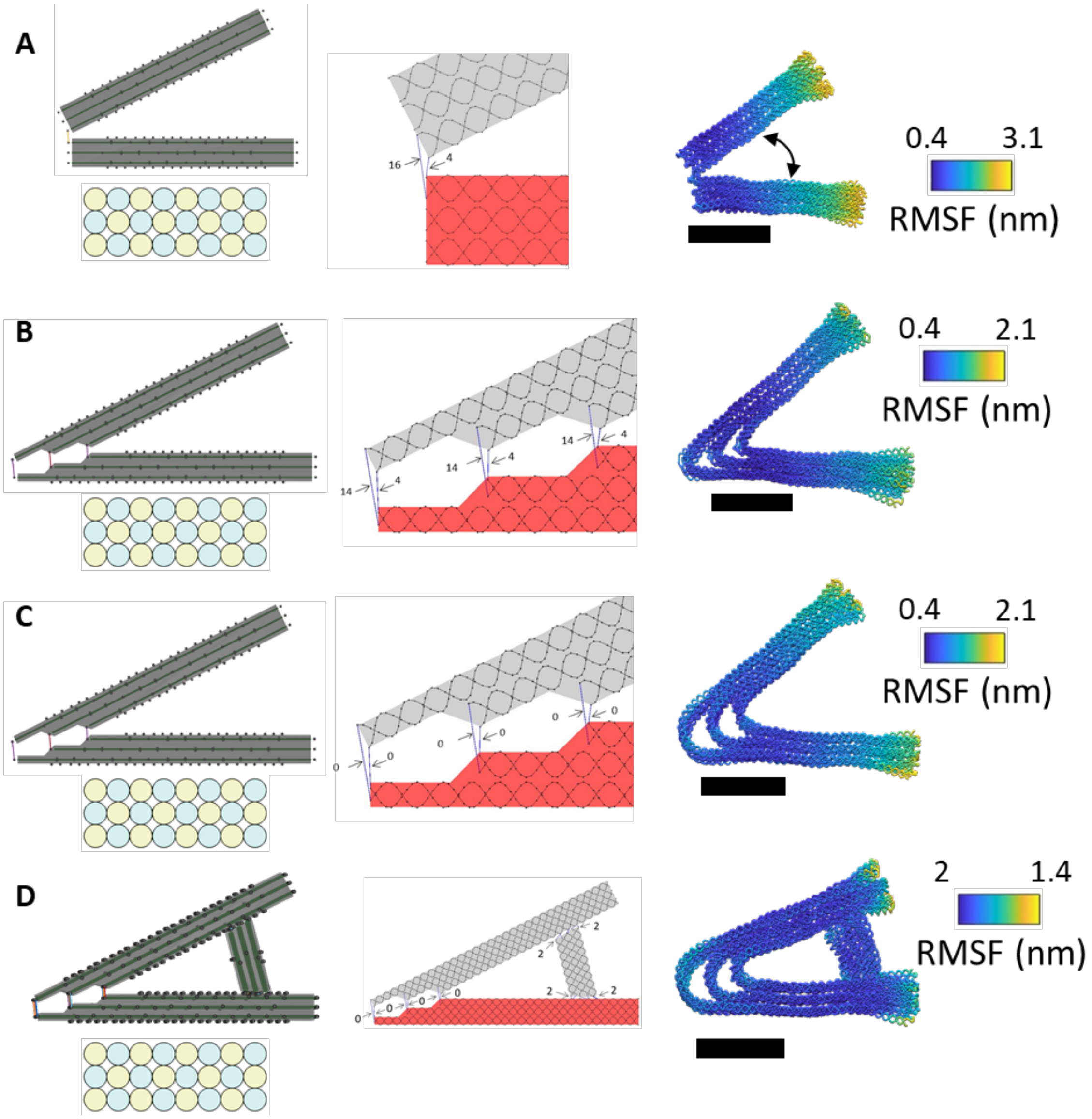

**Supplementary Figure 31: Joint design for static or dynamic cases.** oxDNA simulations reveal how the design of connections between bundles affects the mechanical properties of the nanostructure. (A) Typical design strategy of a dynamic hinge joint[2,23], where only the inner layer is connected to the other arm. Due to the helical orientation, the some connections are given a shorter ssDNA length to facilitate motion primarily along one rotational degree of freedom. (B)(C) To design a static joint of certain angle, the cylinder model should be adjusted to the corresponding lengths, according to different layers. If a sharp corner is desired, leaving some ssDNA on scaffold connections is recommended and the staple routing algorithm keeps the staple strands in two links separate. On the other hand, without ssDNA scaffold bases, the continuous double-stranded duplexes create a round corner at the vertex. (D) To further enhance the stiffness of the joint, adding another bundle as a truss at a location away from the vertex can constrain the motion more precisely. Scale bars = 20 nm.

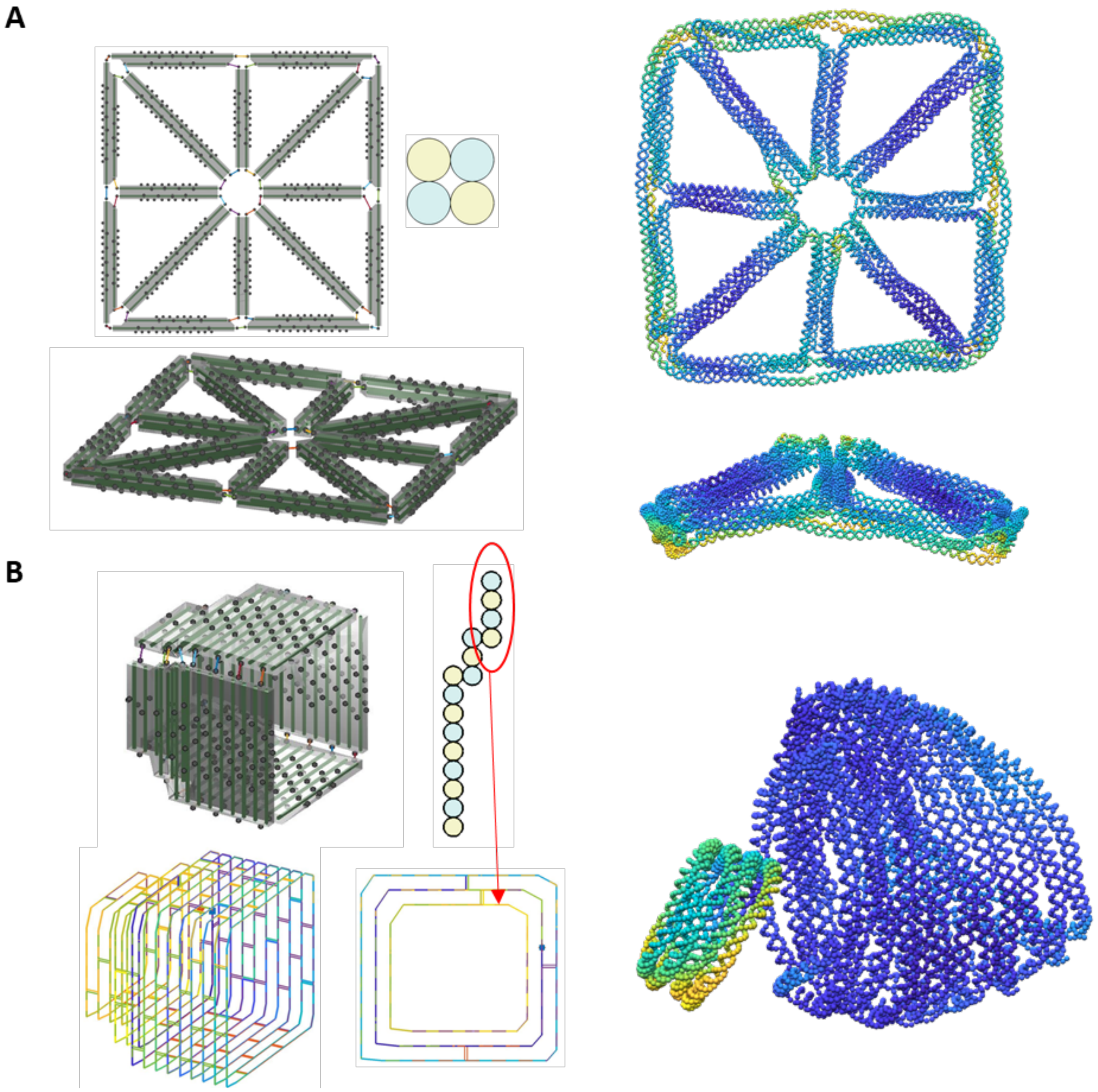

**Supplementary Figure 32: Insufficient local structural stability:** Out-of-plane motion of thin designs and insufficient crossovers in short bundles. (A) A wireframe structure with 2×2 square-lattice cross-sections was designed as a planar structure. Similar to Benson et al.[29], the wireframe structure may not have enough stiffness to constrain the intended planar shape. (B) Due to the lattice-based design practice, the staple algorithm ignores staple crossovers from both ends within a certain clearance (default is 8 bps, which can be changed through a UI) for each bundle. For short cylinders in the inner layer of this turbine, both the scaffold and staple strands had only one crossover to constrain the shape, leading to excessive separation between layers. The side view of the routing shows the lack of crossovers to the middle layer while the 4 cylinders in the inner layer still formed a ring. This effect happens in short bundles in surface modeling and is more likely to happen in square-lattice bundles because the crossover density from one cylinder to its neighbors is 1 crossover/32bps, instead of 1 crossover/21bps.

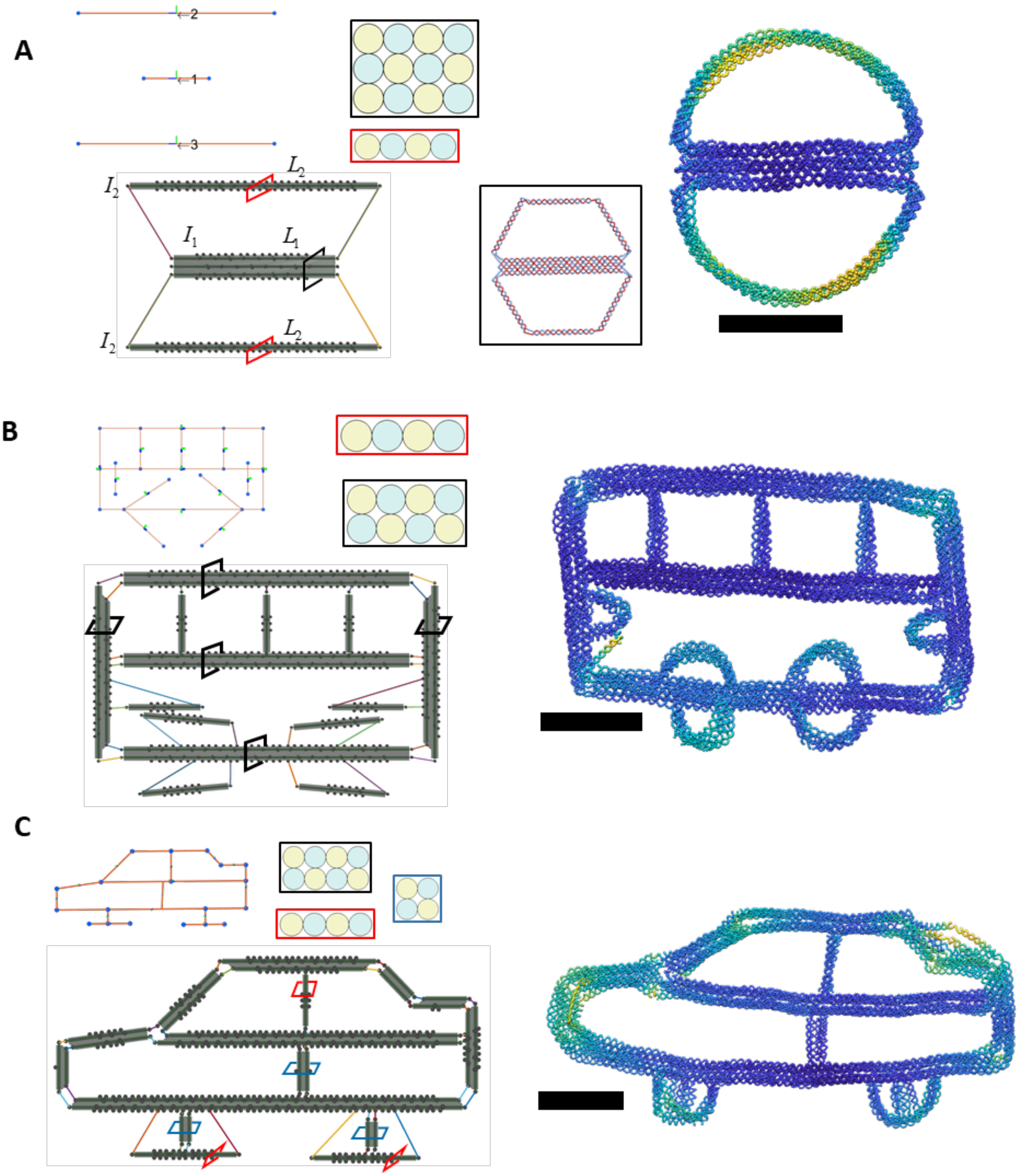

**Supplementary Figure 33: Design of curved shapes by intentionally connecting bundles with different lengths and rigidities.** (A) It is well-know that the stiffness of bundle components depends on the cross-section[28]. Connecting thin and long bundles with a thick and short bundle at both ends can bend the thin bundles into a curved shape. To make the relaxation process easier before running the oxDNA simulation, the bundle indices were assigned to split the bases in those bundles into three parts to transform the configuration (bottom middle inset). Examples structures with curved shapes in design of tires and lights on the (B) bus and (C) sedan structures. Scale bars = 20 nm.

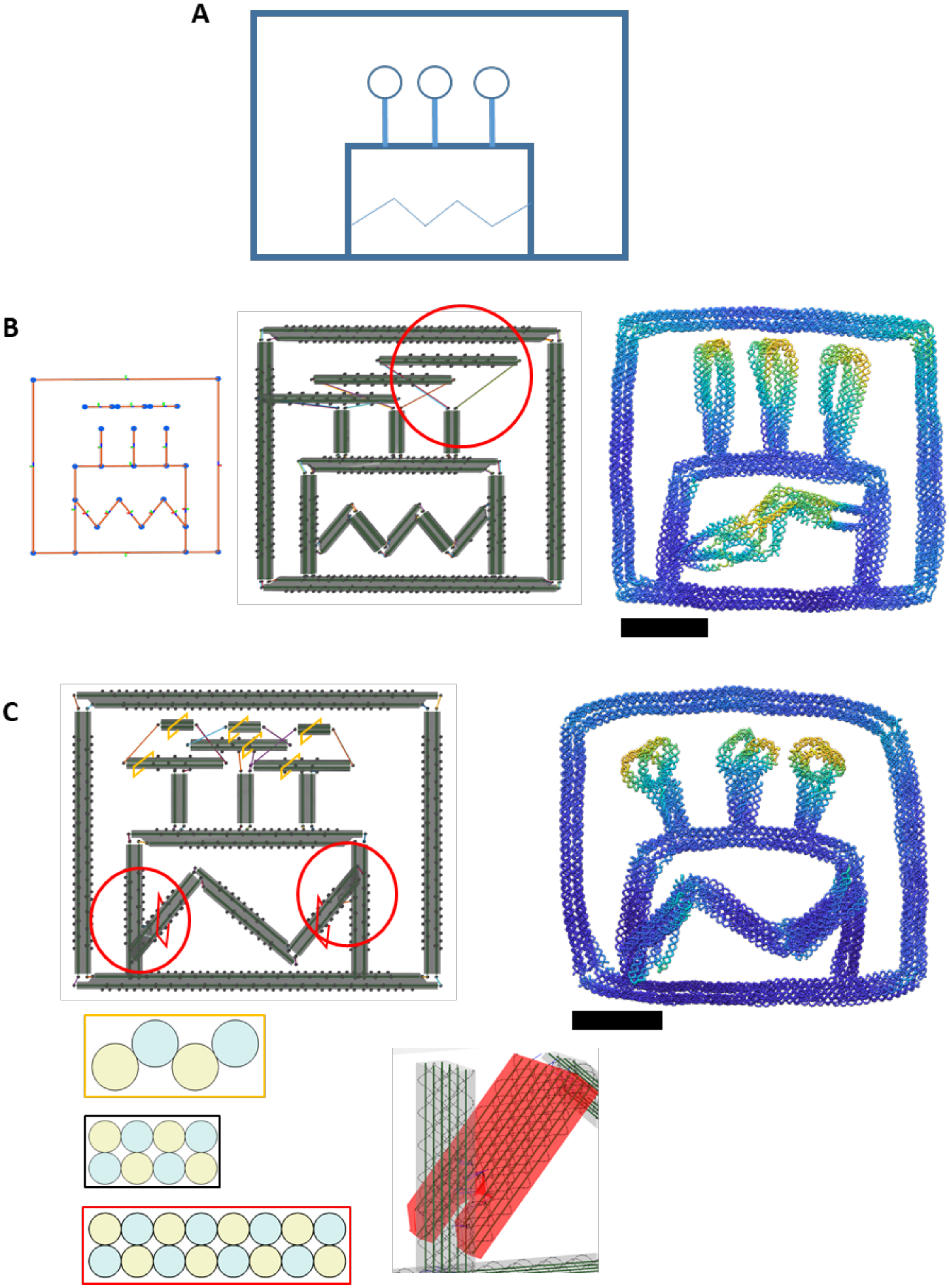

**Supplementary Figure 34: An example of iterative design illustrating success and failure at the sub-component level**. (A) The sketch of a cake to be designed. (B) Since the sketch contains many lines, a .STEP file was created in CAD software and then lines were converted into bundles in MagicDNA. This design has three design sub-systems: the outer and inner rectangle frames, the three candles, and the wavy feature on the cake. After running the simulation, we observed that the outer and inner frames were successfully designed; however, the flame of the candles and the wavy feature were not. (C) In the next design iteration, the previous design profile can be loaded and the bundles (the flame and the wavy features) that were not suitable can be replaced with improved designs using the bottom-up approach of inserting bundle components. This included increasing or decreasing the number of bundles to achieve the feature at the sub-system level and changing the geometry of the component such as lengths and cross-sections. Alternatively, the candle could be designed, simulated, and inserted back to the birthday cake design with duplication.

The results of the hierarchical design of complex static structures with experimental validation

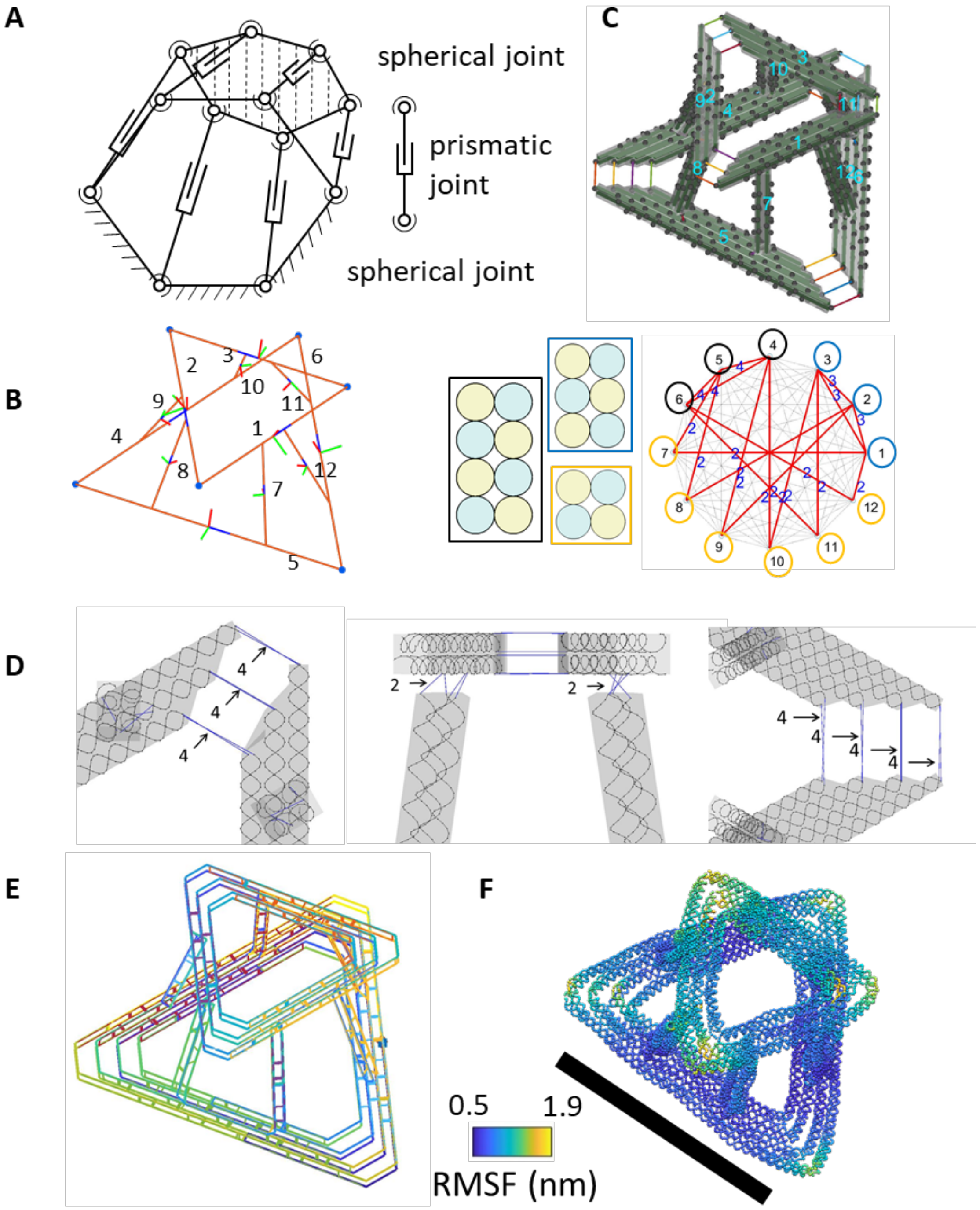

**Supplementary Figure 35: The design details of the Stewart Platform structure.** (A) Schematic of the Stewart platform[30] formed by 6 parallel limbs. (B) The line model, the connectivity matrix for assembly, and the cross-sections of each component. (C) The design model in the MagicDNA assembly panel. (D) The length of scaffold ssDNA connections between individual bundles. (E) The scaffold and staple routing of the final design. (F) The oxDNA simulation result. Scale bar = 50 nm.

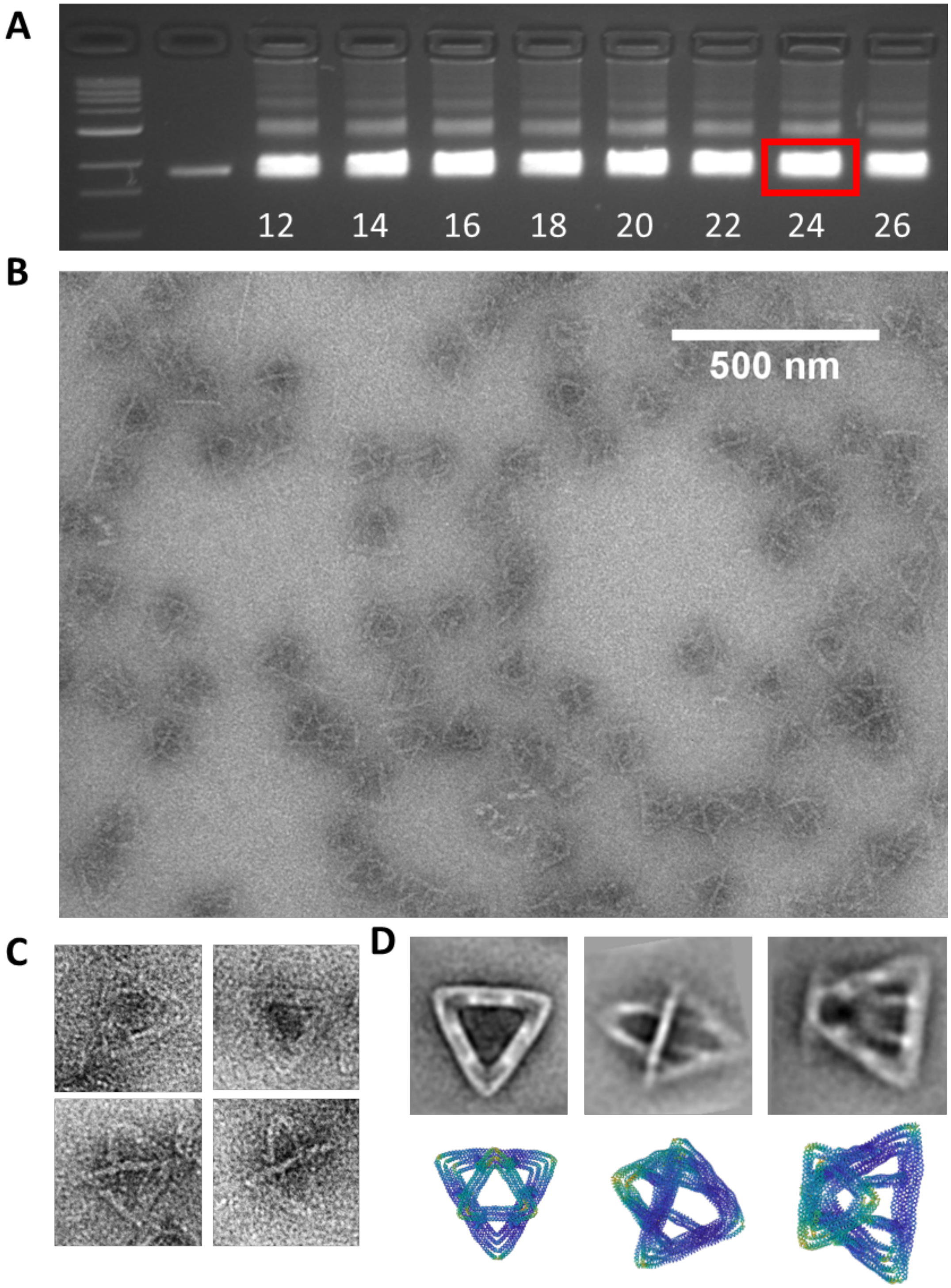

**Supplementary Figure 36: Experimental validation of the Stewart platform structure.** (A) Stewart platform structures were folded using a two-and-a-half-day thermal annealing protocol while testing a range of magnesium chloride concentrations from 12-26 mM and purified using agarose gel electrophoresis (buffer conditions 0.5x TBE 4 mM $MgCl_2$). (B) Zoomed out TEM image of gel-purified Stewart platform folded at 24 mM $MgCl_2$ with K10 peptide added post-purification for contrast. (C) Zoomed in TEM images of gel-purified Stewart platform structures folded at 24 mM $MgCl_2$. (D) Image averages of gel-purified Stewart platform structure folded at 24 mM $MgCl_2$ (Top) and oxDNA simulation results for reference (Bottom).

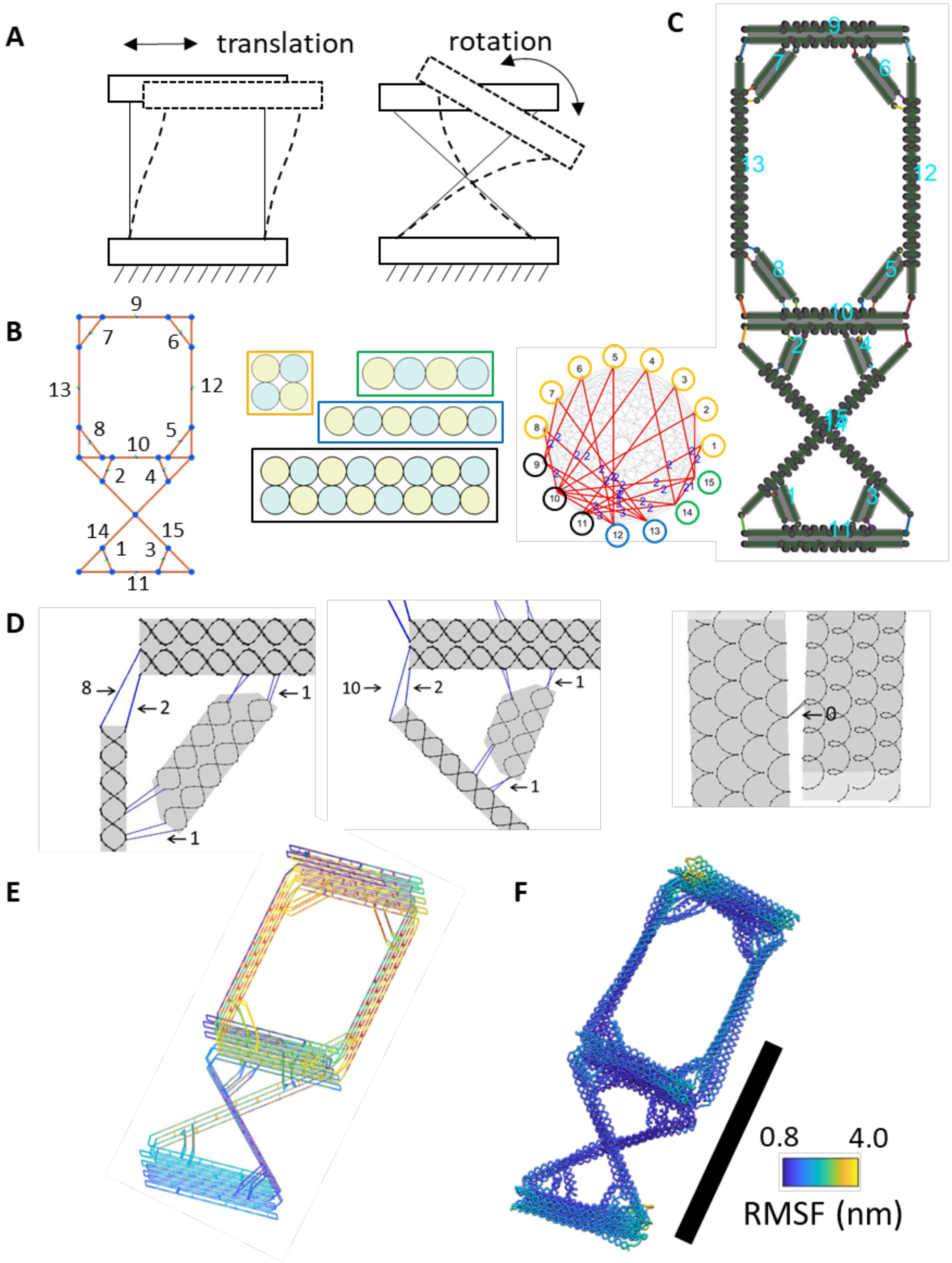

**Supplementary Figure 37: The design details of the compliant compound joint structure.** (A) Schematic of the translation and rotation compliant joints. (B) The line model, the connectivity matrix for assembly, and the cross-sections of each component. (C) The design model in the MagicDNA assembly panel. (D) The length of scaffold ssDNA connections between individual bundles. (E) The scaffold and staple routing of the final design. (F) The oxDNA simulation result. Scale bar = 50 nm.

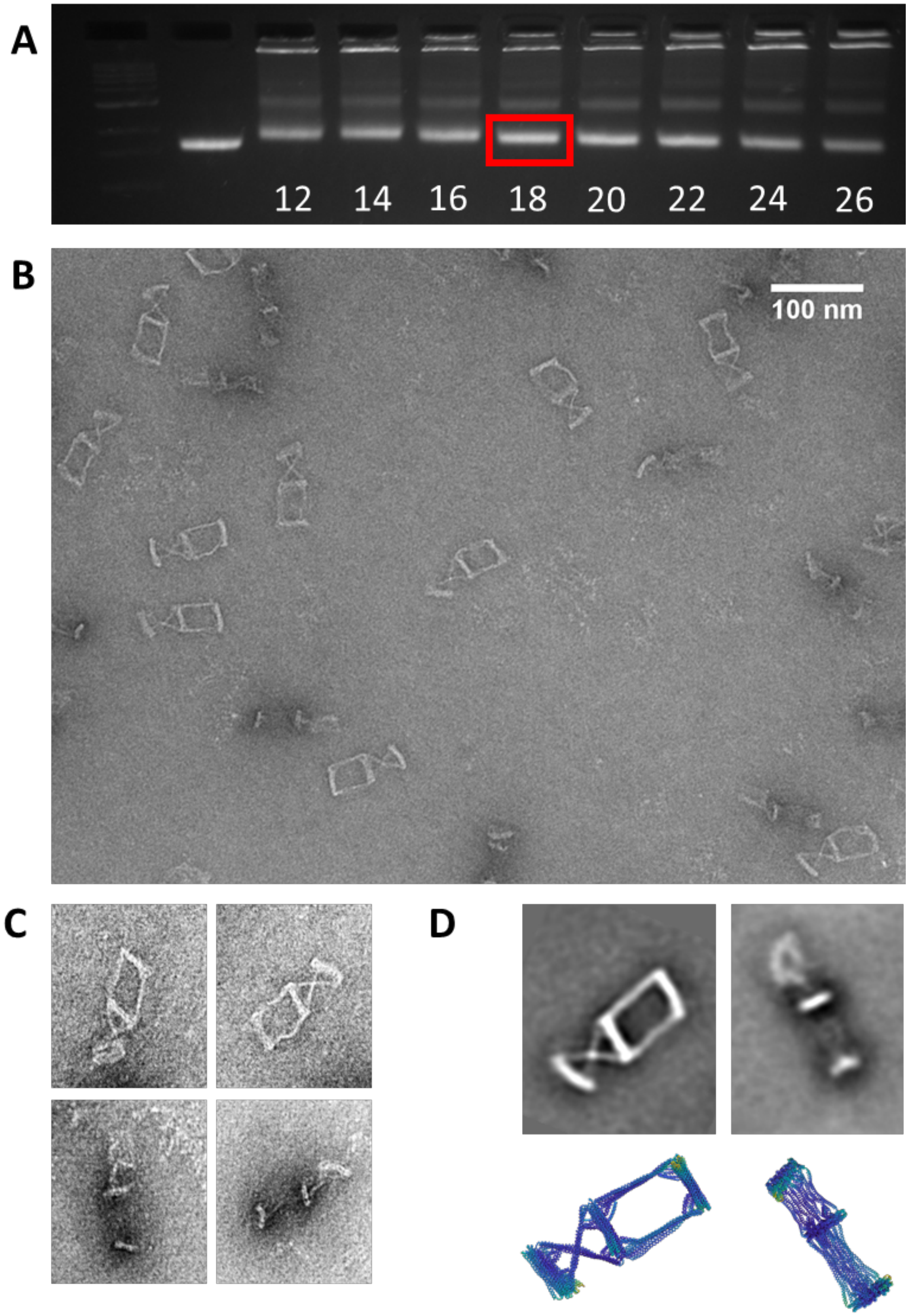

**Supplementary Figure 38: Experimental validation of the compliant compound joint structure.** (A) Compliant joint structures were folded using a two-and-a-half-day thermal annealing protocol while testing a range of magnesium chloride concentrations from 12-26 mM and purified using agarose gel electrophoresis (buffer conditions 0.5xTBE 11 mM MgCl2). (B) Zoomed out TEM images of gel-purified compliant joint folded at 18 mM MgCl2. (C) Zoomed in TEM images of gel-purified compliant joint folded at 18 mM MgCl2. (D) Image averages of gel-purified compliant joint folded at 18 mM MgCl2 (Top) and oxDNA simulation results (Bottom).

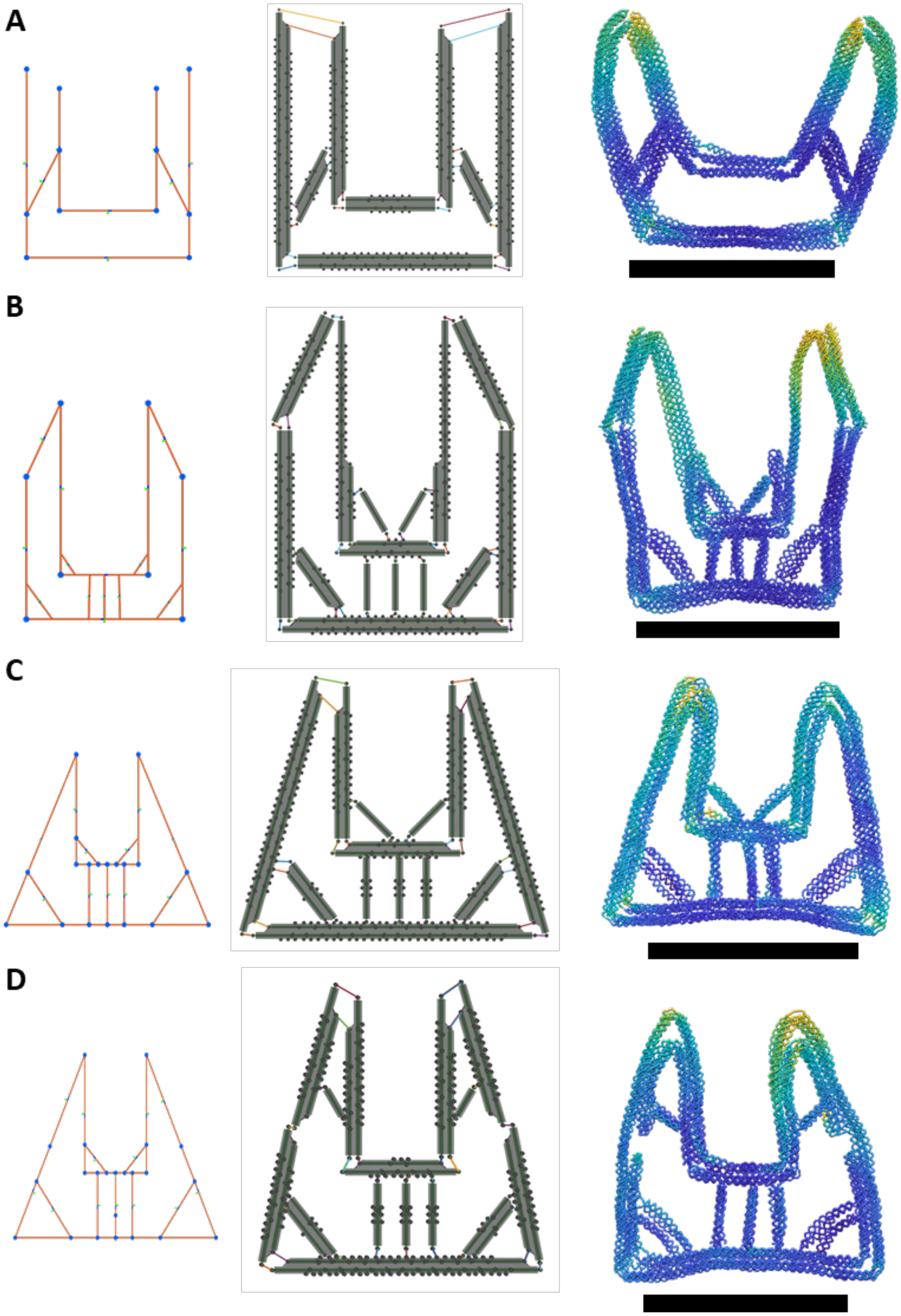

**Supplementary Figure 39: Representative designs and simulation results illustrating the iterative design process of the gripper structure.**

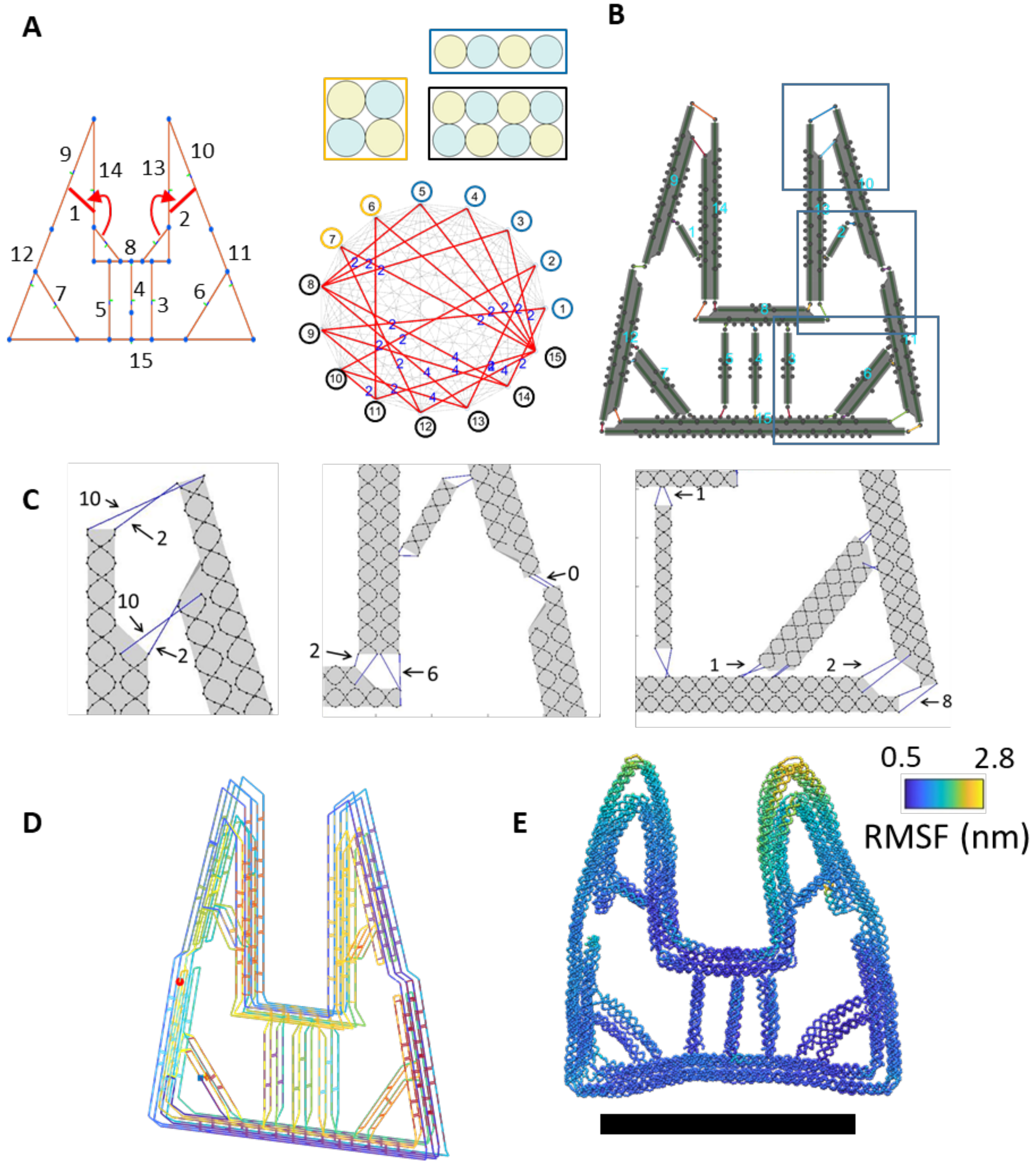

**Supplementary Figure 40: The design details of the gripper structure.** (A) The line model, the connectivity matrix for assembly, and the cross-sections of each component. (B) The design model in the MagicDNA assembly panel. (C) The length of scaffold ssDNA connections between individual bundles. (D) The scaffold and staple routing of the final design. (E) The oxDNA simulation result. Scale bar = 50 nm.

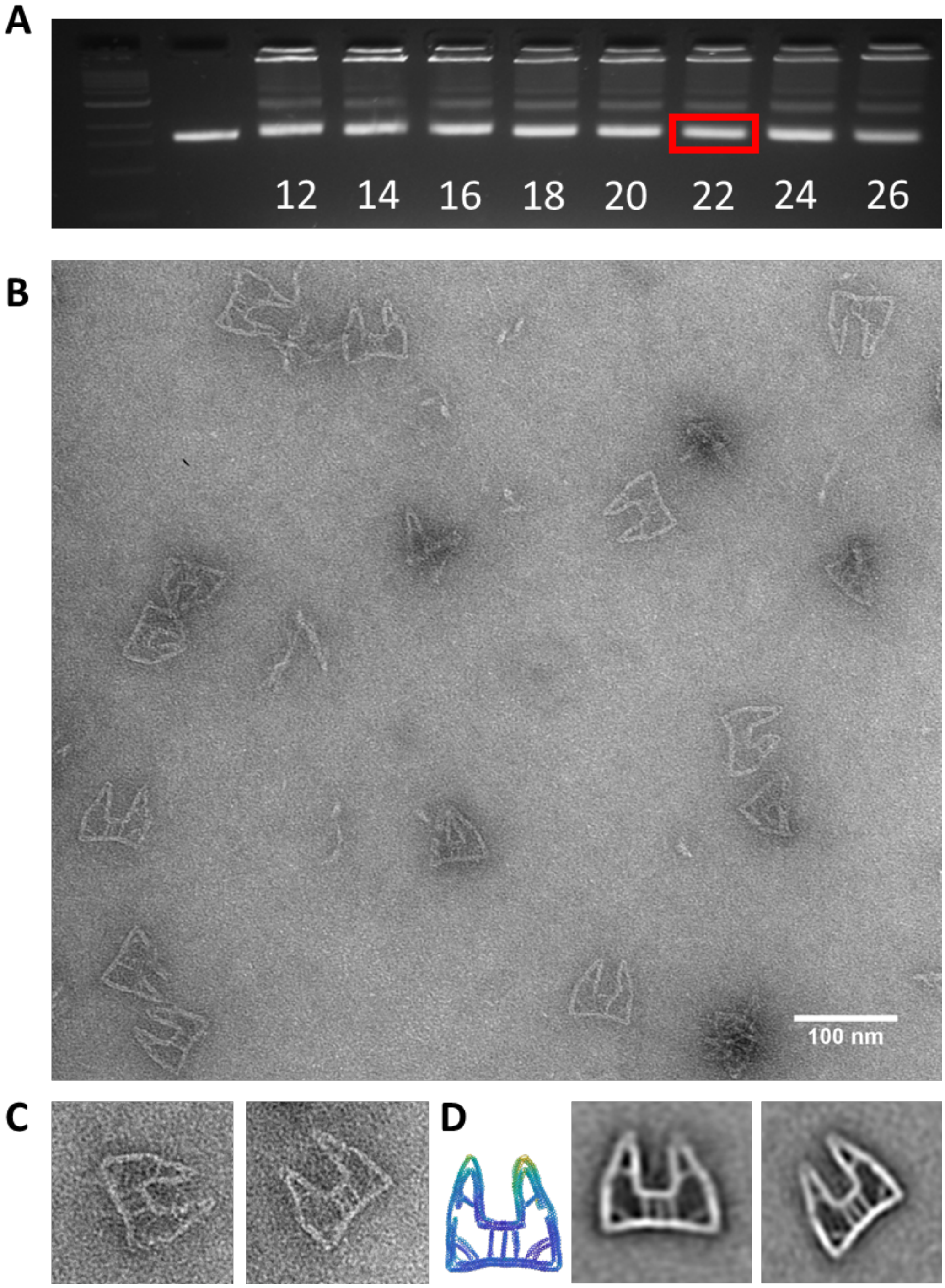

**Supplementary Figure 41: Experimental validation of the gripper structure.** (A) Gripper structures were folded using a two-and-a-half-day thermal annealing protocol while testing a range of magnesium chloride concentrations from 12-26 mM and purified using agarose gel electrophoresis (buffer conditions 0.5xTBE 11 mM MgCl2. (B) Zoomed out TEM image of gel-purified gripper structure folded at 22 mM MgCl2. (C) Zoomed in TEM images of gel-purified gripper structures folded at 22 mM MgCl2. (D) oxDNA simulation result of gripper structure (Left) and image averages of the gripper structure (Right).

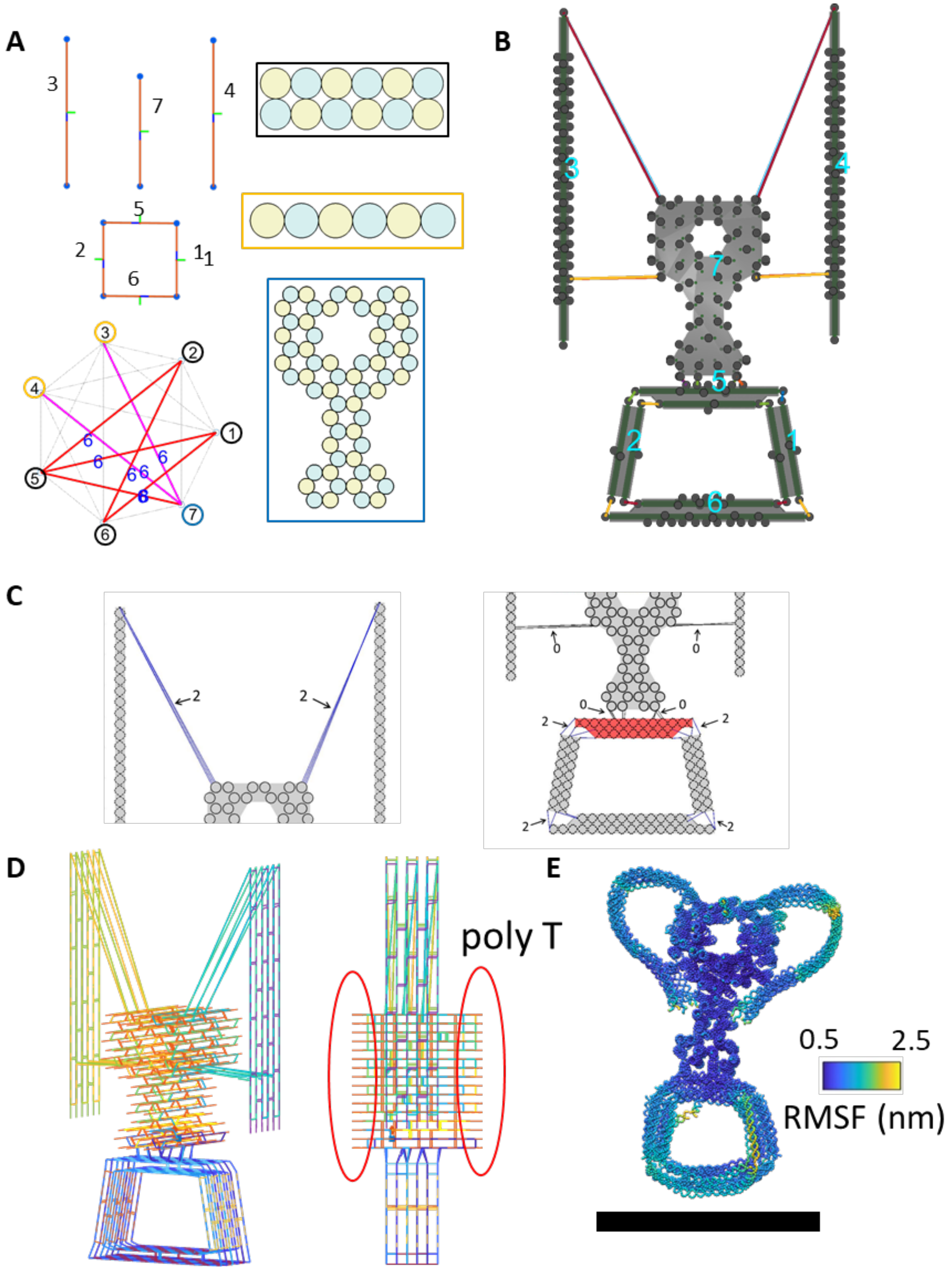

**Supplementary Figure 42: The design details of the trophy structure.** (A) The line model, the connectivity matrix for assembly, and the cross-sections of each component. Pink lines mean the connection between bundles were made by the manual GUI. (B) The design model in the MagicDNA assembly panel. (C) The length of scaffold ssDNA connections between individual bundles. (D) The scaffold and staple routing of the final design. (E) The oxDNA simulation result. Scale bar = 50 nm.

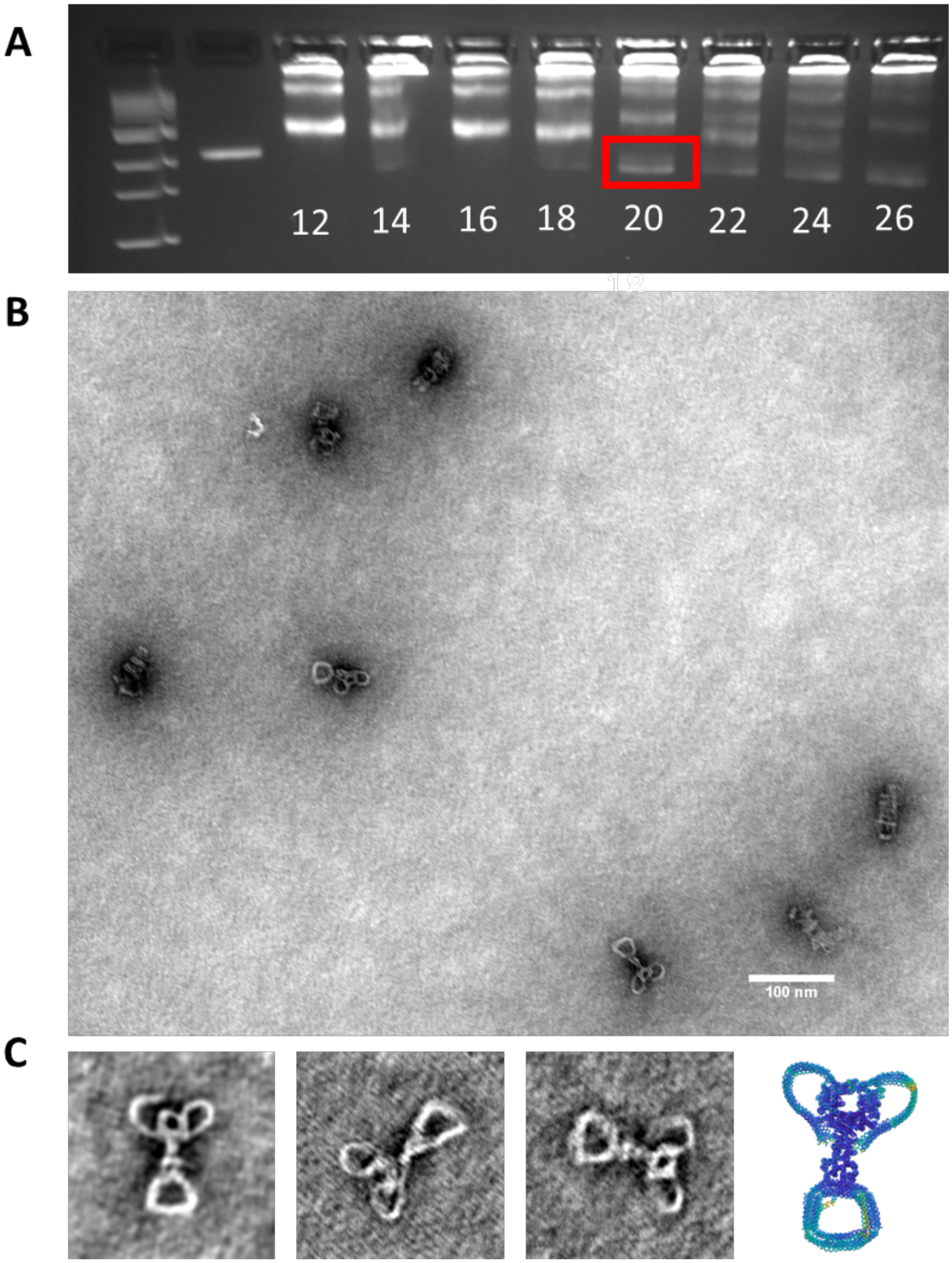

**Supplementary Figure 43: Experimental validation of the trophy structure.** (A) Trophy structure was folded using a two-and-a-half-day thermal annealing protocol while testing a range of magnesium chloride concentrations from 12-26 mM and purified using agarose gel electrophoresis (buffer conditions 0.5xTBE 5.5 mM MgCl2. (B) Zoom-out TEM image of gel-purified trophy structure folded at 20 mM MgCl2. (C) Zoomed-in TEM images of gel-purified trophy structure folded at 20 mM MgCl2 (Left) and oxDNA simulation result of trophy structure (Right).

#### Supplementary Section 4: Bottom-up and hierarchical design of reconfigurable assemblies

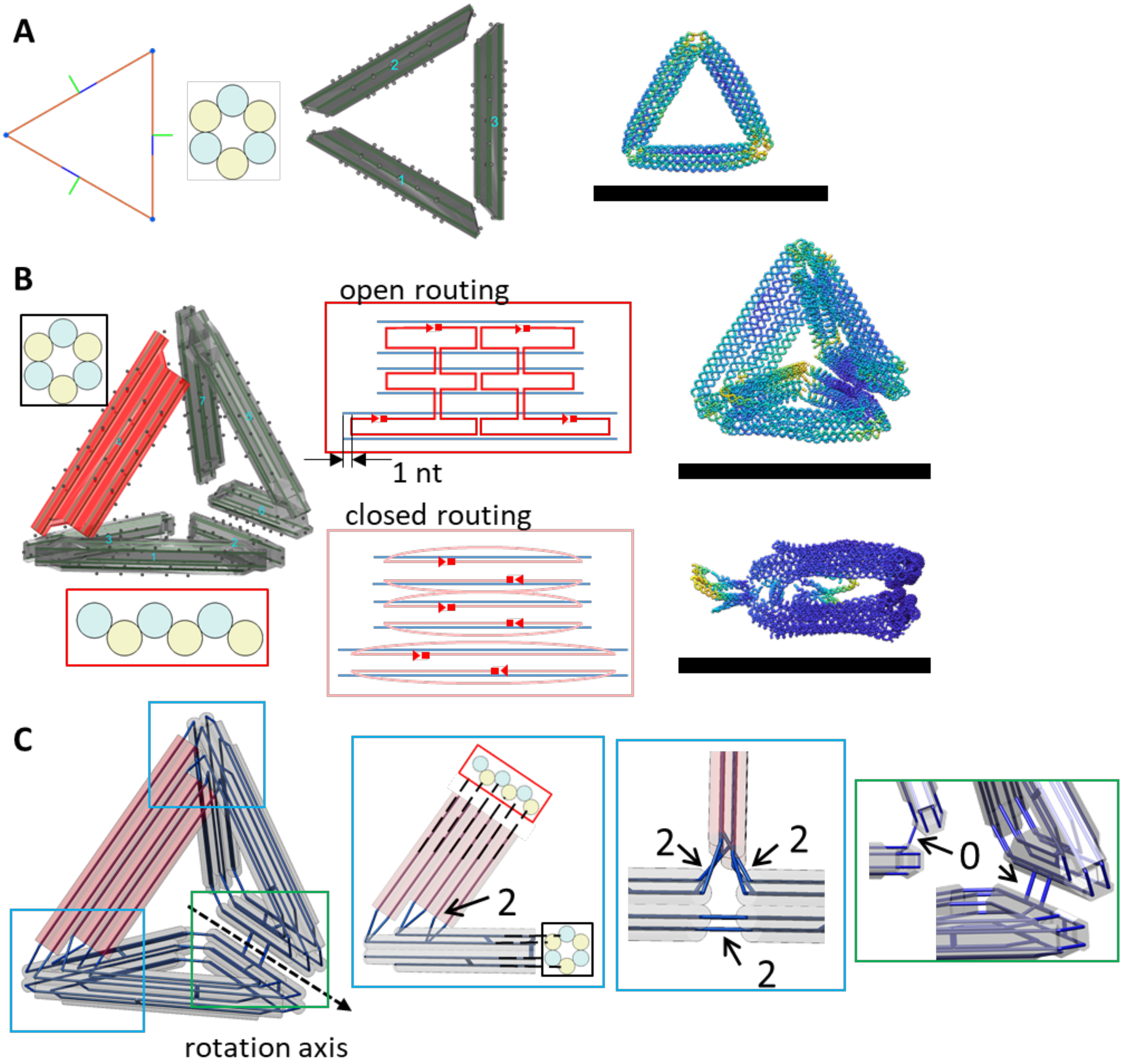

**Supplementary Figure 44: The basic unit design of the serial tetrahedron structure.** (A) A triangle plate made by three bundles with gradients at both ends. (B) One tetrahedron design with two triangle plates and one controller blade bundle (red). Here we purposely designed the blade bundle without scaffold crossover in scaffold algorithm. The staple routing for the blade in the open configuration is depicted schematically (middle top). Later, the caDNAno interface was used to edit the staple routing in the blade component such that both ends of each cylinder were pinned together for the closed configuration (middle bottom). (C) The rotation axis between two triangle plates is determined by two double-scaffold crossover connections (green boxes) while the opposite vertices were connected to the blade bundle (light blue boxes). The lengths of scaffold ssDNA are labeled and the staples all remain locally within bundles. Scale bars = 50 nm.

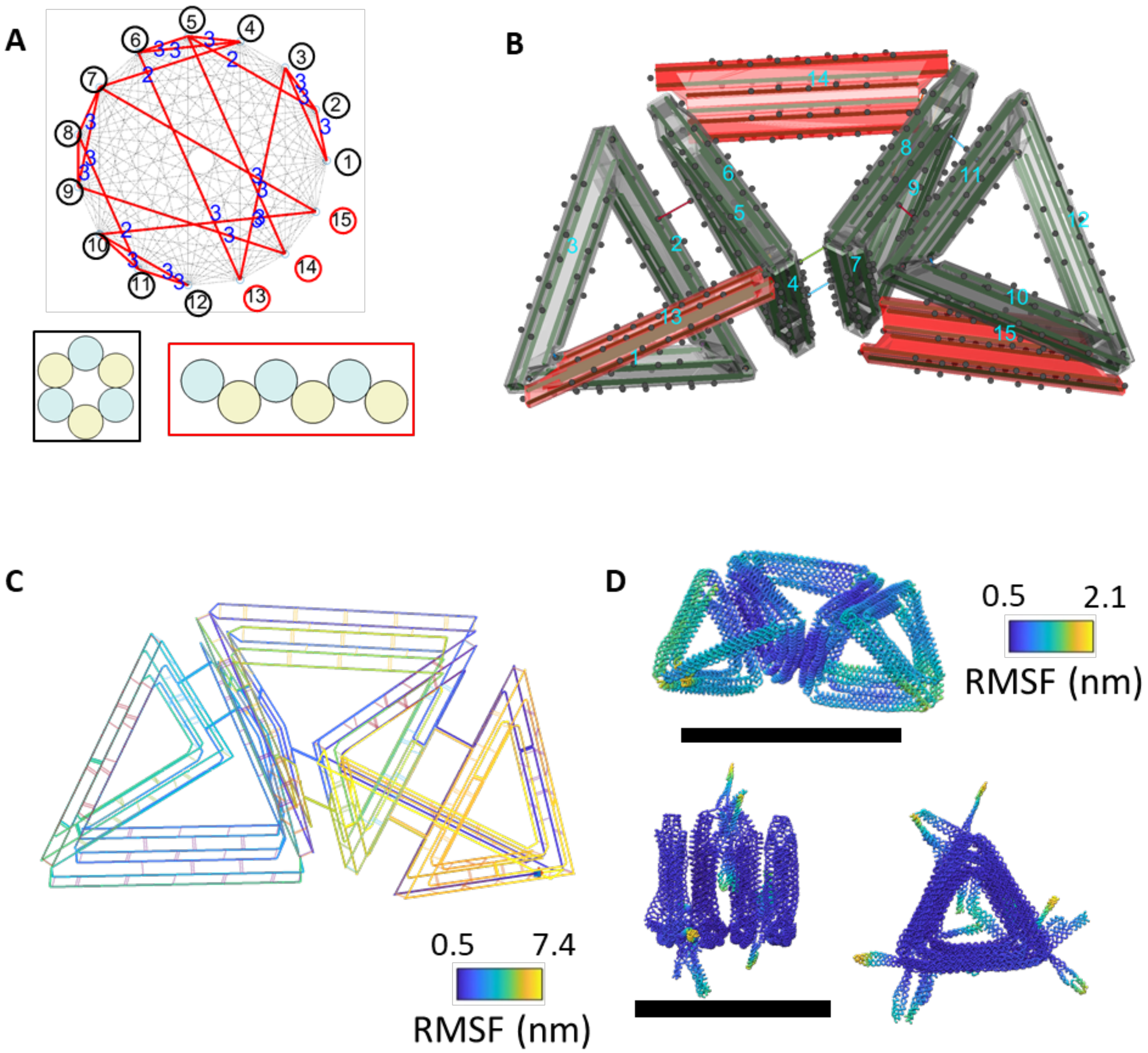

**Supplementary Figure 45: The design details of the serial tetrahedron structure.** (A) The connectivity matrix for assembly and the cross-sections of each component. (B) The design model in the MagicDNA assembly panel. (C) The scaffold and staple routing of the final design shown in chicken-wire representation. (D) The oxDNA simulation results of the open and close configurations. Scale bars = 50 nm.

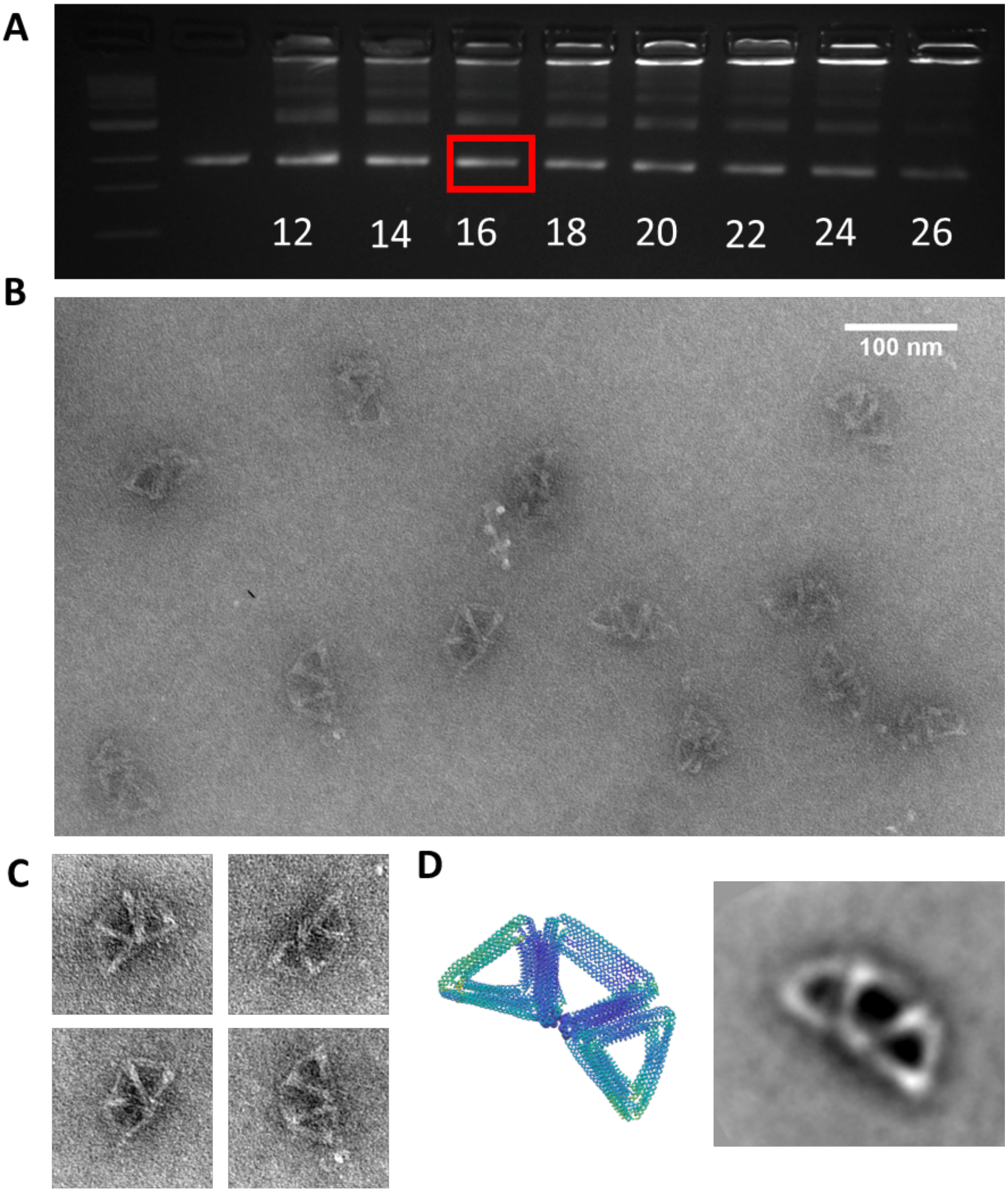

**Supplementary Figure 46: Experimental validation of the deployable tetrahedrons in the open configuration.** (A) Tetrahedron open configuration was folded using a two-and-a-half-day thermal annealing protocol while testing a range of magnesium chloride (mM) concentrations from 12-26 mM and purified using agarose gel electrophoresis (buffer conditions 0.5xTBE 11 mM MgCl2). (B) Zoomed-out TEM image of gel-purified tetrahedron open configuration folded at 16 mM MgCl2. (C) Zoomed-in TEM images of gel-purified tetrahedron open configuration folded at 16 mM MgCl2. (D) (Left) oxDNA simulation of tetrahedron open configuration (Right) image average of gel-purified tetrahedron open configuration folded at 16 mM MgCl2.

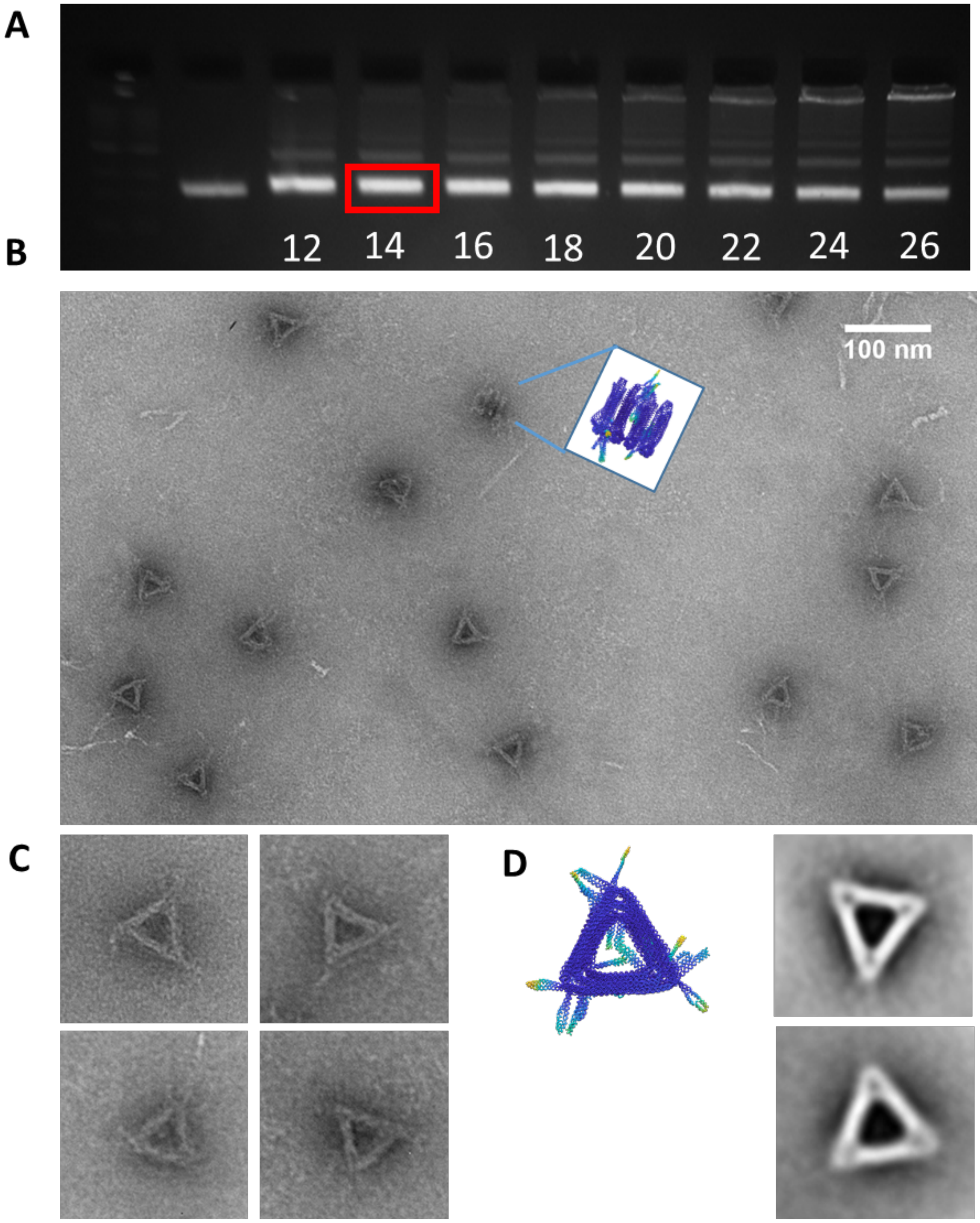

**Supplementary Figure 47: Experimental validation of the deployable tetrahedrons in the closed configuration.** (A) Tetrahedron closed configuration was folded using a two-and-a-half-day thermal annealing protocol while testing a range of magnesium chloride (mM) concentrations from 12-26 mM and purified using agarose gel electrophoresis (buffer conditions 0.5xTBE 11 mM MgCl2). (B) Zoomed-out TEM image of gel-purified tetrahedron closed configuration folded at 14 mM MgCl2. (C) Zoomed-in TEM images of gel-purified tetrahedron closed configuration folded at 14 mM MgCl2. (D) (Left) OxDNA simulation of tetrahedron closed configuration (Right) image average of gel-purified tetrahedron closed configuration folded at 14 mM MgCl2.

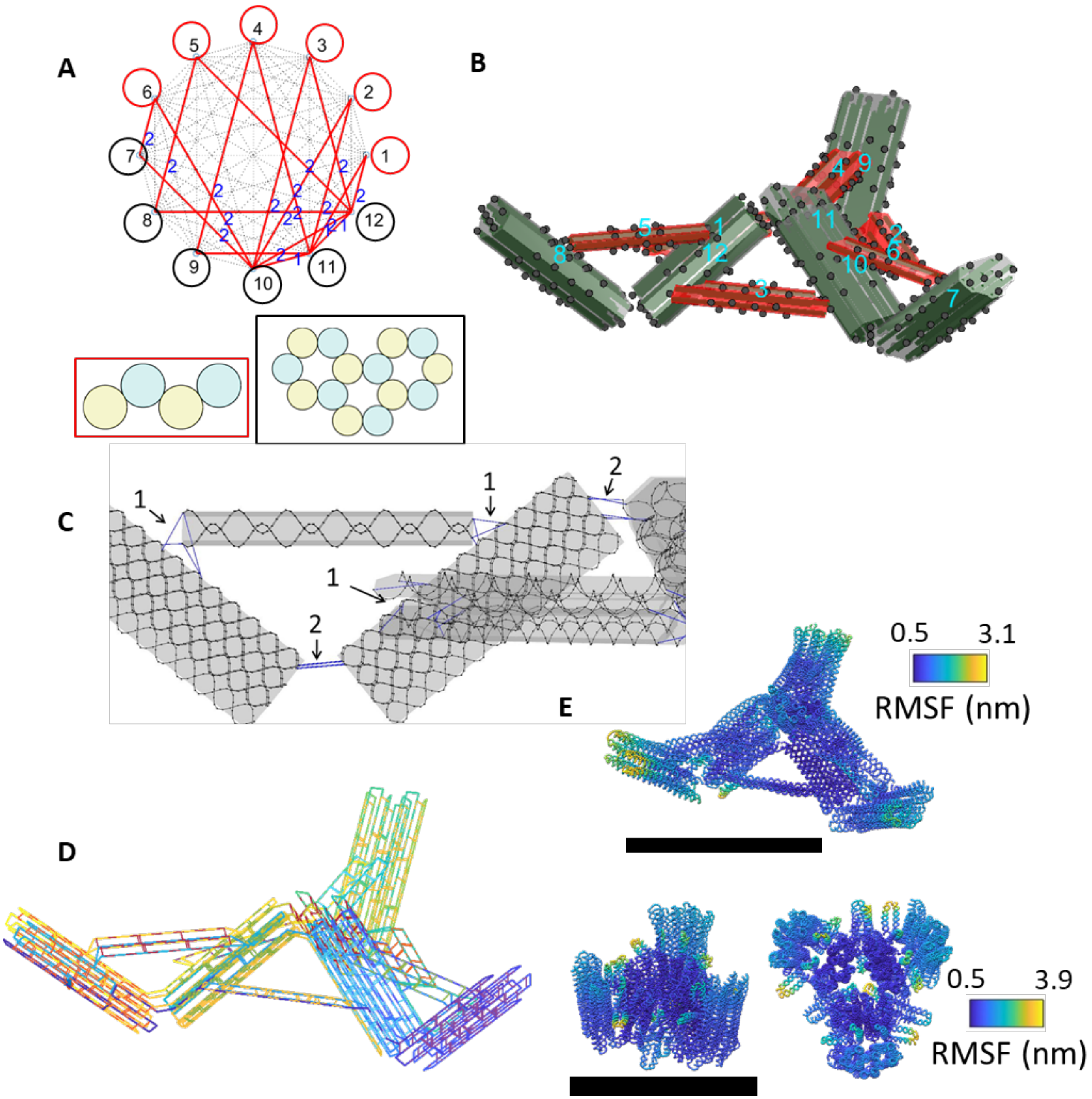

**Supplementary Figure 48: The design details of the umbrella structure.** (A) The connectivity matrix for assembly and the cross-sections of each component. (B)The design model in the MagicDNA assembly panel. (C) The length of scaffold ssDNA connections between individual bundles. (D) The scaffold and staple routing of the final design shown in chicken-wire representation. (E) The oxDNA simulation results of the open and close configurations. Scale bars = 50 nm.

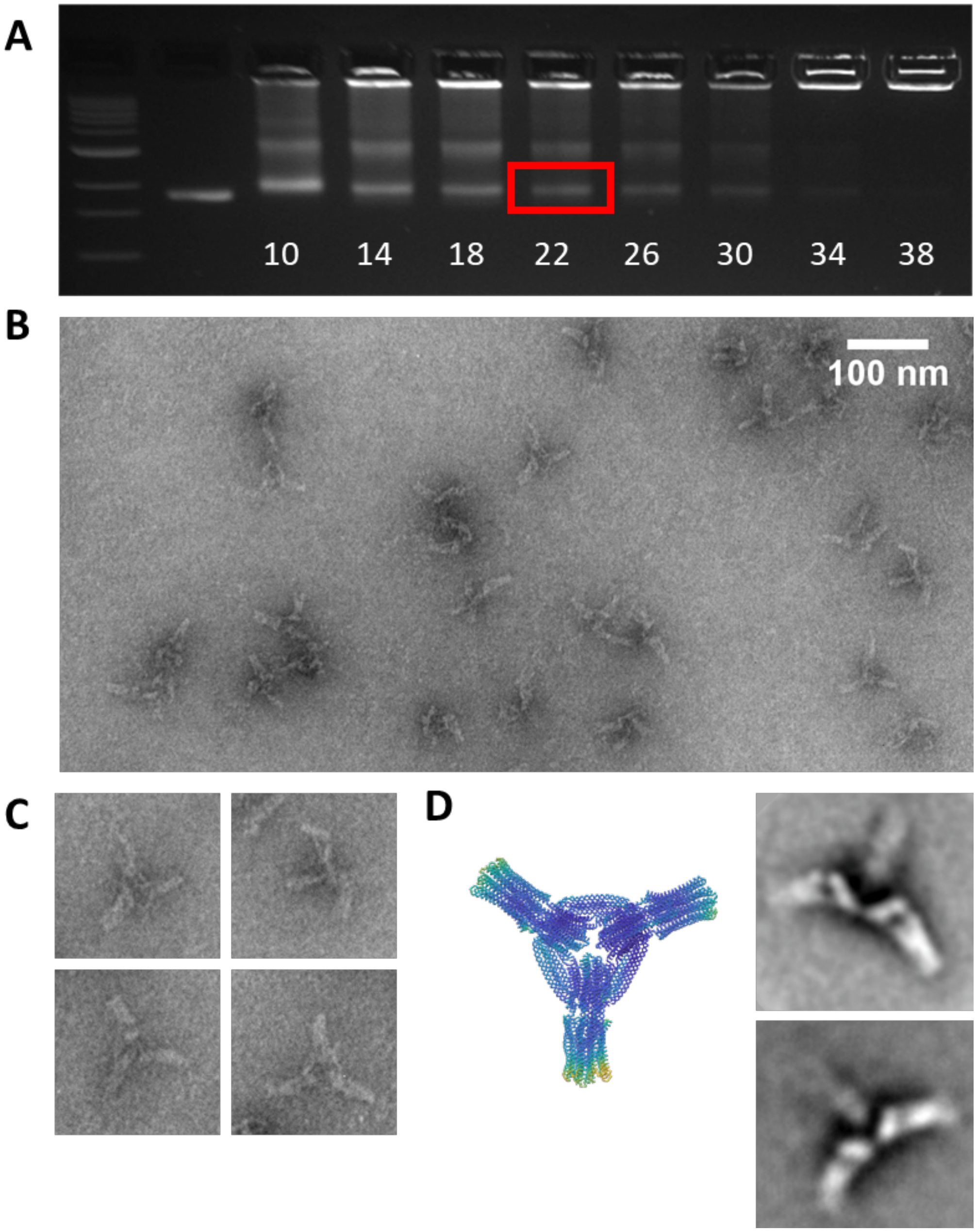

**Supplementary Figure 49: Experimental validation of the umbrella structure in the open configuration.** (A) The umbrella structure in the open configuration was folded using a four-and-a-half-day thermal annealing protocol while testing a range of magnesium chloride (mM) concentrations from 10-38 mM and purified using agarose gel electrophoresis (buffer conditions 0.5x TBE 11 mM MgCl2). (B) Zoom-out TEM image of gel-purified deployable mechanism open configuration folded at 22 mM MgCl2. (C) Zoomed-in TEM images of gel-purified The umbrella structure in the open configuration folded at 22 mM MgCl2. (D) (Left) oxDNA simulation of a deployable mechanism open configuration (Right) image averages of gel-purified open umbrella structure folded at 22 mM MgCl2.

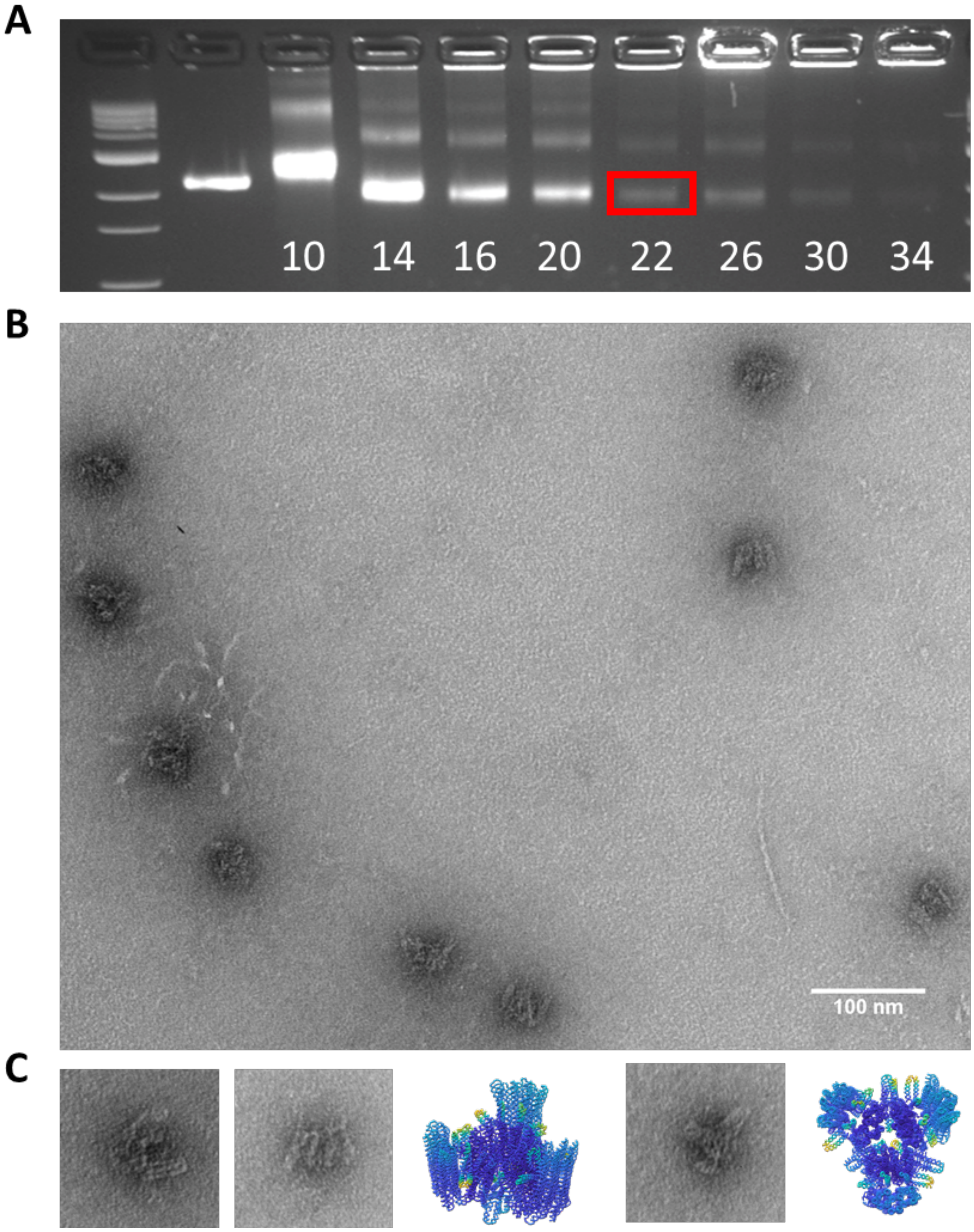

**Supplementary Figure 50: Experimental validation of the umbrella structure in the close configuration.** (A) The umbrella structure in the close configuration was folded using two-and-a-half-day modified thermal annealing protocol while testing a range of chloride (mM) concentrations from 10-34 mM and purified using agarose gel electrophoresis (buffer conditions 0.5x TBE 5.5 mM MgCl2). (B) Zoomed-out TEM image of gel-purified closed umbrella structures at 22 mM MgCl2. (C) (Left)Zoomed-in TEM images of gel-purified closed umbrella structures folded at 22 mM MgCl2. (Right) oxDNA simulation of the closed umbrella structure.

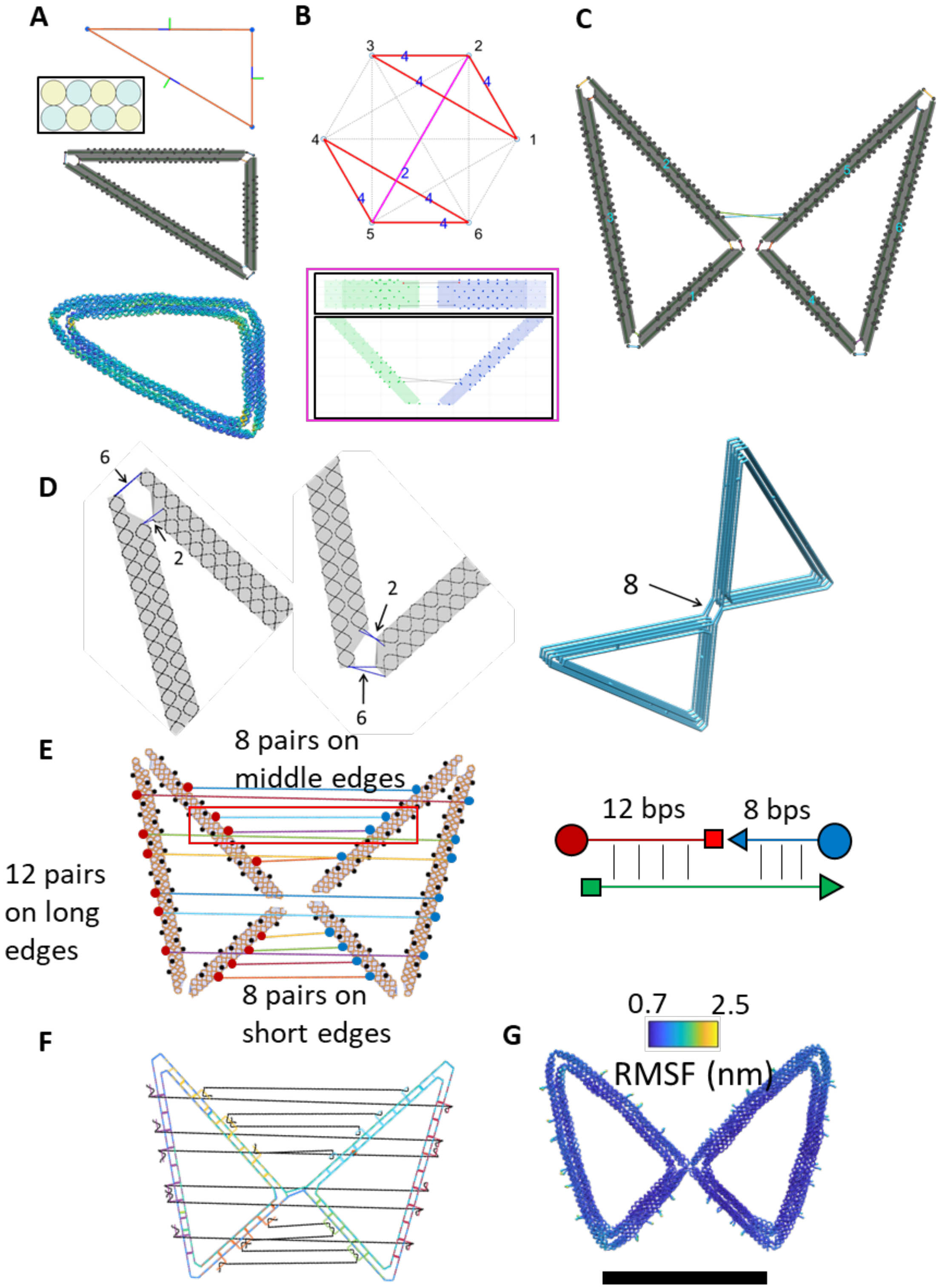

**Supplementary Figure 51: The design details of the butterfly mechanism.** (A) The basic unit of a triangle was designed and simulated by oxDNA, especially for the shape of vertices. (B) The connectivity matrix for assembly and the cross-sections of each component. The pink line indicates the connection between bundles 2 & 5 were made manually in MagicDNA (inset). (C) The design model in the MagicDNA assembly panel. Two double-scaffold crossover connections were intentionally created on the first and the last out-of-plane layers between the 2nd and 5th bundles in the assembly step. Later, after scaffold routing algorithm, these two double-crossover connections were shifted in caDNAno to the vertex to form two four-way junctions at the first and the last layers. (D) The length of scaffold ssDNA connections between individual bundles. (E) The snapshot of the overhang design tool where a total of 28 pairs of overhangs were added for actuation and polymerization. (F) The scaffold and staple routings of the final design shown in chicken-wire representation. (G) The oxDNA simulation results of the free configuration without closing strands. Scale bar = 50 nm.

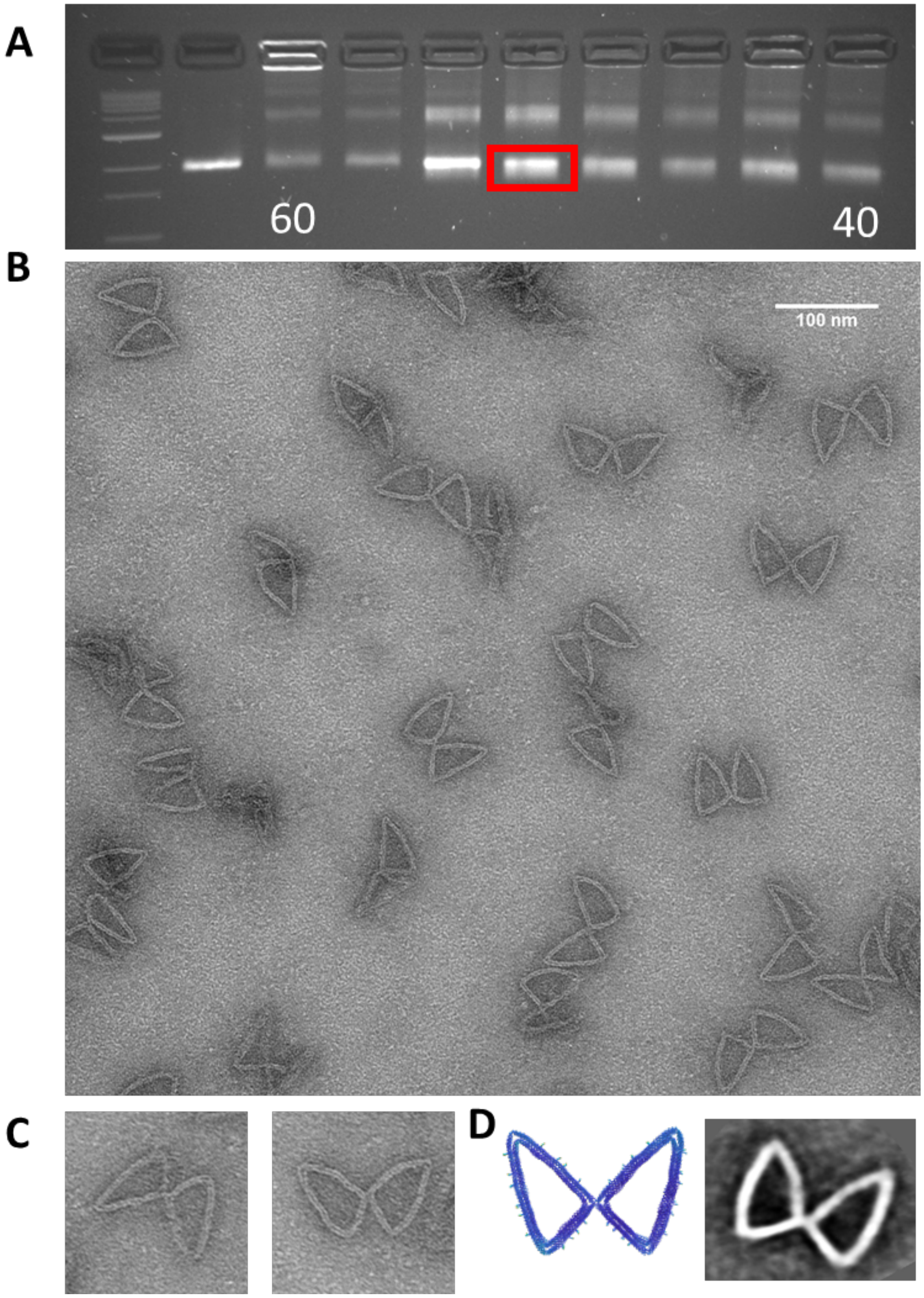

**Supplementary Figure 52: Experimental validation of the butterfly mechanism without actuation.** (A) Butterfly mechanism 4-hour modified thermal annealing protocol from 60°C-40°C at a salt concentration of 20 mM MgCl2 and purified using agarose gel electrophoresis (buffer conditions 0.5xTBE 11 mM MgCl2). (B) Zoom-out TEM image of gel-purified butterfly mechanism folded at 20 mM MgCl2. (C) Zoomed-in TEM images of gel-purified butterfly mechanism folded at 20 mM MgCl2. (D) (Left) OxDNA simulation of butterfly mechanism (Right) image average of gel-purified butterfly mechanism folded at 20 mM MgCl2.

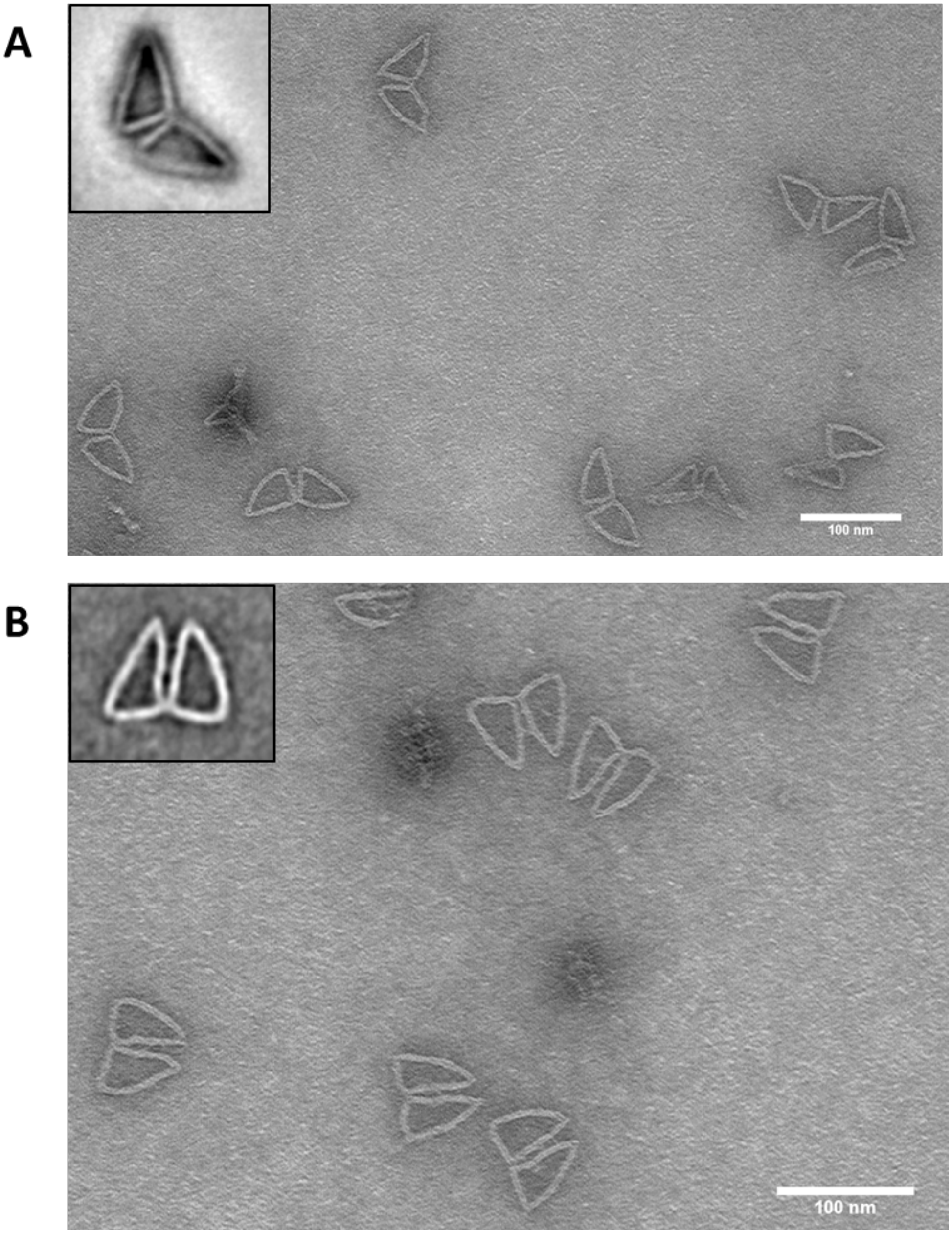

**Supplementary Figure 53: Experimental validation of the butterfly structure after actuation.** Butterfly mechanisms were actuated by adding strands. To close along A) the short edges B) the middle edge to actuate. Image averages of structures after actuation are shown in the inset. The butterfly mechanism was actuated after gel purification with 10x excess concentration of actuation staples relative to the concentration of the structure. The mixture was then incubated at 37°C for 2 hours.

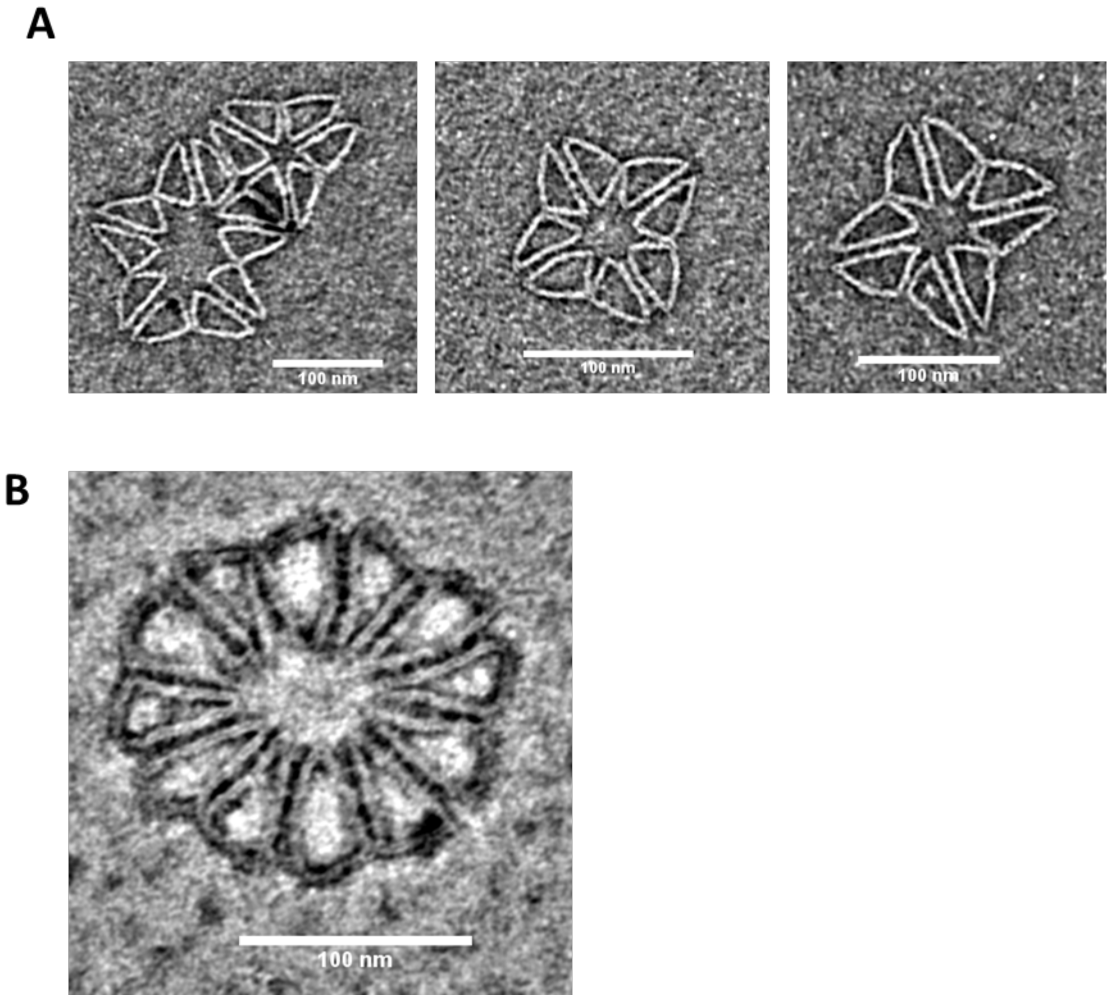

**Supplementary Figure 54: Experimental validation of polymerization of the butterfly mechanism after actuation.** (A) Polymerization of the actuated butterfly on short edges. (B) Polymerization of the actuated butterfly on middle edges. The actuated butterfly mechanism was polymerized into rings with 10x excess concentration of polymerization staples relative to the concentration of the structure. The mixture was then incubated at 37□C for 2 hours.

**Supplementary Section 5: Expanding the design domain of complex DNA assemblies**

Figures 2 to 4 and S23 to S54 have shown a variety of designs assembled by multiple bundles with different cross-sections and experimentally validated by fabrication and TEM imaging, including simple devices with demonstrated functions, complex designs with many components, and reconfigurable designs. Moreover, the hybrid design process implemented in MagicDNA enables the design of assemblies with more diversity and complexity than previous approaches due to an integration of five major perspectives (Fig. S55): 1) the ability to increase the number of components, 2) diversity in component geometries, 3) 3D manipulation of components and versatile assembly, 4) general routing algorithms and caDNAno interface, and 5) robust design by CG simulation at different levels.

1) Incorporating many components (Fig. S55A): Our comprehensive design processes, including the top-down (convert line model to bundles parametrically by .STEP files or sketching in MagicDNA), bottom-up (save and insert single bundles or sub-systems consisting of groups of bundles) and hybrid approaches, provide a way to quickly insert a large amount of bundles into the 3D assembly. Previously developed top-down design methods allow the inclusion of many components, but they do not encompass any bottom-up assembly, which is extremely useful in hierarchical design processes, in designs with repeated features, and in the process of iterating designs based on simulation feedback, which could require swapping in/out components. Furthermore, the 3D design models in MagicDNA remain straightforward to investigate and manipulate even with the increasing number of components, which is also especially helpful for displaying the current design profile and modification in the design iterations if needed.

2) Diversity in component geometry (Fig. S55B): Prior bottom-up design approaches allow versatile control over component geometry, but those approaches are not scalable to many components; whereas previous top-down methods can include many components they do not allow flexibility in individual component design. We combine a capability for 3D assembly with many components with versatility in component geometry. The built-in options in MagicDNA already allow for versatile component design (e.g. varied geometry across components, hybrid lattices, and edge gradients). MagicDNA also has GUIs to allow for custom cross-sections in both square and honeycomb lattices, while the bundle editing GUI allows the inward or outward extrusion of each helix in a bundle to precisely control non-uniform edge gradients or internal cavities. This allows users to design across the three major classes of DNA nanostructures including lattice-based, surface-based, and wireframe. Furthermore the ability to combine all these into one assembly significantly expands the structure design spectrum.

3) Versatility in 3D assembly and connectivity with tunable mechanical properties (Fig. S55C): The process of component assembly mimics assembly operations in commercial CAD software and provides intuitive real-time 3D visualization of the configuration. Again, different visualizations of the connectivity enable complex designs with many components connected in 3D. In particular, the ability to manipulate (translate and/or rotate) individual components or groups of components (i.e. sub-systems) facilitates straightforward manual assignment of connectivity or automated assignment of connectivity based on distances between connection sites. The connectivity matrix quantitatively displays the connections between bundles, which is extremely useful for designs with a large number of bundles. Furthermore, categorizing the connection site nodes into end and side nodes enables flexible design of rigid, compliant, or

flexible connections with a wide variety of geometries (e.g. vertex, hinge, T-junction, etc.). This ability to tune the joint properties (example in Fig. S31) is key to enabling dynamic devices with well-controlled motion and mechanical properties, which is critical for DNA-based robots.

4) Generalized routing algorithms and interface to caDNAno for fine-tuning (Fig. S55D): Given such versatility in design at component and assembly levels and the ability to include a large number of components arranged and connected in 3D, the development of a general scaffold routing algorithm (Fig. S55D) is necessary to handle this process that would be tedious and error-prone to carry out manually. Feasibility checks, such as pairing of the cylinders in the custom cross-section GUIs are embedded to ensure compatibility with the algorithm requirements (e.g. even number of helices in a component). To enhance the generality of the algorithm, MagicDNA also collects user-inputs to define relevant parameters such as routing at ends or direction of cylinder pairing. The computational efficiency was also a key consideration. The most complex design in this study, the airplane with single-scaffold routing (~32 kbps) in the center of Fig. 5, took less than 20 seconds to find a scaffold routing. On the other hand, for some specific cases, while the general routing algorithm can produce an initial routing, it is also useful to leverage the interface to caDNAno fine-tune the design details with the assistance of the visualization tools in MagicDNA such as the mapping between 2D diagram, 3D visualization, and the staple list.

5) Robust design by CG simulation at different levels (Fig. S55E): Based on the previous four perspectives, this hybrid design framework implemented in MagicDNA enables diverse and complex assemblies. Furthermore, interfacing with coarse-grained simulation, specifically oxDNA simulations, allows for rapid virtual prototyping, which is critical especially for complex assemblies. With simulation feedback, the ease of modifying the design parameters in the previous four perspectives is crucial to address local design flaws and eventually assure the target design match the requirement for the application. For multiple cases, we demonstrated a multi-scale iterative design optimization that improves the efficiency and robustness of the design process.

We illustrate this broad and versatile design spectrum enabled by hybrid top-down and bottom-up design in MagicDNA in Figs. 5, S56 (lattice-based), S57 (surface-modeling), S58 (wireframe), and S59 to S69.

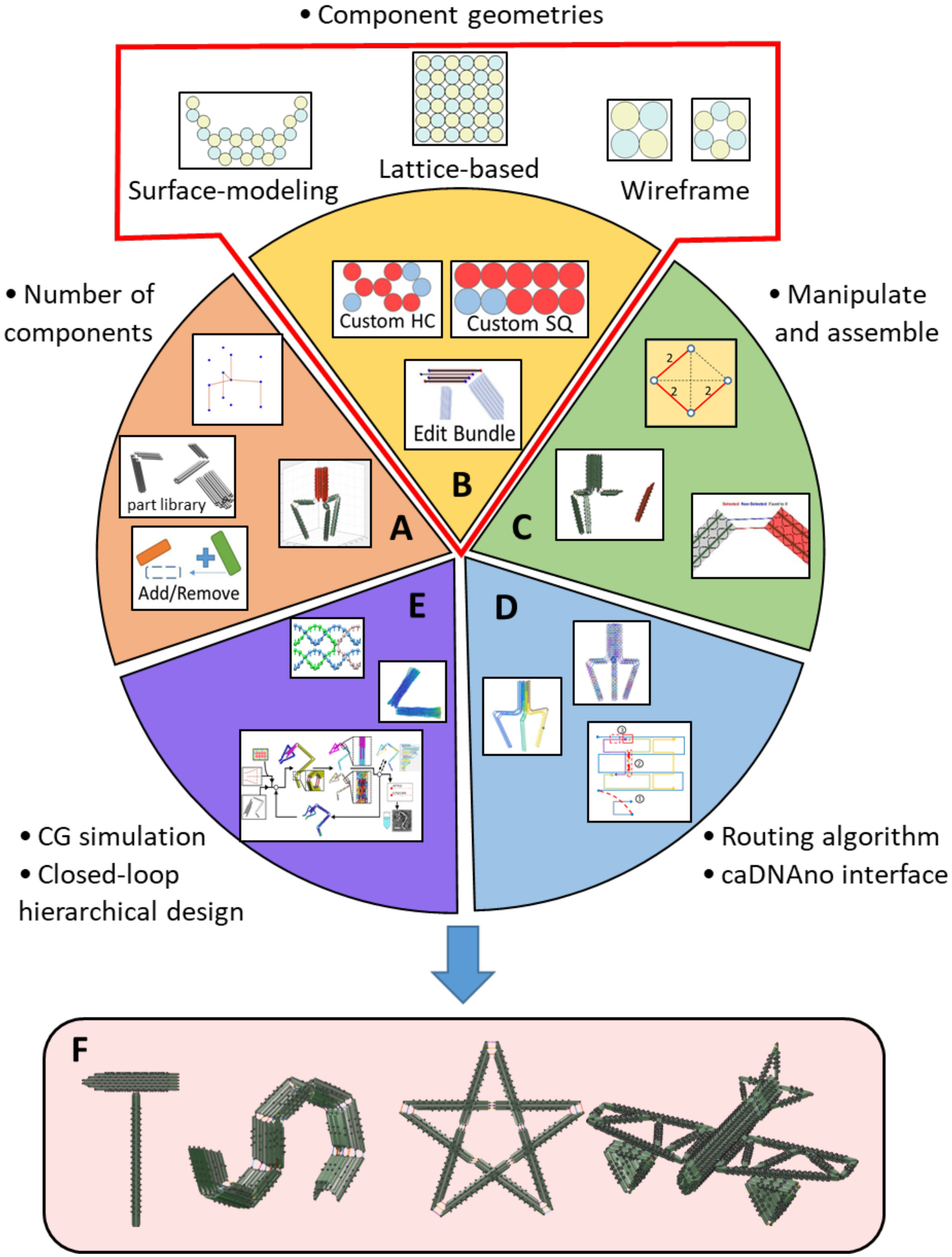

**Supplementary Figure 55: The functionality and features in MagicDNA to broaden the design spectrum.** (A) Due to the hybrid top-down and bottom-up approach, many bundles can be inserted into the assembly for designing more complex structures with increased number of components. Direct 3D models are also helpful with forming connections between more components in complex 3D assemblies. (B) Combining wireframe, honeycomb and square lattices and inputting edge gradients allows flexible design of diverse components. Meanwhile, the bundle editing GUI built-in MagicDNA allows changing the lengths of individual helices for non-uniform edge gradients and hollow cavities for further variety in design. (C) Manipulations in 3D, the connectivity matrix, and the ssDNA scaffold GUI enable versatile assembly between components, including end-to-end vertex design, end-to-side, side-to-side (layered crossovers (65)), and tunable mechanical properties of joints. (D) Such a broad design space requires a robust automatic routing algorithms. To deal with some specific applications, the caDNAno interface always can serve as an option to locally fine-tune the routing details. (E) Integrating with coarse-grained simulation enables a closed-loop design framework to computationally evaluate the design profile at different levels and bottom-up assembly of sub-system designs for efficiency. (F) Example designs across the versatile design spectrum that emerges from the integration of these four features of the hybrid design process in MagicDNA.

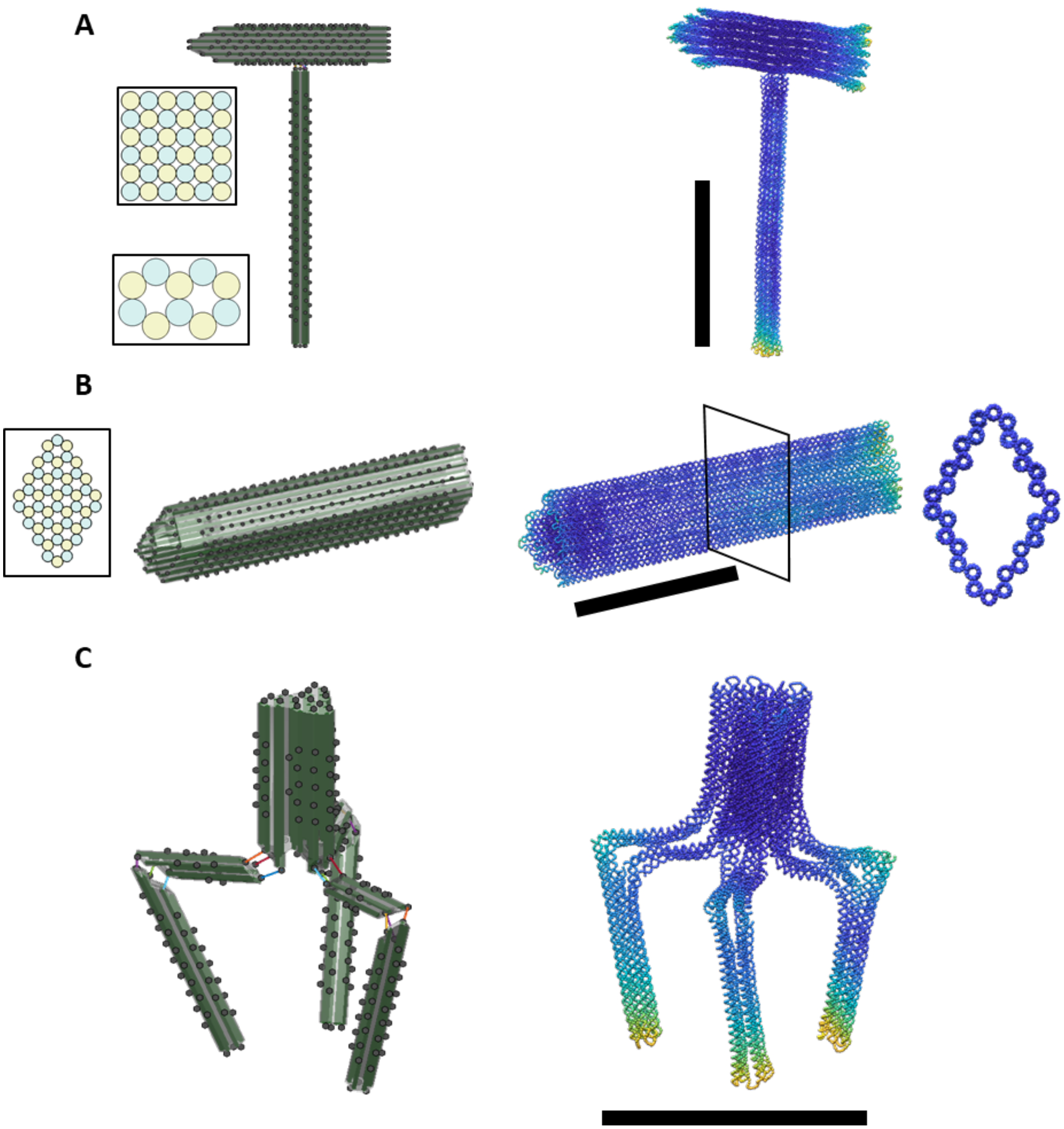

**Supplementary Figure 56: Examples of lattice-based structures.** (A) A hammer design. Scaffold length = 8916 nt. (B) The main body (i.e. fuselage) of the airplane design. The outermost layer was extended to have shell feature. Scaffold length = 10786 nt. (C) A claw design. Scaffold length = 8063 nt. Scale bars = 50 nm.

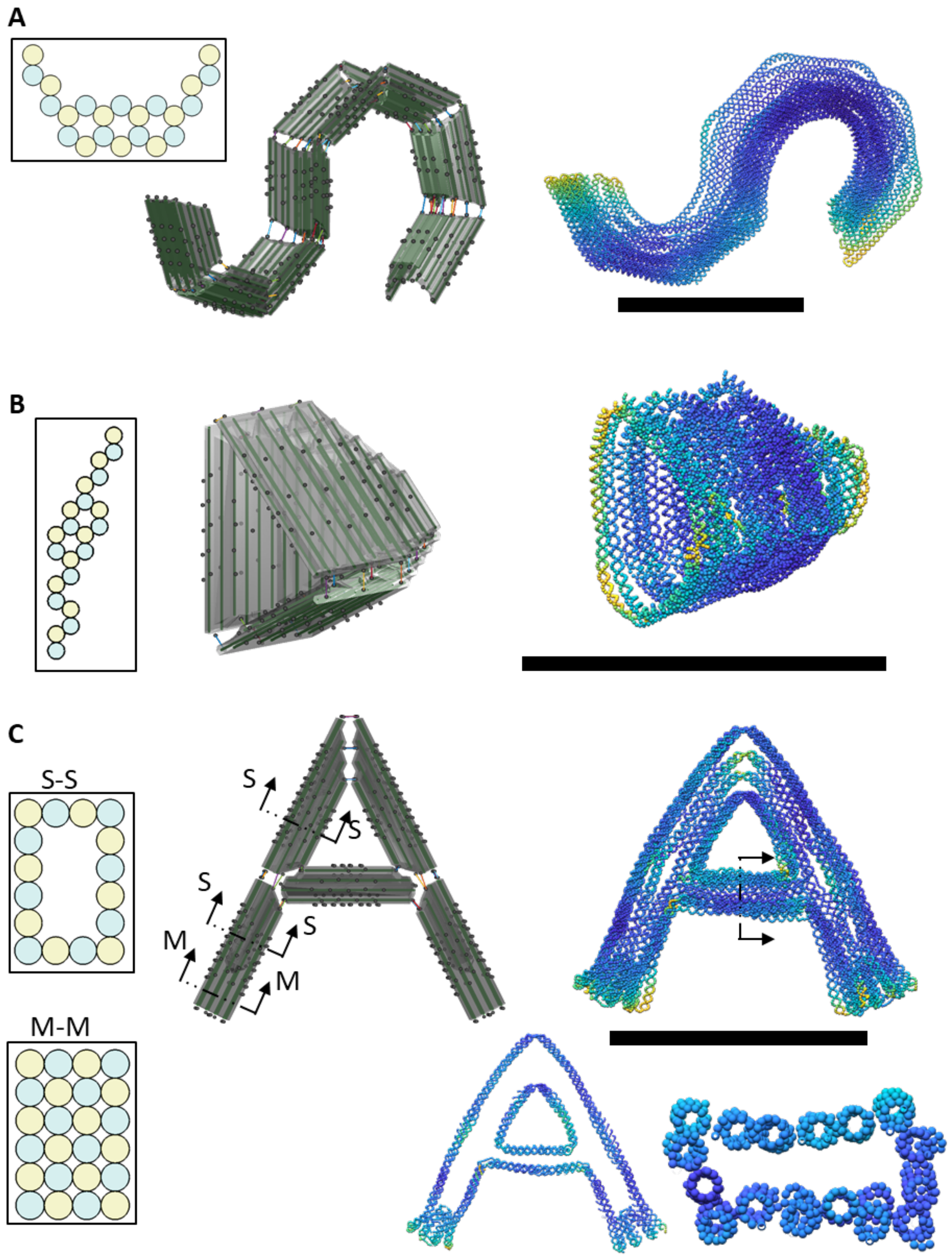

**Supplementary Figure 57: Example of surface-based structures.** Surface-modeling follows two steps: 1) arranging the cylinders with one or two layers along a specific direction in a plane, and 2) extruding or revolving or sweeping the in-plane geometry along a specified path to obtain a 3D structure. Variable cross-sections and gradients at ends (cylinders with different lengths) can also be applied. (A) A curved cross-section was swept along an S-shaped path. Scaffold length = 11232 nt. (B) Approximating a revolved cross-section with three bundles for the turbine sub-system in the airplane design. Scaffold length = 4380 nt. (C) A shell A-shaped structure with hollow cavity. Scaffold length = 6797 nt. Scale bars = 50 nm.

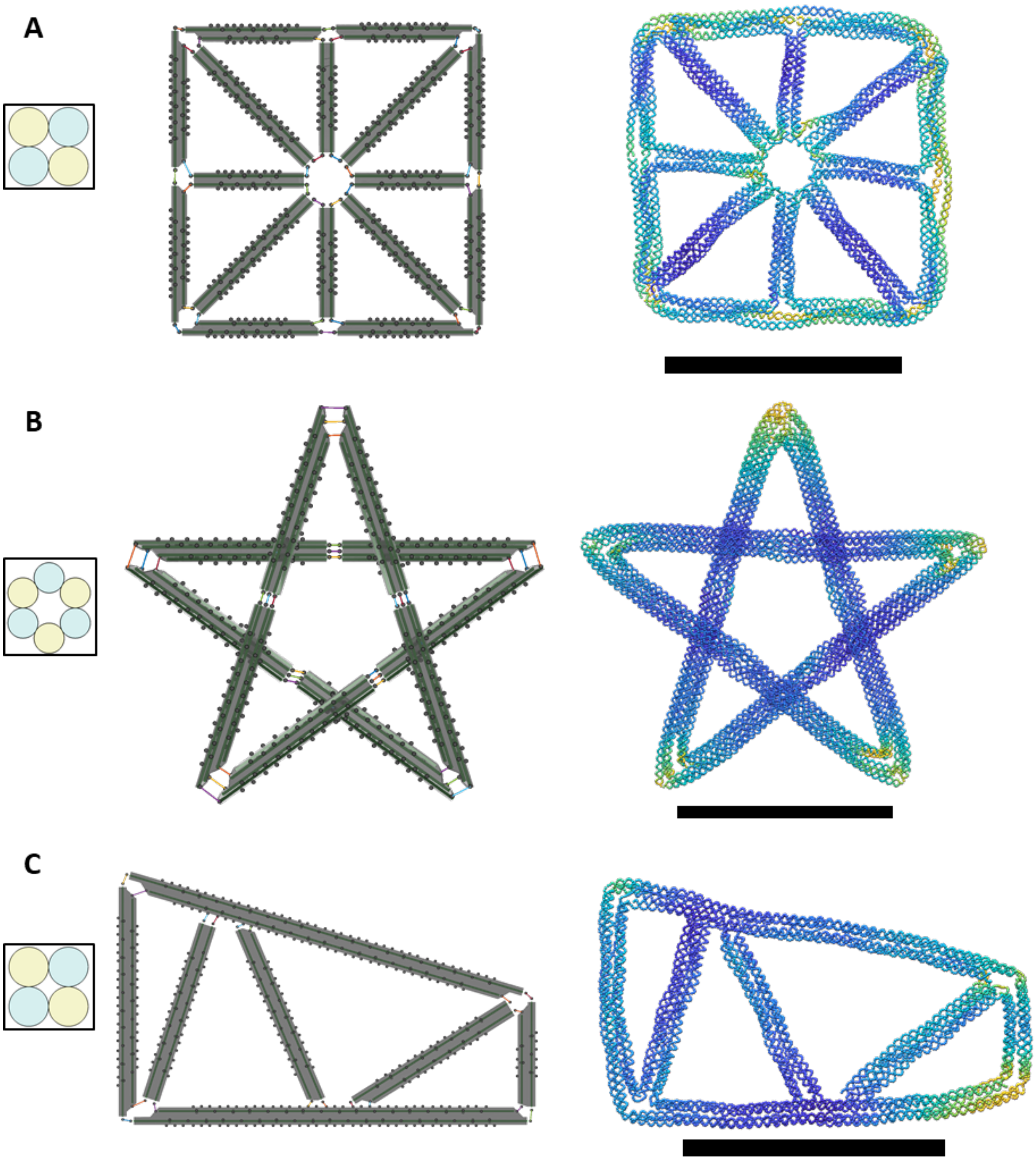

**Supplementary Figure 58: Examples of wireframe structures**. The wireframe modeling approach for creating large structures is based on using relatively long structures but with small cross-sections like 2×2 and 6-HB. (A) A triangular grid with 2×2 cross-sections with extended cylinders at the vertices with the bundle editing GUI. Scaffold length = 5768 nt. (B) A five-star structure with ten 6-HB bundles where five side-to-side double-scaffold crossovers (layered crossovers) constrain the out-of-plane shape. Scaffold length = 7800 nt. (C) The wing in the airplane design. Scaffold length = 3955 nt. Scale bars = 50 nm.

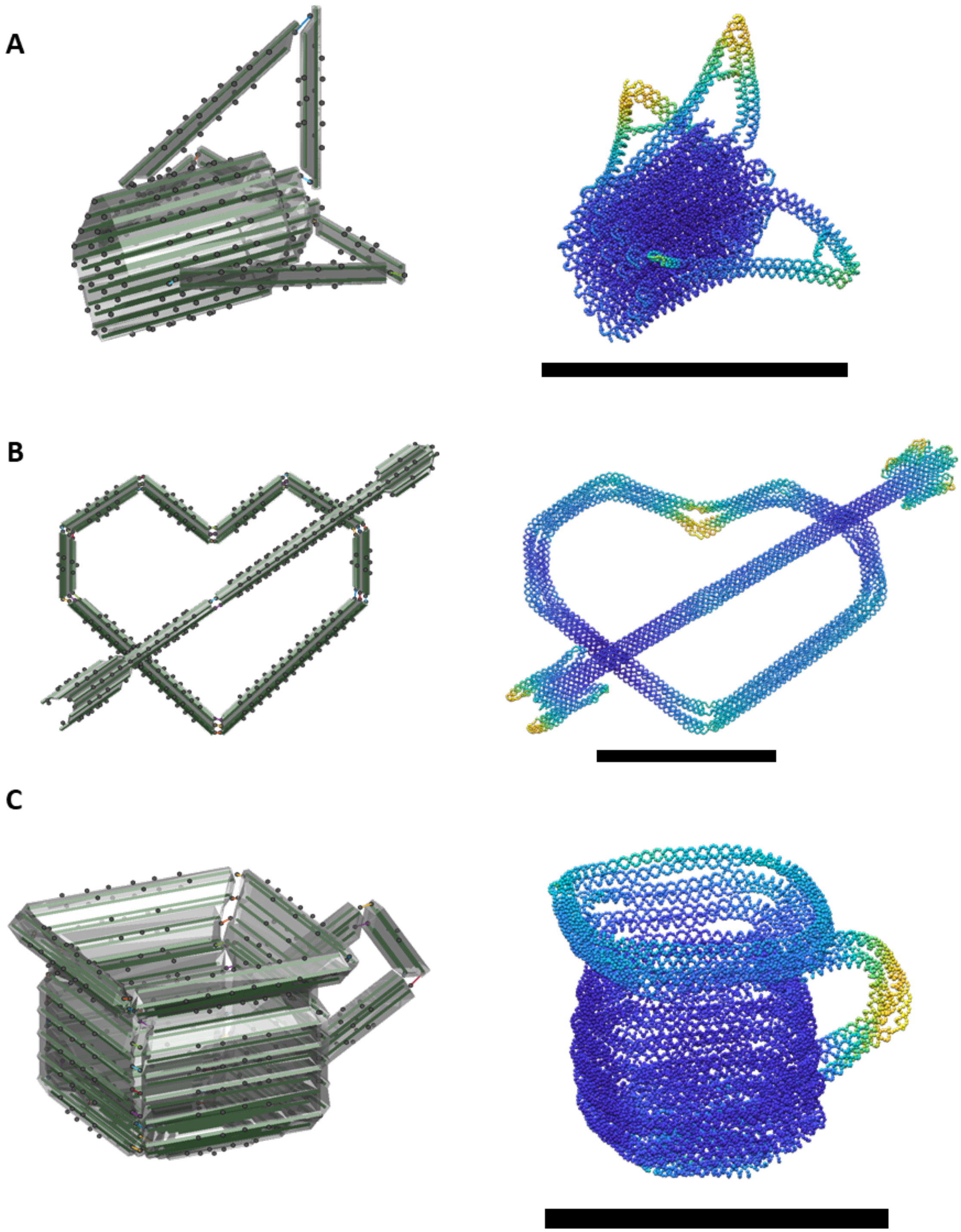

**Supplementary Figure 59: Examples of hybrid structures.** (A) The tail of the airplane design with lattice body and wireframe tail wings. Scaffold length = 5882 nt. (B) A design of the symbol of the Cupid heart with a lattice-based arrow and wireframe heart. Scaffold length = 8148 nt. (C) A mug design with surface-based cup and curved wireframe handle. Scaffold length = 8334 nt. Scale bars = 50 nm.

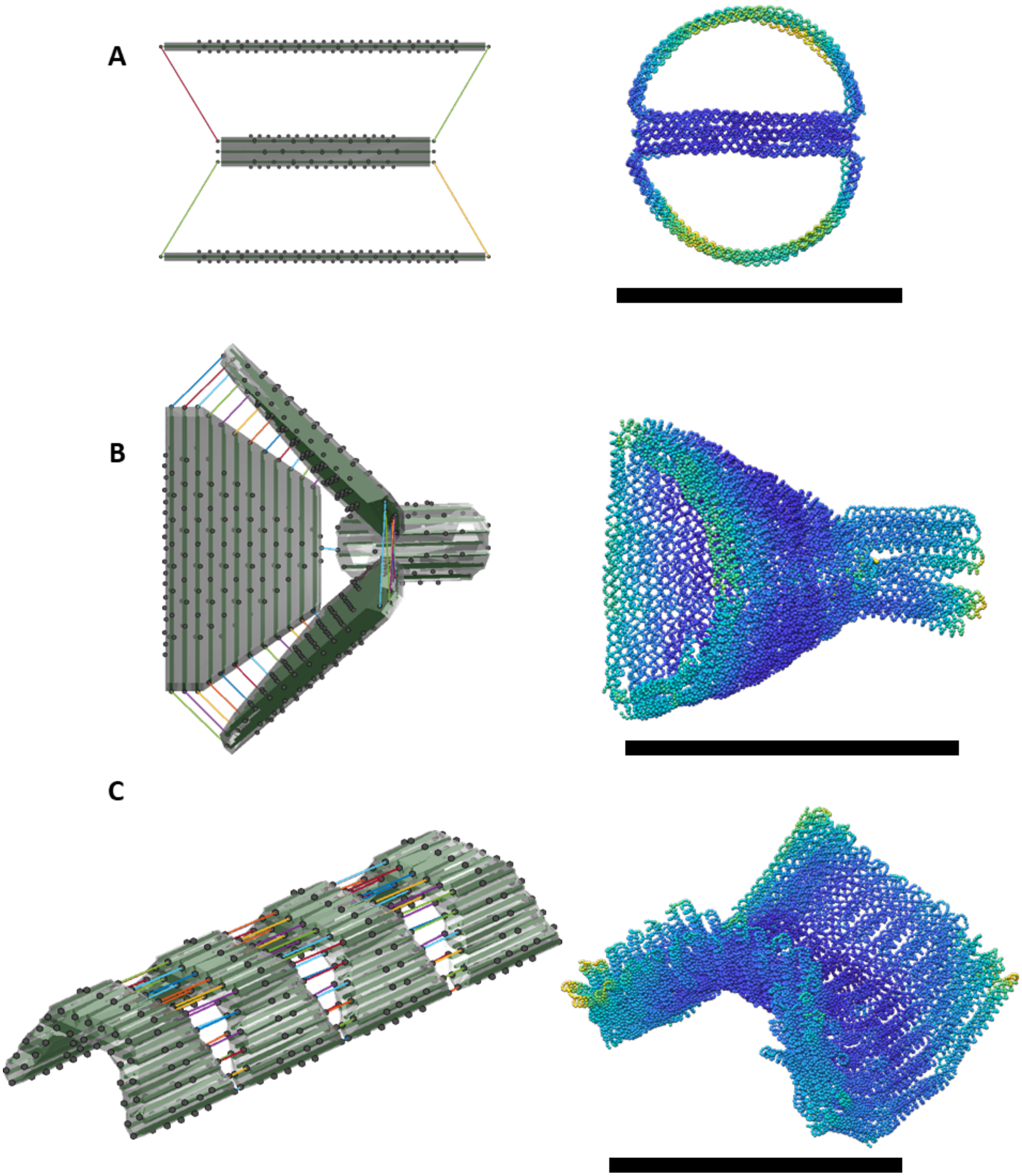

**Supplementary Figure 60: Other complex design examples.** (A) A wheel design with lattice-based frame and curved wireframe profile. Scaffold length = 2872 nt. (B) A design with surface-based funnel and lattice-based central pore. Scaffold length = 8085 nt. (C) A surface-based saddle design. Scaffold length = 13520 nt. Scale bars = 50 nm.

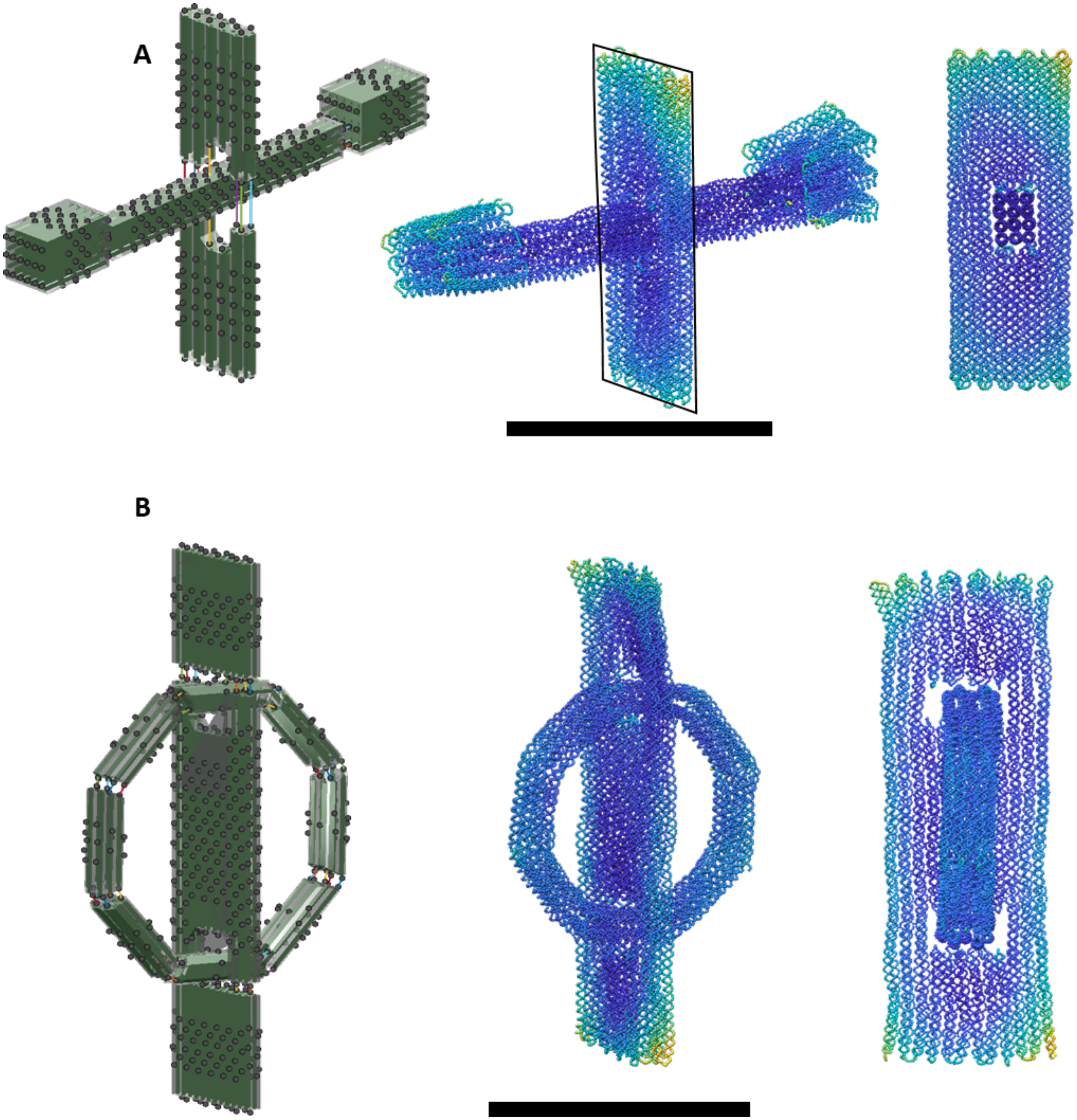

**Supplementary Figure 61: Examples of designs with interlocking features.** (A) A design of a plate with an interlocked bar. Scaffold length = 11226 nt. (B) A design of a plate with an interlocked ring. Scaffold length = 10910 nt. Scale bars = 50 nm.

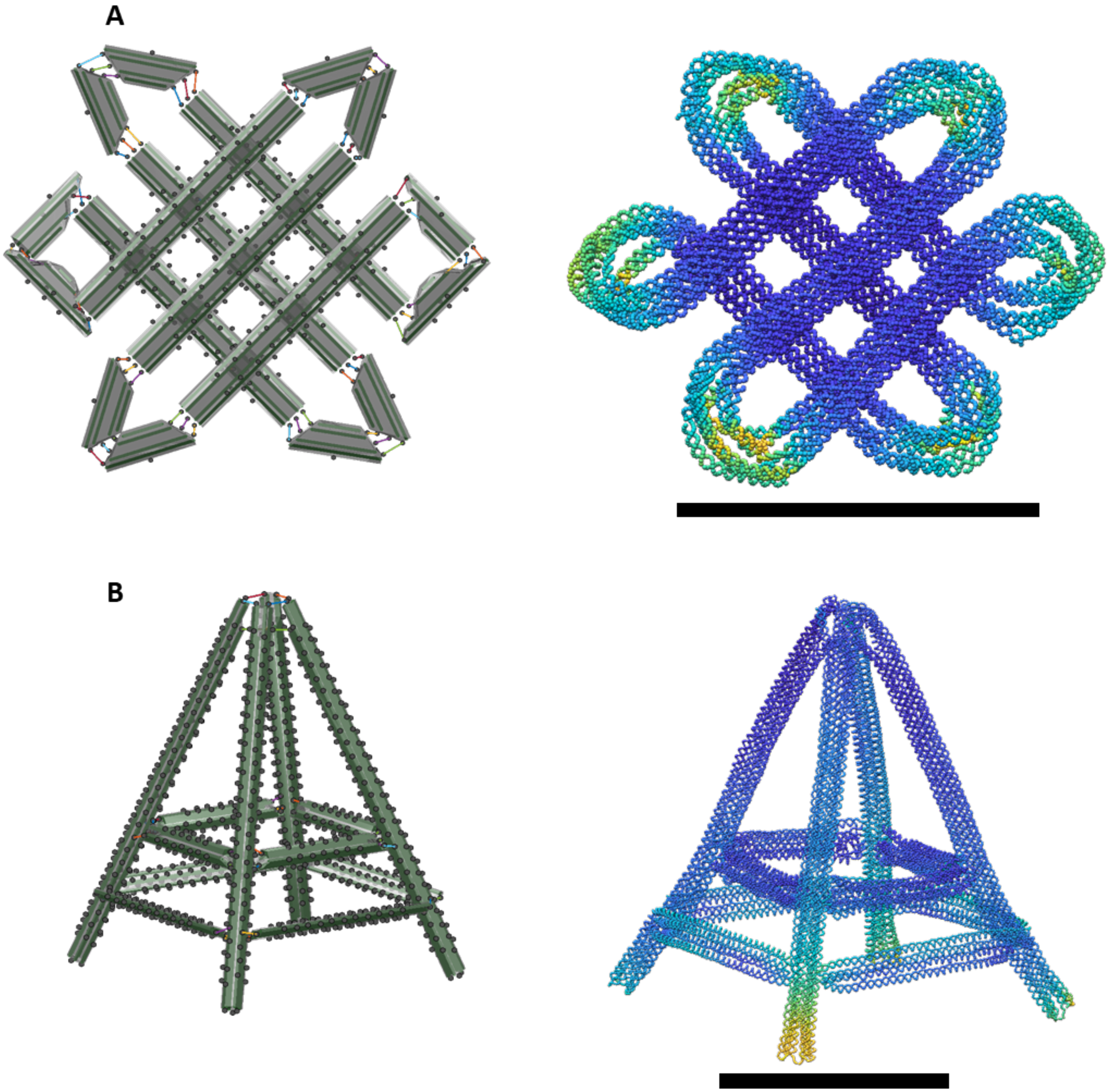

**Supplementary Figure 62: Examples of wireframe structures with complex 3D arrangement and connectivity.** (A) A design of Chinese knot where the cylinders in 45º and -45º are in different layer and connected by external double-scaffold crossovers. Scaffold length = 7416 nt. (B) A tower design. Scaffold length = 12256 nt. Scale bars = 50 nm.

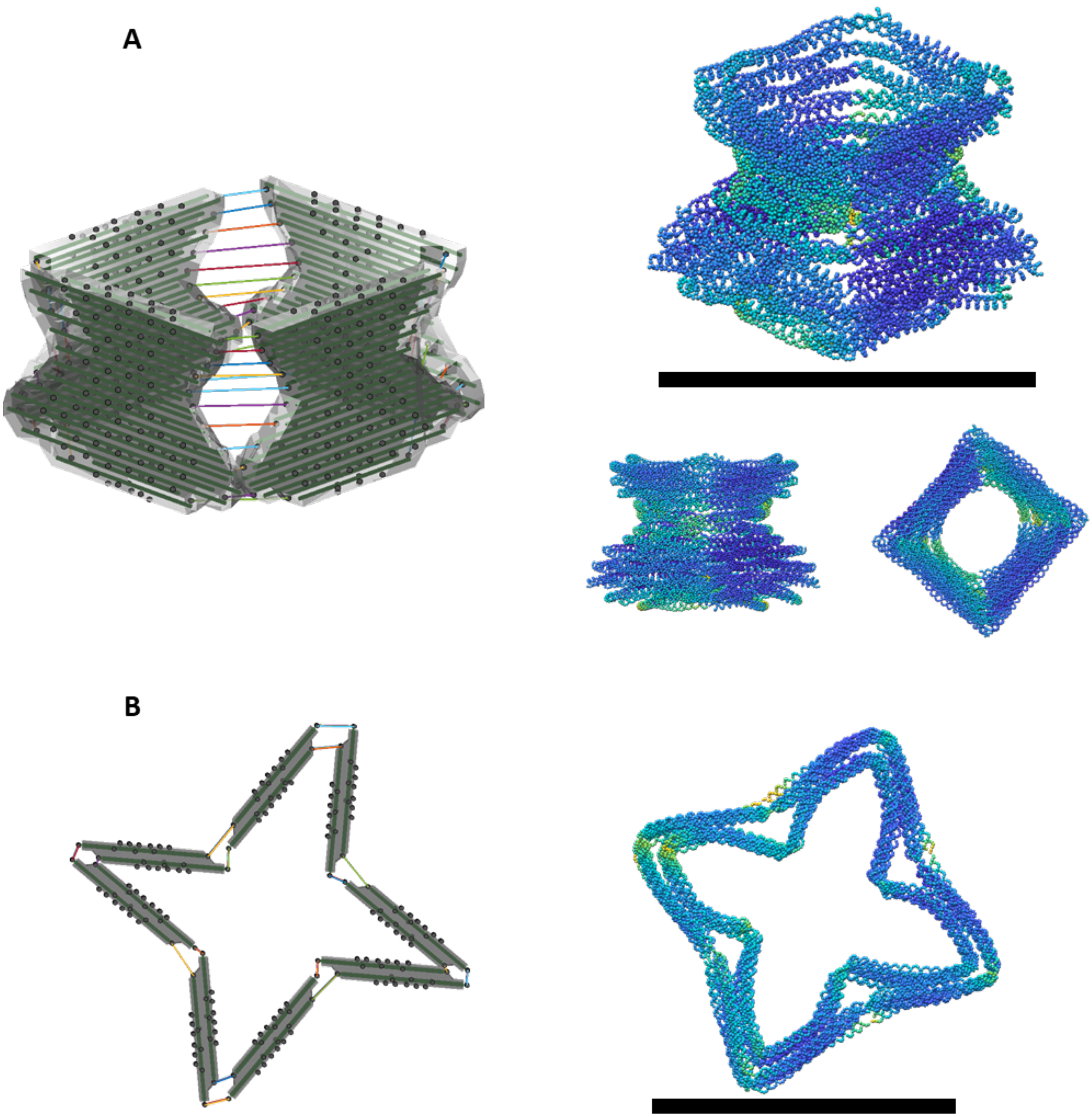

**Supplementary Figure 63: Other examples.** (A) A vase design. Scaffold length = 7440 nt. (B) A design of a 4-pointed ninja star. Scaffold length = 6504 nt. Scale bars = 50 nm.

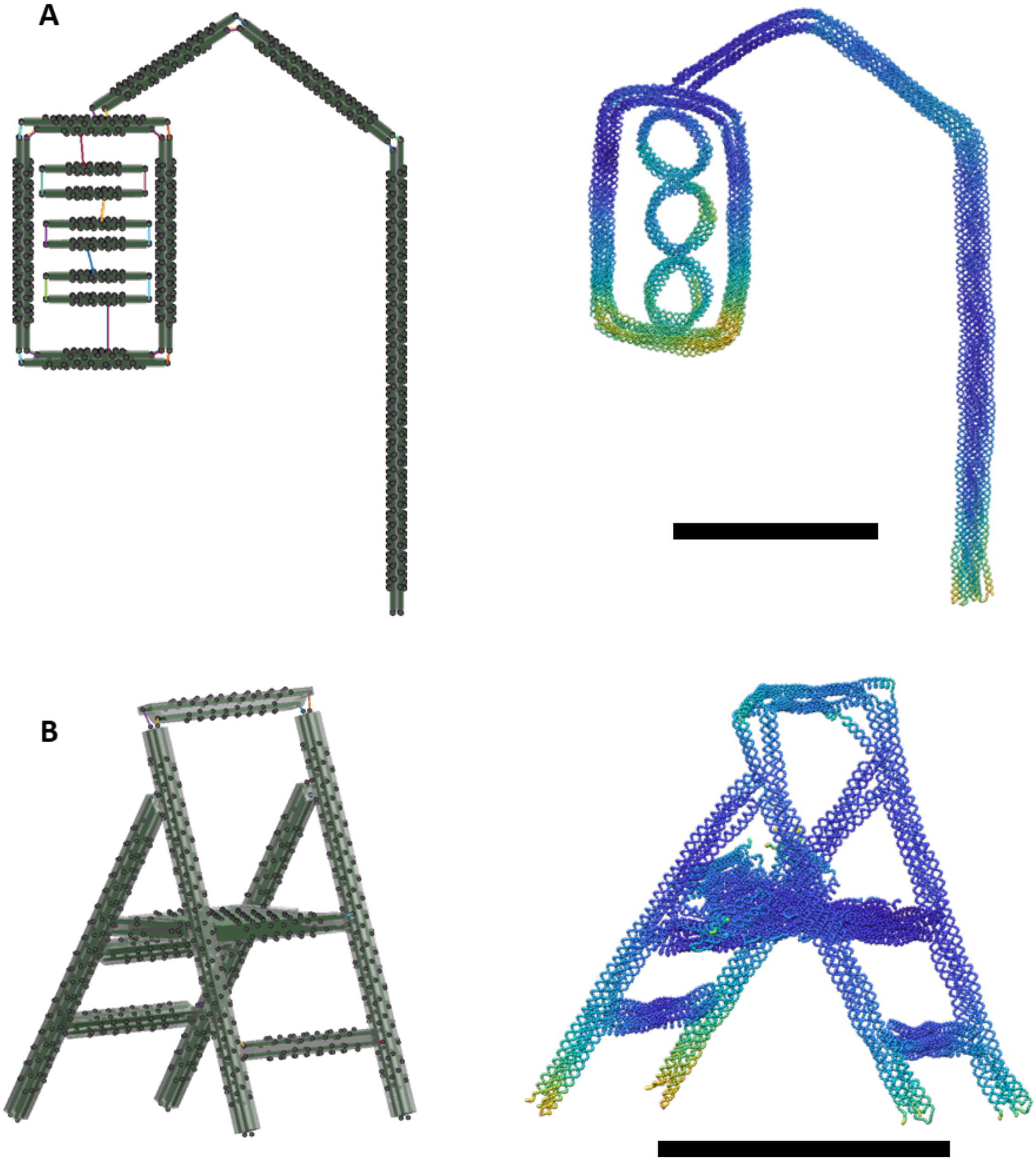

**Supplementary Figure 64: Other examples.** (A) A design of a traffic light. Scaffold length = 10836 nt. (B) A design of a foldable stand. Scaffold length = 7868 nt. Scale bars = 50 nm.

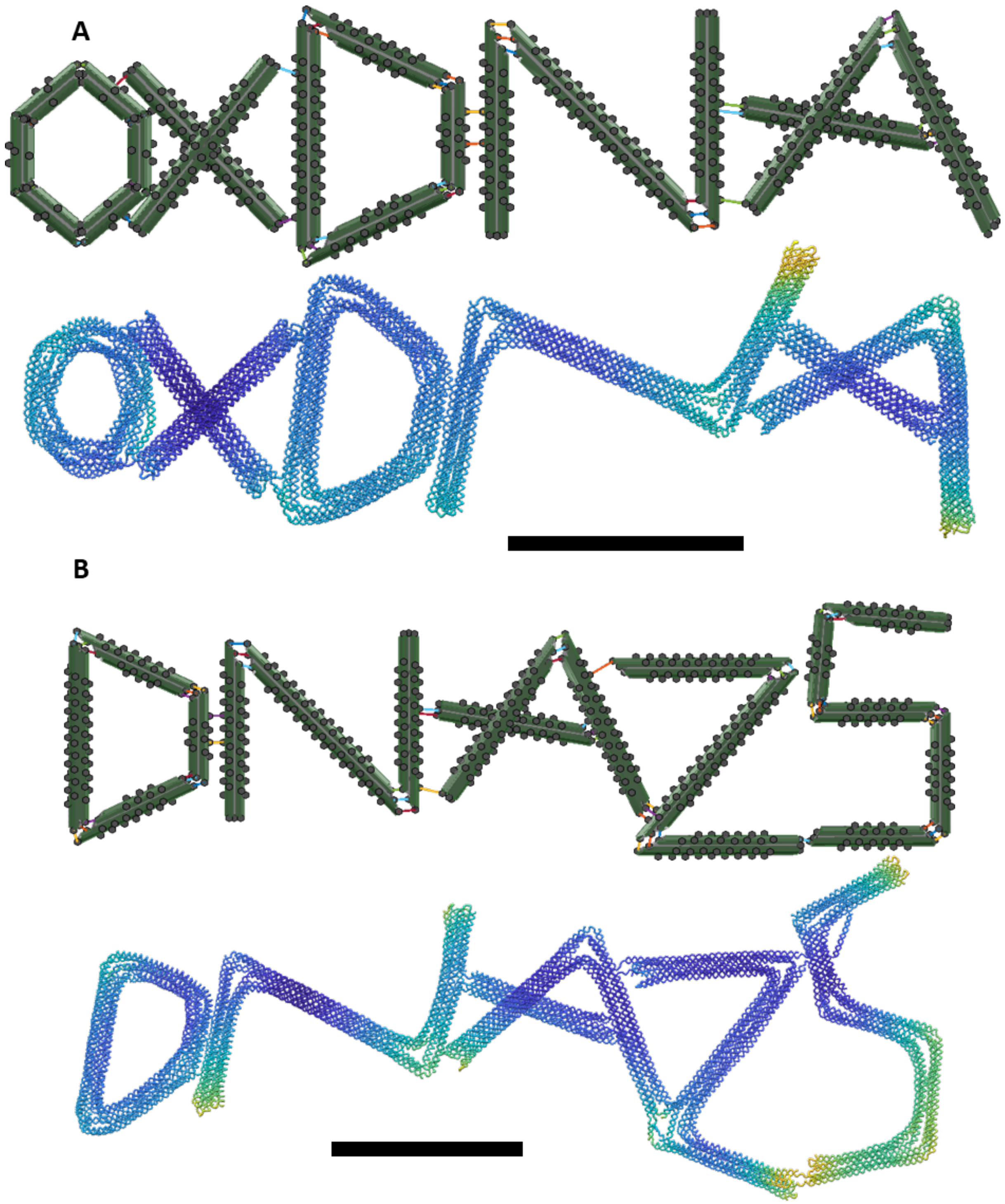

**Supplementary Figure 65: Examples of written designs.** (A) A design of script "oxDNA". Scaffold length = 10670 nt. (B) A design of script "DNA25". Scaffold length = 12610 nt. Scale bars = 50 nm.

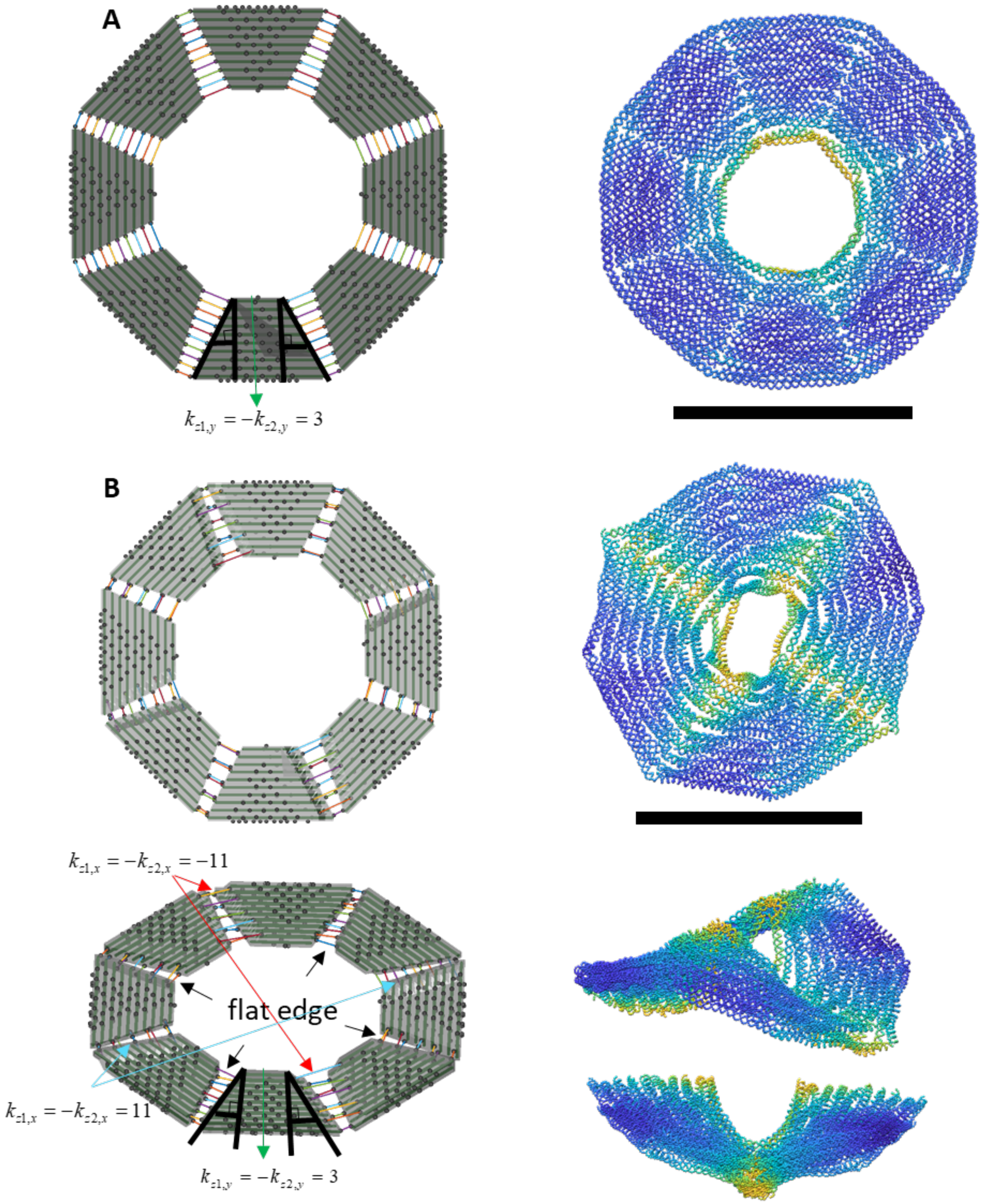

**Supplementary Figure 66: Other examples.** (A) A design of a flat disc by 8 bundles. Scaffold length = 10544 nt. (B) A design of a curved flipping disc. Scaffold length = 10226 nt. Scale bars = 50 nm.

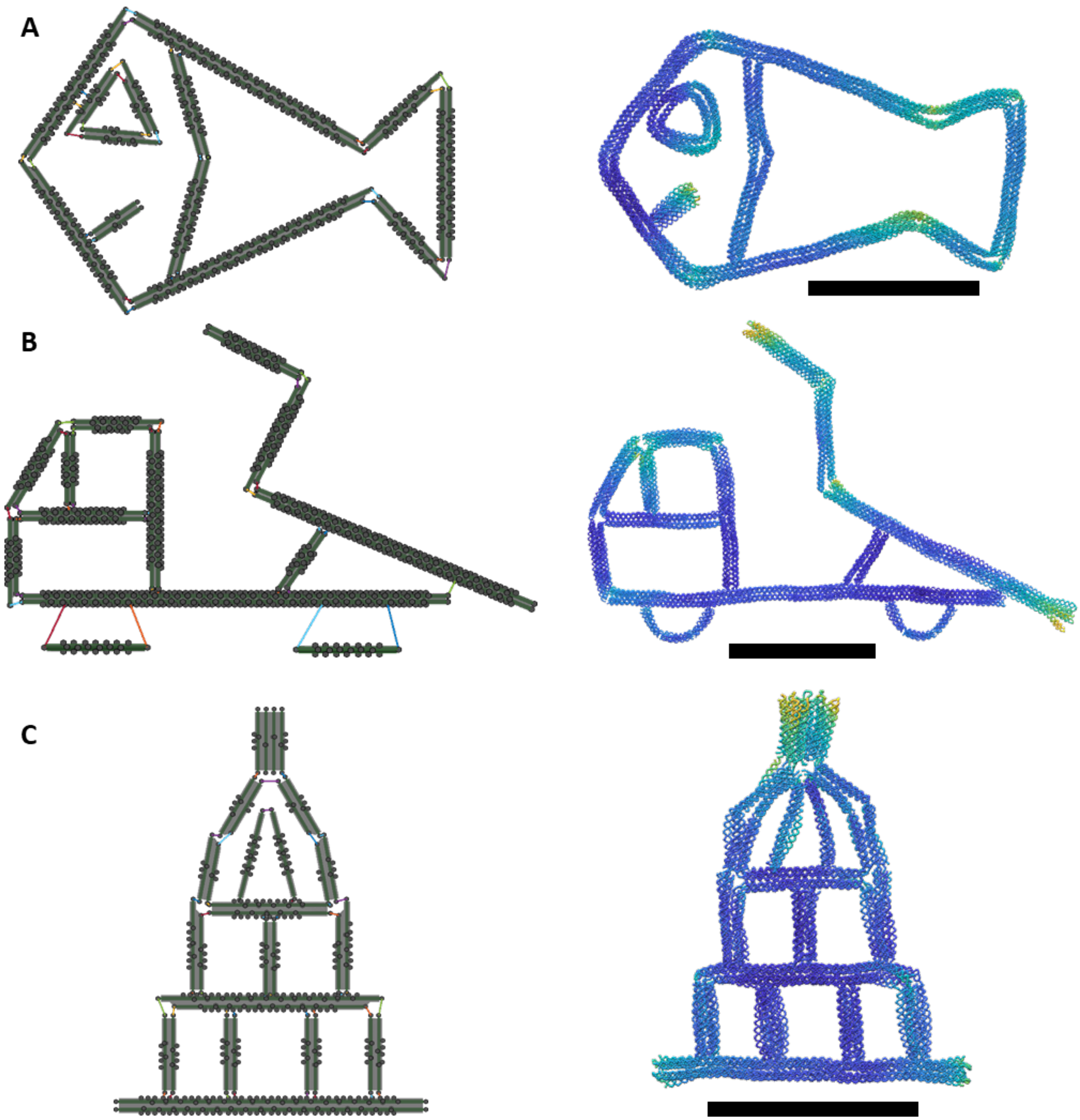

**Supplementary Figure 67: Other examples.** (A) A design of fish. Scaffold length = 11102 nt. (B) A design of a dump truck. Scaffold length = 13230 nt. (C) A design of a capital building. Scaffold length = 9804 nt. Scale bars = 50 nm.

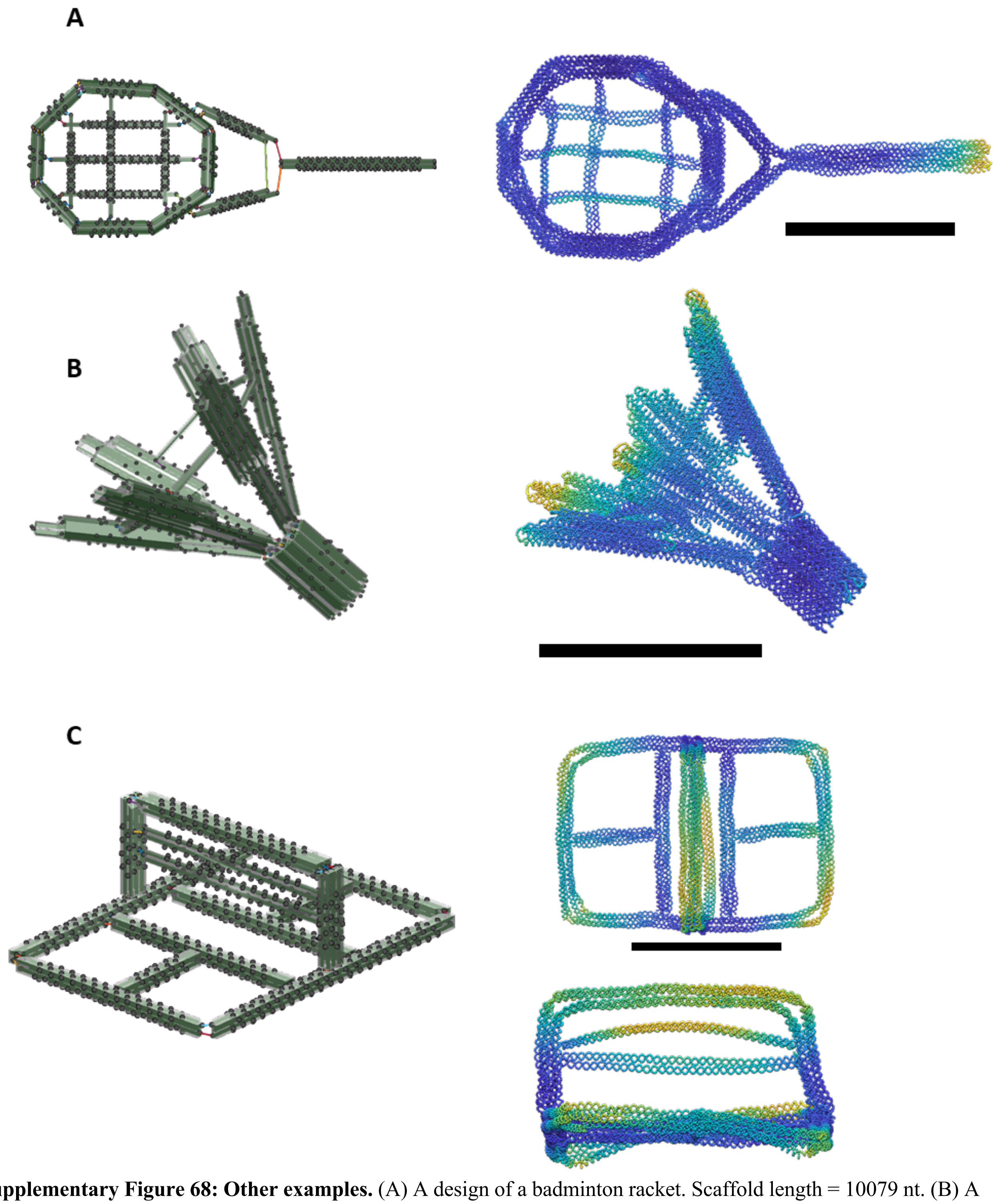

**Supplementary Figure 68: Other examples.** (A) A design of a badminton racket. Scaffold length = 10079 nt. (B) A design of shuttlecock. Scaffold length = 11130 nt. (C) A design of badminton court. Scaffold length = 8118 nt. Scale bars = 50 nm.

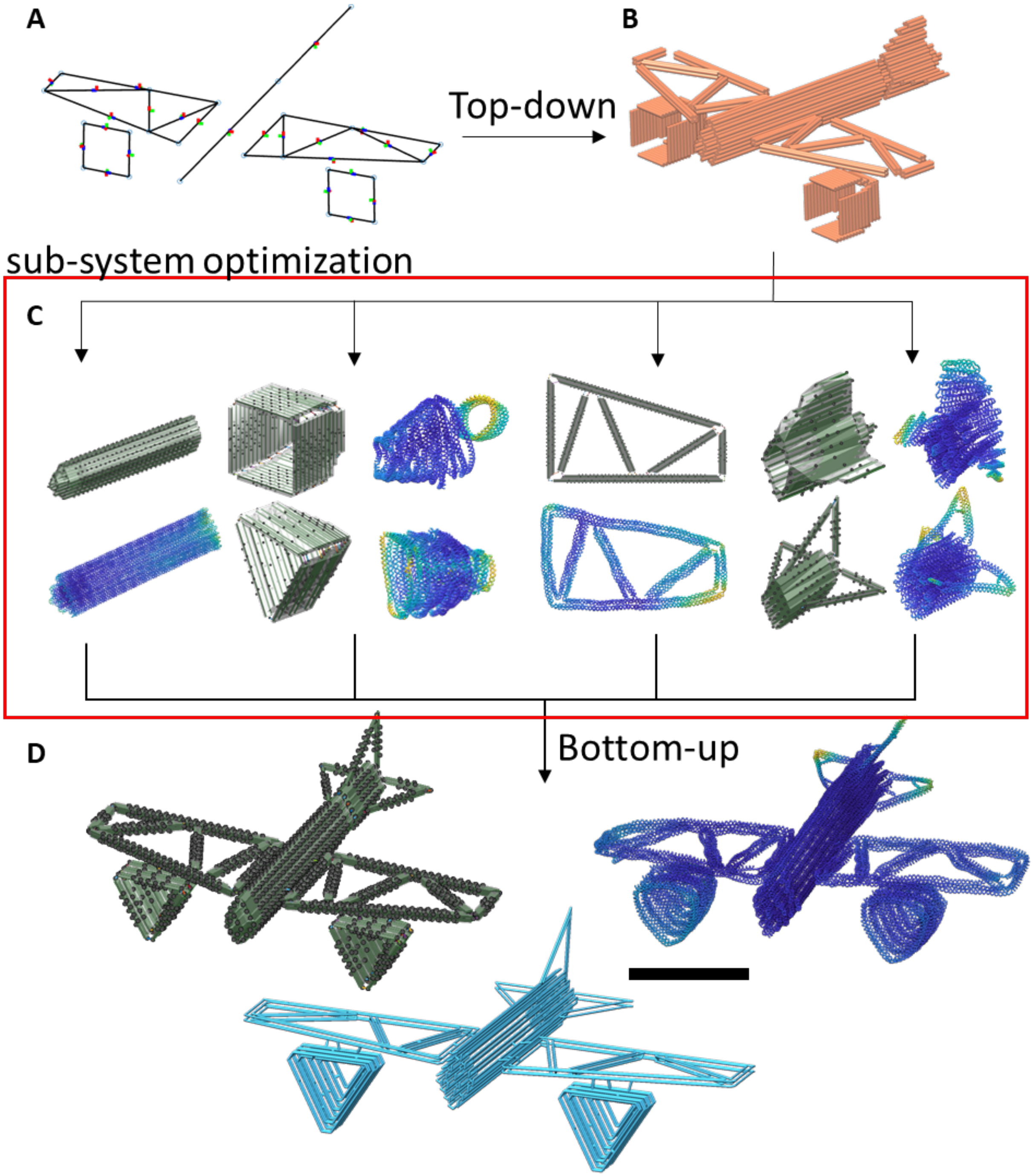

**Supplementary Figure 69: The hybrid hierarchical design process with sub-system optimization starting with top-down initial design, followed by simulation guided iteration of sub-systems, and then bottom-up integration into the larger assembly.** (A) The airplane design is the most complex structure in this study. The top level design strategy is to first apply the top-down approach in which the system is divided into five sub systems: one fuselage, two wings and two turbines. A 3D sketch with 24 lines was created in CAD software and imported into MagicDNA. (B) Each sub system was embodied from a line model into bundles of cylinders. (C) Each sub-system is individually optimized through iterative design process leveraging simulation feedback. (D) Finally, the bottom-up approach was applied to assemble each optimized sub-system into the final design. In this step, each sub-system was imported and manipulated to a desired position and orientation. Also, the connectivity between sub-systems was defined and edited and then the routing algorithms were executed to obtain a single-scaffold routing. The overhang design tool was used to constrain the angles of the tail wings. Scaffold length = 32800 nt. Scale bar = 50 nm.

**Supplementary Section 6: Multi-scaffold and modular designs**

We extended the design process and features of MagicDNA to enable design of multi-scaffold DNA origami following on recent progresses in hierarchical assembly and orthogonal scaffolds[8]. We implemented two approaches in MagicDNA specific for design of multi-scaffold assemblies: (1) Specifying defined interfaces between scaffolds by collecting additional user-inputs for scaffold routings to steer the spanning tree algorithm. (2) Split a single looped scaffold into *K* by searching and applying *K*-1 crossovers with length constraints on the loop sizes.

For the first approach, we created GUIs for users to control the adjacency matrix and transformed the spanning tree algorithm to a spanning forest with multiple trees. This can be achieved in multiple ways. The first is by deactivating crossovers along two parallel cylinders in the bundle editing GUI (Figs. S70B) to force the adjacency matrix to be dispersed into two groups of nodes/cycles. Hence, the spanning forest algorithm returns two trees which represent two scaffold routings by integrating the cycles separately, leaving an interface between the cylinders as specified by the user. In addition to separating two scaffold routings between cylinders, one can also spilt the routings across a component cross-section by specifying a scaffold seam at a slice of the cylinder model (Fig. S71). This list of user-defined internal crossovers is appended to the list of external ones from the assembly GUI, guaranteeing that the final scaffold routings have those user-defined double-crossovers. In order to prevent the integration along the cylinder direction (i.e. finding a double-crossover between those seams), another GUI operation allows users to specify a region over which the internal crossovers are deactivated (Fig. S71B) leading to separation of the spanning trees at the desired interface to create a spanning forest. Other examples for this multi-scaffold algorithm are the simultaneous design process for finishing multiple designs in one MagicDNA assembly, such as shape-complementary and sticky-end assembles.

Details of the second approach for multi-scaffold routing are described in Fig. S81.

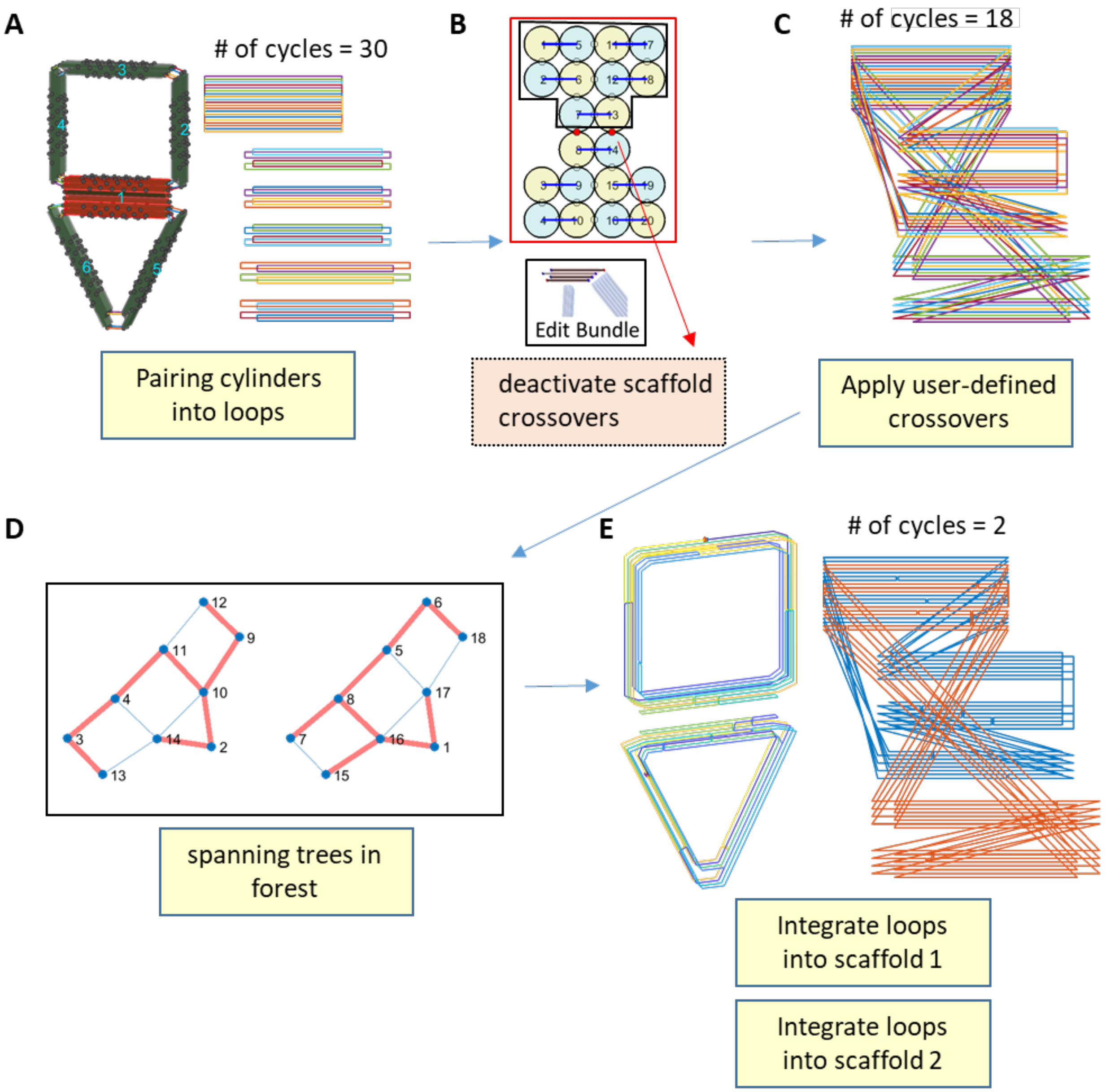

**Supplementary Figure 70: The scaffold algorithm for a multi-scaffold design with defined interface for the case where scaffold routings are spilt between cylinders in a bundle.** (A) Pairing the cylinders when the mechanism is initially generated. (B) In the bundle editing GUI, the cross-section view allows users to deactivate all scaffold crossovers between cylinders to split the routing. (C) Applying external crossovers from the result of the assembly GUI. (D) Compared with Fig. S11, the adjacency matrix between cycles is intentionally split at the user-defined interface between layers of a bundle (i.e. between cylinders 7&8 and 13&14 in the bundle 1), thereby creating two spanning trees in a spanning forest. (E) Each spanning tree guides the integration of cycles into one scaffold routing, and the full routing is visualized in both 2D and 3D representations.

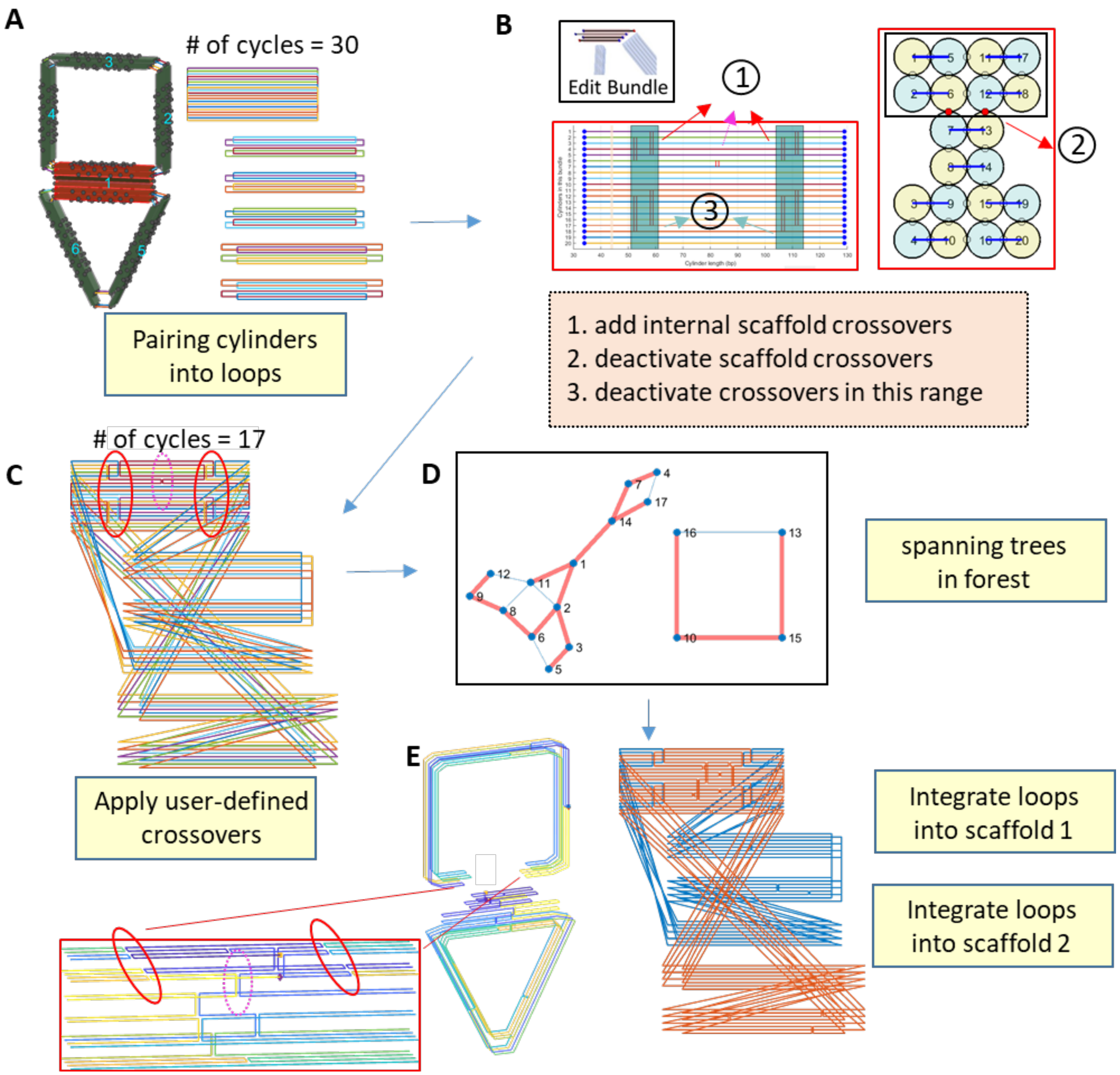

**Supplementary Figure 71: The scaffold algorithm for a multi-scaffold design with defined interface for the case where users form a lock-and-key type interface that cuts across and between cylinders.** (A) Pairing the cylinders when the mechanism is generated. (B) In the bundle editing GUI, in addition to deactivating scaffold crossovers between cylinders, two more operations are introduced to specify the locations of the interface between scaffolds. One is to allow users to specify the internal crossovers (labeled as 1), which split the regions along the helical direction. The other is to deactivate the crossovers in a specific range (labeled as 3, cyan) in order to avoid integration of multiple scaffolds. (C) Applying external crossovers from the result of the assembly GUI and 9 internal crossovers from the bundle editing GUI in (B), 8 of which form two interfaces perpendicular to the cylinder direction (shown in solid red circles) and 1 of which merges the scaffold routing in the bump to the bottom scaffold routing of the triangle (shown in dashed pink circles). (D) After examining the adjacency matrix between these cycles, the two split spanning trees divide two sets of cycles to be integrated separately resulting in a spanning forest with two trees. (E) After the integrations, the routing result is visualized in 2D and 3D representations. Inset shows the user-defined internal crossovers which are the seams and the interface of the two scaffold routings.

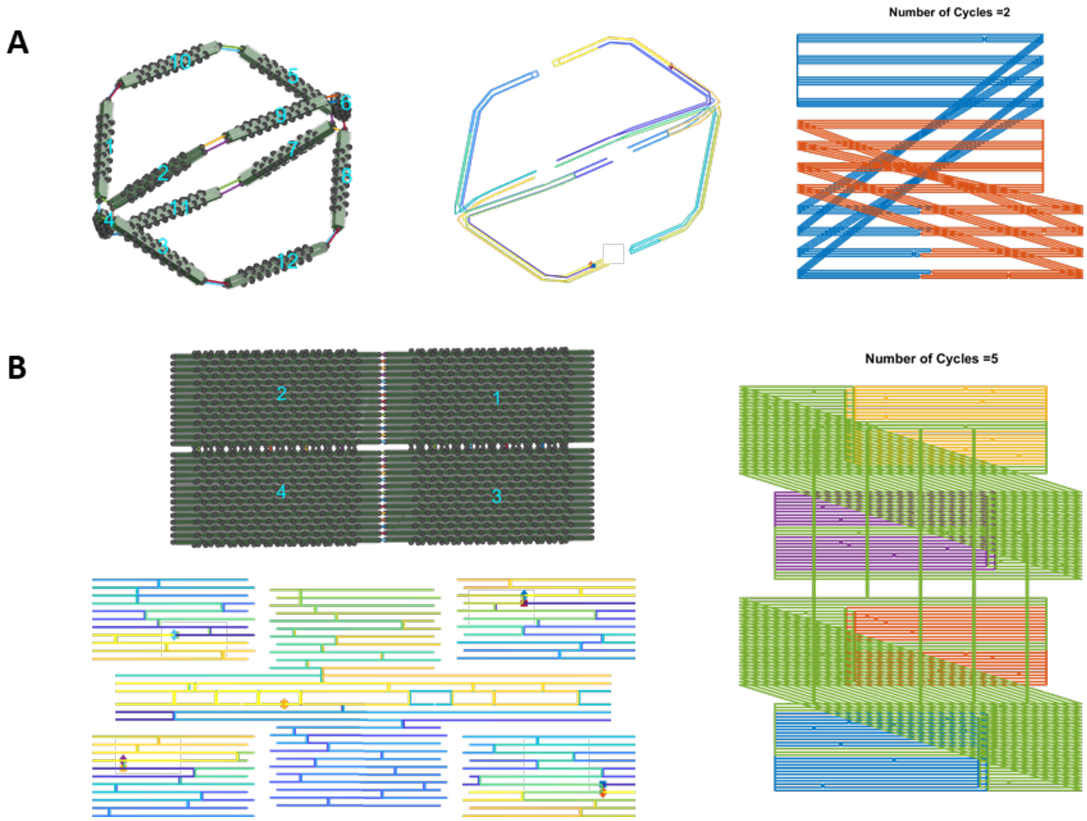

**Supplementary Figure 72: Examples of multi-scaffold routings.** (A) A two-scaffold-routing case of a wireframe ball (interfaces on bundles 9, 10, 11, and 12). (B) A five-scaffold-routing case of a surface-based plate by defining the interfaces in each bundle.

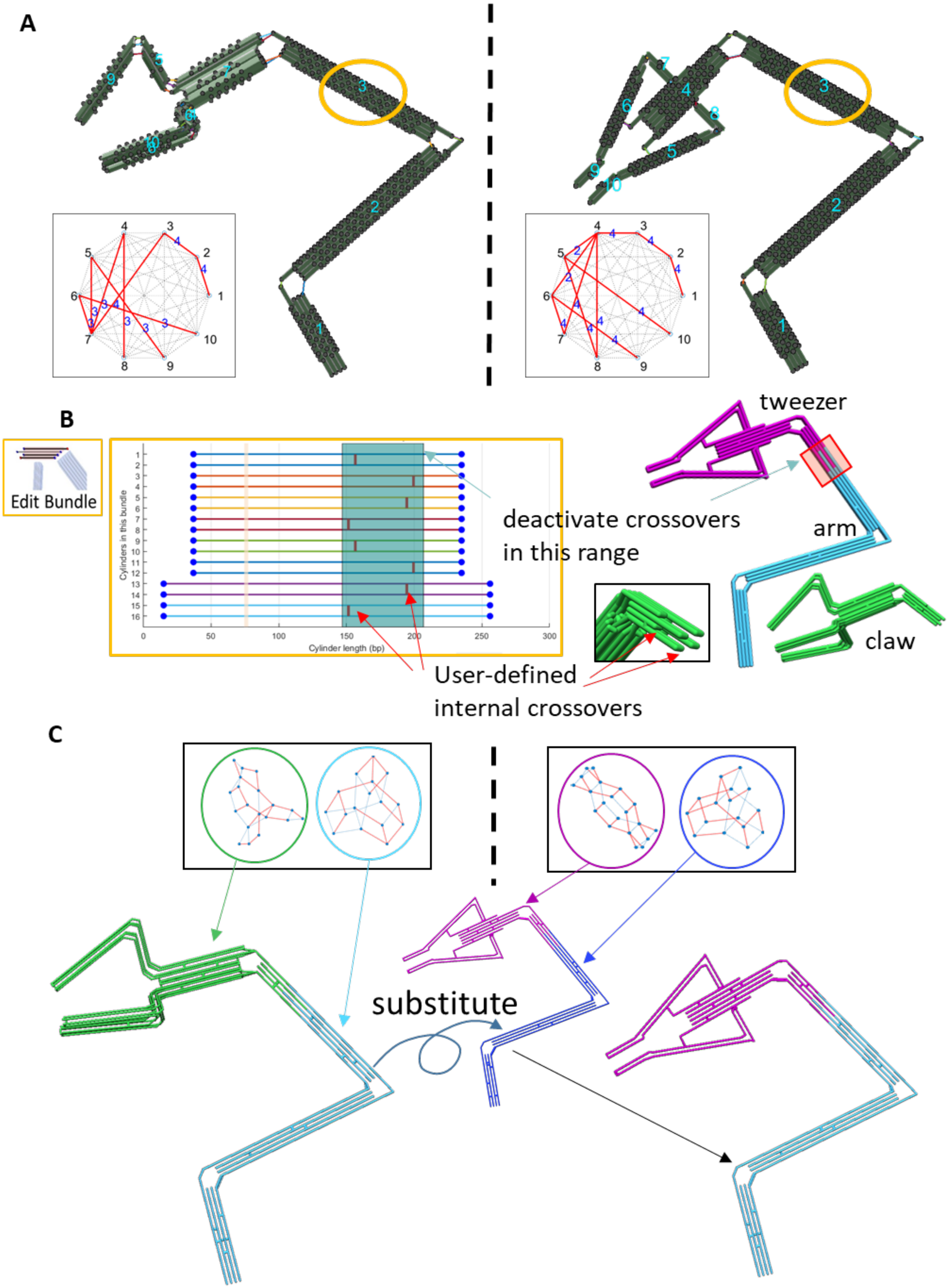
A
B
Edit Bundle
Cylinders in this bundle
Cylinder length (bp)
deactivate crossovers in this range
User-defined internal crossovers
tweezer
arm
claw
C
substitute

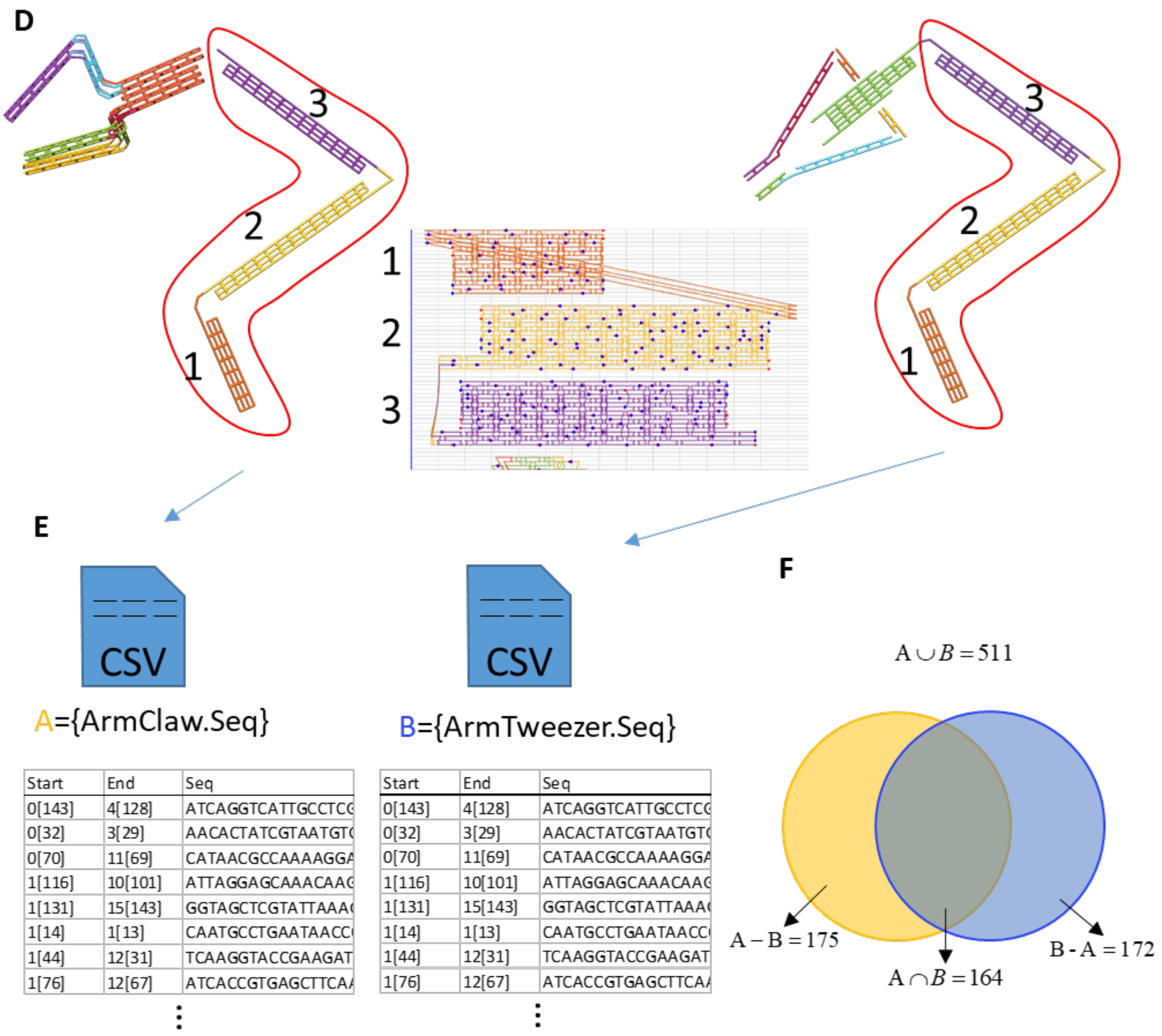

**Supplementary Figure 73: Multi-scaffold design process for the exchangeable robotic manipulator**. (A) After scaling and iterative optimization, two MagicDNA programs were launched side-by-side and the arm bundles (1, 2, and 3) are identical with the same geometries and assembly. (B) Using the bundle editing GUI, the interface of two scaffolds was assigned by specifying the internal crossovers and the region with deactivated crossovers in bundle 3 for both designs. (C) Due to the specified interface, the scaffold algorithm obtained a spanning forest comprising two separated spanning trees, which guided the determination of two scaffold routings. To obtain the same sequences for the three-component robot arm, the scaffold routing of the arm in the Tweezer design was substituted by the one from the Claw design in MATLAB command line. (D) Similarly, the staple routings on the repeated units (bundles 1, 2, 3) have to be the same by duplicating one to the other using a MATLAB script. We provide an example script for these processes for a very similar case (Fig. S80) with the software package. (E) After specifying the sequences of two scaffolds, the staple sequences were exported individually. (F) The sequence set analysis of the two exported files shows the same164 staples are used in both designs to make up most of the arm, and 175 staples are included to form the Claw-Arm or a different set of 172 staples are included to form the Tweezer-Arm.

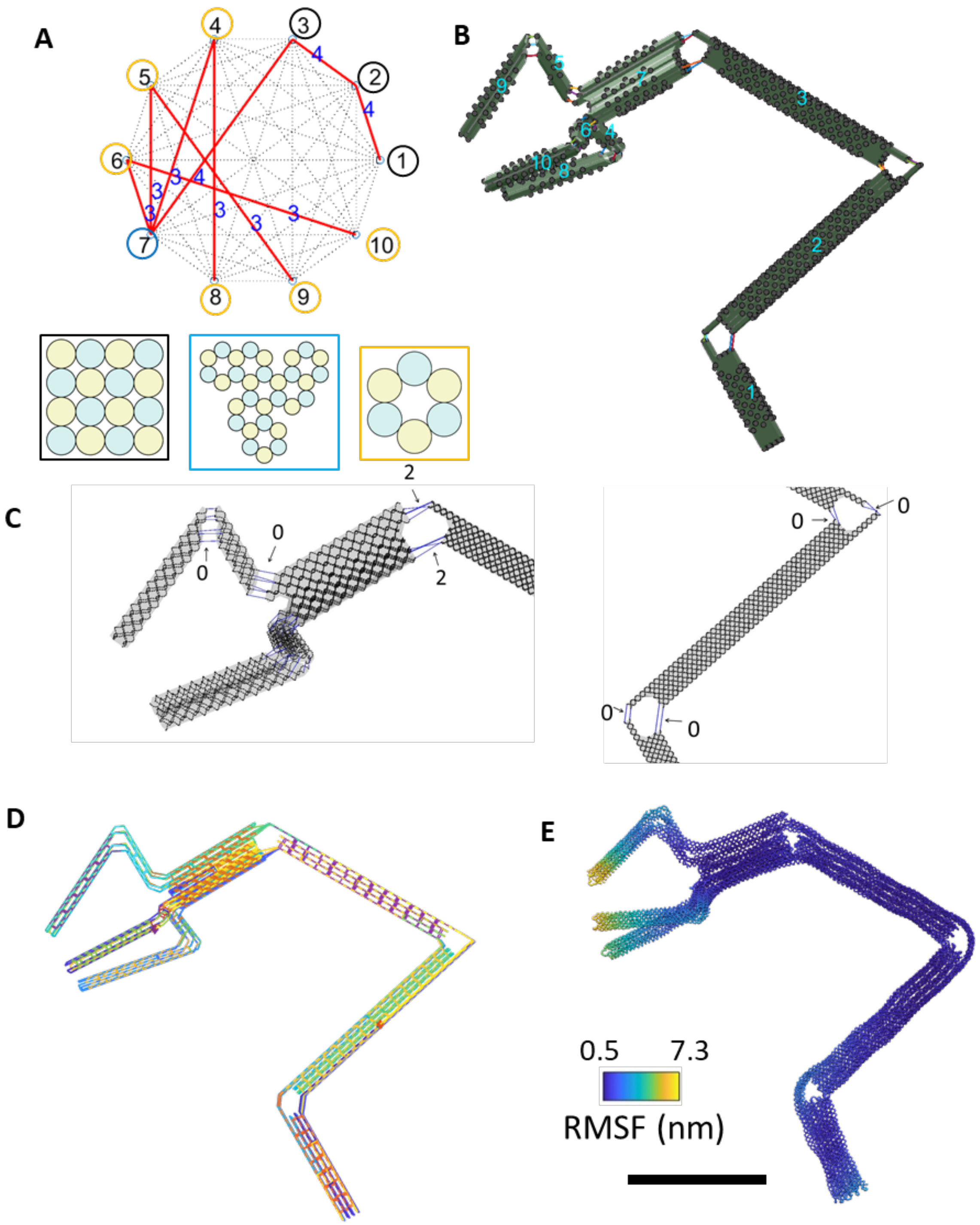

**Supplementary Figure 74: Design details of the Claw-Arm structure.** (A) The connectivity matrix for assembly and the cross-sections of each component. (B) The design model in the MagicDNA assembly panel. (C) The length of scaffold ssDNA connections between individual bundles. (D) The scaffold and staple routing of the final design. (E) The oxDNA simulation result. Scale bar = 50 nm.

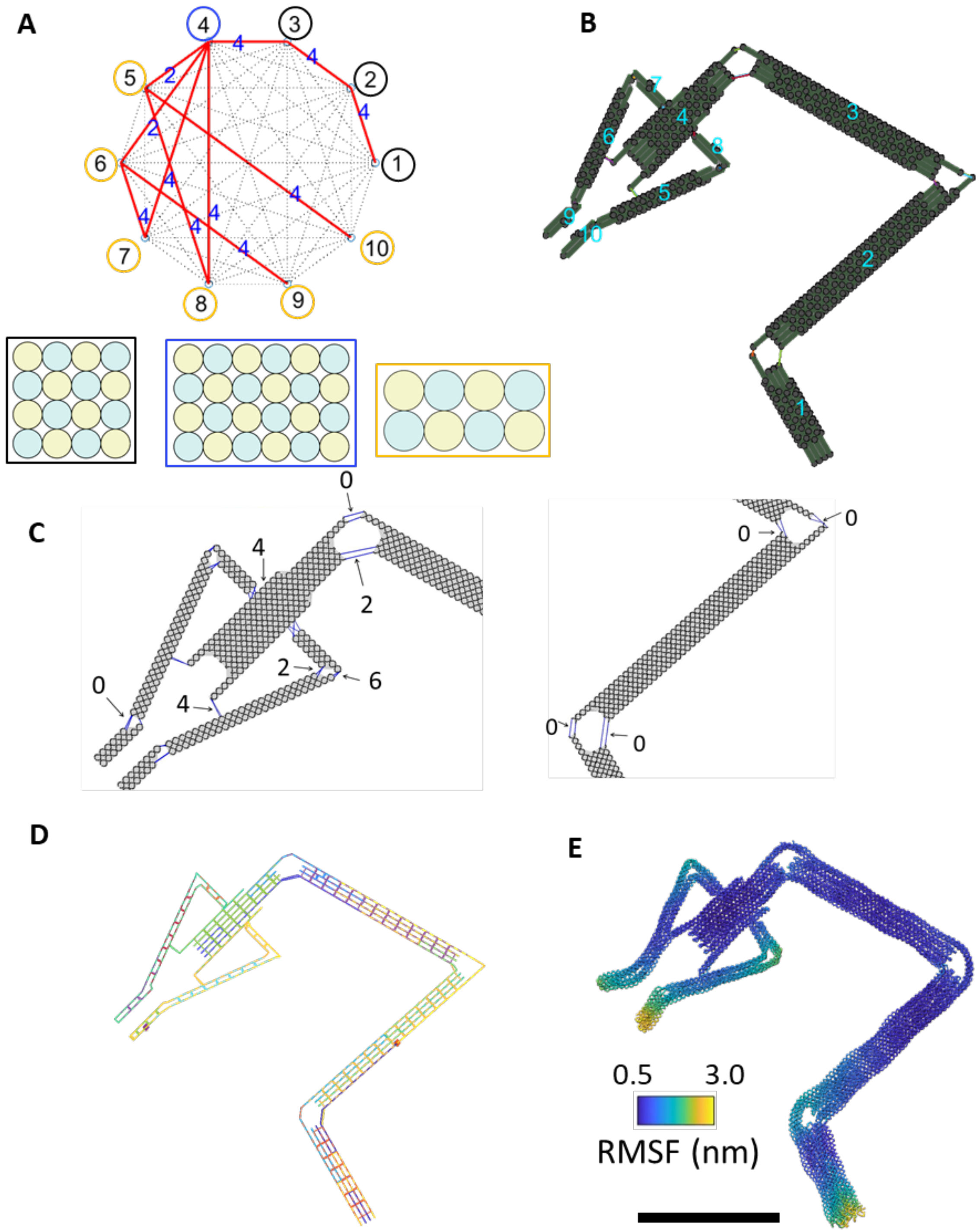

**Supplementary Figure 75: Design details of the Tweezer-Arm structure.** (A) The connectivity matrix for assembly and the cross-sections of each component. (B) The design model in the MagicDNA assembly panel. (C) The length of scaffold ssDNA connections between individual bundles. (D) The scaffold and staple routing of the final design. (E) The oxDNA simulation result. Scale bar = 50 nm.

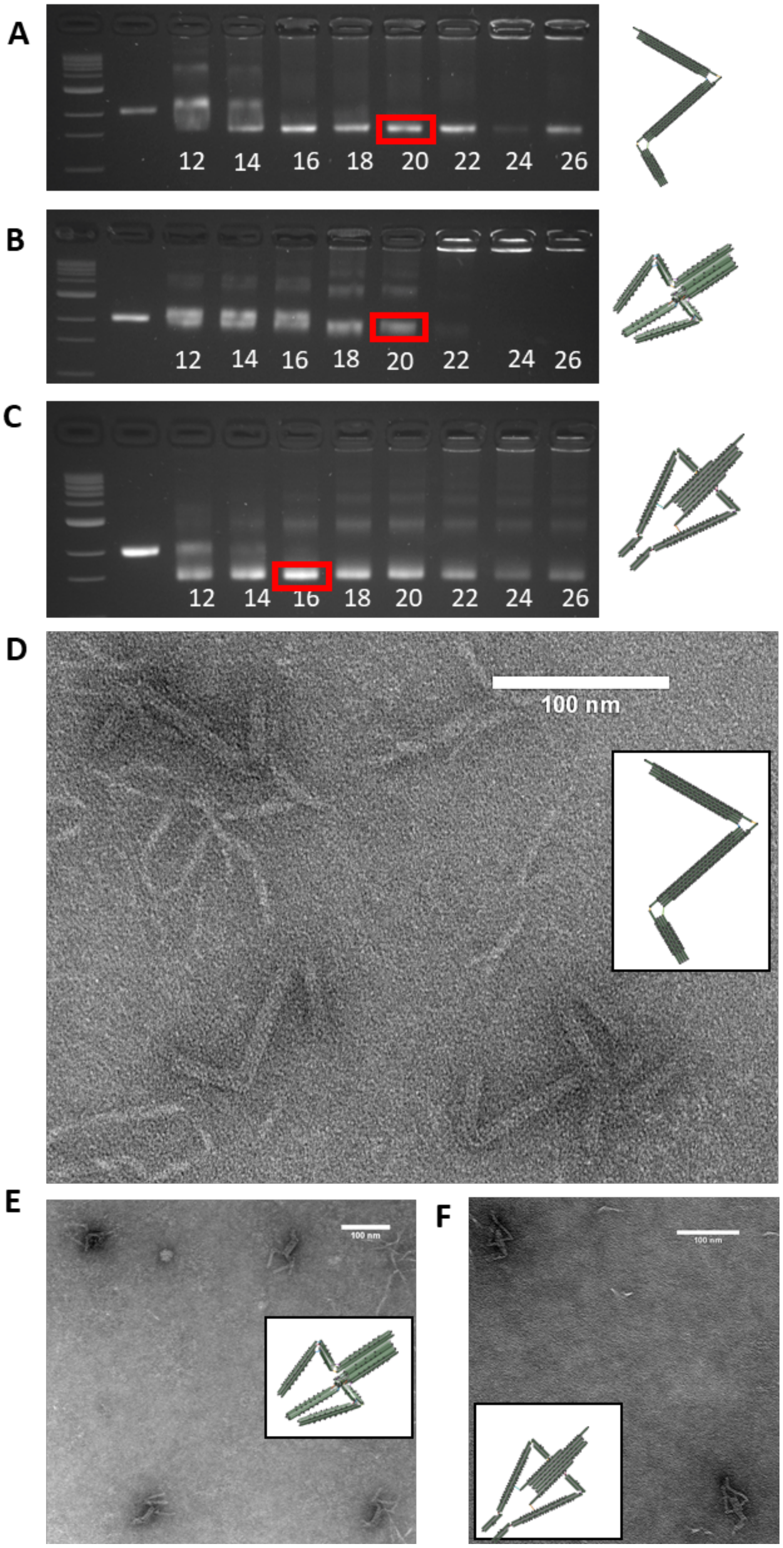

**Supplementary Figure 76: Experimental validation of the components in the robotic manipulator.** (A)-(C) Robot arm, Claw, and Tweezer was folded using a two-and-a-half-day modified thermal annealing protocol while testing a range of magnesium chloride concentrations from 12-26 mM and purified using agarose gel electrophoresis (buffer conditions 0.5xTBE 5.5 mM MgCl2). (D)-(F) Zoomed-out TEM image of gel-purified Robot arm, Claw, and Tweezer folded at 20 mM, 20 mM, and 16 mM respectively MgCl2.

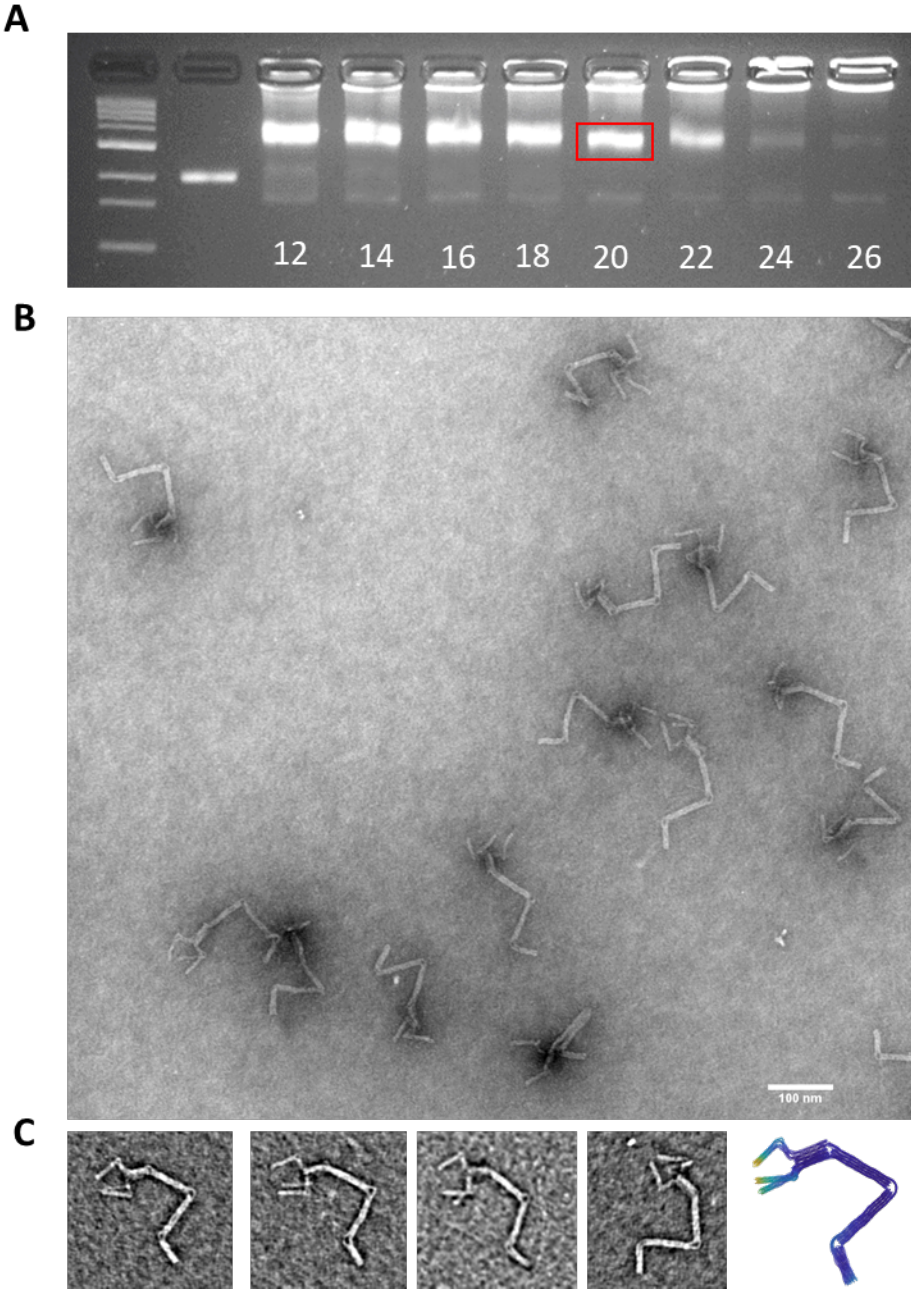

**Supplementary Figure 77: Experimental validation of the robotic manipulator with the claw End-Of-Effector (EOE).** (A) Claw-Arm was folded using a two-and-a-half-day modified thermal annealing protocol while testing a range of magnesium chloride concentrations from 12-26 mM and purified using agarose gel electrophoresis. (B) Zoomed-out TEM image of gel-purified Claw-Arm folded at 20 mM MgCl2. (C) (Left) Zoomed-in TEM images of gel-purified robot arm with claw folded at 20 mM MgCl2. (Right) oxDNA simulation of Claw-Arm.

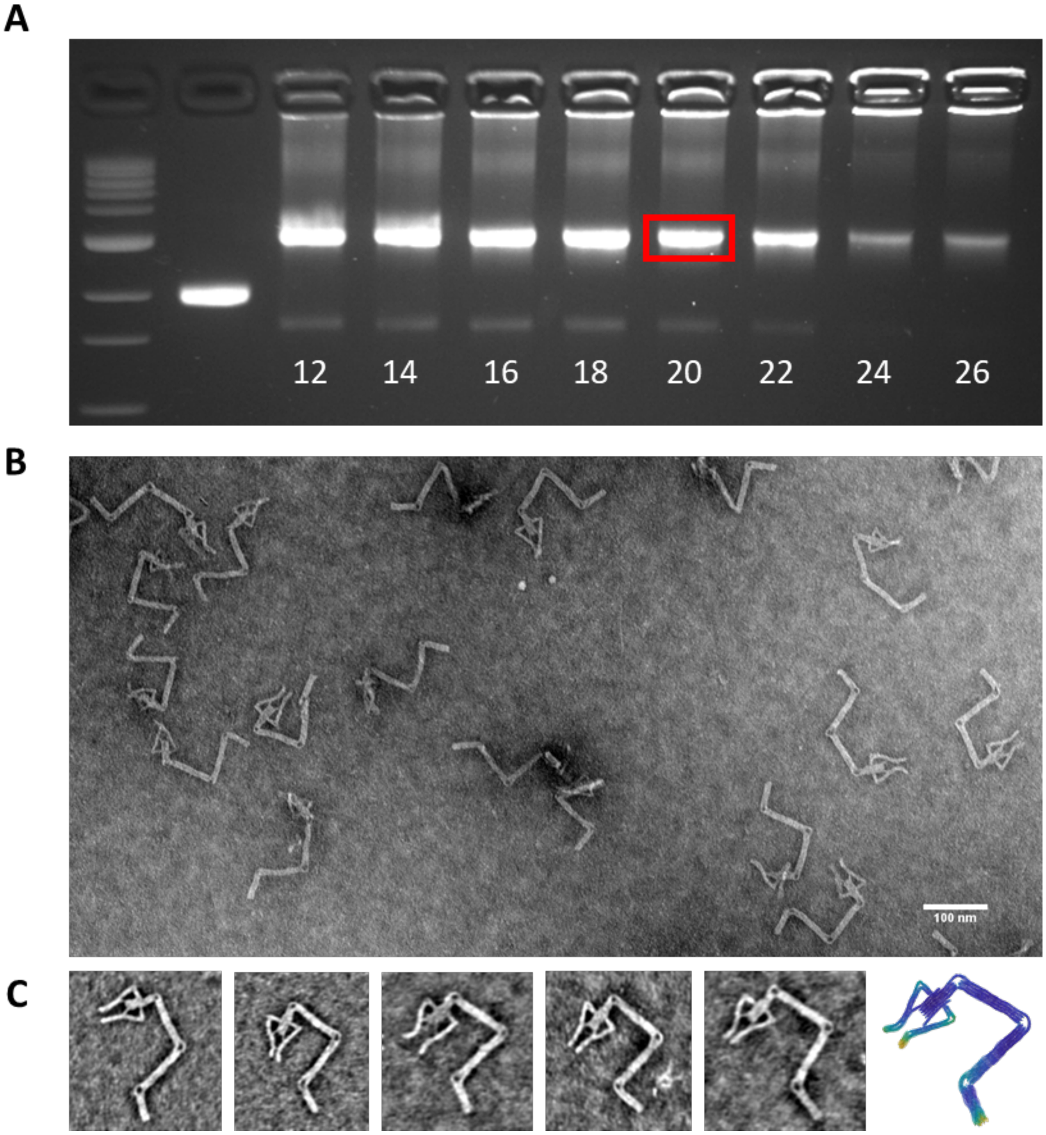

**Supplementary Figure 78: Experimental validation of the robotic manipulator with the tweezer EOE.** (A) Tweezer-Arm was folded using a two-and-a-half-day modified thermal annealing protocol while testing a range of magnesium chloride (mM) concentrations from 12-26 mM and purified using agarose gel electrophoresis (buffer conditions 0.5xTBE 5.5 mM MgCl2). (B) Zoomed-out TEM image of gel-purified Tweezer-Arm folded at 20 mM MgCl2. (C) (Left) Zoomed-in TEM images of gel-purified Tweezer-Arm folded at 20 mM MgCl2. (Right) oxDNA simulation of Tweezer-Arm.

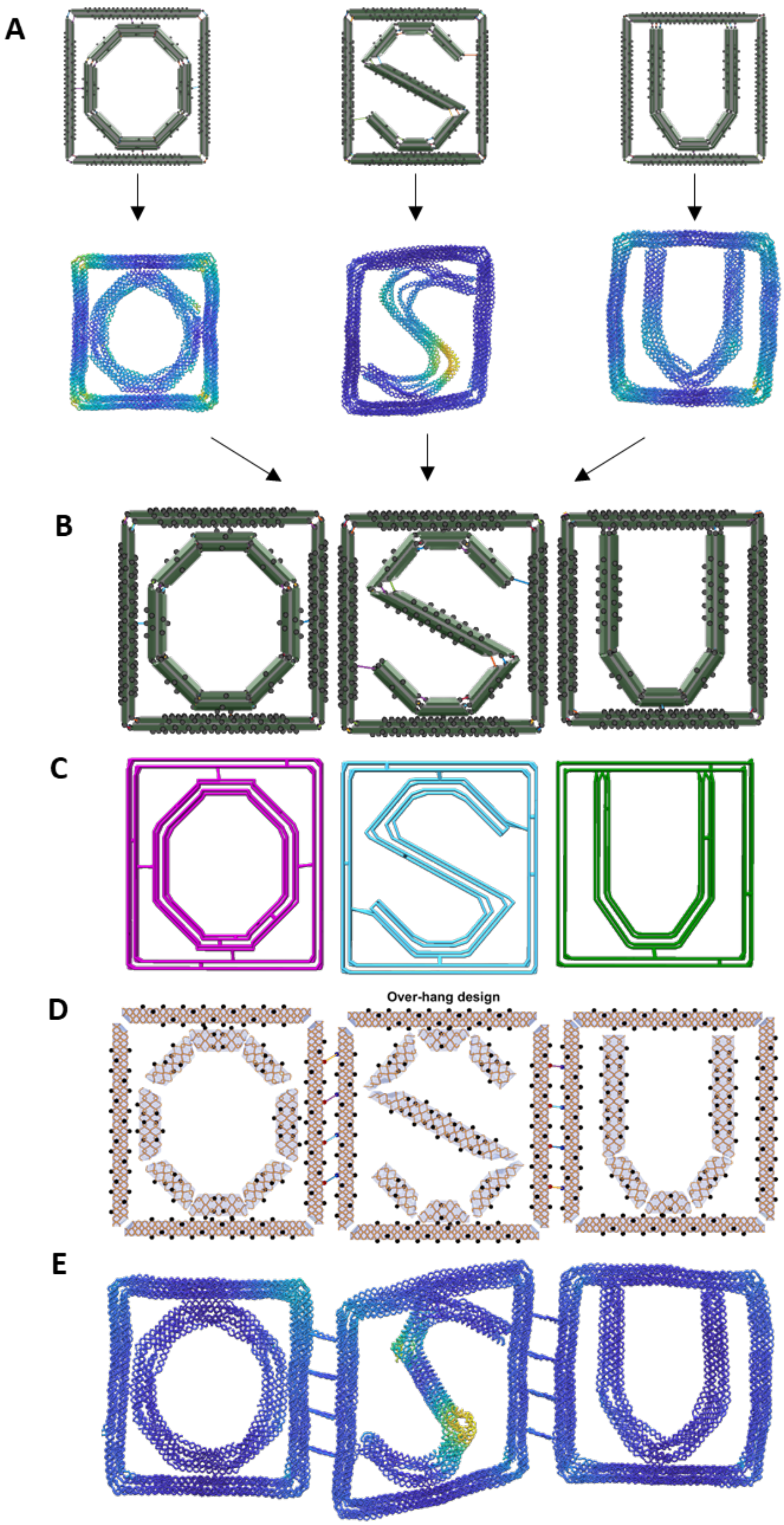

**Supplementary Figure 79: Example of design for hierarchical assembly of multiple DNA nanostructures using the multi-scaffold algorithm.** (A) Individual design and iterative optimization. (B) Insert the design geometries of each nanostructure, “O”, “S”, and “U” into one assembly design. Without adding new connectivity between each nanostructure, manipulate and assemble them with staple overhangs. (C) Due to the disconnection of the connectivity matrix, the multi-scaffolding algorithm first obtained a spanning forest consisting of three separate spanning trees, which guided the determination of three separate scaffold routings, one for each structure. (D) The overhang design tool was used for designing the staple overhangs and binding staples to enable hierarchical assembly of multiple nanostructures. (E) oxDNA simulation of the hierarchical structure.

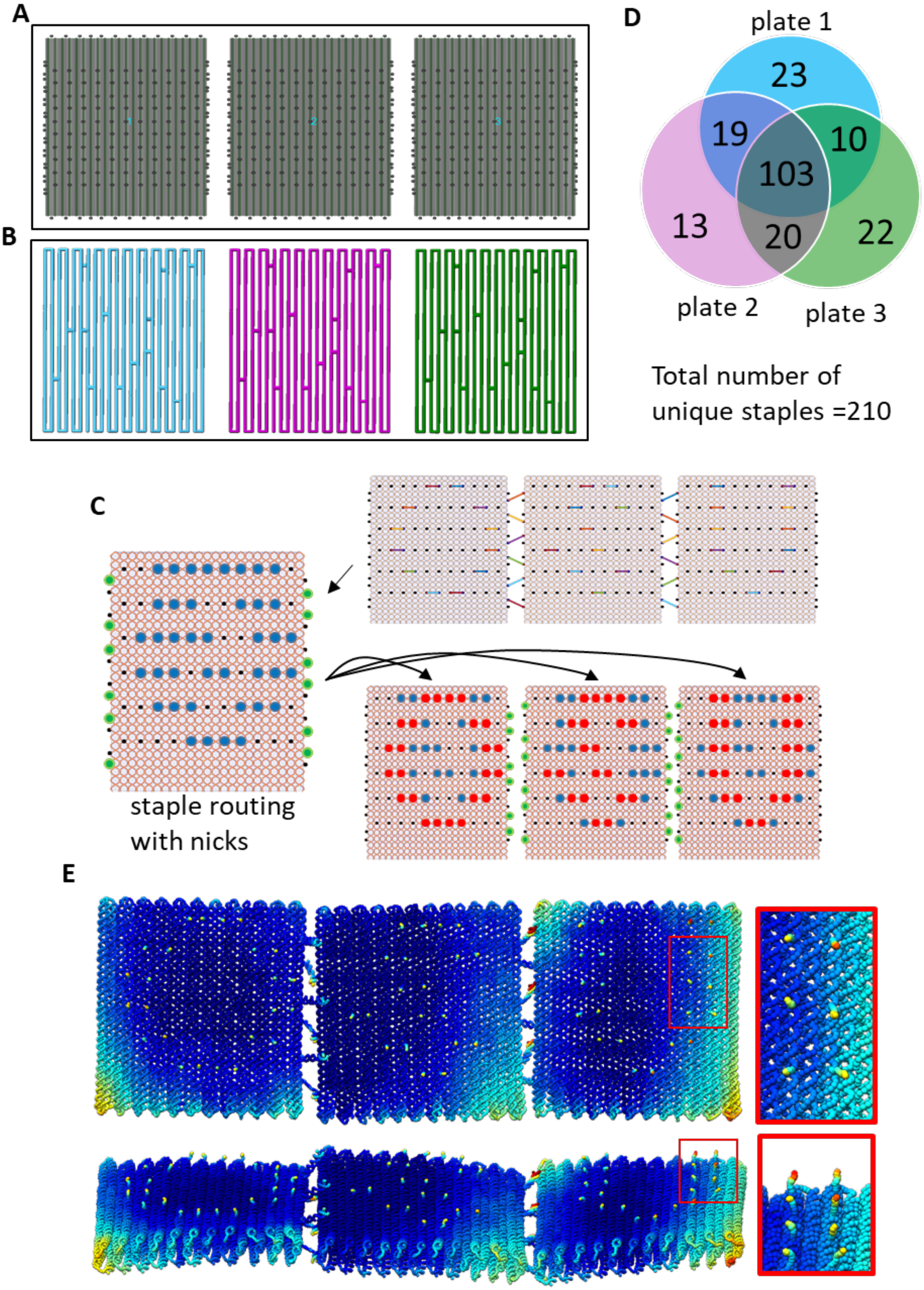

**Supplementary Figure 80: Example of design for hierarchical assembly using repeated staple sequences for cost saving.** (A) A plate template was created and duplicated into multiple identical instances. (B) Without adding any connectivity, the scaffold algorithm yields a spanning forest with three separate spanning trees, which guides the determination of three different scaffold routings, one for each plate. The scaffold routing of one plate was then duplicated to replace the routing of the other two plates, so all three plates had the identical scaffold routing. (C) Here we used our overhang design tool in 3D to decorate the surfaces of the three plates. (Top-right) The snapshot of clicking the shape of three letters (“O”, “S”, and “U”) on the surfaces. (Left) To guarantee the same routing in staples before extending the overhangs, the union of the nick positions (green and blue dots) on three plates was taken into account for the staple routing template and then was duplicated to the staple routing for all three plates using a custom script. The script of these processes is provided with the software package as well as the example design file. (Bottom-right) With the original overhang design algorithm, the overhangs were extended at the specified locations (red dots for decoration and green dots for connections) from each plate. (D) The staple set analysis where 103 staples (49%) are repeated in three plates. (E) Single-stranded overhangs are expected to have higher RMSF values in simulation. Hence, the target shape, “O-S-U”, can be seen on the three surfaces.

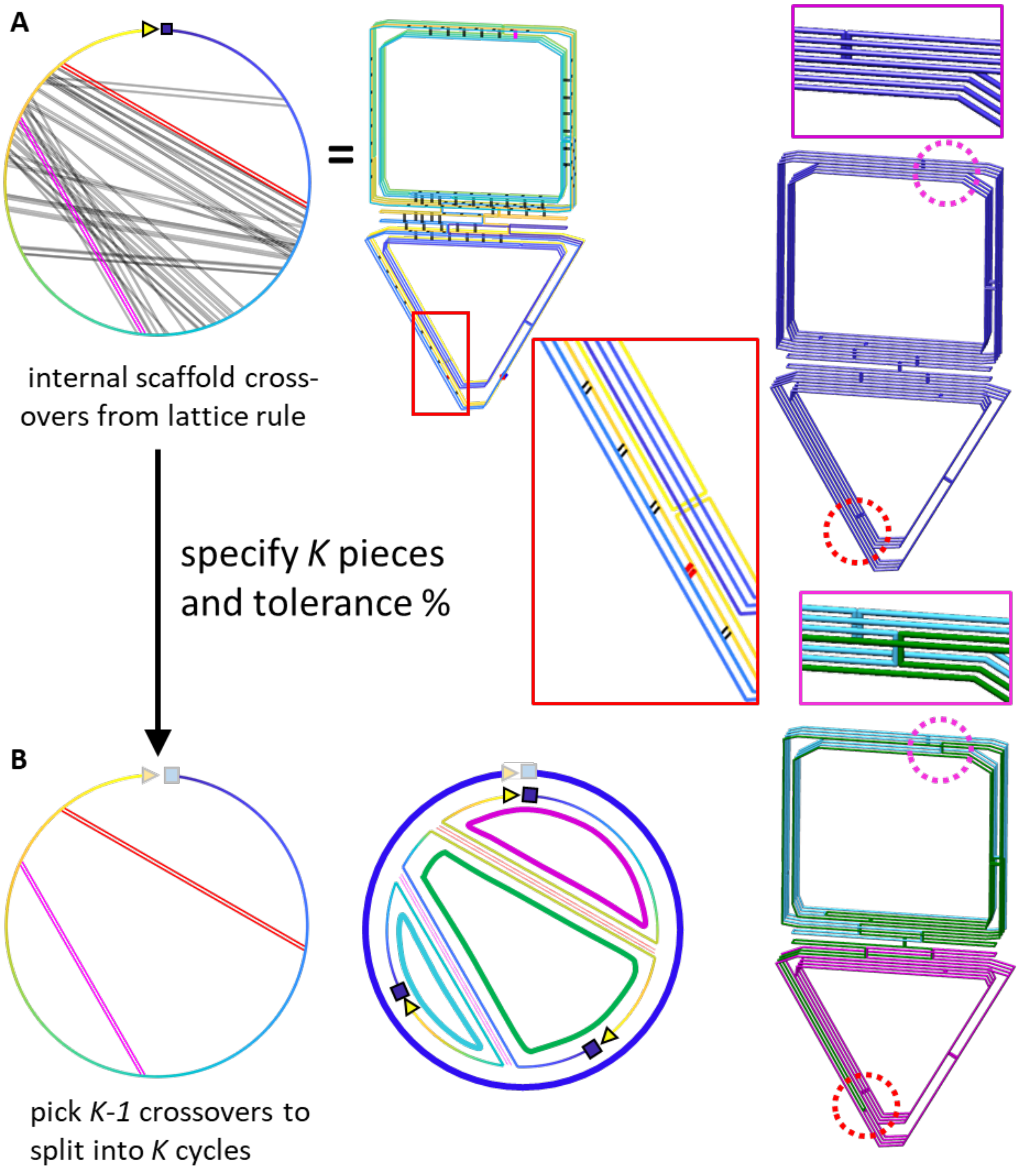

**Supplementary Figure 81: The second approach for multi-scaffold routing by searching and applying K-1 internal crossovers to one long scaffold routing.** (A) The circular diagram (Left) is a schematic representation of the 3D routing (right) to illustrate the locations of the potential internal crossovers from the lattice rule (inset, black pairs of parallel lines) that would split the single cycle into two cycles. Not all possible crossovers are shown in the circular diagram. Some crossovers were excluded if their two ends on the circular diagram are too close. Others were removed for clarity of visualization, but all are still shown in the 3D routing on the right. This approach also considers the lengths of split scaffolds as a constraint when randomly selecting and applying K-1 crossovers to the original one scaffold. (B) Circular diagram illustrates an example of splitting one large cycle into three scaffold cycles. The essential goal is to find two internal crossovers such that the three split cycles satisfy the length constraint according to what scaffolds are intended for use. In addition, the selected crossovers cannot cross each other. Otherwise, the number of total cycles may not always increase, which can result in an incorrect number of cycles.

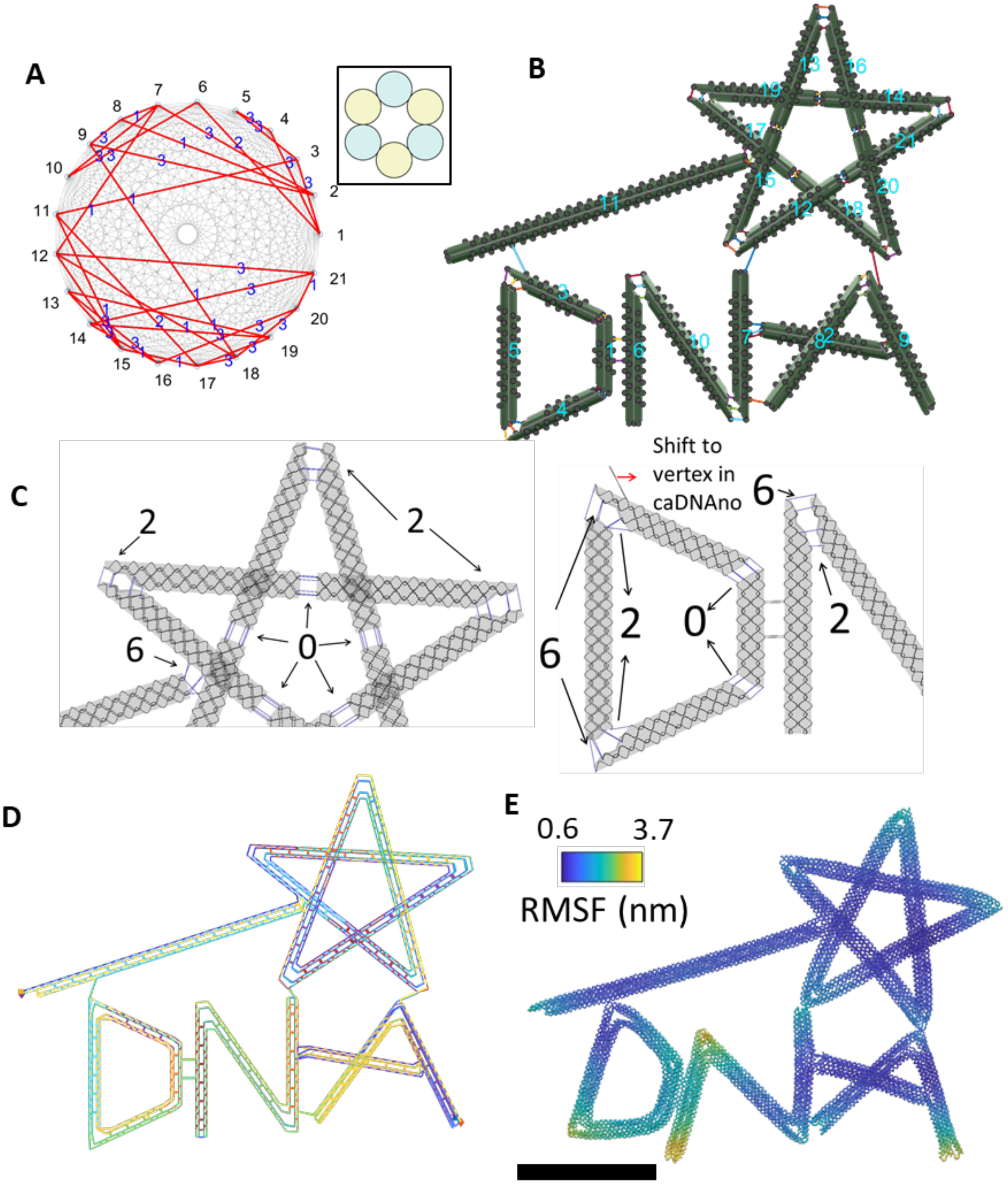

**Supplementary Figure 82: Design details for the MagicDNA logo structure.** (A) The connectivity matrix for assembly and the cross-section (6-HB) for all bundles. (B) The design model in the MagicDNA assembly panel. (C) The length of scaffold ssDNA connections between individual bundles. (D) The scaffold and staple routing of the final design. (E) The oxDNA simulation result. Scale bar = 50 nm.

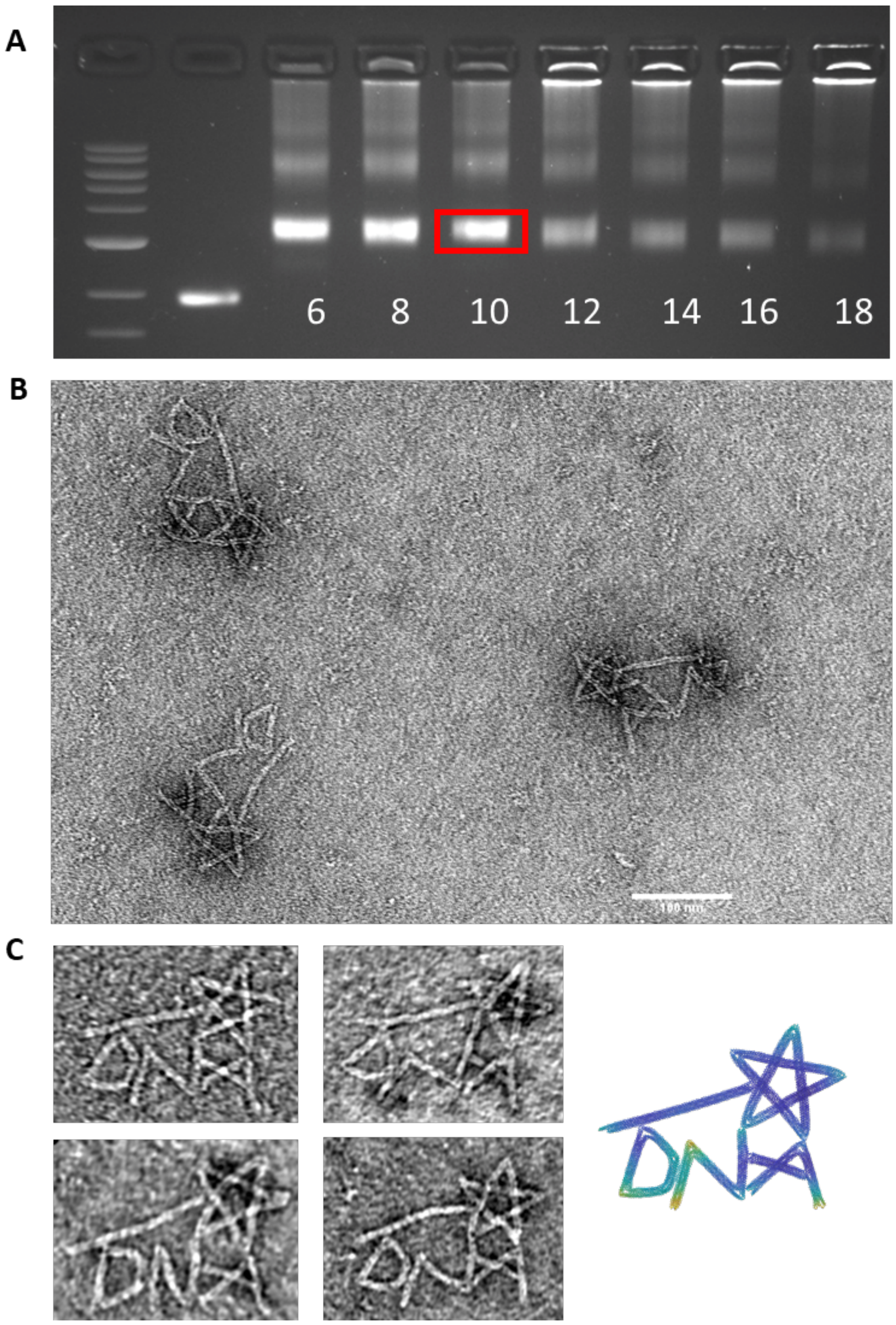

**Supplementary Figure 83: Experimental validation of the MagicDNA logo structure.** (A) Logo was folded using a two-and-a-half-day modified thermal annealing protocol while testing a range of magnesium chloride concentrations from 6-18 mM purified using agarose gel electrophoresis (buffer conditions 0.5xTBE 5.5 mM MgCl2). (B) Zoomed-out TEM image of gel-purified logo folded at 10 mM MgCl2. (C) (Left) Zoomed-in TEM images of gel-purified logo folded at 10 mM MgCl2. (Right) oxDNA simulation of logo.

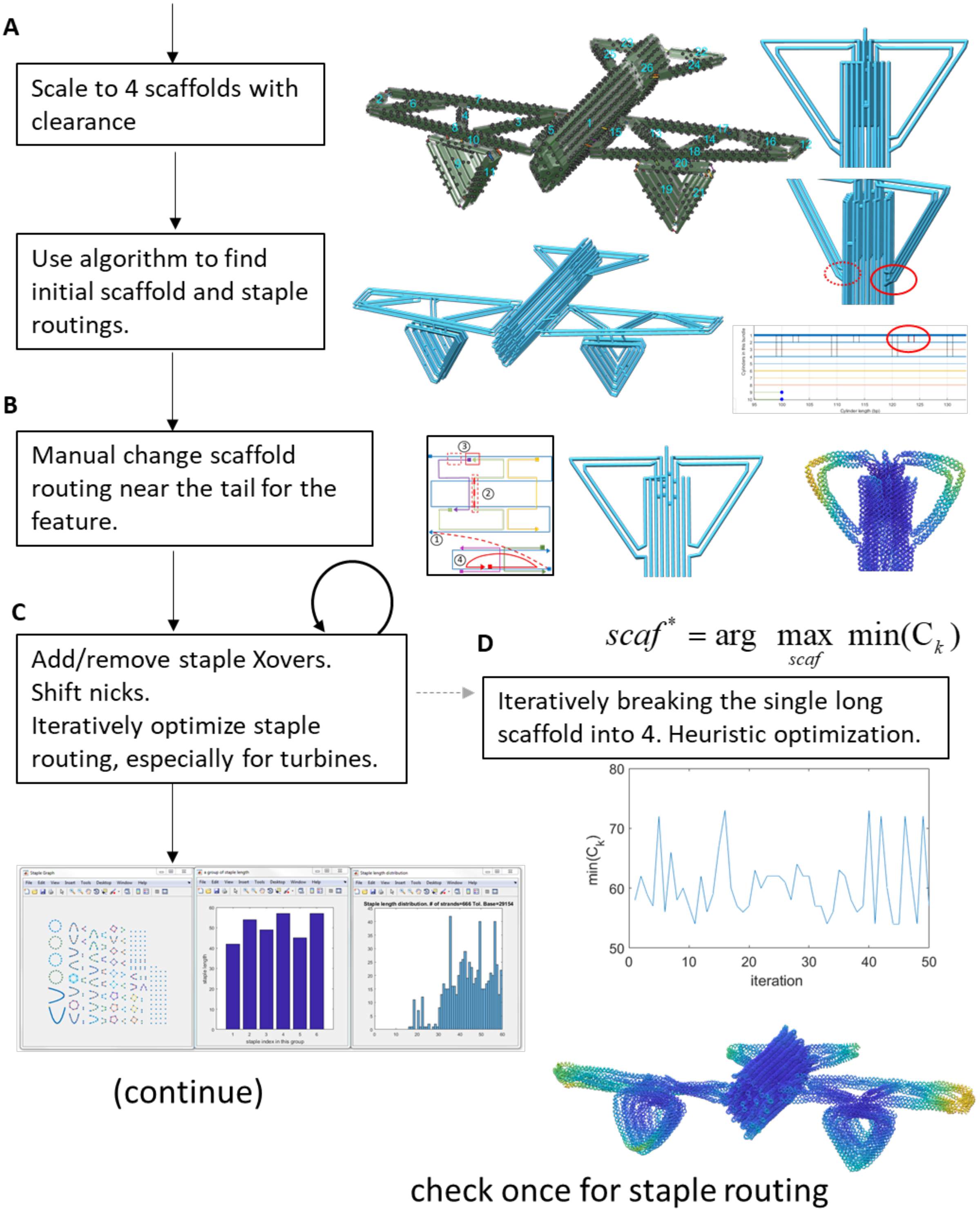
A
Scale to 4 scaffolds with clearance
Use algorithm to find initial scaffold and staple routings.
B
Manual change scaffold routing near the tail for the feature.
C
Add/remove staple Xovers.
Shift nicks.
Iteratively optimize staple routing, especially for turbines.
(continue)
D
scaf* = arg max min(C_k)
scaf
Iteratively breaking the single long scaffold into 4. Heuristic optimization.
min(C_k)
iteration
check once for staple routing

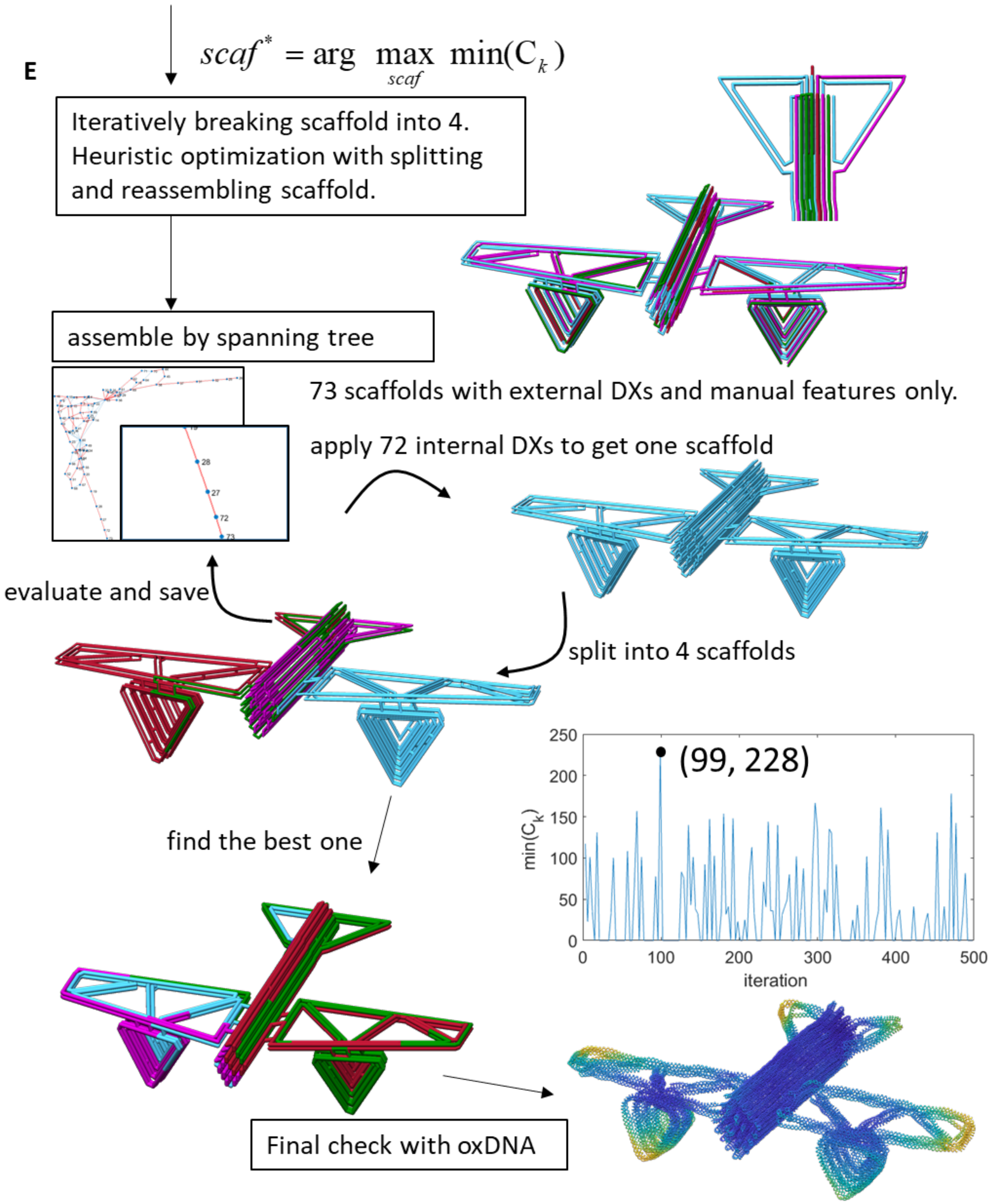

**Supplementary Figure 84: The process of designing the airplane made up of four scaffolds.** (Continue from Supplementary Figure 69.) (A) We planned to use 4 scaffolds (M13-8064, and orthogonal scaffolds CS3_L_7560, CS4_7557, CS5_7559 (41)) to fabricate this structure. The airplane design in Fig. S69 was around 33 kbps, which was over the maximum limit of scaffold lengths; hence, we used the bottom-up approach to scale with the bundle editing GUI. Specifically, we shortened some cylinders and simplified the tail design. (B) For reducing the flexibility of the horizontal stabilizers on the tail, three-way junctions were introduced between the stabilizers and the body. Since this type of routing is not directly supported by the scaffold algorithm, we assigned two user-defined internal crossovers between the two external double-scaffold crossover connections (B24-B26 and B25-B26) and later used caDNAno to shift the half-crossovers as shown in the insets. (C) Before splitting the single-scaffold routing, the staple algorithm obtained an initial staple routing which was modified with caDNAno with the assistance of visualization and mapping GUIs of MagicDNA (Fig. S18). (D) To ensure stable binding between all the scaffolds in the multi-scaffold structure, we developed a heuristic optimization process (Eqs. 8 to 10) where the optimization seeks to maximize the objective function, which is the minimum number of staples connecting from one scaffold to the others. In this process, while the staple routings remained the same, the multi-scaffold routings were iteratively tested by searching and applying three internal scaffold crossovers to the initial single cycle while considering the scaffold length constraint. The results were evaluated by inspecting results using the visualizations of multi-scaffold results shown in Fig. S19. The optimal multi-scaffold routing that was obtained by dividing up one single initial large single cycle yielded a minimum of 73 staples that connected one scaffold to the others. (E) To further improve the heuristic search process, we aimed to enlarge the domain space of the initial single-scaffold routing. However, the stabilizers at the tail had manual modifications which restricts direct use of the scaffold routing algorithm. Alternatively, we detected and removed the internal 72 scaffold crossovers in the modified one-scaffold routing and obtained the scaffold routings with 73 cycles (reverse of the spanning tree). Since the spanning tree guides the integration of cycles in a stochastic process (i.e. the selection of crossovers on tree edges is random), repeating the integration of these 73 scaffold cycles allows determination many different initial large single-scaffold routings. Combining this reassembly process with the heuristic optimization process in (D) generated the 4-scaffold routing design with at least 228 staples that connected any two scaffolds. This was used as the final scaffold routing for the experimental fabrication. The script of these processes is provided with the software package.

$$B_{ij}\text{:numbers of complementary bases } \begin{cases} i: \text{ staple index} \\ j: \text{scaffold index} \end{cases} \tag{8}$$

$$\begin{aligned} &C_k: \text{ Scaf } k \text{ has } C_k \text{ staples that connect to other scaffolds} \\ &= \sum_{i=1}^{I} (\text{AND}(B_{ik} > 0, \sum_{p \neq k} B_{ip} > 0)) \end{aligned} \tag{9}$$

$$scaf^* = \arg \max_{scaf} \min(\mathrm{C}_k) \tag{10}$$

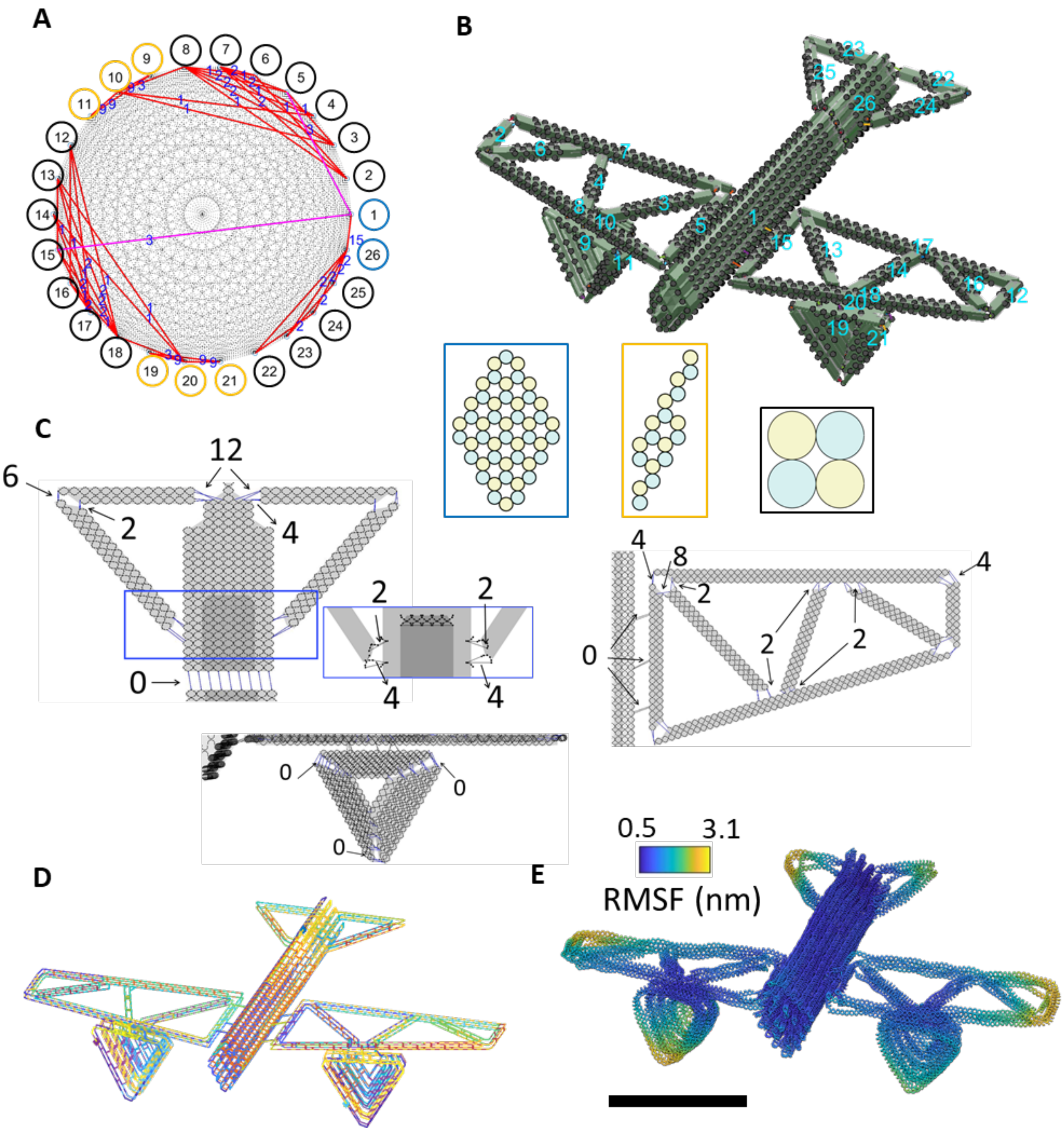

**Supplementary Figure 85: Design details for the airplane structure.** (A) The connectivity matrix for assembly and the cross-sections of each component. (B) The design model in the MagicDNA assembly panel. (C) The length of scaffold ssDNA connections between individual bundles. (Inset) The single-stranded length on the tail was assigned as some values before the algorithm. However, we used the caDNAno interface to change the local routing near the tail shown in right. Hence the correct lengths of scaffold ssDNA were labeled separately. (D) The scaffold and staple routing of the final design. (E) The oxDNA simulation result. Scale bar = 50 nm.

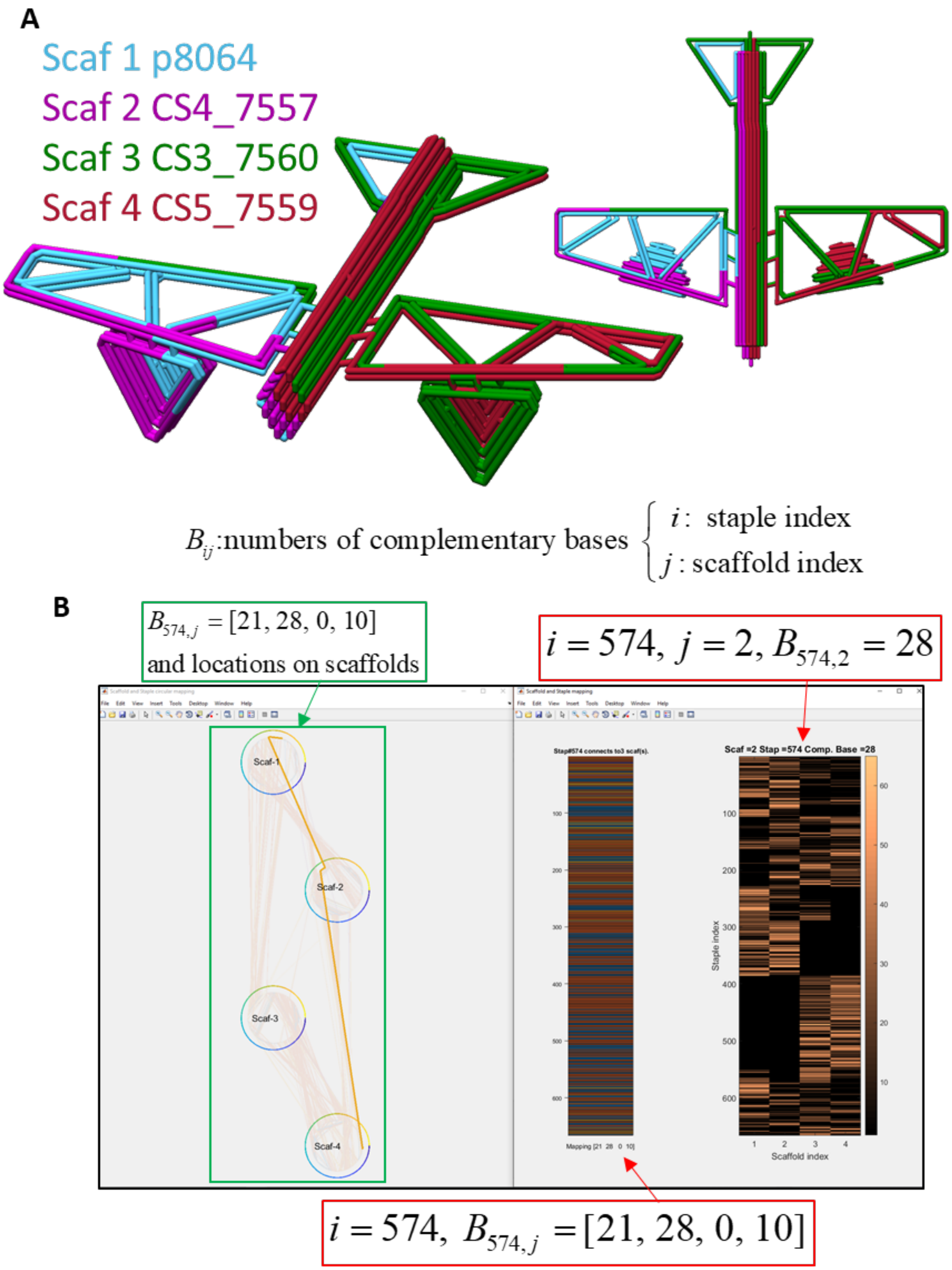

**Supplementary Figure 86: The final four-scaffold routing design and the mapping with staples of the full airplane structure.** (A) The routing results of the heuristic method to maximize the level of "interconnection" between scaffolds. (B) Use the inspection function in the GUI in Fig. S20 to visualize the mapping between 4 scaffolds and 666 staples. The 666×4 complementary matrix describes the number of complementary bases between staples and scaffolds. Note that there are additional 16 staples for binding with unused scaffold sections which are not shown here. It is visualized in a planar color map which indicates the value of the selected element through GUI function. Meanwhile, the complementary bases on scaffolds are also shown and highlighted in another window. In this window, one can see that the number of crossing staples between the two cases (scaffold-1 to scaffold-4, and scaffold-2 to scaffold-3) are significant less than the other four cases.

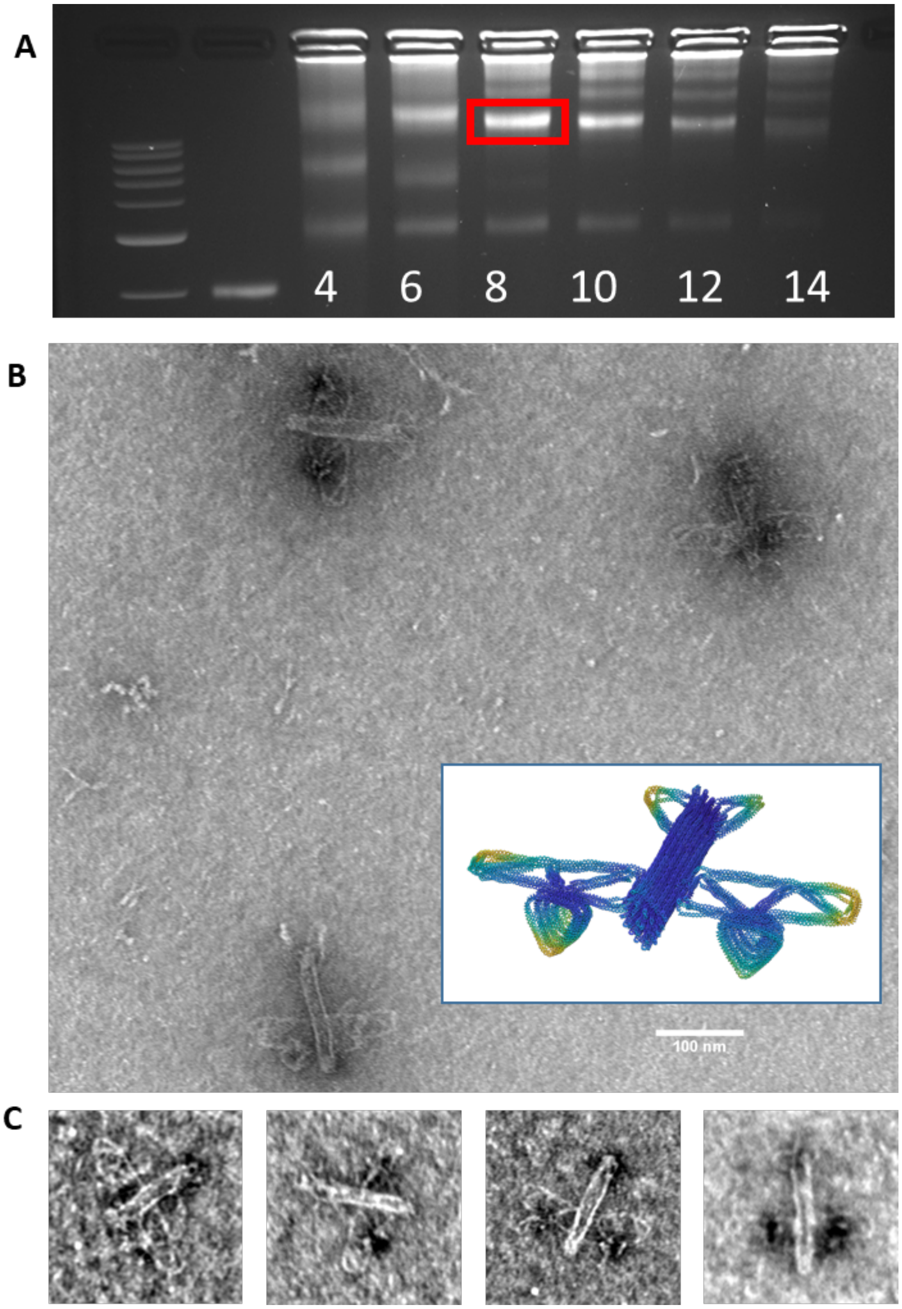

**Supplementary Figure 87: Experimental validation of the airplane structure.** (A) Airplane was folded using a two-and-a-half-day modified thermal annealing protocol while testing a range of magnesium chloride concentrations from 4-14 mM and purified using agarose gel electrophoresis (buffer conditions 0.5xTBE 5.5 mM MgCl2). (B) Zoomed-out TEM image of gel-purified logo folded at 8 mM MgCl2. (C) Zoomed-in TEM images of gel-purified logo folded at 8 mM MgCl2. and oxDNA simulation of the airplane.

#### Movie captions:

Movie S1. Top-down parametric design for a hinge structure performed by converting two lines into bundles and specifying the connectivity to assemble the components.

Movie S2. The final design profile of the airplane and the CG simulation with $3\times10^8$ steps

#### Additional supplementary Materials:

The pdf file for the software user manual

The excel sheets for the staple list of the 14 structures for fabrication

**Supplementary References**

1. Douglas, S. M. *et al.* Rapid prototyping of 3D DNA-origami shapes with caDNAno. *Nucleic Acids Res* **37**, 5001–5006 (2009).
2. Huang, C.-M., Kucinic, A., Le, J. V., Castro, C. E. & Su, H.-J. Uncertainty quantification of a DNA origami mechanism using a coarse-grained model and kinematic variance analysis. *Nanoscale* **11**, 1647–1660 (2019).
3. Snodin, B. E. K., Schreck, J. S., Romano, F., Louis, A. A. & Doye, J. P. K. Coarse-grained modelling of the structural properties of DNA origami. *Nucleic Acids Res* **47**, 1585–1597 (2019).
4. Doye, J. P. K. *et al.* Coarse-graining DNA for simulations of DNA nanotechnology. *Phys. Chem. Chem. Phys.* **15**, 20395–20414 (2013).
5. Snodin, B. E. K. *et al.* Introducing improved structural properties and salt dependence into a coarse-grained model of DNA. *J. Chem. Phys.* **142**, 234901 (2015).
6. Pettersen, E. F. *et al.* UCSF Chimera—A visualization system for exploratory research and analysis. *J. Comput. Chem.* **25**, 1605–1612 (2004).
7. Castro, C. E. *et al.* A primer to scaffolded DNA origami. *Nat Meth* **8**, 221–229 (2011).
8. Engelhardt, F. A. S. *et al.* Custom-Size, Functional, and Durable DNA Origami with Design-Specific Scaffolds. *ACS Nano* **13**, 5015–5027 (2019).
9. Sobczak, J.-P. J., Martin, T. G., Gerling, T. & Dietz, H. Rapid Folding of DNA into Nanoscale Shapes at Constant Temperature. *Science* **338**, 1458–1461 (2012).
10. Ponnuswamy, N. *et al.* Oligolysine-based coating protects DNA nanostructures from low-salt denaturation and nuclease degradation. *Nature Communications* **8**, 15654 (2017).
11. Tang, G. *et al.* EMAN2: An extensible image processing suite for electron microscopy. *Journal of Structural Biology* **157**, 38–46 (2007).

12. Abramoff, M. D., Magalhães, P. J. & Ram, S. J. Image processing with ImageJ. *Biophotonics international* http://localhost/handle/1874/204900 (2004).
13. Williams, S. *et al.* Tiamat: A Three-Dimensional Editing Tool for Complex DNA Structures. in *DNA Computing* 90–101 (Springer, Berlin, Heidelberg, 2008).
14. Matthies, M., Agarwal, N. P. & Schmidt, T. L. Design and Synthesis of Triangulated DNA Origami Trusses. *Nano Lett.* **16**, 2108–2113 (2016).
15. Benson, E. *et al.* DNA rendering of polyhedral meshes at the nanoscale. *Nature* **523**, 441–444 (2015).
16. Veneziano, R. *et al.* Designer nanoscale DNA assemblies programmed from the top down. *Science* **352**, 1534–1534 (2016).
17. Jun, H. *et al.* Automated Sequence Design of 3D Polyhedral Wireframe DNA Origami with Honeycomb Edges. *ACS Nano* (2019).
18. Shi, Z., Castro, C. E. & Arya, G. Conformational Dynamics of Mechanically Compliant DNA Nanostructures from Coarse-Grained Molecular Dynamics Simulations. *ACS Nano* **11**, 4617–4630 (2017).
19. Kim, D.-N., Kilchherr, F., Dietz, H. & Bathe, M. Quantitative prediction of 3D solution shape and flexibility of nucleic acid nanostructures. *Nucleic Acids Res* **40**, 2862–2868 (2012).
20. Martin, T. G. & Dietz, H. Magnesium-free self-assembly of multi-layer DNA objects. *Nature Communications* **3**, 1103 (2012).
21. Poppleton, E. *et al.* Design, optimization, and analysis of large DNA and RNA nanostructures through interactive visualization, editing, and molecular simulation. *bioRxiv* 2020.01.24.917419 (2020).

22. Reshetnikov, R. V. *et al.* A coarse-grained model for DNA origami. *Nucleic Acids Res* **46**, 1102–1112 (2018).

23. Marras, A. E., Zhou, L., Su, H.-J. & Castro, C. E. Programmable motion of DNA origami mechanisms. *PNAS* **112**, 713–718 (2015).

24. Marras, A. E., Zhou, L., Kolliopoulos, V., Su, H.-J. & Castro, C. E. Directing folding pathways for multi-component DNA origami nanostructures with complex topology. *New J. Phys.* **18**, 055005 (2016).

25. Sharma, R., Schreck, J. S., Romano, F., Louis, A. A. & Doye, J. P. K. Characterizing the Motion of Jointed DNA Nanostructures Using a Coarse-Grained Model. *ACS Nano* (2017).

26. Maffeo, C. & Aksimentiev, A. MrDNA: a multi-resolution model for predicting the structure and dynamics of DNA systems. *Nucleic Acids Res*.

27. Han, D. *et al.* DNA Origami with Complex Curvatures in Three-Dimensional Space. *Science* **332**, 342–346 (2011).

28. E. Castro, C., Su, H.-J., E. Marras, A., Zhou, L. & Johnson, J. Mechanical design of DNA nanostructures. *Nanoscale* **7**, 5913–5921 (2015).

29. Benson, E. *et al.* Effects of Design Choices on the Stiffness of Wireframe DNA Origami Structures. *ACS Nano* (2018).

30. Stewart, D. A Platform with Six Degrees of Freedom. *Proceedings of the Institution of Mechanical Engineers* **180**, 371–386 (1965).
