## Supplementary material for "Integrating computer-aided engineering and computer-aided design for DNA assemblies": MagicDNA software manual

(**M**ulti-component **A**ssembly in a **G**raphical **I**nterface guided by **C**omputation for **DNA** origami)

The Ohio State University

Playlists for tutorial movies (TM), all available on the MagicDNA YouTube channel (Fig. 1A) (https://go.osu.edu/magicdna_tutorial ):

A. MagicDNA GUI tutorials (The movies in this playlist are also embedded in the software package for each GUI. Press [h] in any GUI window to play the corresponding movie in MATLAB, or all movies are also accessible in the 'InstructionMovie" folder.)
   - A1 Sketch in MagicDNA to create lines.
   - A2 Basic editing of line model from .STEP files.
   - A3 Convert the line model into bundles.
   - A4 The bundle editor GUI
   - A5 The Assembly GUI in MagicDNA
   - A6 The single-stranded scaffold GUI
   - A7 The algorithm for scaffold routing
   - A8 The options for scaffold routing
   - A9 The algorithm for staple routing without overhangs.
   - A10 The overhang design tool
   - A11 The sequence and design diagram GUI
   - A12 MagicDNA to oxDNA
   - A13 oxDNA trajectory to MagicDNA

B. MagicDNA design examples (This playlist contains step-by-step instructions to design structures, covering different levels of complexity and using optional tools in MagicDNA for specific cases.)
   - B1 Example: a series of hinges from two lines.
   - B2 Example: a series of ring from a circular pattern of lines/bundles
   - B3 Example: the design of the Stewart platform in DNA origami
   - B4 Example: the butterfly mechanism (modify in caDNAno, the overhang design tool)
   - B5 Example: the airplane structure (Bottom-up approach from sub-systems, split a long scaffold into 4)
   - B6 Example: the interlock ring. (Top-down in parallel, bottom-up, edit bundle GUI)
   - B7 Example: the tower structure. (.STEP top-down, edit bundle GUI, modify in caDNAno)
   - B8 Example: the triangle plate. (multi-scaffold)

C. oxDNA simulations (This playlist is the collection of oxDNA trajectories for compliant or dynamic structures. The simulations were conduct with 3e8 steps and 300 frames, which is 30 times of normal simulation length for static structures.)
   - C1 The oxDNA simulation of the hollow hinge
   - C2 Compliant compound joint
   - C3 The oxDNA simulation of the butterfly mechanism
   - C4 The design and simulation trajectory of the airplane structure with four scaffolds

### 1. Introduction

Structural DNA nanotechnology[1,2] has offered a reliable nanofabrication method for devices with diverse applications ranging from nanorobotics[3–5] to medicine[6,7]. In general, DNA self-assembly leverages the sequence-specific base-pairing of DNA strands to program anywhere from a two up to hundreds or even thousands of strands to assemble into various geometries ranging from basic helices to 2D sheets, 3D lattice-based bundles, and wireframe structures. Scaffolded DNA origami[8,9] is arguably the most robust design approach that, in addition to complex nanoscale geometry, enables multi-component structures with programmed motion, mechanical properties, and dynamic behavior. In this approach, a long (typically ~7000-8000 nucleotides) “scaffold” strand is folded into a compact shape by base-pairing in a piecewise manner to ~100-200 “staple” strands (typically ~30-60 nucleotides long). The general design process starts by approximating a desired geometry out of helices, or at this stage they are usually abstracted as cylinders. The detailed design then involves scaffold routing (i.e. routing the scaffold strand through the target geometry), staple routing with frequent crossovers to constrain the shape, and finally, generating a list of staple sequences based on the complementary binding to the scaffold. The details of the experimental fabrication process can be found in many manuscripts[10–12].

The most widely used design software, caDNAno[13], has a user-friendly interface to design 3D lattice-based structures; however, the bottom-up design process limits the number of components and the complexity of structures owing to the mapping of a 3D structure onto a 2D design diagram and the lack of scaffold routing algorithm. Recent semi- or fully-automatic softwares[14–17] focus on developing a scaffold routing algorithm to increase the number of component and speed up the design process. However, these softwares are limited to particular types of geometry, and in particular, they are not amenable to design of multi-component dynamic devices. Due to the increasing demand for complex designs, we present a versatile robust framework in which the target shape is approximated by multiple bundles and their connections in 3D space. Users can mainly focus on component geometries and their assembly in our graphical user interfaces (GUIs) with the help of the routing algorithm to finalize the design. Since this software tool enables design of increasingly complex structures, we facilitate an iterative design process by enabling straightforward interfacing with the coarse-grained simulation tools oxDNA[18–21] and mrDNA[22] to obtain simulation feedback (e.g. average shape, fluctuations, and motion analysis) that can be used to guide design modifications. We also provide an interface to caDNAno for fine tuning of underlying strand routing, and we included features that allow for design of overhangs for actuation or functionalization, direct visualization in the software or exporting a variety of 3D structure visualizations, and capabilities to incorporate multiple scaffolds into a single structure[23] or directly design higher order assemblies[24,25]. We believe the combination of these features can facilitate expanding the design domain of DNA origami self-assembly into new regimes of complexity and function.

This document with high-level instructions is meant to outline the corresponding tutorial movies on YouTube that contain low-level step-by-step detailed processes with the advantage of visualizing steps and interaction with UI components. Here we illustrate the general design process, going through each GUI, and we show several specific examples of designs covering different complexities. The GUI tutorial movies illustrate the complete design process in detail and also go through design examples including, but not limited to, the examples in this manual. The movies can be accessed directly in the software (Playlist A: GUI tutorial, details below) or the YouTube link (All movies). We highly recommend watching and following along through design examples with the movies since they illustrate directly steps of the design process.

#### 2. Installation

This software is called MagicDNA (**M**ulti-component **A**ssembly in a **G**raphical **I**nterface guided by **C**omputation for **DNA** assemblies) and requires MATLAB 2017a or later versions to run. It also requires the Bioinformatics Toolbox and the Statistics and Machine Learning Toolboxes for MATLAB. MagicDNA has been packaged as a MATLAB app (MagicDNA_v2.0.mlappinstall) and can be installed with the attached package (Fig. 1B).

After installing the app (MagicDNA_v2.0.mlappinstall), MagicDNA, the software shows up as an app in MATLAB apps list. But the folder is located in somewhere else (Ex: ….\Documents\MATLAB\Add-Ons\Apps\MagicDNA_v20\code\). Users can either: 1) go to there as the working directory or 2) copy the entire MagicDNA folder to a preferred location, and directly change the MATLAB directory to work in that folder. Once the directory is set to that folder, the user can click on the icon in the app list or type 'main' in the command line to start the app.

Required: MATLAB v2017a, Bioinformatics Toolbox, Statistics and Machine Learning Toolbox

Resource: GitHub link: https://github.com/cmhuang2011/MagicDNA

YouTube link: https://go.osu.edu/magicdna_tutorial

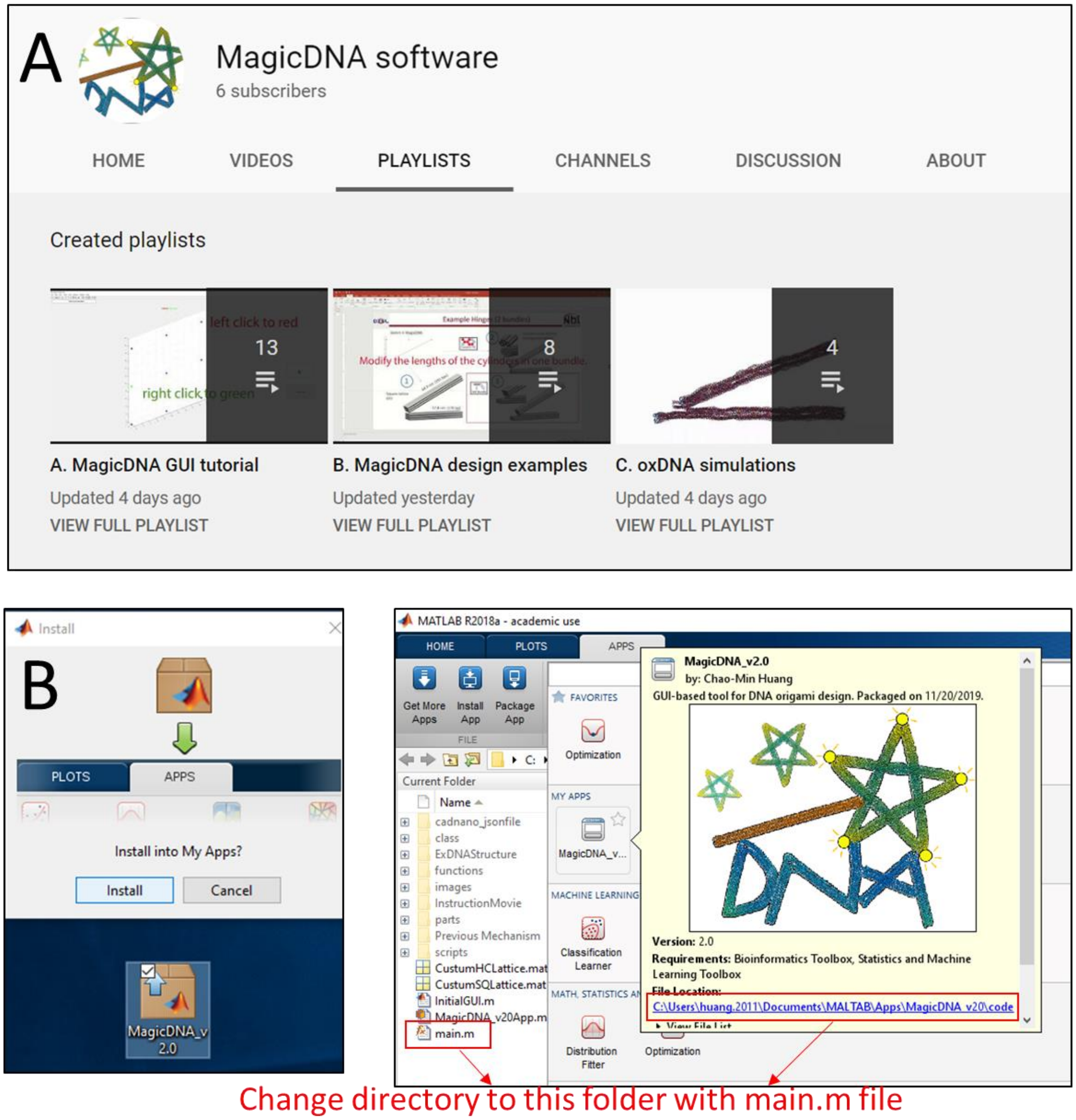

*Figure 1. (A) YouTube channel snapshot for tutorial movies. (B) Install MagicDNA to MATLAB app list.*

#### 3. Getting started

MagicDNA is a GUI-based software in which most of operations are performed by interacting with the GUIs, or program window (a.k.a. "figure" in MATLAB). Fig. 2 shows a screenshot of the "Assembly" GUI. Opening multiple windows is possible and sometime is helpful for designing large structures with several components. We have created instructions for each GUI that can be accessed by clicking on the tab at the top-left corner (red box in Fig. 2), where there are descriptions about the proper operations and GUI functions, including steps triggered by mouse clicks and keyboard keys. The "Part" tab shown in Fig. 2 contains a way to create parts. However, it was created for purely bottom-up approach in the beginning of the software development and we do not recommend using this feature as we have not optimized the interface, and the same designs can be achieved directly in the Sketch and Assembly GUIs.

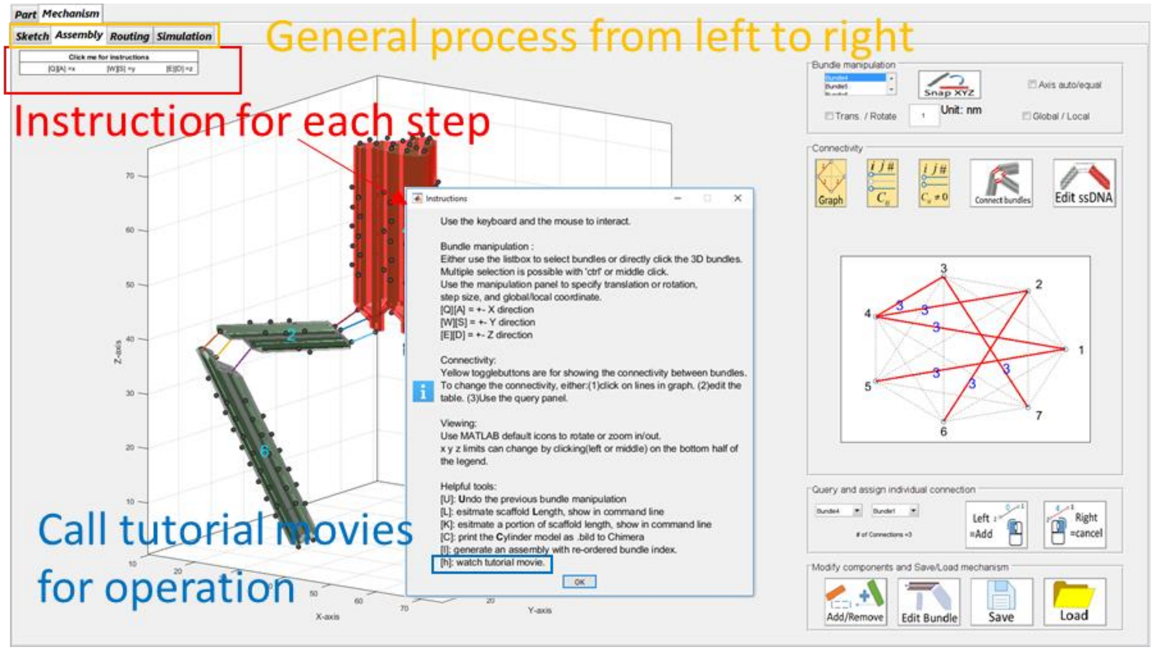

*Figure 2. MagicDNA interface and instructions.*

### 4. General Design Process

The goal of the MagicDNA software is to simplify the design process and enable a user-friendly and versatile tool that covers a wide range of design domains from simple to complex. Some tools in MagicDNA are necessary to use for every design process, and some are additional functions to expand the range of design capabilities. For detailed operations of the processes, we recommend watching the GUI tutorial movies (Playlist A) along with this section.

The general process has four major steps:

(1) Geometries of each bundle (top-down or bottom-up), (2) Assembly, (3) Routing algorithm, and (4) Simulation, as shown in Fig. 3.

#### 4.1. Defining/Editing the geometry

The first step of the design process is to define the Geometry. In MagicDNA, defining the Geometry refers to specifying the number of components, the overall layout of components (i.e. bundles) or initial configuration of the design (note this can be adjusted later in Assembly), and the detailed geometry of individual bundle components.

##### 4.1.1. The top-down approach for the geometry (see Tutorial Movies(TM) A1-A3)

For defining the geometry (Fig. 3A), when we mention a top-down approach, we refer to the use of a line model to define the overall configuration and the conversion of individual into bundles by assigning required parameters, such as cross-section and length. A line model can be created by inputting a .STEP (Standard for the Exchange of Product Data) file, which is a standard computer-aided design (CAD) file format for line models, or by directly sketching a line model in MagicDNA. This step generates a defined number of components, or bundles, in a user-specified configuration (this configuration can be fine-tuned or adjusted later), which is then passed to the Assembly GUI for additional operations.

4.1.2. The bottom-up approach for the geometry

On the other hand, when we mention a bottom-up approach, we refer to saving a bundle or multiple bundles (using "Save bundle" icon in the bundle editor GUI or "Save" icon depicted in Fig. 4) into a library and inserting (using "Add/Remove" icon depicted in Fig. 4) and combining multiple bundles or assemblies into a larger design.

4.1.3. Fine-tuning bundle geometry using the edit bundle GUI (optional) (see TM A4)

Fine tuning the geometry of individual bundles using the "edit bundle" GUI to enhance control over local the geometry. Features in the "edit bundle" GUI include visualizing the selected bundle, extruding the lengths of individual or multiple helices, changing the pairing between helices (a pair refers to two helices connected together by scaffold crossovers at both ends), and specifying internal crossover locations for one type of multi-scaffold algorithm. If any modifications related to extrusion and pairing are carried out in the "edit bundle" GUI, the user should export all bundles by using the "add/remove" button to update the entire cylinder model before going the routing step.

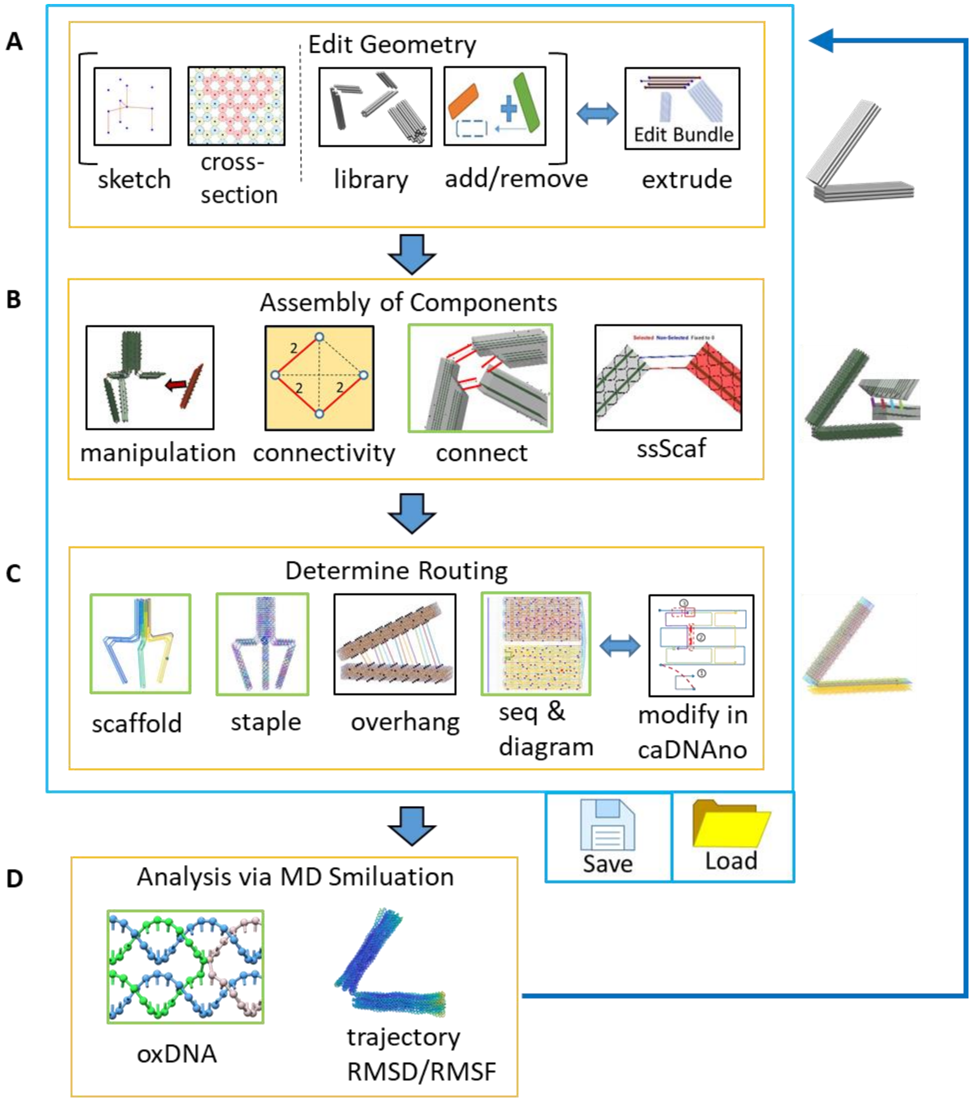

*Figure 3. General design process of this robust design framework.*

#### 4.2. Assembly of components

The next step is Assembly of Components, which we refer to here just as Assembly (Fig. 3B).

##### 4.2.1. Manipulate components (TM A5 0:00~1:16)

The first step of Assembly is to translate and rotate the components in 3D space to the approximate position to achieve a desired configuration (Fig. 4). The user can select one bundle in the list box or directly click on the bundle/component with the mouse in the 3D solid model window. Users can also select multiple components at once by using the middle click button on the mouse or selecting bundles in the list box while holding ‘Ctrl’ or ‘Shift”. This allows grouped manipulation, which can be useful for designs with a large number of components. Note the first-selected bundle is the reference in grouped manipulation along the local coordinate.

##### 4.2.2. Define the connectivity between components (TM A5 1:16~end)

The second step of Assembly is to define the connectivity between components. Every connection made in the MagicDNA software will consist of two scaffold strand connections, whose length can be adjusted and does not need to be equal. The dots depicted on the surface of the bundle components denote positions where connections can be made between bundles. The dots on the cylinder ends denote a scaffold crossover between paired helices that can be broken to make a connection to another component. Similarly, the dots on the sides of the bundles denote locations where the helical position of the scaffold is oriented at the surface, which is a location where a break could be introduced to make a double-scaffold connection to another component. These connections can be made automatically where the software searches for the nearest connection sites with the number and type (end or side) defined by user. Alternatively, the connections can be manually assigned between two selected components. The manual option pulls up a new window where users can click pairs of two nodes to form a connection regardless of separation distances.

##### 4.2.3. Specify the lengths of single-stranded scaffold connections (TM A6)

After defining the number and location of the connections, users can edit the lengths of individual scaffold connections using the “Edit ssDNA” function, which depicts the finer helical model allowing users to select connections and specify the single-stranded lengths. This is useful, for example, to tune joint mechanical properties[4].

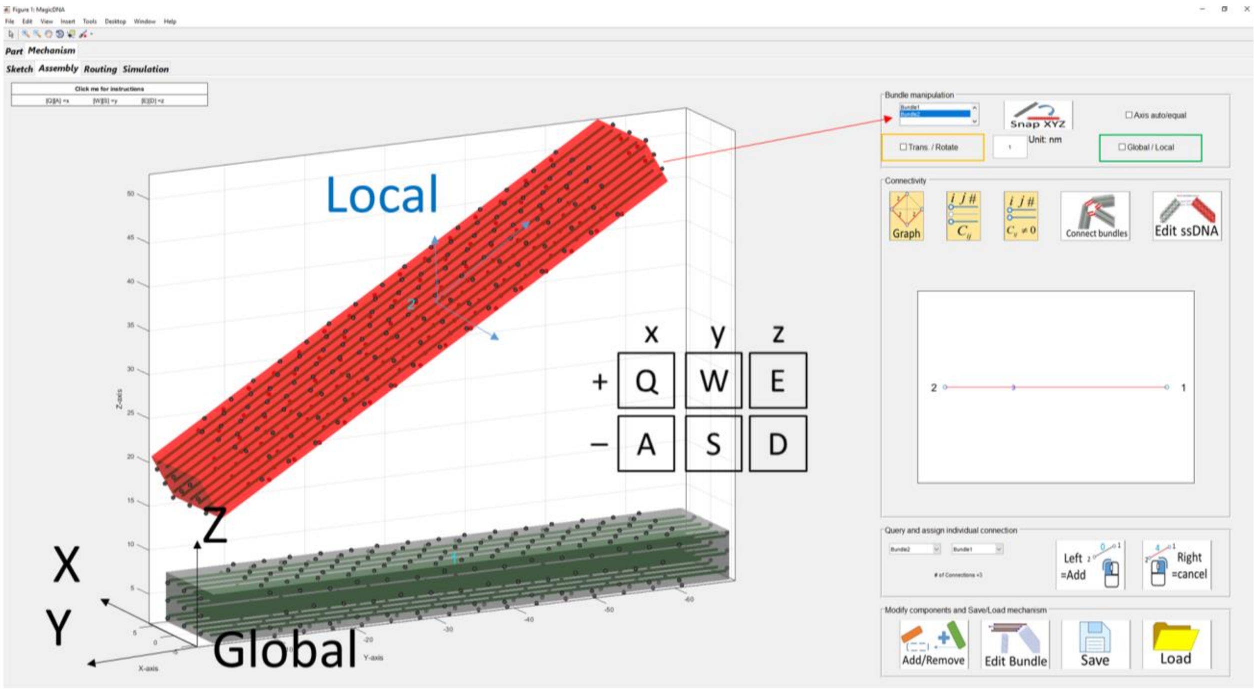

Figure 4. Bundle manipulation.

#### 4.3. Determine scaffold and staple routing

##### 4.3.1. The scaffold routing algorithm (TMs A7 and A8)

After defining the Geometry and Assembly properties, the routing algorithms embedded in MagicDNA can automatically find the scaffold and staple routings and visualize them in 3D space (Fig. 3C). The scaffold routing algorithm is based on the spanning tree algorithm to guide the integration of cycles into one loop and further expand to multi-scaffold designs by creating a spanning forest. To steer the spanning forest into multiple trees, users can go back to the bundle editor GUI to specify three features: 1) adding internal crossovers at designated positions, 2) ignoring potential crossovers between pairs of cylinders, and 3) ignoring potential crossovers within a specific portion of a bundle (See TM B5, B8 and Sec. 5.3). Alternatively, multiple (*K*) scaffold routings can also be achieved by applying (*K*-1) double-scaffold crossovers to one single scaffold with length constraint of split scaffold cycles (Fig. 5A). The triangular plate example (Section 5.3) describes the details of multi-scaffold designs.

##### 4.3.2. The staple routing algorithm and the overhang design tool (TMs A9 and A10)

In addition to the typical staple routing algorithm, the overhang design tool utilizes the advantage of 3D model to allow users to specify the locations to extend staples to form overhangs. The user can select overhang locations, 5' or 3' ends, and lengths of the overhangs in 3D model (TM B4). This provides extra inputs for the staple overhang algorithm to complete the staple design.

##### 4.3.3. Visualize the routings and assign the sequences to obtain the staple sequence list (TM A11)

Last, both scaffold and staple routings can be visualized in the sequence and diagram GUI with 2D (path diagram) and 3D (helical or straight-line) representations. By assigning the sequence(s) of the scaffold(s), the staple sequence list is generated for simulation and fabrication. This GUI also helps with the modification in caDNAno and users can upload designs that have been modified in caDNAno back into MagicDNA for further use.

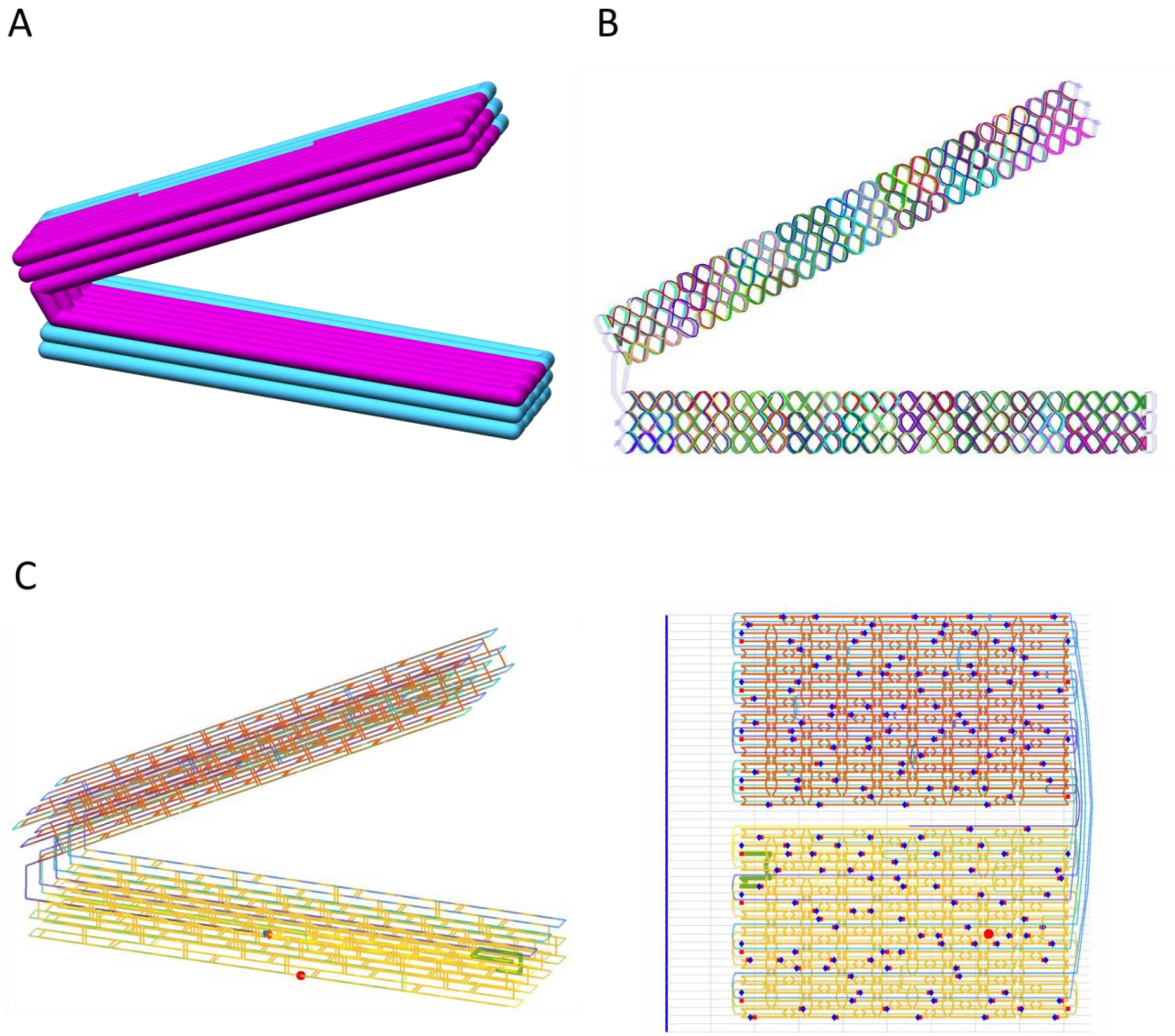

*Figure 5. (A) Scaffold routing with two scaffolds. (B) Staple routing. (C) The routing result in the GUI.*

#### 4.4. Interfacing with simulation

##### 4.4.1. Prepare the oxDNA topology and initial configuration for simulation (TM A12)

The last step is to export the oxDNA topology and configuration files for simulation. The topology file contains the sequence and connectivity between bases which can be obtained from the routings. The configuration file indicates the 3D positions and orientations of each base which are the linear composition of the chicken-wire (thin cylinder model) and helical representations. MagicDNA also allows users to decide the initial configuration by fine tuning the 3D manipulation of each component. It is highly recommended that users save the entire design profile (including geometry, assembly, and routing) with the simulation into one folder for managing multiple iterations and other designs.

###### 4.4.2. Visualize and analyze the trajectory file of the simulation (TMs A13, C1~C4)

After the simulation is completed, we provide visualization tool and basic analysis like root-mean-square deviation (RMSD) and root-mean-squared fluctuations (RMSF). Users can direct visualize one or multiple configurations in the trajectory in MATLAB or export a .BILD file to load into other software like UCSF Chimera for higher quality visualization.

### 5. Design examples

We have mentioned the general design process to design a simple hinge device (Sec. 4 and TMs. A1~A13). We encourage new users to practice one or two simple designs a few (2-3) components first (See TM. B1~B2 with step-by-step instructions.) to get familiar with the interface and the instructions of each tab. In fact, one of MagicDNA's features is the wide range of design capacity. Hence, here we present a few examples with medium-level complexity (TMs. B3, B4, B6, and B7) to demonstrate the versatile design process. Moreover, with the multi-scaffold algorithm, extremely complex structures can be designed (TMs. B8 and B5), simulated, and experimentally fabricated.

#### 5.1 Interlocked ring (TM B6)

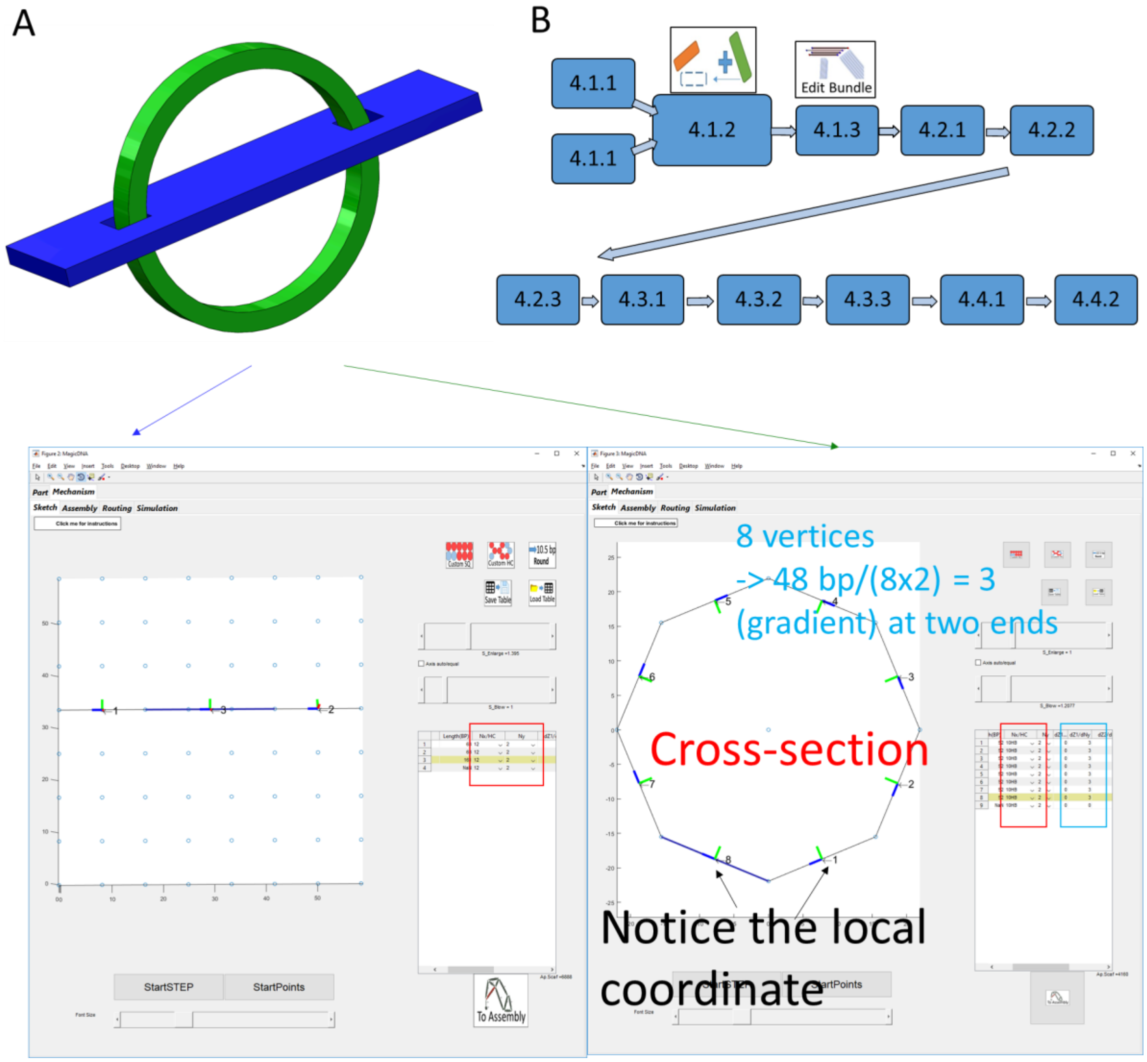

*Figure 6. Interlock ring (A) The target structure. (B) The overview of schematic design processes as TM (GUI) indices.*

The first structure is the interlocked ring shape shown in Fig. 6. It is clear that we can separate the structure into two portions: the plate (blue) and the ring (green).

- Design strategy and the top-down approach for geometry (see Sec. 4.4.1).

The first step is to mimic the rigid model with the cylinder model, especially paying attention to the number of bundles and cross-sections. For the plate, because there are two holes in the plates, we need to use three lines to form the three bundles/sections and then edit the middle bundle by shrinking a few to create two holes. The sketch is created with three serial lines using the line model sketch tool with Cartesian coordinates. One the other hand, the ring can be approximated by a circular pattern of tilted bundles with calculated gradients. The line model sketch tool with cylindrical coordinate is suitable for this task. Now we plan to convert two sets of lines to two sets of bundles (top-down) and later merge two sets of bundles (bottom-up) in the Assembly GUI. Remember, the only thing the user cannot change in Assembly and bundle editor GUIs is the cross-section property of each bundle. Hence, depend on how clearly the user is aware of the desired dimensions, it may take a few rounds between converting, merging, manipulation, and assigning the connectivity matrix. A small tip for estimating the length of the scaffold is to use keys: [k] and [l] in Assembly GUI and the rough double-stranded size will print out on the command line. Note this estimated length does not consider ssDNA connections and scaffold loops since the user may assign different values in the following steps, so it is only an approximate estimate.

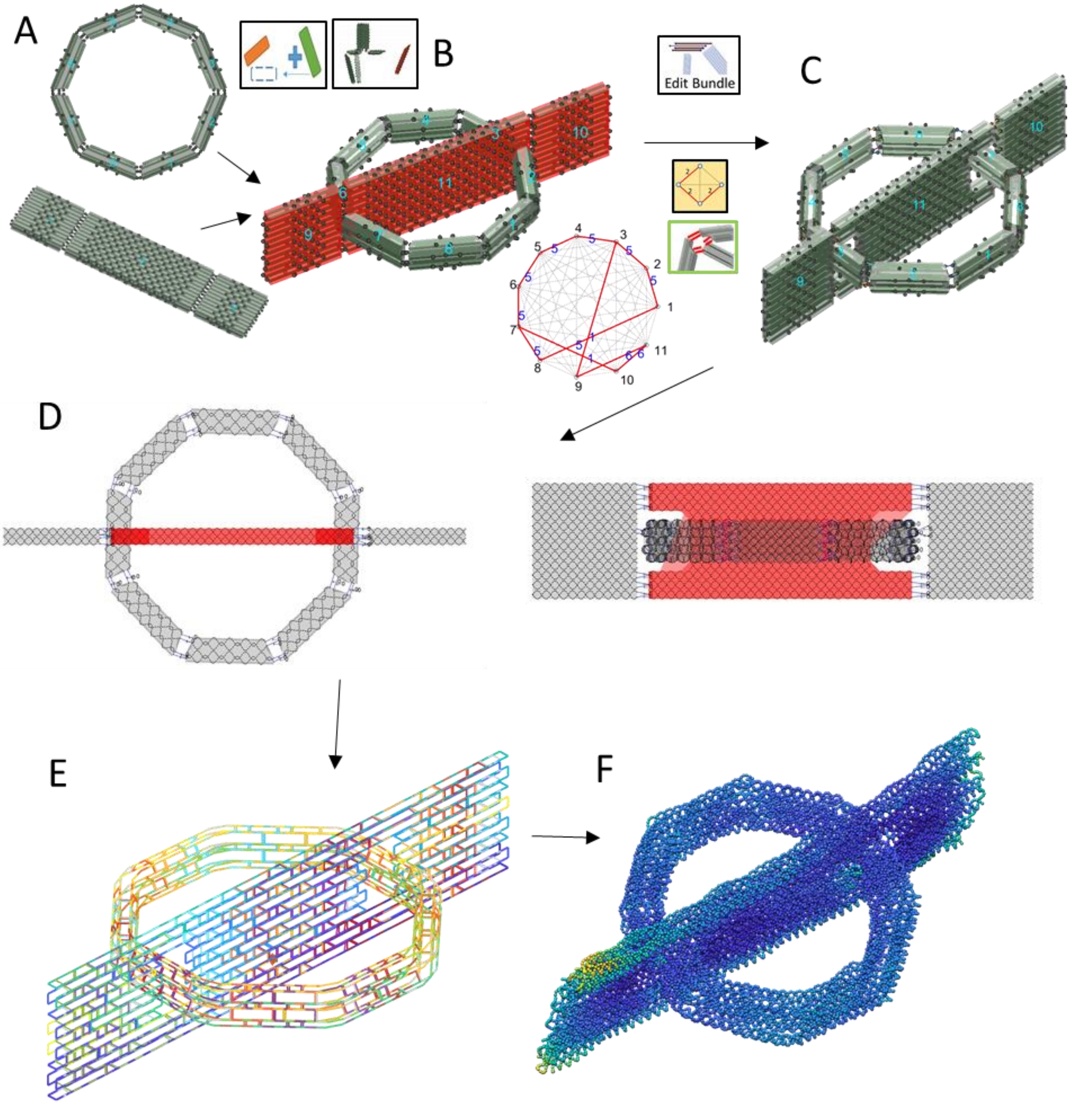

*Figure 7. Design process of the interlock structure*

- Bottom-up to merge the geometry (see Sec. 4.1.2 and 4.1.3)

After using the top-down approach to convert lines to bundles (Fig. 7A), we insert two sets of bundles and manipulate them in group to approximated position (Fig. 7B). Then, we want to modify the lengths of some cylinders to create two holes in the plate while the ring is still in the space with transparent appearance as the reference for the holes in the edit bundle GUI. Again, the user should remember to export all the bundles to update the assembly because the cylinder model has been changed.

- Complete the design (Sec. 4.2, 4.3, 4.4)

The rest of following processes are similar to the general design process, including assign connectivity matrix to connect bundles with double-scaffold crossovers (Fig. 7C), single-stranded scaffold length GUI (Fig. 7D), routing algorithm to find the scaffold and staple routing (Fig. 7E), and coarse-grained oxDNA simulation (Fig. 7F).

#### 5.2 Tower (see TM B7)

- Use a .STEP file and the bundle edit GUI for geometry (Sec 4.1.1 and 4.1.3)

The second structure is the wireframe tower structure formed by either 4HB or 6HB bundles, shown in Fig. 8. Because it contains 12 edges/bundles in 3D space axially symmetric to the axis, we chose to use CAD software, such as Solidworks or FreeCAD, to generate the line model (.STEP files) and later converted to bundles as the top-down approach (Fig. 8 C and D). It is still possible to change the lengths of the cylinders in a bundle by using the edit bundle GUI. To maintain the vertex angle at the top, the lengths of the cylinders have been adjusted for the 3D four-way junction. The geometry of this structure (the cylinder model) is basically completed in Fig. 8F.

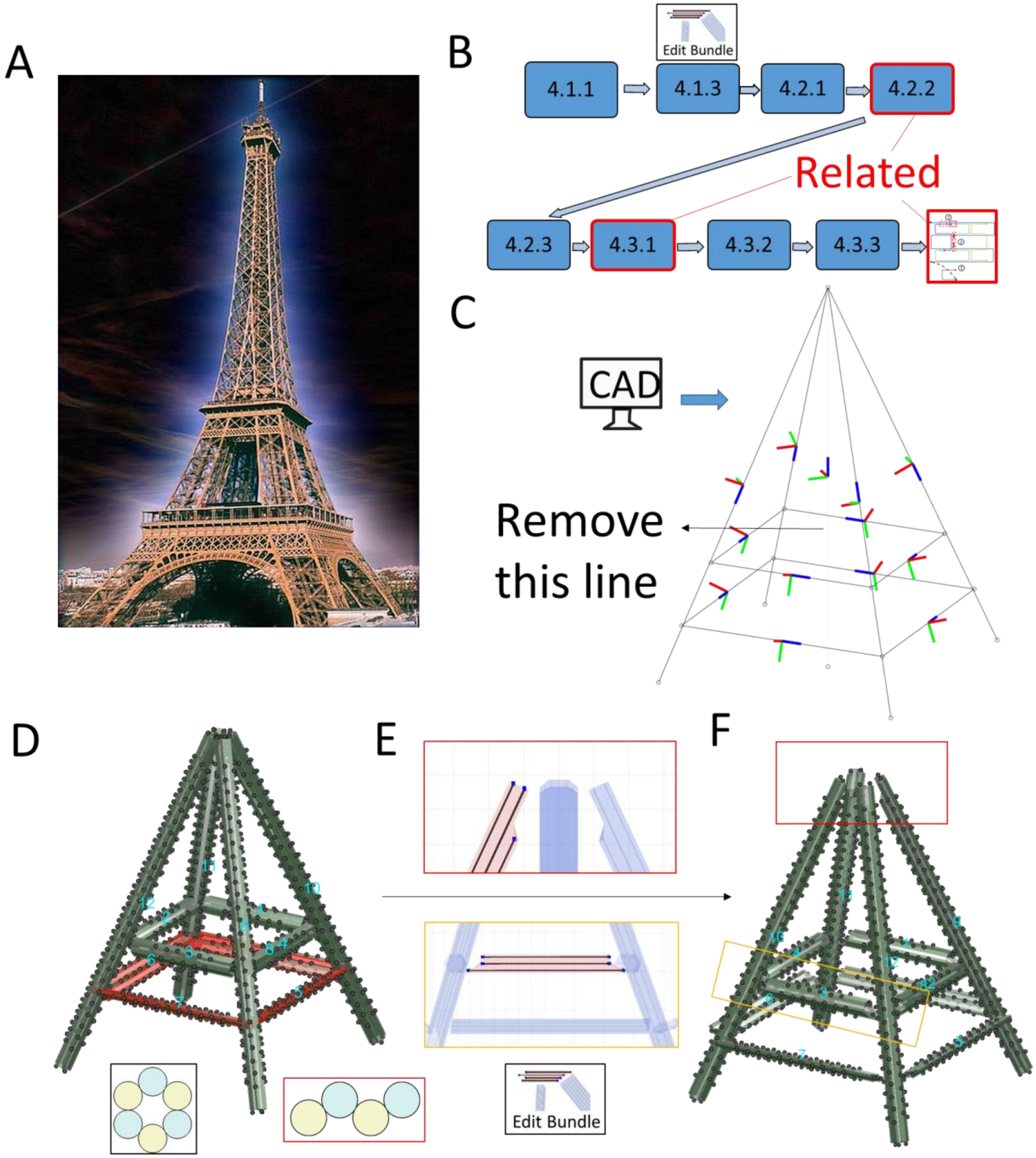

*Figure 8. Geometry of the tower structure*

- Consider the connectivity with scaffold algorithm and modification in caDNAno (Sec. 4.2 and 4.3)

After the geometry is created from the line model and fine-tuned in the bundle editor GUI, the next steps are assembly and scaffold/staple routing. Here we want to consider these two steps together, especially for the four-way junction. The rest of the connectivity between components are end-to-side double-scaffold crossover connections (DSCC) and intuitively related to the cross-sections, i.e. 4HB -> 2 DSCCs and 6HB -> 3DSCCs. On the other hand, for each bundle at the four-way junction, we extended four of the cylinders (blue in Fig. 9B and 9D) which corresponds to two connection nodes (pairs of cylinders), while leaving the other two cylinders shorter (orange in Fig. 9B and 9E) which corresponds to one connection node. For the top portion (4 extended cylinders for each bundle), even numbers of nodes are easy to count how many

DSCC are needed (Fig. 9C). Later, we can assign longer ssDNA lengths on the longer connection in the single-stranded scaffold GUI (Fig. 9D). As for the bottom section, the algorithm does not allow multiple connections from a single node (i.e. one node can only connect to one other node.). Hence, we only assign the connectivity of components 9 and 10 (Fig. 9E). According to the routing algorithm, the auto-generated initial scaffold routing will only have an external DSCC across bundles 9 and 10. The ends of bundles 11 and 12 remain as scaffold loops.

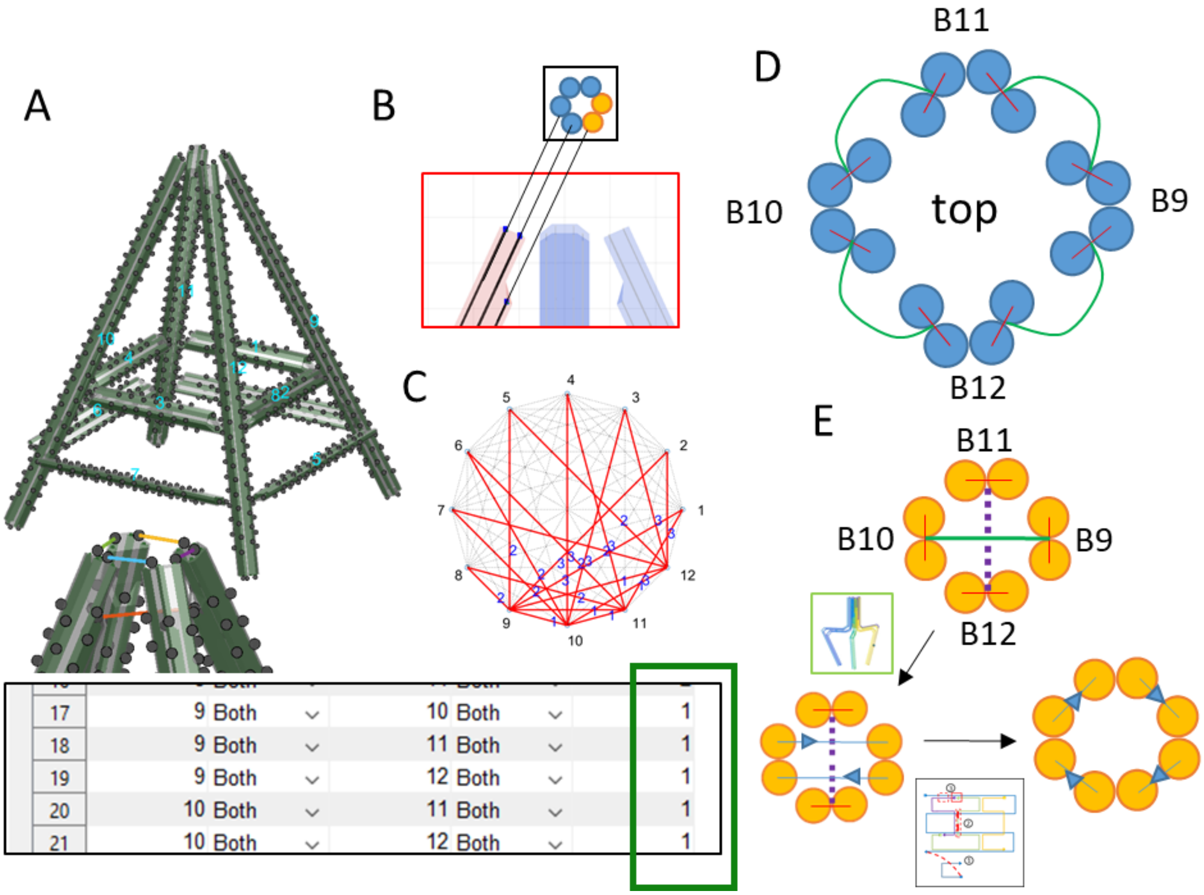

*Figure 9. Assembly and routing of the tower structure.*

- Modification in caDNAno (Sec 4.3.3)

The next step is to modify the routing at the four-way junctions by applying two crossovers between the DSCCs and the scaffold loops (Figs. 9E and 10). The exported .JSON file has been arranged so that all components are arranged diagonally in the cross-section view and two dummy cylinders are inserted between the components as spacing. In MagicDNA, the visualization in the sequence and diagram GUI is designed for assisting modification in caDNAno. Now the user may want to edit/add scaffold crossovers between components. Hence, by clicking on the 3D model, the location in the .JSON file is shown in graphical interface as well as the text (Fig. 10B). After adding two DSCCs as forced connections in caDNAno from one-scaffold routing, the number of total scaffolds is either 1 or 3 according to the double-crossover rule. Hence, adding or removing two scaffold internal crossovers may be necessary if users want to use single scaffold to fold the structure. Then, in the staple tab, use the button to upload the modified .JSON file. Both scaffold and staple routing will be updated and can be visualized in 3D to make sure the modification is reasonable.

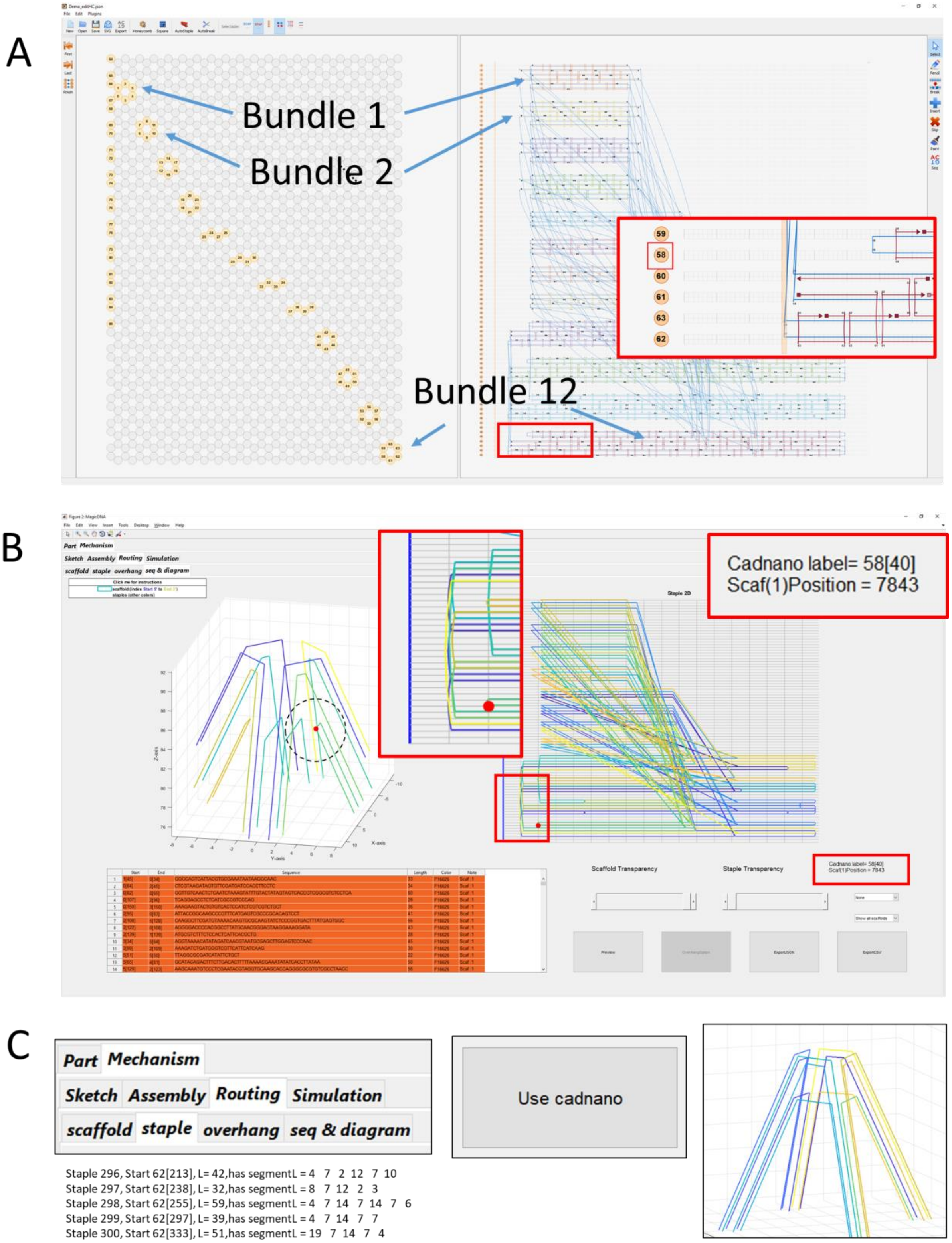

*Figure 10. Use caDNAno to update the routings.*

#### 5.3 A triangle plate (see TM B8 Multi-scaffold)

- Geometry and Assembly (Sec 4.1 and 4.2)

In this example, the topic will focus on how to design multi-scaffold structures in MagicDNA. Let's take a simple triangle plate made by 4×4 square-lattice bundles as illustrated in Fig. 11. For the routing algorithm, Figs. S70~S72 and S81 in the supplement material have more descriptions for multi-scaffold. In general, the first approach is to allow users to specify more inputs to affect the routing algorithm to obtain the desired routing, which may need some level of understanding the algorithm. The second approach is to search and apply *K*-1 internal crossovers to the original single scaffold to obtain *K* scaffold routings while considering the length constraints of the spilt scaffolds (Remember the double-crossover rule for total number of cycles.).

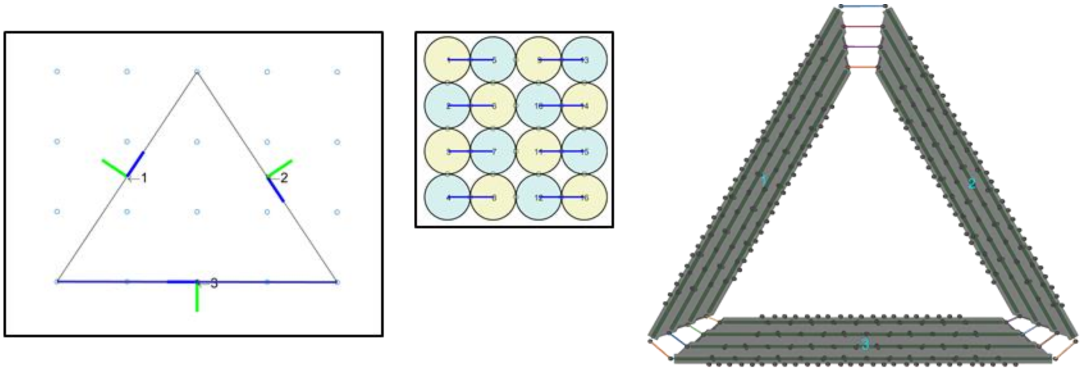

| | Edge | Length(BP) | Nx/HC | Ny | dZ1/dNx | dZ1/dNy | dZ2/dNx | dZ2/dNy |
|---|---|---|---|---|---|---|---|---|
| 1 | 1 | 159 | 4 | 4 | 0 | -8 | 0 | 8 |
| 2 | 2 | 159 | 4 | 4 | 0 | -8 | 0 | 8 |
| 3 | 3 | 159 | 4 | 4 | 0 | -8 | 0 | 8 |
| 4 | NaN | NaN | 4 | 4 | 0 | 0 | 0 | 0 |

*Figure 11. Geometry and assembly of the triangle plate.*

- The 2nd approach for multi-scaffold (Sec 4.3.1)

From the viewpoint of user operations, the second approach is relatively easy since users only need to assign the number of scaffolds, *K*, and a tolerance percentage for the length constraints (Fig. 12). The initial single scaffold cycle can be either directly from the MagicDNA algorithm or the modified routing from caDNAno. One thing to notice is that this process has to start with one single scaffold routing.

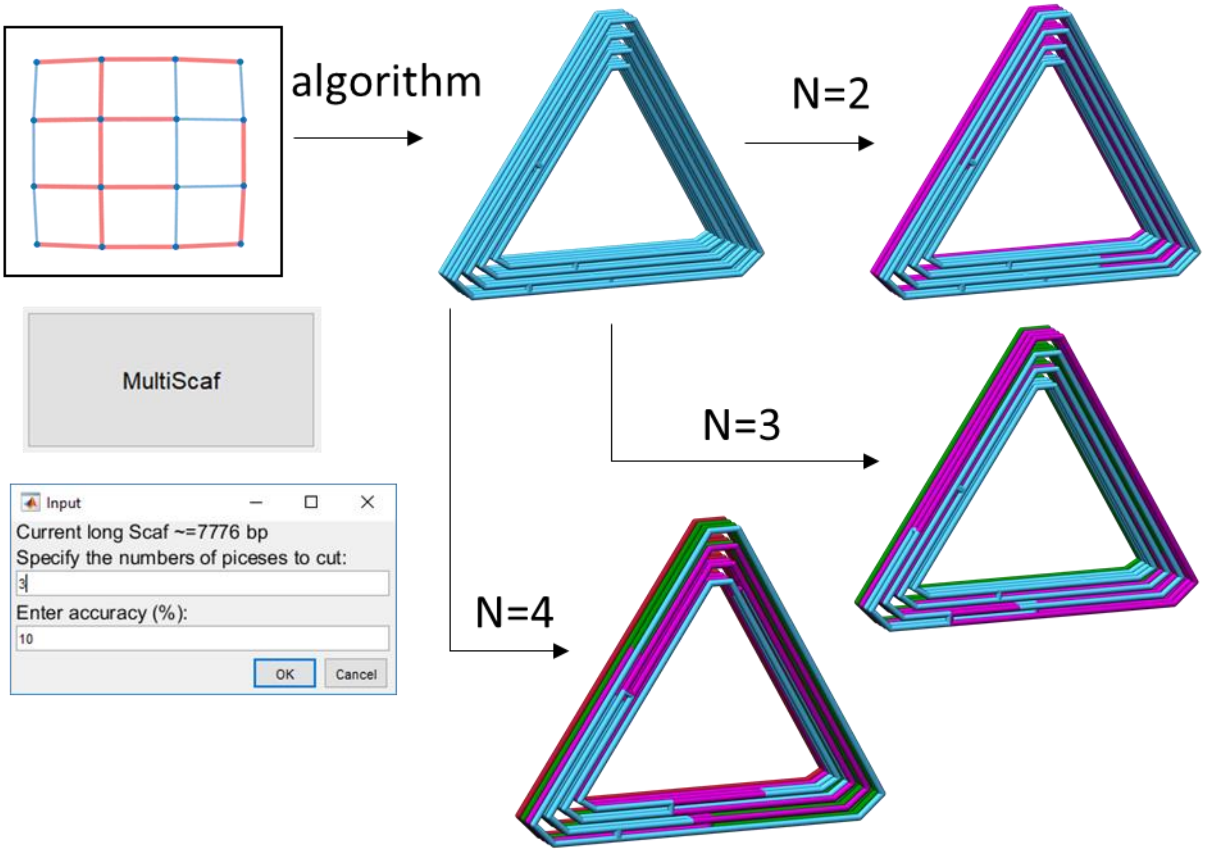

*Figure 12. Multi-scaffold using 2nd approach*

- The 1st approach for multi-scaffold designs, only splitting between cylinders (Sec. 4.1.3 and 4.3.1)

As for the first approach, specifying more inputs to the algorithm, the bundle editor GUI provides interfaces to: 1) add internal crossovers at designated positions, 2) ignore potential crossovers between pairs of cylinders, and 3) ignore potential crossovers within a specific portion of a bundle. Fig. 13 shows the cross-section view of the cylinders where the red dots mean that the potential crossovers between cylinders will be ignored in the algorithm later. Hence, after applying the external DSCCs to the cycles as paired cylinders, this creates some disconnections between the cycles (Fig. 13 middle) in the graph compared to Fig. 12. (It can be observed that the total number of cycles/nodes (=16) after applying external DSCCs remains the same as Fig. 12.) The function in MATLAB Graph and Network Algorithms Toolbox returns multiple spanning trees in a forest instead of a single tree. Later, each tree guides the integration of corresponding cycles into one scaffold routing by searching one internal crossover for each edge (between two cycles) on the spanning tree (bold red lines).

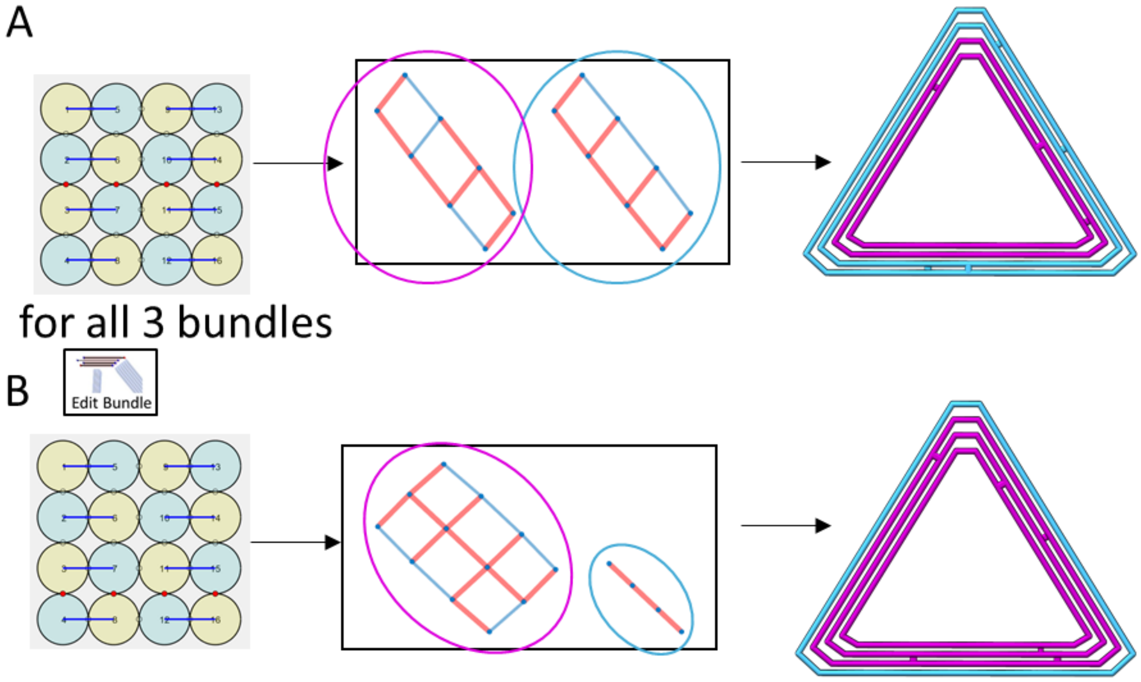

*Figure 13. Multi-scaffold using 1st approach ignoring potential crossovers between cylinders*

- Use the 1st approach for multi-scaffold, split along the cylinder direction. (Sec. 4.1.3 and 4.3.1)

In addition to <u>(2) ignore potential crossovers between pairs of cylinders</u>, the other two inputs for multi-scaffold are also assigned through the edit bundle GUI (Fig. 14). The GUI suggests the internal crossovers based on either the selected cylinder (top) or the pairs of cylinders (bottom). To define the interface of multiple scaffolds, we suggest to assign internal crossovers through pairs of cylinders because it has to have 8 concentrated crossovers on all cylinders in the 4×4 cross-section to form the seam. To prevent the examining of neighboring cycles in the graph across the interface, it also has to assign a specific portion of a bundle (green rectangles) where the potential crossovers for integrating cycles are ignored. Fig. 14 B and C show how to define two kinds of interfaces between scaffolds: one has almost flat interfaces and the other can better interlock the scaffolds.

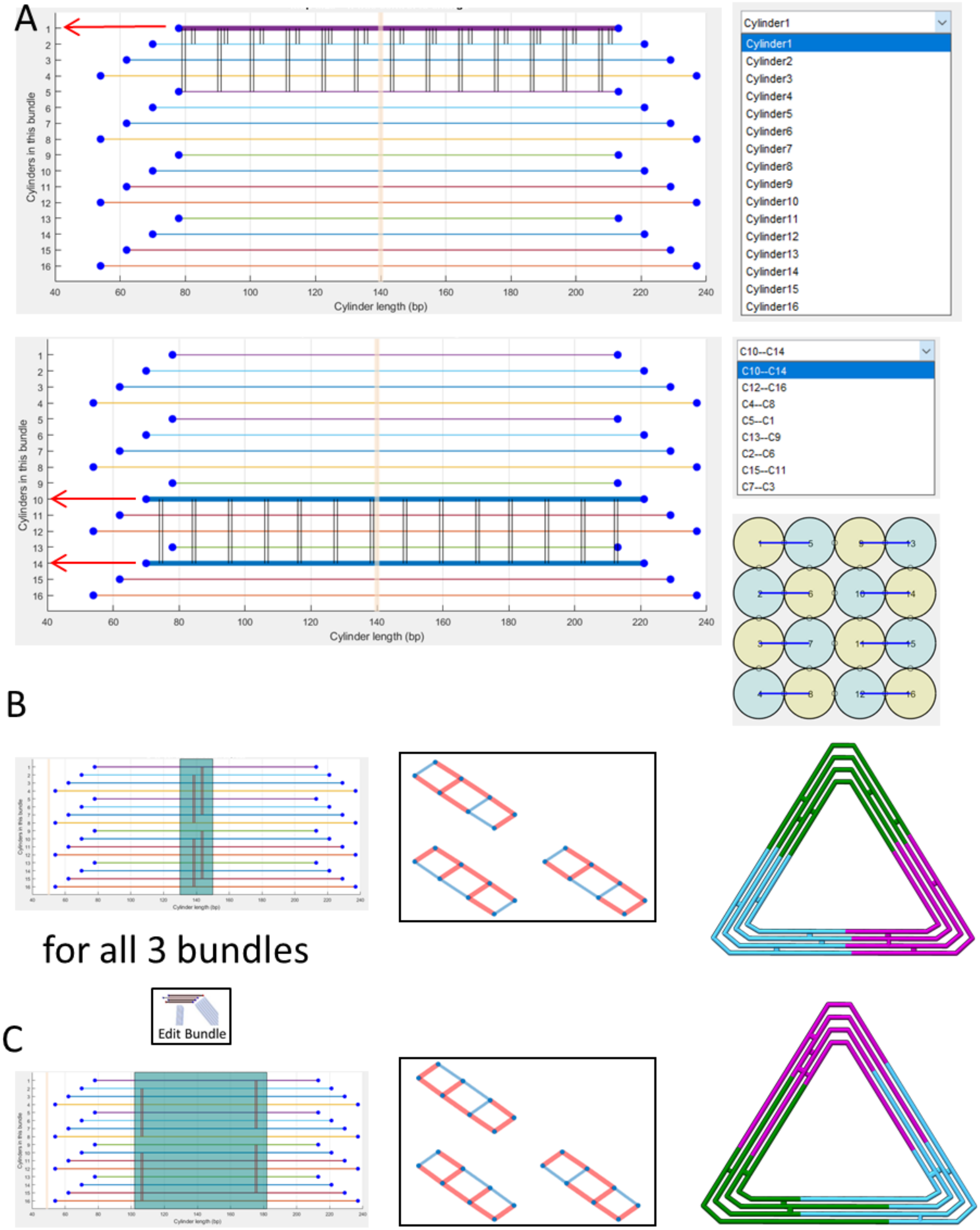

*Figure 14. Multi-scaffold using 1st approach that the user defines internal crossovers and assign in a portion of a bundle which ignores the potential crossovers. (A) The snapshots in the edit bundle GUI to show the scaffold crossovers from a cylinder (Top) or between a pair of cylinders (Bottom). (B)(C)The corresponding 3-scaffold routings by defining the interfaces.*

### Reference:

(1) Seeman, N. C. Nucleic Acid Junctions and Lattices. *J. Theor. Biol.* **1982**, *99* (2), 237–247. https://doi.org/10.1016/0022-5193(82)90002-9.

(2) Seeman, N. C.; Sleiman, H. F. DNA Nanotechnology. *Nat. Rev. Mater.* **2017**, *3* (1), 1–23. https://doi.org/10.1038/natrevmats.2017.68.

(3) Gerling, T.; Wagenbauer, K. F.; Neuner, A. M.; Dietz, H. Dynamic DNA Devices and Assemblies Formed by Shape-Complementary, Non–Base Pairing 3D Components. *Science* **2015**, *347* (6229), 1446–1452. https://doi.org/10.1126/science.aaa5372.

(4) Marras, A. E.; Zhou, L.; Su, H.-J.; Castro, C. E. Programmable Motion of DNA Origami Mechanisms. *Proc. Natl. Acad. Sci.* **2015**, *112* (3), 713–718. https://doi.org/10.1073/pnas.1408869112.

(5) DeLuca, M.; Shi, Z.; E. Castro, C.; Arya, G. Dynamic DNA Nanotechnology: Toward Functional Nanoscale Devices. *Nanoscale Horiz.* **2019**. https://doi.org/10.1039/C9NH00529C.

(6) Linko, V.; Ora, A.; Kostiainen, M. A. DNA Nanostructures as Smart Drug-Delivery Vehicles and Molecular Devices. *Trends Biotechnol.* **2015**, *33* (10), 586–594. https://doi.org/10.1016/j.tibtech.2015.08.001.

(7) Ke, Y.; Castro, C.; Choi, J. H. Structural DNA Nanotechnology: Artificial Nanostructures for Biomedical Research. *Annu. Rev. Biomed. Eng.* **2018**, *20* (1), null. https://doi.org/10.1146/annurev-bioeng-062117-120904.

(8) Rothemund, P. W. K. Folding DNA to Create Nanoscale Shapes and Patterns. *Nature* **2006**, *440* (7082), 297–302. https://doi.org/10.1038/nature04586.

(9) Douglas, S. M.; Dietz, H.; Liedl, T.; Högberg, B.; Graf, F.; Shih, W. M. Self-Assembly of DNA into Nanoscale Three-Dimensional Shapes. *Nature* **2009**, *459* (7245), 414–418. https://doi.org/10.1038/nature08016.

(10) Wagenbauer, K. F.; Engelhardt, F. A. S.; Stahl, E.; Hechtl, V. K.; Stömmer, P.; Seebacher, F.; Meregalli, L.; Ketterer, P.; Gerling, T.; Dietz, H. How We Make DNA Origami. *ChemBioChem* **2017**, *18* (19), 1873–1885. https://doi.org/10.1002/cbic.201700377.

(11) Castro, C. E.; Kilchherr, F.; Kim, D.-N.; Shiao, E. L.; Wauer, T.; Wortmann, P.; Bathe, M.; Dietz, H. A Primer to Scaffolded DNA Origami. *Nat. Methods* **2011**, *8* (3), 221–229. https://doi.org/10.1038/nmeth.1570.

(12) Halley, P. D.; Patton, R. A.; Chowdhury, A.; Byrd, J. C.; Castro, C. E. Low-Cost, Simple, and Scalable Self-Assembly of DNA Origami Nanostructures. *Nano Res.* **2019**, *12* (5), 1207–1215. https://doi.org/10.1007/s12274-019-2384-x.

(13) Douglas, S. M.; Marblestone, A. H.; Teerapittayanon, S.; Vazquez, A.; Church, G. M.; Shih, W. M. Rapid Prototyping of 3D DNA-Origami Shapes with CaDNAno. *Nucleic Acids Res.* **2009**, *37* (15), 5001–5006. https://doi.org/10.1093/nar/gkp436.

(14) Benson, E.; Mohammed, A.; Gardell, J.; Masich, S.; Czeizler, E.; Orponen, P.; Högberg, B. DNA Rendering of Polyhedral Meshes at the Nanoscale. *Nature* **2015**, *523* (7561), 441–444. https://doi.org/10.1038/nature14586.

(15) Veneziano, R.; Ratanalert, S.; Zhang, K.; Zhang, F.; Yan, H.; Chiu, W.; Bathe, M. Designer Nanoscale DNA Assemblies Programmed from the Top Down. *Science* **2016**, *352* (6293), 1534–1534. https://doi.org/10.1126/science.aaf4388.

(16) Jun, H.; Shepherd, T. R.; Zhang, K.; Bricker, W. P.; Li, S.; Chiu, W.; Bathe, M. Automated Sequence Design of 3D Polyhedral Wireframe DNA Origami with Honeycomb Edges. *ACS Nano* **2019**. https://doi.org/10.1021/acsnano.8b08671.

(17) Matthies, M.; Agarwal, N. P.; Schmidt, T. L. Design and Synthesis of Triangulated DNA Origami Trusses. *Nano Lett.* **2016**, *16* (3), 2108–2113. https://doi.org/10.1021/acs.nanolett.6b00381.

(18) Ouldridge, T. E.; Louis, A. A.; Doye, J. P. K. Structural, Mechanical, and Thermodynamic Properties of a Coarse-Grained DNA Model. *J. Chem. Phys.* **2011**, *134* (8), 085101. https://doi.org/10.1063/1.3552946.

(19) Doye, J. P. K.; Ouldridge, T. E.; Louis, A. A.; Romano, F.; Šulc, P.; Matek, C.; Snodin, B. E. K.; Rovigatti, L.; Schreck, J. S.; Harrison, R. M.; et al. Coarse-Graining DNA for Simulations of DNA Nanotechnology. *Phys. Chem. Chem. Phys.* **2013**, *15* (47), 20395–20414. https://doi.org/10.1039/C3CP53545B.

(20) Snodin, B. E.; Randisi, F.; Mosayebi, M.; Šulc, P.; Schreck, J. S.; Romano, F.; Ouldridge, T. E.; Tsukanov, R.; Nir, E.; Louis, A. A. Introducing Improved Structural Properties and Salt Dependence into a Coarse-Grained Model of DNA. *J. Chem. Phys.* **2015**, *142* (23), 06B613_1.

(21) TacoxDNA: A user-friendly web server for simulations of complex DNA structures, from single strands to origami - Suma - 2019 - Journal of Computational Chemistry - Wiley Online Library https://onlinelibrary.wiley.com/doi/full/10.1002/jcc.26029 (accessed Jan 21, 2020).

(22) MrDNA: A multi-resolution model for predicting the structure and dynamics of nanoscale DNA objects | bioRxiv https://www.biorxiv.org/content/10.1101/865733v1 (accessed Jan 17, 2020).

(23) Engelhardt, F. A. S.; Praetorius, F.; Wachauf, C. H.; Brüggenthies, G.; Kohler, F.; Kick, B.; Kadletz, K. L.; Pham, P. N.; Behler, K. L.; Gerling, T.; et al. Custom-Size, Functional, and Durable DNA Origami with Design-Specific Scaffolds. *ACS Nano* **2019**, *13* (5), 5015–5027. https://doi.org/10.1021/acsnano.9b01025.

(24) Tikhomirov, G.; Petersen, P.; Qian, L. Fractal Assembly of Micrometre-Scale DNA Origami Arrays with Arbitrary Patterns. *Nature* **2017**, *552* (7683), 67–71. https://doi.org/10.1038/nature24655.

(25) Wagenbauer, K. F.; Sigl, C.; Dietz, H. Gigadalton-Scale Shape-Programmable DNA Assemblies. *Nature* **2017**, *552* (7683), 78. https://doi.org/10.1038/nature24651.
